# The effects of inversion polymorphisms on patterns of neutral genetic diversity

**DOI:** 10.1101/2023.02.23.529778

**Authors:** Brian Charlesworth

**Affiliations:** Institute of Ecology and Evolution, School of Biological Sciences, University of Edinburgh, Edinburgh EH9 3FL, United Kingdom

**Keywords:** Inversion polymorphisms, genetic diversity, sequence divergence, recombination rate, population subdivision

## Abstract

The strong reduction in the frequency of recombination in heterozygotes for an inversion and a standard gene arrangement causes the arrangements to become partially isolated genetically, resulting in sequence divergence between them and changes in the levels of neutral variability at nucleotide sites within each arrangement class. Previous theoretical studies on the effects of inversions on neutral variability have either assumed that the population is panmictic or that it is divided into two populations subject to divergent selection. Here, the theory is extended to a model of an arbitrary number of demes connected by migration, using a finite island model with the inversion present at the same frequency in all demes. Recursion relations for mean pairwise coalescent times are used to obtain simple approximate expressions for diversity and divergence statistics for an inversion polymorphism at equilibrium under recombination and drift, and for the approach to equilibrium following the sweep of an inversion to a stable intermediate frequency. The effects of an inversion polymorphism on patterns of linkage disequilibrium are also examined. The reduction in effective recombination rate caused by population subdivision can have significant effects on these statistics. The theoretical results are discussed in relation to population genomic data on inversion polymorphisms, with an emphasis on *Drosophila melanogaster*. Methods are proposed for testing whether or not inversions are close to recombination-drift equilibrium, and for estimating the rate of recombinational exchange in heterozygotes for inversions; difficulties involved in estimating the ages of inversions are also discussed.

## Introduction

Naturally occurring genetic factors that massively reduce the rate of crossing over in *Drosophila melanogaster* when heterozygous were discovered by A.H. Sturtevant over 100 years ago (Sturtevant 1917), who later showed them to be inversions of segments of chromosomes (Sturtevant 1926). Inversion polymorphisms were long regarded as a curiosity of species of *Drosophila* and other “higher” Diptera, where they can readily be detected cytologically using the polytene chromosomes of the larval salivary glands. The largest body of information concerning the properties of inversions has been accumulated in studies of numerous *Drosophila* species (Krimbas and Powell 1992a; Kapun and Flatt 2019). The recent application of genome sequencing technology to the natural populations of many organisms, including humans, has revealed that inversions are much more abundant than was previously thought. They are sometimes associated with striking phenotypic polymorphisms, such as the social chromosomes of ants and the behavioral and color polymorphisms of the ruff and white-throated sparrow (Wellenreuther and Bernatchez 2018; Villoutreix et al. 2021). This has led a surge in interest in the evolutionary significance of inversion polymorphisms; however, the *Drosophila* studies suggest that the vast majority of polymorphic inversions have little or no effects on visible phenotypes, although effects on quantitative traits such as body size and some fitness components have been detected (Krimbas and Powell 1992a; Kapun and Flatt 2019).

It is evident that the genetic isolation of different gene arrangements due to the suppression of crossing over in heterokaryotypes must play a major role in the processes that lead to the evolution of inversion polymorphisms, although the nature of these processe is still an open research question, which may well have multiple answers (Krimbas and Powell 1992b; Wellenreuther and Bernatchez 2018; Kapun and Flatt 2019; Villoutreix et al. 2021). The present paper is concerned with the consequences of the suppression of crossing over in heterozygotes for an inversion and a standard arrangement (heterokaryotypes) for the levels of diversity within and between the two arrangements at neutral nucleotide sites contained inside or close to the genomic region covered by the inversion. A critical parameter in determining these genomic features is the rate of recombinational exchange between arrangements in heterokaryotypes, the “gene flux” of Navarro et el. (1997). Evidence about this rate has been provided by genetic studies of recombination in heterokaryotypes. An important mechanism by which crossing over is suppressed or reduced in frequency during *Drosophila* female meiosis in heterozygotes for a paracentric inversion and the standard arrangement was elucidated by Sturtevant and Beadle (1936), whose genetic data suggested that single crossovers produce dicentric and acentric chromosomes that fail to be included in the egg nucleus.

Recent studies of heterozygous *Drosophila* inversions suggest a near-total suppression of crossing over within the regions covered by a heterozygous inversion and for a substantial distance outside it (Li et al. 2022; Koury 2023). Recombination suppression is, however, often incomplete, with gene conversion and/or double crossing over causing exchanges of alleles between arrangements in heterokaryotypes (Sturtevant and Beadle 1936; Chovnick 1973; Krimbas and Powell 1992b; Navarro et al. 1997; Crown et al. 2018; Korunes and Noor 2019; Li et al. 2022). The conversion of double strand breaks into non-crossover associated gene conversion events apparently plays a major role in the suppression of crossing over, as well as allowing exchange to occur via gene conversion (Gong et al. 2005; Crown et al. 2018; Li et al. 2022). But the low rate of occurrence of such gene conversion events and double crossovers (a consensus value for *Drosophila* is about 10^−5^ per basepair in female meiosis: Korunes and Noor [2019]) means that different arrangements are likely to become substantially genetically differentiated from each other, as a result of the interplay between mutation, genetic drift and recombination (Ishii and Charlesworth 1977; Navarro et al. 1997, 2000; Andolfatto et al. 2001; Guerrero et al. 2012).

Several lines of evidence, including studies of changes in inversion frequencies in experimental populations, temporal fluctuations in inversion frequencies and clinal patterns of inversion frequencies, as well as direct measurements of fitness components, show that many inversions are maintained at intermediate frequencies by natural selection (Krimbas and Powell 1992b; Mérot et al. 2018; Kapun and Flatt 2019). An initial complete loss of variability within haplotypes carrying a newly arisen, selectively favored inversion (the hitchhiking effect of Maynard Smith and Haigh [1974]), will be eroded by the occurrence of new mutations that spread as a result of genetic drift and recombinational exchange with the standard arrangement. Once an inversion subject to balancing selection has established itself at an intermediate frequency within a population, there will be a gradual approach to mutation-drift-recombination equilibrium with respect to neutral or nearly neutral variants (Navarro et al. 2000). It is therefore important to consider the effects on variability of both the approach to equilibrium and the equilibrium situation.

The equilibrium properties of an inversion polymorphism with respect to neutral variability are similar to those for neutral loci linked to a single locus subject to balancing selection, first modeled by Ohta and Kimura (1970) using diffusion equations and by Strobeck (1983) using identity probabilities. Over the following two decades, subsequent modeling work using the structured coalescent process shed further light on patterns of neutral variability at sites linked to loci under balancing selection or divergent local selection, e.g., Kaplan et al. (1988), Hudson and Kaplan (1988), Hudson (1990), Takahata (1990), Charlesworth et al. (1997), Nordborg (1997) and Navarro et al. (2000), reviewed by Charlesworth et al. (2003). These studies showed that, when a balanced polymorphism has been maintained for a time that is much longer than the mean coalescent time for a neutral locus, we expect to see linkage disequilibrium (LD) between the target of selection and neutral sites closely linked to the target of selection, LD among these neutral sites, sequence divergence between haplotypes carrying the different alleles at the target of selection, and enhanced variability in the population as a whole around the target of selection.

These patterns are all reflections of the same process of divergence by drift and mutation among the partially isolated populations represented by the alleles at the target of selection, with a strong analogy with the outcome of mutation and drift in a geographically structured population (Hudson 1990; Charlesworth et al. 2003). Inversions are only special because of the strong suppression of recombination in inversion heterozygotes, which extendes to regions outside the inversion that are close to the inversion breakpoints (Navarro et al. 1997; Crown et al. 2018; Korunes and Noor 2019; Li et al. 2022; Koury 2023). The patterns just described are thus much more likely to be detected with inversions than with single locus polymorphisms, due to the much larger region of the genome involved.

A basic finding of the early theoretical work was that significant associations between a balanced polymorphism and variants at a neutral locus at statistical equilibrium under drift and recombination are only likely to be observed if the rate of recombinational exchange in double heterozygotes for the loci involved is of the order of the reciprocal of the effective population size, *N*_*e*_, as is also the case for a pair of neutral loci (Hill and Robertson 1968; Ohta and Kimura 1969, 1971). Andolfatto et al. (2001) applied the theoretical results of Navarro et al. (2000) to the available data on *D. melanogaster* inversions, and concluded that large effects of inversions on patterns of variability are likely to be seen only for loci close to inversion breakpoints, where exchange is probably most strongly inhibited, and that the observed patterns suggested that the inversions in question were of relatively recent origin compared with the age of the species.

Interest in the question of the effect of inversion polymorphisms on patterns of variability at sites that are either inside inversions or closely linked to inversion breakpoints has increased with the advent of whole genome sequencing, which has both greatly increased our ability to detect and characterize inversions, and allowed much more fine-scaled analyses of patterns of variability in genomic regions associated with inversions, e.g., Corbett-Detig and Hartl (2012), Cheng et al. (2012), Mérot et al. (2021) and Kapun et al. (2023). Recent population genomic analyses of several classic inversion polymorphisms in *Drosophila melanogaster* suggest the following patterns (Corbett-Detig and Hartl 2012; Kapun et al. 2023). (1). There is a modest increase in overall nucleotide site diversity in the regions covered by inversions and adjacent to them, reflecting a low level of sequence divergence between arrangements relative to the genome-wide average diversity. (2). If genetic differentiation between inverted and standard arrangements relative to within-arrangement diversity is measured by *F*_*ST*_-like statistics, a much stronger effect is seen, with mean between-arrangement *F*_*ST*_ values of the order of 0.1 to 0.2 in the interior of the inversion and in the regions adjacent to the inversions, with a sharp increase in regions close to the breakpoints. (3). For a low frequency inversion such as *In(3R)P* in the Zambian population (Kapun et al. 2023), there is a lower than genome-wide average diversity at sites within the inversion and a higher than average diversity within the standard arrangement. Much more information of this kind is likely to become available, and its interpretation requires a solid basis in population genetics theory.

A number of theoretical investigations on the effect of balanced polymorphisms on variability at linked sites have extended the older work described above, without, however, greatly modifying the basic conclusions, e.g., Innan and Nordborg (2003), Guerrero et al. (2012), Rousset et al. (2014) and Zeng et al. (2021). A limitation of most of the theoretical work on the effects of selectively maintained polymorphisms on neutral diversity is that it assumes a single, randomly mating population, with the exception of Charlesworth et al. (1997), Nordborg (1997), Guerrero et al. (2012) and Rousset et al. (2014), who considered the case of divergent and/or balancing selection in a pair of populations. Nordborg and Innan (2003) examined a more general model of population structure but relied on coalescent process simulations to generate predictions.

While it is often considered that population subdivision in organisms like *Drosophila* is likely to have only minor effects on genetic diversity, given the generally low levels of *F*_*ST*_ among populations (Singh and Rhomberg 1987; Schaeffer 2002; Lack et al. 2015, 2016), there is evidence from studies of allelism of recessive lethals that local deme sizes are in reality somewhat restricted in size, with limited migration among them (Wright et al. 1942; Mukai and Yamaguchi 1974; Ives and Band 1986); this conclusion has recently been confirmed by a resequencing study of a single US population over time (Lange et al. 2022). It is known that population subdivision with a large number of demes increases the amount of linkage disequilibrium among neutral loci when genomes are sampled from the same deme (Wakeley and Lessard 2003), because population subdivision increases local homozygosity, thereby reducing the effectiveness of recombination. This effect should also apply to associations between a locus under balancing selection and linked neutral sites. It is therefore important to examine the consequences of such subdivision for the effect of a diallelic balanced polymorphism on variability at a linked neutral site, and this is the main topic of the present paper.

For brevity, the locus under selection is referred here to exhibiting an inversion polymorphism, but the results apply to any Mendelian locus with two alleles maintained by balancing selection. An island model of a metapopulation of large size, divided into a finite number of demes of equal size, is assumed, with the same migration rate between all pairs of demes. As shown by Wakeley and Aliacar (2001), the properties of such a model are likely to provide a good approximation to more realistic scenarios, such as a two-dimensional stepping stone model, provided that the number of demes is large. In order to obtain simple results for equilibrium populations, it is assumed that the inversion is the derived state, and that selection on the inversion is sufficiently strong that it has risen quickly to an equilibrium frequency that is constant across demes. The properties of variability at the neutral locus, LD between the neutral locus and the inversion, and the extent of divergence between karyotypes at drift-mutation-recombination equilibrium, are studied first, followed by an examination of the approach to equilibrium. Here, “karyotype” is used to denote the state of a haplotype with respect to the arrangement which it carries.

Recursion relations for mean pairwise coalescent times are used here to obtain simple approximate expressions for the expected diversity and divergence statistics relevant to an inversion polymorphism at equilibrium under recombination and drift, and for the approach to their equilibrium values following the sweep of an inversion to a stable intermediate frequency. The effects of an inversion polymorphism on patterns of linkage disequilibrium are also examined. The reduction in effective recombination rate caused by population subdivision can have significant effects on these statistics, and hence on estimates of the ages of inversions. Methods are proposed for testing whether or not inversions are close to recombination-drift equilibrium, and for estimating the rate of recombinational exchange in heterozygotes for inversions, and a new method for determining the variances of pairwise coalescences times is described. It is concluded that many of the observed patterns of diversity at putatively neutral sites associated with inversion polymorphisms in *Drosophila melanogaster* are consistent with their being close to mutation-recombination-drift equilibrium.

### The model and its analysis

Assume that an autosomal inversion (*In*) is maintained at a frequency *x* and the standard arrangement (*St*) has frequency *y* = 1 – *x*. Without loss of generality, we can assume *x* ≤ ½ when considering equilibrium results; if this is not the case, then *In* and *St* can simply be interchanged. Parameters for *In* and *St* are denoted by subscripts 1 and 2, respectively. The population is assumed to be divided into local populations (demes) that are at equilibrium under mutation, genetic drift and migration at loci independent of the inversion. A Wright-Fisher model of reproduction is assumed, so that the effective population size of a deme is equal to its adult population size. An island model with a large number of demes, *d*, each with population size *N* is assumed, so that the migration effective population size (Nagylaki 1998) is *N*_*T*_ = *Nd*. Migration between populations occurs at rate *m* per generation.

The level of equilibrium neutral differentiation at autosomal loci that are independent of the inversion is measured by *F*_*ST*_ (Wright 1951), which is defined here as one minus the ratio of the mean coalescent time for pairs of alleles sampled from within a population to the mean coalescent time for pairs of alleles sampled randomly from the population as whole. With a large number of demes, *F*_*ST*_ ≈ 1/(1 + 4*Nm*) (Charlesworth 1998).

Random mating within local populations and with respect to arrangement status is assumed. The migration and recombination parameters, *m* and *r*, are assumed to be so small that second-order terms can be neglected. At a given neutral site within the region covered by the arrangement (or just outside it), recombinational exchange between arrangements caused by gene conversion and/or double crossing over occurs at rate *r* per generation. In the gametes produced by a heterokaryotype, *In*/*St*, there is a probability *r* that a given neutral site associated with the *In* haplotype came from the *St* haplotype, and that the homologous site associated with the *St* haplotype came from the *In* haplotype. It is likely that the value of *r* will depend on the location of the site within the inversion, with the largest values for sites within the inversion that are remote from the breakpoints, with the smallest values at sites close to the inversion breakpoints, due to the effects of the breakpoints in disturbing synapsis (Navarro et al. 1997), although the extent to which gene conversion events are influenced by proximity to the breakpoints is uncertain (Li et al. 2022). Sites outside the inversion will experience increasingly high rates of recombination with distance from the breakpoints as the effect of crossover suppression dies out (Koury 2023).

Inversion-carrying and standard haplotypes are denoted by indices 1 and 2, respectively. We need to distinguish between a pair of haplotypes that are sampled from the same deme (denoted by subscript *w*), and a pair of haplotypes sampled from two different demes (denoted by subscript *b*). The expected coalescent time for a sample of class *i*/*j* for a within-deme sample is denoted by *t*_*ijw*_ and the equivalent for a between-deme sample is *t*_*ijb*_. For a between-deme sample, the probability that a migrant haplotype came from the same deme as the non-migrant haplotype is 1/(*d –* 1) and the probability that it came from a different deme is (*d* – 2)/(*d –* 1). Simple recursions for the *t*’s can be obtained, which are given by Equations (A1) of the Appendix. Further simplification is provided by neglecting the products of 1*/Nx* and 1/*Ny* with *m* and *r* (Equations A2).

It is convenient to scale the migration rate and recombination rate by four times the deme size, writing *M* = 4*Nm* and *R* = 4*Nr*. In addition, coalescent times can be expressed relative to the expected coalescent time for a pair of alleles sampled from the same deme at a locus that is independent of the inversion, *T*_*Sn*_ = 2*N*_*T*_ = 2*dN*. Upper case *T*’s are used to denote *t*’s divided by *T*_*Sn*_. As shown in the Appendix, manipulation of the resulting equations leads to simple explicit approximate expressions for the equilibrium values of the *T*_*ijb*_ and *T*_*ijw*_, assuming a large number of demes and *M* >> *R* (Equations A5 and A6).

It is useful to consider the mean scaled coalescent time for pairs of alleles sampled randomly across karyotypes. For alleles from different demes, this is given by:

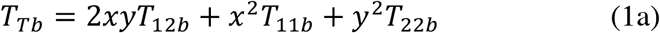

Similarly, following Charlesworth et al. (1997), the mean scaled mean coalescent time for a pair of alleles sampled within karyotypes, but from different demes, is:

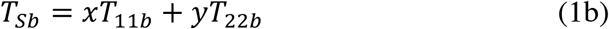

For measuring differentiation between sequences associated with alleles maintained by selection, a between-karyotype analogue of *F*_*ST*_ was defined by Charlesworth et al. (1997), which is analogous to the *K*_*ST*_ measure of Hudson, Boos and Kaplan (1992):

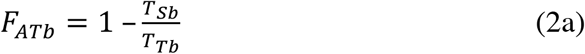

A related quantity, analogous to the <*F*_*ST*_> statistic of Hudson, Slatkin and Maddison (1992), is:

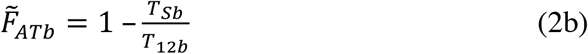

For equilibrium under recombination and drift, for which 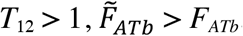, since *T*_12*b*_ > *T*_*Tb*_.

It should be borne in mind that different authors use different estimators for *F*_*ST*_ and *F*_*AT*_; the widely used methods of Weir and Cockerham (1984) and Hudson, Slatkin and Maddison (1992), which are mathematically equivalent, have much larger expected values than *K*_*ST*_ when the number of populations being compared is small (Charlesworth 1998; Gammerdinger et al. 2020), so that caution needs to be used when comparing *F*_*ST*_ or *F*_*AT*_ estimates from different studies.

Corresponding expressions apply to alleles sampled within demes, replacing subscript *b* by *w*. On general grounds, the Maruyama invariance principle that applies to structured coalescent processes with conservative migration (Maruyama 1974; Nagylaki 1982) implies that at equilibrium we have:

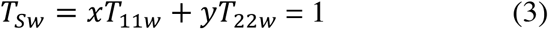

This result breaks down when *N*_*T*_*r* is close to zero; with *r* = 0, the two karyotypes behave as separate populations, with migration effective population sizes of *N*_*T*_*x* and *N*_*T*_*y*, respectively, so that *T*_*Sw*_ = *x*^2^ + *y*^2^.

Equation (3) implies that the equilibrium expressions for *F*_*ATw*_ and 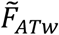 simplify to:

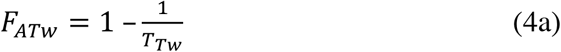

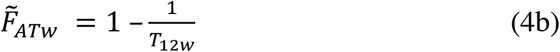

For equilibrium populations, therefore, the two *F*-statistics for within-population/between-karyotype differentiation contain no more information than do *T*_*Tw*_ and *T*_12*w*_, respectively; we have *T*_*Tw*_ =1/(1 – *F*_*ATw*_) and 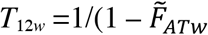. For applications of these formulae to population genomic data, the divergence statistics corresponding to the *T*’s need to be scaled by an estimate of mean neutral nucleotide site diversity at sites independent of the inversion, or by the weighted mean of the two within-karyotype diversities, as described in the *Discussion, Interpreting population genomic data on inversion polymorphisms*. The latter has the advantage that potential differences in mutation rates among different genomic regions are eliminated.

When the scaled migration rate *M* tends to infinity and *F*_*ST*_ tends to zero, the case of a panmictic population with population size *N*_*T*_ is approached, and the subscripts *w* and *b* can be dropped. The recursion relations for the within-deme statistics can be used for this case (Equations A2), setting *m* to zero and the deme size to *N*_*T*_. Application of the above method to this case, writing *ρ* = 4*N*_*T*_*r* for the scaled recombination rate, and assuming that *ρ* > 0, gives the following expressions for equilibrium:

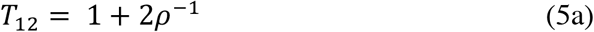

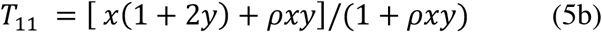

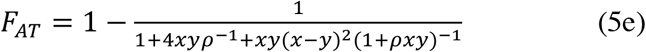

These are equivalent to Equations (A3) and (A4) of Nordborg (1997), Equations (6) of Navarro et al. (2000), and Equation (8) of Zeng et al. (2021), which were derived by more complex methods. Consistent with Equation (3), *T*_*S*_ = *xT*_11_ + *yT*_22_ = 1 at equilibrium, so that:

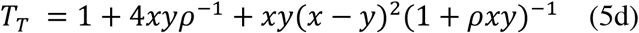

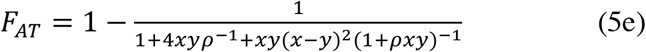

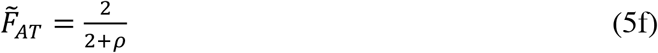

### Theoretical results: equilibrium populations

#### General considerations

Navarro et al. (2000) described results for the case of an equilibrium panmictic population, which is the limiting case when *F*_*ST*_ = 0. The focus here is therefore on subdivided populations, using the approximations for the equilibrium *T*’s given by Equations (A5) and (A6), which assume large *d* and *m* >> *r*. When comparing the theoretical results with observations, it is useful to note that the *T*’s under the infinite sites model of Kimura (1971) are proportional to the corresponding mean pairwise diversities or divergences per nucleotide site, taken over large numbers of neutral sites with the same mutation and recombination rates (e.g., Hudson 1990). The mean levels of nucleotide site diversity and divergence between arrangements within *D. melanogaster* populations are such that the infinite sites model fits the data reasonably well (Langley et al. 2012; Corbett-Detig and Hartl 2012; Kapun and Flatt 2023), so that the *T* values and their ratios presented in the figures below indicate the corresponding expected diversity and divergence values when interpreting data on inversion polymorphisms. Most multicellular organisms have similar or lower diversity values compared to *D. melanogaster* (Buffalo 2021), and so should present even less of a problem of interpretation.

#### Numerical results for subdivided populations

The results presented here are intended to represent a set of populations in a single geographical region that is isolated from other regions, with a within-deme neutral nucleotide site diversity value (*π*) similar to that found in population genomic surveys. The results from the *Drosophila* Genome Nexus Project, which has assembled genome sequence data for a large number of *D. melanogaster* genomes from natural populations (Lack et al. 2015, 2016), show that *F*_*ST*_ between pairs of populations within a region is generally low, around 0.05 or even less, and is only about 0.2 between continents (Fig. 3 of Lack et al. 2016). *F*_*ST*_ values in animals (Roux et al. 2016) and outcrossing flowering plants (Charlesworth 2003) rarely exceed 0.25, so that *F*_*ST*_ in the figures was restricted to the range 0 – 0.25 for purposes of illustration, consistent with the approximations used to generate the results.

In what follows, special attention is paid to the results with *F*_*ST*_ = 0.05, although the mean pairwise *F*_*ST*_ for inversion-free genomes between the Zambian *D. melanogaster* population (ZI) and two populations from South Africa (SP and SI) is only 0.007 (Lack et al. 2016, Fig. 3). This suggests that panmixia is a good approximation for populations in this region, which is thought be close to the center of origin of the species (Sprengelmeyer et al. 2020). The ZI population was a focus of the intensive study of the *In(3R)P* inversion by Kapun et al. (2023), discussed below in relation to the theoretical results. It is important to note that, under the assumptions made here, the equations for the *T*’s involve only the parameters *M* = 4*Nm* and *ρ* = 4*N*_*T*_*r* = 4*Ndr*; in turn, with large *d* we have *M* ≈ (1 – *F*_*ST*_)/*F*_*ST*_, where *F*_*ST*_ corresponds to the *K*_*ST*_ measure of differentiation among local populations for neutral loci independent of the inversion. Under these conditions, if *F*_*ST*_, *r* and *N*_*T*_ are specified, the results are not affected by the number of demes, the deme size (*N*) or the migration rate (*m*).

Results are displayed for four different values of the rate of recombination (*r*) between a neutral site and the arrangement in *In*/*St* heterokaryotypes, which fall within the range reported for single inversions in *Drosophila* (Navarro et al. 1997; Korunes and Noor 2019). Given that *r* is expected to be highest in the central regions of inversions, and lowest near their breakpoints (Navarro et al. 1997, 2000), increasing *r* in the figures can be interpreted as reflecting increasing distance from a breakpoint. Fig. 1 plots several expected pairwise coalescent times for within- and between-deme samples against *F*_*ST*_ for the case of an inversion frequency of 0.1. Fig. 2 shows the corresponding ratios of *T*_11_/*T*_22_ for within- and between-deme samples, as well as the *F*_*AT*_ statistics. Figs. S1 and S2 in the Supplementary Information give the results for an inversion frequency of 0.5.

**Figure 1.**
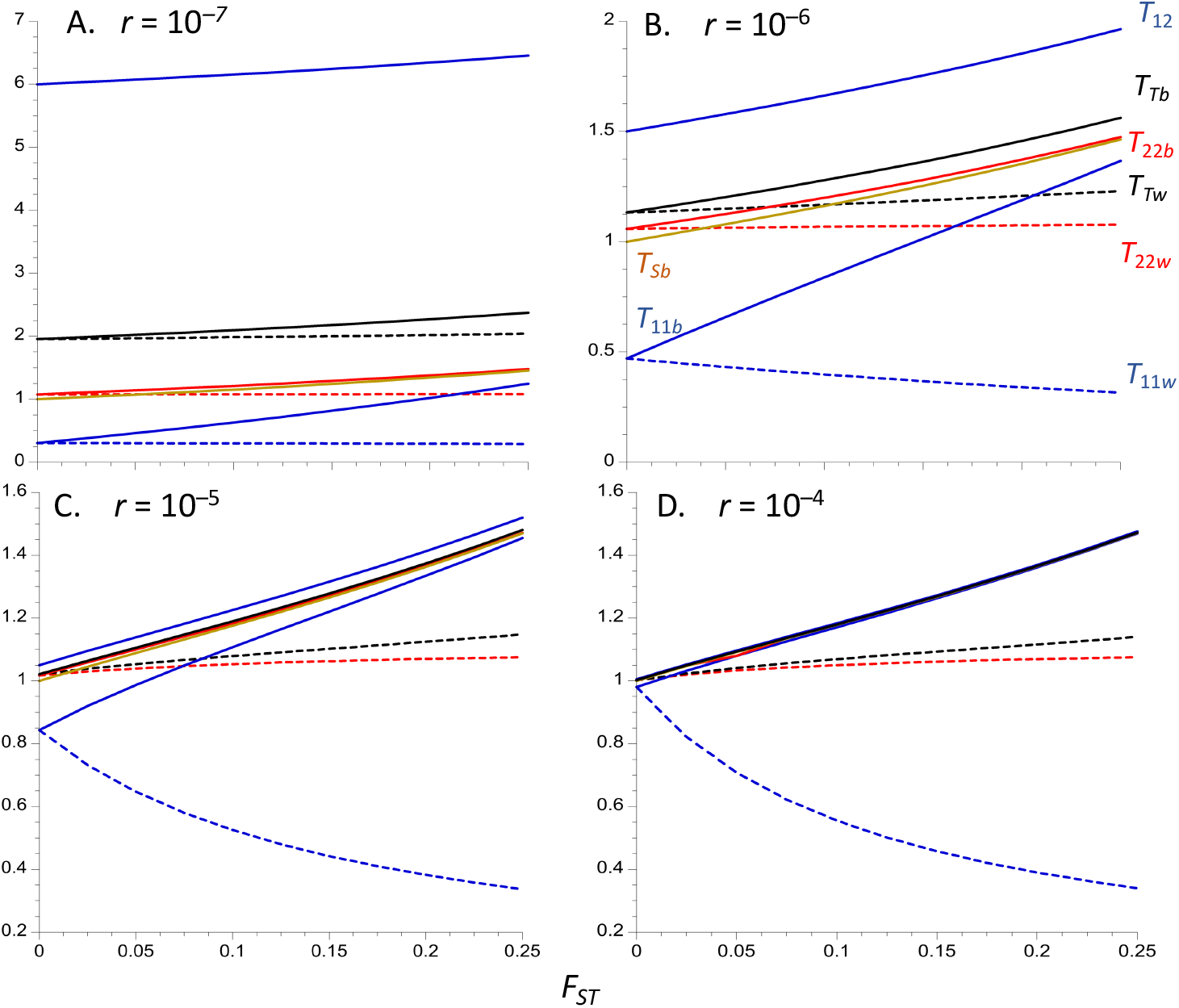
Equilibrium expected coalescence times (relative to 2*N*_*T*_) for an inversion polymorphism where the inversion is maintained at a constant frequency of 0.1. Four different recombination rates in heterokaryotypes (*r*) are modeled, as indicated at the top left of each panel. An island model of population structure is assumed, with 200 demes and a total population size of *N*_*T*_ = 10^6^, so that individual demes have a population size of *N* = 5000. The X axis is the equilibrium *F*_*ST*_ for neutral sites unlinked to the inversion. Subscripts 1 and 2 denote alleles sampled from the inversion and standard arrangement, respectively; subscripts *w* and *b* denote alleles sampled from the same and from separate demes, respectively; subscript *T* denotes pairs of alleles sampled without regard to karyotype. The dashed curves represent within-deme coalescent times, and the solid curves are between-deme coalescent times; red is *T*_12_, black is *T*_*Tw*_ and *T*_12_, blue is *T*_11_ and red is *T*_22_ (for *T*_12_, there is no significant difference between with- and between-deme values.) The mean within-karyotype values for between-deme samples (*T*_*sb*_) is the solid beige curve; the within-deme equivalent (*T*_*sw*_) is equal to 1 for all r and *F*_*ST*_ values and is not displayed.

**Figure 2.**
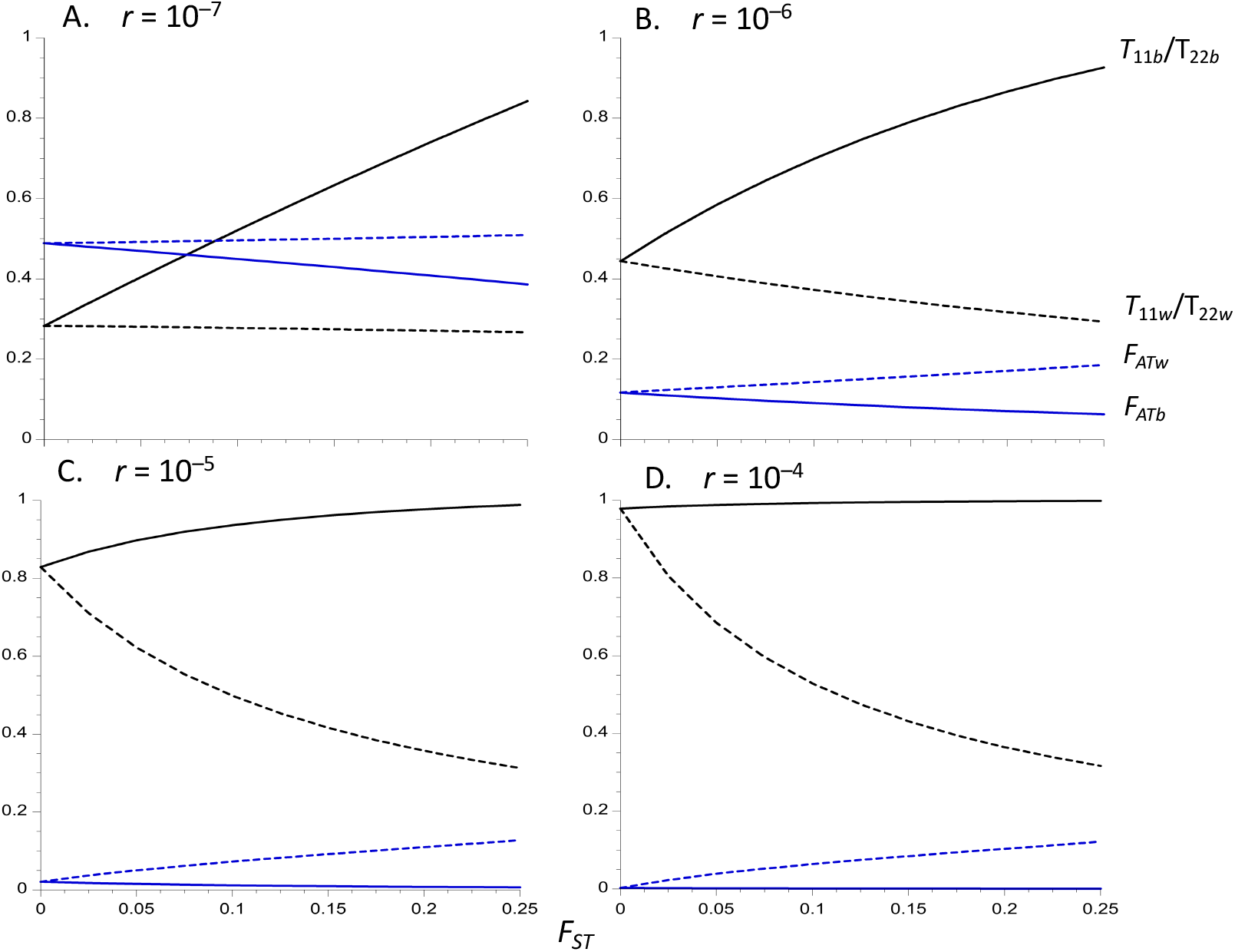
Equilibrium values of *F*_*ATw*_ (blue dashed curves), *F*_*ATb*_ (blue solid curves), *T*_11*w*_/*T*_22*w*_ (black dashed curves) and *T*_11*b*_/*T*_22*b*_ (black solid curves), for an inversion polymorphism where the inversion is maintained at a constant frequency of 0.1. The population and recombination parameters in Figure 1 are used.

The mean coalescent time between *In* and *St* relative to 2*N*_*T*_, *T*_12_, is almost the same for within-deme pairs of alleles as for alleles from separate demes (see Equations A6d and A6e), so that only the between-deme values are shown. This property reflects the fact that, with large *d*, the probability that two alleles sampled from two distinct demes were derived from the same deme in the recent past (where they have an opportunity to recombine) is very small compared with the probability that alleles sampled from the same deme were derived from separate demes (and cannot recombine); given that *m* is assumed to be >> *r*, migration is the major factor affecting alleles sampled from the same deme. This causes the properties of pairs of alleles sampled from the same or different demes (other than their probabilities of coalescing) to converge; there is, of course, no immediate chance of coalescence for two alleles sampled from different arrangements.

*T*_12_ provides a measure of the expected net sequence divergence between a pair of alleles from the two different karyotypes, relative to the expected within-deme neutral diversity for loci independent of the inversion, or to the expected neutral diversity averaged over *In* and *St* haplotypes sampled randomly from the same deme. This because these diversity measures are both equal to *θ* = 4*N*_*T*_*u*, where *u* is the mutation rate per basepair (Nagylaki 1998)– see Equation (3). For *u* = 5 × 10^−9^ (Assaf et al. 2017) and *N*_*T*_ = 10^6^, a value of *T*_12_ = 1 corresponds to a sequence divergence of *T*_12_*θ* /2 = 0.01 per basepair, which is approximately equal to the within-population synonymous site diversity in *D. melanogaster* ancestral range populations (Langley et al. 2012). As can be seen from Fig. 1, *T*_12_ is quite close to 1 for *r* = 10^−5^ or 10^−4^ with *F*_*ST*_ ≤ 0.05, but takes much larger values for the two lower recombination rates. *T*_12_ increases with *F*_*ST*_, but only slowly at the two lower recombination rates, and is at most about 1.5 for the two higher *r* values.

Unless *r* is very low, as may be the case near inversion breakpoints (Navarro et al. 1997), the equilibrium level of sequence divergence between *In* and *St* is thus likely to be less than 50% larger than the genome-wide mean neutral diversity, corresponding to an 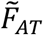 of 0.33. Equation (A6d) shows that the effect of *F*_*ST*_ on *T*_12_ is caused by the increase in divergence between alleles sampled from different karyotypes and different populations as the between-deme coalescent times increase. The effect of recombination on *T*_12_ is controlled by the reciprocal of *ρ*, the recombination rate scaled by 4*N*_*T*_, which is invariant with respect to the level of population subdivision under the assumptions used here.

A related statistic is the net mean coalescent time for a pair of alleles sampled randomly within demes without regard to karyotype (*T*_*Tw*_), which determines the overall mean within-deme *π* for nucleotide sites located within the region covered by the inversion. Figs. 1 and S1 show that *T*_*Tw*_ is much less sensitive to *F*_*ST*_ than *T*_12_, but increases slightly with increasing *F*_*ST*_, especially at high recombination rates, reflecting the fact that it is heavily influenced by the within-karyotype coalescent times *T*_11*w*_ and *T*_22*w*_, which either change in opposite directions as *F*_*ST*_ increases (for *x* = 0.1, or remain constant (for *x* = 0.5). This also means that *T*_*Tw*_ < *T*_12_, even for an intermediate frequency inversion, so that the corresponding diversity statistic provides a less useful index of between-karyotype differentiation than the between-karyotype divergence. The corresponding statistic for alleles sampled from different demes (*T*_*Tb*_) has similar properties, but is always somewhat larger in magnitude, especially at high *F*_*ST*_ values, due to the inflation of beween-deme coalescent times with restricted migration. Unless *r* is very low, an inversion polymorphism at recombination-drift equilibrium is not expected to have a large effect on the overall level of sequence diversity in the population unless there is a high degree of population subdivision, consistent with the results for chromosome arm 3R of *D. melanogaster* shown in Fig. 4 of Corbett-Detig and Hartl (2012).

When the inversion is present at a low frequency, as in Figs. 1 and 2, the mean coalescent times for pairs of alleles sampled within the inversion (*T*_11_) are smaller than those for alleles sampled within *St* (*T*_22_), due to the lower effective population size of carriers of *In*; this of course also applies to the corresponding *π* values (Nordborg 1997; Navarro et al. 2000). This difference disappears when *In* and *St* have equal frequencies, since their effective sizes are necessarily equal, and is smaller, the smaller the difference in frequencies. The converse is true if *In* is more frequent than *St*. Figure 2 shows that the ratio *T*_11*w*_/*T*_22*w*_ decreases considerably as *F*_*ST*_ increases when *r* is relatively large, whereas *T*_11*b*_/*T*_22*b*_ increases slightly. Conversely, *T*_11*b*_/*T*_22*b*_ increases considerably with *F*_*ST*_ when *r* is small, but *T*_11*w*_/*T*_22*w*_ hardly changes. *T*_11*w*_/*T*_22*w*_ is, however, always greater than the value of 0.11 expected from the relative frequencies of *In* and *St*, reflecting the effect of recombination in reducing allele frequency differences; consistent with this, both *T*_11*w*_/*T*_22*w*_ and *T*_11*b*_/*T*_22*b*_ increase with *r*.

All other things being equal, therefore, the departure of estimates of *T*_11*w*_/*T*_22*w*_ from the ratio of the frequencies of *In* and *St* provides an inverse measure of the extent of differentiation between karyotypes when the frequency ratio departs considerably from unity. The behavior of *T*_11*w*_/*T*_22*w*_ as a function of *r* reflects the fact that, for a low frequency inversion, *T*_11w_ increases considerably with increasing *r* for a given *F*_*ST*_ whereas *T*_22w_ decreases slightly, as seen in Figure 1. This complementary behavior arises from the fact that *xT*_11w_ + *yT*_22*w*_ = 1 at equilibrium, by Maruyama’s invariance principle (Equation 3). An increase in *F*_*ST*_ results in a reduced effective rate of recombination for within-deme samples, due to increased homozygosity (Wakeley and Lessard 2003) so that *T*_11*w*_ decreases with *F*_*ST*_, whereas *T*_22*w*_ increases. *T*_11*b*_ and *T*_22*b*_ both increase with *F*_*ST*_, especially with high *r* values, reflecting the effects of reduced migration rates on coalescence times; for between-deme samples, population subdivision of the kind modeled here does not affect the effective recombination rate (Wakeley and Lessard 2003).

The between-karyotype analogues of *F*_*ST*_, *F*_*AT*_ and 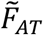 (see Equations 4), are often used as a measure of the extent of genetic differentiation among karyotypes, rather than the absolute divergence measures just discussed. As noted by Charlesworth et al. (1997) in the context of a single-locus balanced polymorphism (see also Zeng et al. [2021]), *F*_*AT*_ is equivalent to the 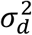 measure of linkage disequilibrium of Ohta and Kimura (1971), treating the neutral site and karyotype as a pair of loci, but it is considerably less tedious to estimate from population genomic data. Its properties thus differ somewhat from those of *T*_12_ or *T*_*Tw*_, especially as it is heavily influenced by the within-karyotype coalescent time in the numerator of Equation (4a) (Charlesworth 1998). Figs. 2 and S2 show that *F*_*ATw*_ and *F*_*ATb*_ are only weakly affected by the extent of population subdivision at the lower two recombination rates, with *F*_*ATw*_ increasing slightly with *F*_*ST*_ whereas *F*_*ATb*_ decreases; *F*_*ATw*_ is always larger than *F*_*ATb*_, especially when *F*_*ST*_ and *r* are large, and is thus more useful as a measure of between-karyotype differentiation. This difference reflects the effect of population subdivision on within-deme samples, and lack of such an effect for between-deme samples, described above in connection with the ratio *T*_11*w*_/*T*_22*w*_. Both statistics are highly sensitive to *r*, with small values at the two higher *r* values, especially for 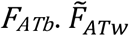 for equilibrium populations is given by 1 – 1/*T*_12_ (Equations 4b and 5f), so that it behaves in a similar fashion to *T*_12_ as a function of *r* and *F*_*ST*_ and is therefore not shown here.

A moderately high level of population subdivision in an equilibrium system thus makes it much easier to detect between-karyotype differentiation at low recombination rates, as measured by 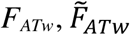 or *T*_12_, but makes *F*_*ATb*_ a less useful measure. Comparisons of the mean between-karyotype divergence (either within or between demes) with the mean within-deme and within-karyotype diversity for samples at loci independent of the inversion (corresponding to a *T*_*Sw*_ of 1), or with the mean within-deme and within-karyotype diversity (corresponding to a *T*_*Sn*_ of 1), are probably the most useful measures of the extent of between-karyotype divergence, if equilibrium can be assumed.

It is of some interest to compare the values of *F*_*AT*_ for a polymorphism maintained at constant frequencies with the value of 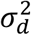 for a pair of neutral loci at statistical equilibrium under recombination and drift in a subdivided population. Table 1 shows results for the within-deme measure of *F*_*AT*_ compared with 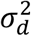 for within-deme samples calculated from the equations in the Appendix to Wakeley and Lessard (2003). For the intermediate and highest recombination rates (*ρ* = 4 and *ρ* = 40), *F*_*AT*_ for *x* = 0.5 is very close to 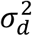, whereas *F*_*AT*_ for *x* = 0.1 is substantially smaller, except for the panmictic case (*F*_*ST*_ = 0); all three variables increase considerably with increasing *F*_*ST*_, reflecting the reduced effective recombination rate when there is extensive population subdivision (Wakeley and Lessard 2003). In contrast, for the lowest recombination rate (*ρ* = 0.4), *F*_*AT*_ for *x* = 0.1 is closer to 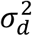, and there is only a small increase in the three measures with *F*_*ST*_. For a pair of neutral loci, 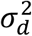 for a sample of haplotypes taken from separate demes (which is equivalent to random sampling from the whole population with large *d*) is independent of the extent of subdivision with the model used here (Wakeley and Lessard 2003), in contrast to the behavior of *F*_*ATb*_ shown in Figs 1 and 2. The results for neutral loci thus shed only limited light on what is to be expected for an inversion polymorphism maintained at a constant frequency. The discrepancy between the statistics for the neutral case and the case with a constant inversion frequency could in principle be used as a test for selection, although this would require accurate knowledge of *r*.

**Table 1.** Comparisons of within-deme *F*_*AT*_ for a balanced polymorphism versus 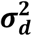 for a pair of neutral loci

| $F_{ST}$ | $F_{AT}$<br>$x = 0.1$ | $F_{AT}$<br>$x = 0.5$ | $\sigma_d^2$ | $F_{AT}$<br>$x = 0.1$ | $F_{AT}$<br>$x = 0.5$ | $\sigma_d^2$ | $F_{AT}$<br>$x = 0.1$ | $F_{AT}$<br>$x = 0.5$ | $\sigma_d^2$ |
| --- | --- | --- | --- | --- | --- | --- | --- | --- | --- |
| 0 | 0.489 | 0.714 | 0.380 | 0.117 | 0.200 | 0.156 | 0.021 | 0.024 | 0.023 |
| 0.05 | 0.492 | 0.718 | 0.419 | 0.130 | 0.232 | 0.200 | 0.051 | 0.072 | 0.070 |
| 0.10 | 0.496 | 0.723 | 0.454 | 0.143 | 0.265 | 0.241 | 0.073 | 0.120 | 0.114 |
| 0.15 | 0.500 | 0.728 | 0.488 | 0.157 | 0.299 | 0.281 | 0.092 | 0.167 | 0.156 |
| 0.20 | 0.504 | 0.733 | 0.521 | 0.171 | 0.333 | 0.320 | 0.110 | 0.216 | 0.197 |
| 0.25 | 0.509 | 0.739 | 0.552 | 0.186 | 0.368 | 0.358 | 0.128 | 0.263 | 0.238 |

The scaled recombination rates of $\rho = 0.4, 4$ and $40$ correspond to $r = 10^{-7}, 10^{-6}$ and $10^{-5}$ , respectively, when $N_T = 10^6$ , as assumed in the figures.

As expected from Equation (5a), a comparison of Figs. 1 and 2 with Figs. S1 and S2 shows that *T*_12_ is not greatly affected by the inversion frequency; in contrast, *F*_*ATw*_ and *F*_*ATb*_ are larger with equal frequencies of the two karyotypes than when the inversion is either rare or very common. This reflects the smaller *T*_*T*_ values with extreme inversion frequencies, as can be seen from Equation (5d) for the panmictic case, where *T*_*T*_ is approximately 1 + 2*ρ* ^−1^ for *x* = 0.5, but approaches 1 + *x*(4*ρ* ^−1^ +1) as *x* tends to 0. This is another reason for using between-karyotype divergence relative to mean within-deme diversity as a measure of differentiation between karyotypes.

### Theoretical results: Approach to equilibrium

#### General considerations

The above results assume both that the inversion has reached its equilibrium frequency and that there has been sufficient time for the effects of coalescence and recombination to equilibrate. In reality, as was assumed in early hitchhiking models of associations between electrophoretic variants and inversions (Ishii and Charlesworth 1977), a polymorphic inversion is likely to have arisen on a single unique haplotype drawn at random from the initial population. Having survived early stochastic loss, it will have approached its equilibrium frequency over a time that is inversely related to the strength of selection acting on it. Molecular characterizations of inversion breakpoints in *D. melanogaster* and *D. subobscura* strongly support the unique origin hypothesis (Corbett-Detig and Hartl 2012; Orengo et al. 2019; Kapun et al. 2023). After this selective equilibrium has been approached, there will be another extended period during which mutation-recombination-drift equilibrium is approached. In the case of a panmictic population, both of these episodes can be included in the same model, using phase-type theory (Zeng et al. 2021).

But the population genetics of the spread of a new mutation in a subdivided population, and its hitchhiking effects on linked neutral sites, is much more complex (Barton et al. 2013), and has not been applied to the case of balancing selection. In the present treatment of subdivided populations, the initial approach to the equilibrium inversion frequency is assumed to occur effectively instantaneously, so that only the second phase of the approach to equilibrium is studied. Clearly, this is likely to cause the time taken to approach equilibrium and the effects of recombination during this period to be underestimated, due to the additional time needed for a mutation to spread through a subdivided population compared with panmixia, even if the habitat is two-dimensional rather than one-dimensional (Barton et al. 2013).

By using this simplifying assumption, we can set *T*_22*w*_ = *T*_22*b*_ = 0, *T*_12*w*_ = *T*_11*w*_ = 1, and *T*_12*b*_ = *T*_11*b*_, with *T*_11*b*_ ≈ 1/(1 – *F*_*ST*_) when *d* is large. Using these as initial values, the change per generation in the deviations of the *T*’s from their equilibrium values can be calculated by means of the matrix iteration ***x***_*n*_ = ***A x***_*n*–1_, where ***x***_*n*_ is the column vector of deviations of the *T*’s in generation *n* from their equilibrium values (see section S1 of the Supplementary Appendix for details). This approach breaks down in the absence of recombination, since in this case the divergence between *In* and *St* increases without limit as time increases; a separate treatment of this case is given in sections S6 and S7 of the Supplementary Appendix.

In order to speed up calculations, a relatively small total population size (10^5^ for the case of a subdivided population, and 10^4^ for the equivalent panmictic population) was assumed, with rescaling of parameters to keep their products with *N*_*T*_ identical with those used for the equilibrium results. Insight into the asymptotic rate of approach to equilibrium when *d* is large can be obtained from the eigenvalues of the ***A*** matrix. As shown in section S3 of the Supplementary Appendix, if second-order terms in 1/*N, r* and *m* are neglected, the structure of ***A*** is such that its 6 eigenvalues (3 in the panmictic case) are each approximately equal to one of its diagonal elements, i.e., to 1 – 2*m* – 2*ry* – 1/(2*Nx*), 1 – 2*m*/*d* – 2*ry*, 1 – 2*m* – 2*rx –* 1/(2*Ny*), 1 – 2*m*/*d* – 2*rx*, 1 – 2*m* – *r*, and 1 – 2*m*/*d* – *r*, respectively. The asymptotic rate of approach to equilibrium is controlled by the largest of these quantities; which of the six is the largest is determined by the relative values of *Nx, Ny, rx, ry* and *m*. Since *m* >> *r* and *d* >> 1 with the approximations used here, either 1 – 2*m*/*d* – 2*ry* or 1 – 2*m*/*d* – 2*rx* will be the largest eigenvalue, showing that the asymptotic rate of approach to equilibrium is largely controlled by the product of 2*r* and the smaller of the two frequencies *x* and *y* when *d* is very large. In the case of a panmictic population, the system reduces to a three-dimensional one with eigenvalues approximately equal to 1 – 2*ry* – 1/(2*Nx*), 1 – 2*rx* – 1/(2*Ny*), and 1 – *r*, so that a similar conclusion applies.

The timescale of the approach of the coalescence times to equilibrium after the inversion has approached its equilibrium frequency is thus of the order of the inverse of *rx* (if x ≤ 0.5 or *ry* (if *x* > 0.5), unless *r* is very close to zero – see sections S6 and S7 of the Supplementary Appendix for this case. It is nearly independent of population structure with large *d* and *m* >> *r*. The full solution for ***x***_*n*_ in generation *n* in terms of the representation of ***A***^***n***^ by the eigenvalues and eigenvectors of ***A*** is given in section 3 of the Supplementary Appendix. In practice, however, it simpler to iterate the basic matrix recursion – it takes only a few seconds to iterate several million generations on a laptop computer.

It is also of interest to examine the initial rates of change of the *T*s using the starting point of an instantaneous sweep to equilibrium described above. The corresponding initial values of *F*_*ATw*_ and *F*_*ATb*_ are then both equal to 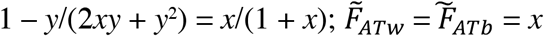. The initial *F*_*AT*_ statistics can thus be substantially different from zero despite the absence of any divergence between *In* and *St*, due to the assumed lack of variability within inverted chromosomes. This is seen in other systems such as X-Y comparisons when a newly evolved Y chromosome lacks variability (Bergero et al. 2019); Toups et al. 2019); Gammerdinger et al. 2020).

Normalizing Equations (A2) by dividing each term by 2*N*_*T*_, neglecting second-order terms in the products of the deviations of the *T*’s from their equilibrium values with *m, r* and 1/*N*, and exploiting the fact that 4*Nm* ≈ (1 – *F*_*ST*_)/*F*_*ST*_ in order to simplify Equations (A6c - A6e), the following simple relations hold for the initial changes per generation in the *T*’s:

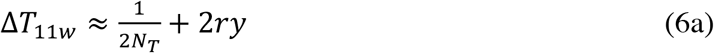

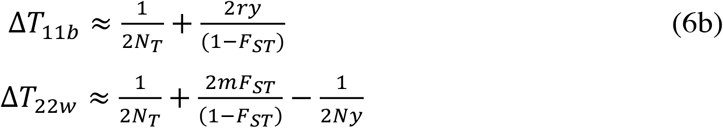

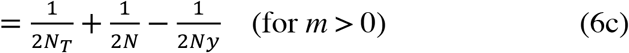

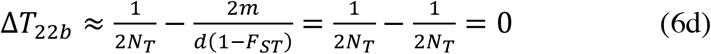

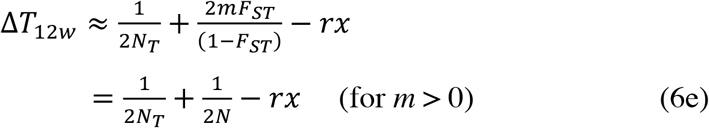

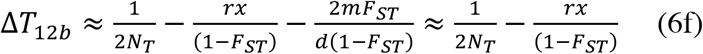

Using the results in section S4 of the Supplementary Appendix, we also have:

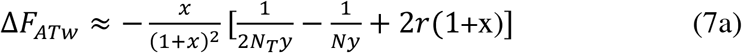

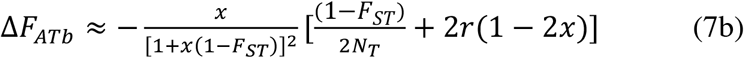

These results require *rx* and *ry* to both exceed 1/2*N*_*T*_, and hence are invalid for very low rates of recombination, as in the case of *ρ* = 0.4 shown in Figures 3-5 below.

**Figure 3.**
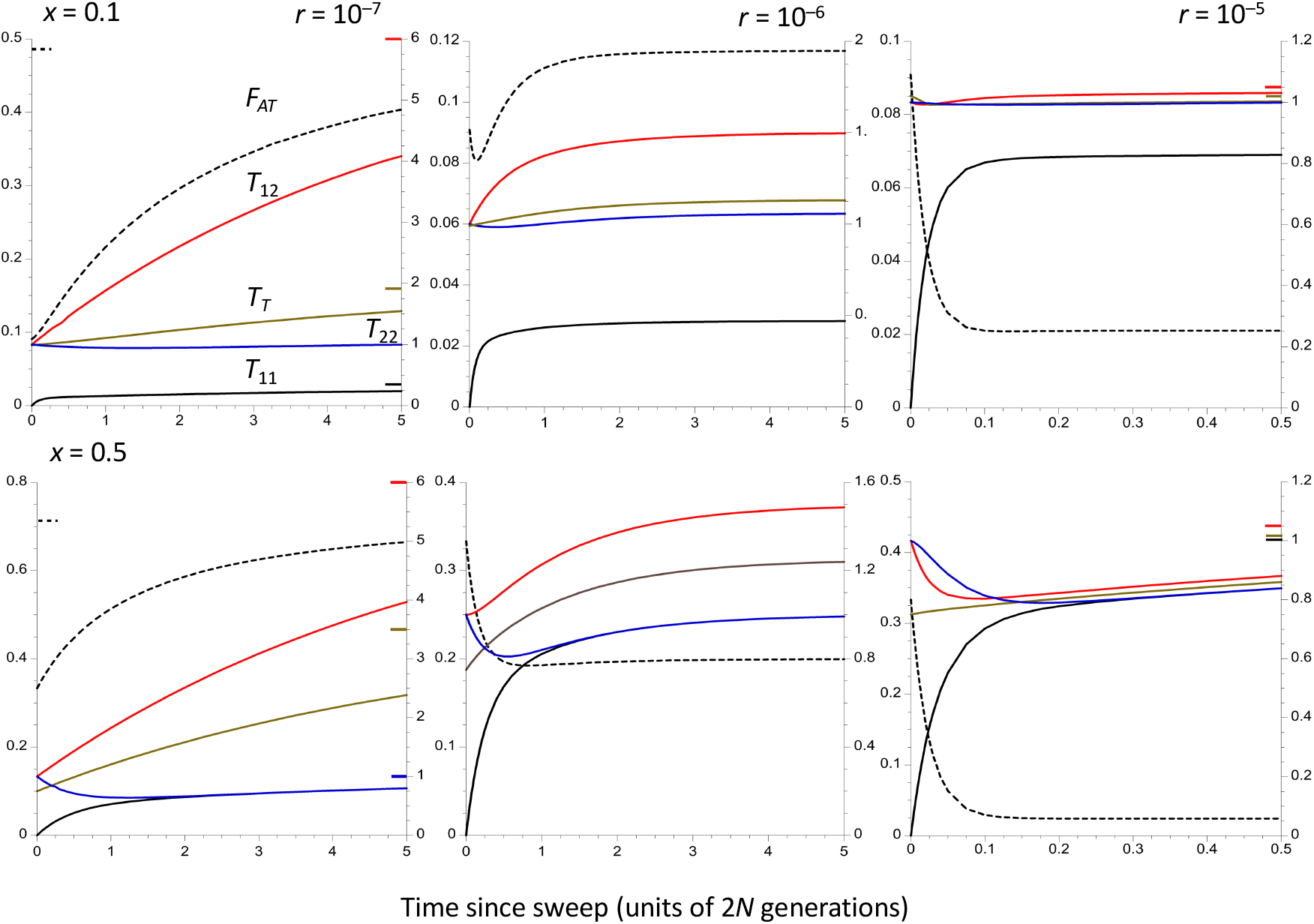
The trajectories of change in the population statistics for the case of a panmictic population of size *N* = 10^6^, assuming that the time taken to approach the equilibrium inversion frequency is negligible compared with the coalescent time of 2*N* generations. The X axes display times in units of coalescent time following the sweep to equilibrium. Three different recombination rates in heterokaryotypes are shown, as well as two different frequencies of the inversion (0.1 in the upper panels and 0.5 in the lower panels). The dashed curves are *F*_*AT*_, whose values are displayed on the left-hand Y axes. The solid curves are mean coalescent times, measured relative to 2*N*; red is T_12_, brown is *T*_*T*_, black is *T*_11_ and blue is *T*_22_ (right-hand Y axes). For the highest rate of recombination (*r* = 10^−5^), only the first *N* generations are shown, in order to capture the rapid changes at the start of the process. The coloured bars inside the Y axes indicate the equilibrium values of the corresponding statistics, for cases when these are substantially different from the final values of the statistics.

Equations (6a), (6c), (6e) and (7a) can be applied to the panmictic case, by setting the right-hand terms involving *m* and 1/2*N* (but not 1/2*Ny*) to zero, and equating *N* and *N*_*T*_.

#### Results for a single randomly mating population

The results for a single, randomly mating population of size *N* = *N*_*T*_ are considered first, assuming an instantaneous sweep of a new inversion to its equilibrium frequency. This case was previously studied by Navarro et al. (2000) using coalescent simulations. Some representative results are shown in Fig. 3 for three different recombination rates adjusted to the values for *N* = 10^6^, i.e., scaled recombination rates of *ρ* = 4*Nr* = 0.4, 4 and 400, respectively), and two different equilibrium frequencies, *x* = 0.1 and *x* = 0.5. Time is measured relative to 2*N* = 2*N*_*T*_, and the results are thus invariant with respect to *N* for constant *ρ* (unless *N* is sufficiently small that the assumptions of the coalescent process are violated). The accuracy of the approximations used to generate these results was checked by coalescent simulations and found to be excellent (see Table S2). Further details of results based on the recursion relations are given in Table S3.

As shown above, the initial value of *F*_*AT*_ after an instantaneous sweep of the inversion to its equilibrium frequency is simply *x*/(1 + *x*). *F*_*AT*_ does not necessarily change monotonically over time, as can be seen for the case of *x* = 0.1 with *r* = 10^−6^, where it first decreases and then increases. With the highest recombination rate, the final direction of change of *F*_*AT*_ can be opposite to that for *T*_12_ because its initial value is greater than its equilibrium value, so that *F*_*AT*_ declines over time, reflecting the negative term in *r* in Equation (7a). In contrast, *T*_12_ increases on the approach to equilibrium, after an initial decline when *r* is large, as predicted by Equation (6e) (see the results for *x* = 0.5 and *r* = 10^−5^).

As expected intuitively, and as is consistent with Equation (6a), the mean coalescent time for the inversion (*T*_11_) always increases with time towards its equilibrium value, reflecting the effect of recombination in causing it to share ancestry with the standard sequence, and the fact that the carriers of the inversion increase in numbers away from a completely bottlenecked haploid population size of one. Even with the lowest frequency of recombination (*ρ* = 0.4), there is a relatively fast approach of *T*_11_ towards its equilibrium value compared with *F*_*AT*_ and *T*_12_, over a timescale that is much less than the neutral coalescent time. Except for the lowest recombination rate, relatively large values of *F*_*AT*_ and *T*_12_ (when compared with the values for the higher two recombination rates) are approached over this timescale, reflecting their large equilibrium values.

With the lowest recombination rate, however, there is a long period of time when *F*_*AT*_ and *T*_12_ are both much lower than their equilibrium values. Consistent with Equation (6c), *T*_22_ decreases initially for all three recombination rates, although very slowly if *Ny* is close to *N*. In some cases (e.g., with *x* = 0.5 and *r* = 10^−5^), *F*_*AT*_ changes non-monotonically, with an initial decrease followed by an increase. This reflects the fact that, for these parameter values, Equation (7a) predicts an initial decrease in *F*_*AT*_, whereas its final value is greater than its initial value.

A simple analytical solution to the trajectories of the mean coalescent times for the case of no recombination can also be derived (see section S6 of the Supplementary Appendix). In this case, *T*_12_ increases linearly with time on the coalescent timescale (Equation S5a), whereas *T*_11_ and *T*_22_ experience an exponential decay of their deviations from their respective equilibrium values of *x* and *y*, with rate constants *x* and *y*. In the case of an instantaneous sweep of *In* to its equilibrium frequency, this implies that *T*_11_ always increases over time, and *T*_22_ always decreases. For sufficiently large *T*, these expressions imply that *F*_*AT*_ increases monotonically towards one, due to the fact that *T*_12_ increases without bound in the absence of recombination. Approximations to these expressions for small *T* show that *F*_*AT*_ always increases initially (see Equations (S5e) – (S5i) in the Supplementary Appendix).

#### Results for a subdivided population

The numerical values of the population statistics were generated by using scaled recombination and migration parameters that matched those used for Figs. 1 and 2. Figs. 4 and 5 show the results for two different values of *F*_*ST*_ at neutral sites independent of the inversion, corresponding to scaled migration rates of 4*Nm* = 19 and 5.67, respectively. Further details are given in Tables S4 and S5.

**Figure 4.**
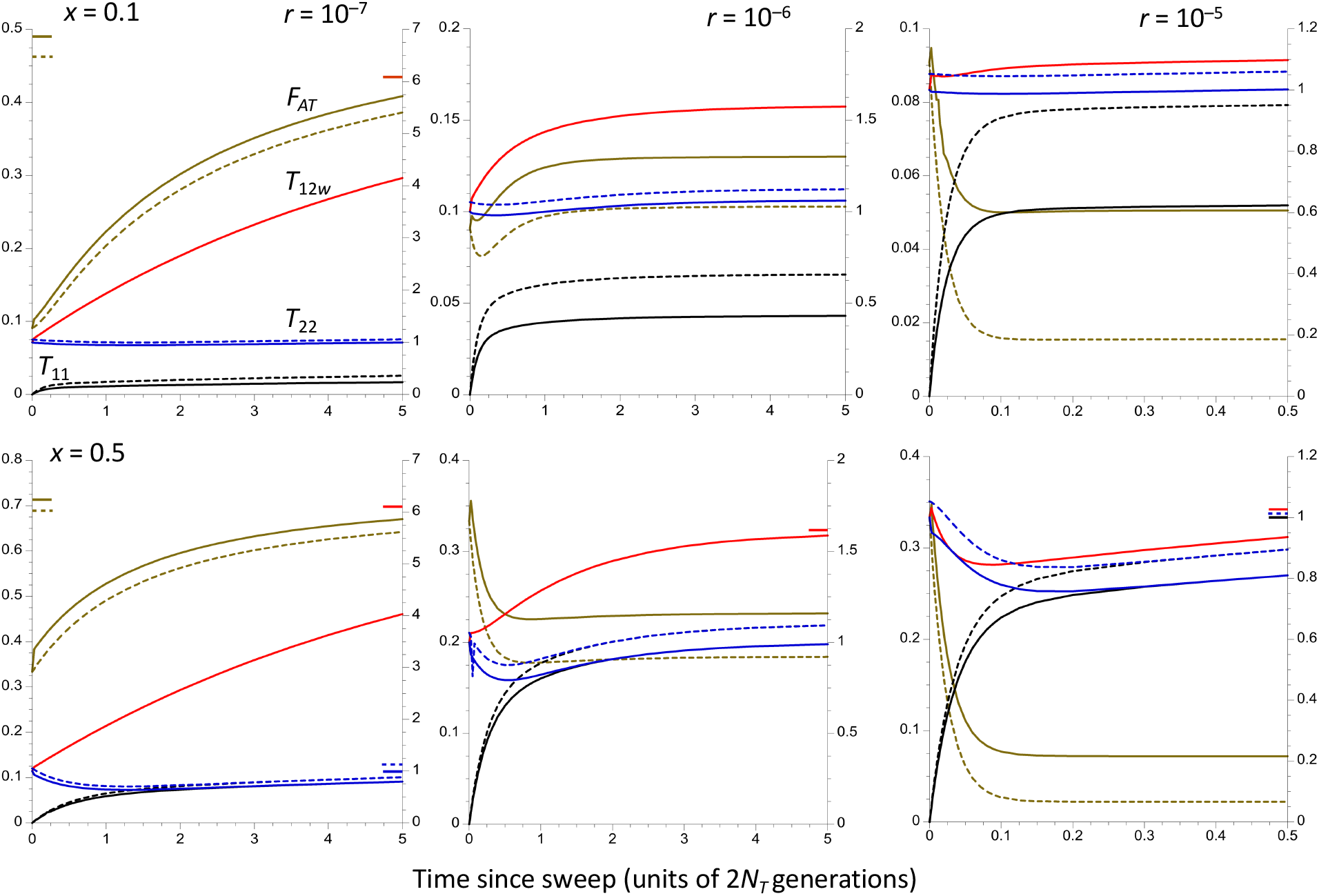
The trajectories of change in the population statistics for the case of a subdivided (island model) population of total size *N*_*T*_= 10^6^, with 200 demes of size *N* = 500 and with an *F*_*ST*_ for neutral sites independent of the inversion of 0.05 (i.e., 4*Nm* = 19). It is assumed that the time taken to approach the equilibrium inversion frequency is negligible compared with the coalescent time of 2*N*_*T*_ generations. The X axes display times in units of coalescent time following the sweep to equilibrium. Three different recombination rates in heterokaryotypes are shown, as well as two different frequencies of the inversion (0.1 in the upper panels and 0.5 in the lower panels). The solid curves represent within-population statistics and the dashed curves are between-population statistics. The values of *F*_*ATw*_ and *F*_*ATb*_ (brown curves) are given by the left-hand Y axes. The other curves are mean coalescent times, measured relative to 2*N*_*T*_; red is *T*_12*w*_ (*T*_12*b*_ behaves almost identically, except for its higher initial value and slower rate of increase when *T* < 0.005); black is *T*_11_ and blue is *T*_22_ (right-hand Y axes). For the highest rate of recombination (*r* = 10^−5^), only the first *N*_*T*_ generations are shown. The coloured bars inside the Y axes indicate the equilibrium values of the corresponding statistics, for cases when these are substantially different from the final values of the statistics.

**Figure 5.**
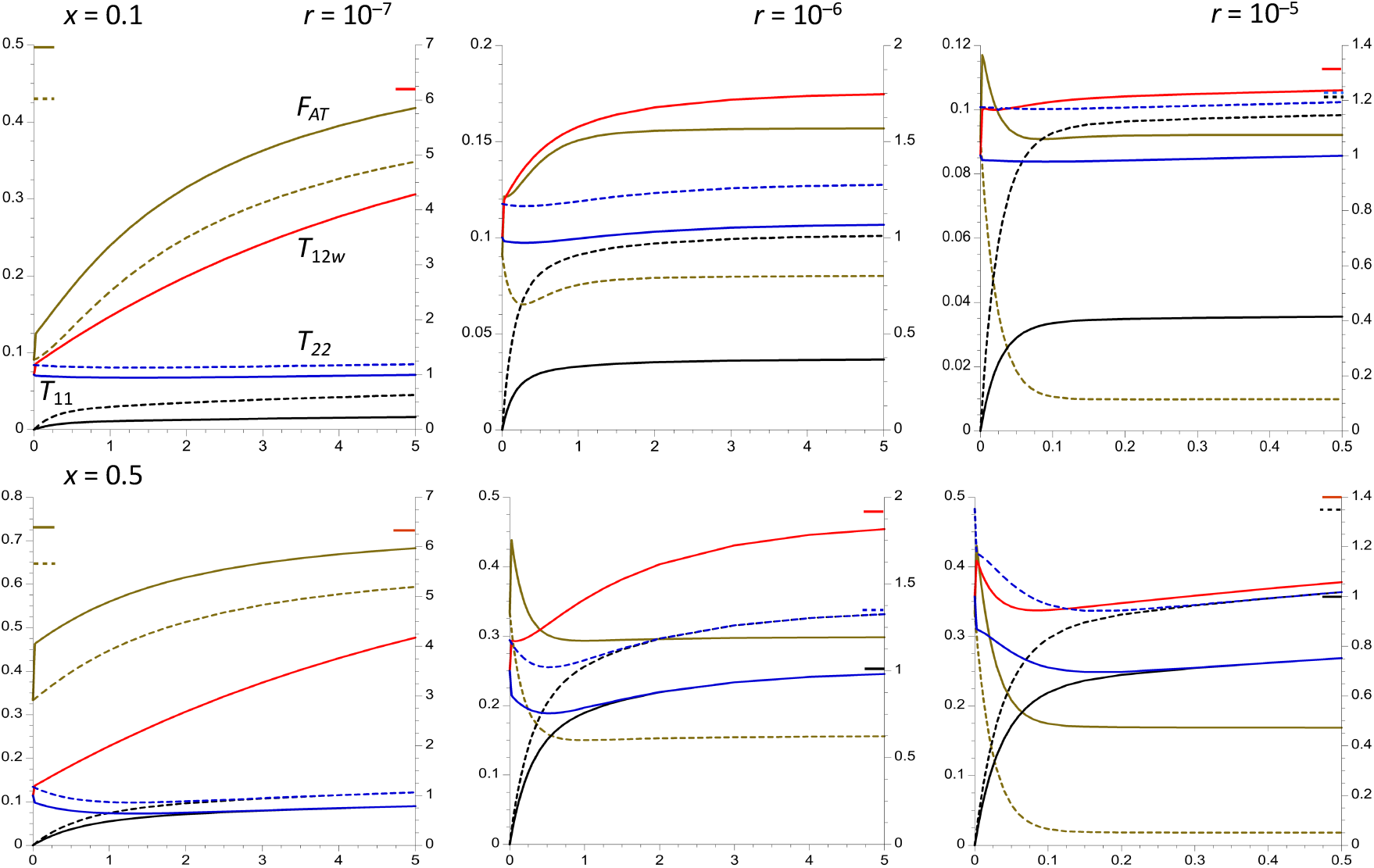
This is the same as Figure 4, except that *F*_*ST*_ = 0.15 (i.e., 4*Nm* = 5.67).

The results are broadly similar to those for the panmictic case described above. The most notable difference is that there is a short initial period with rapid increases in *T*_12*w*_ and *F*_*ATw*_. For *F*_*ATw*_, this period is followed by a monotonic decrease if the equilibrium value of *F*_*ATw*_ is lower than the maximum values it achieves, or (in the case of *x* = 0.1 and *r* = 10^−6^) a decrease followed by an increase) when its equilibrium value exceeds its minimum value. In contrast, *T*_12*b*_ and *F*_*ATb*_ behave initially much like *T*_12_ and *F*_*AT*_ in the panmictic case. For the case of *r* = 0 and a large number of demes (*d*), Equations (S9c) and (S9d) with very small *dMT* show that *T*_12*w*_ ≈ 1 + *dT*, whereas *T*_12*b*_ ≈ (1 + *M*)/*M* + *T*.

The numerical results used in Figs. 4 and 5 show, however, that *T*_12*w*_ and *T*_12*b*_ quickly converge and have nearly identical trajectories after *T* = 0.005 (about 10,000 generations with the parameters used here). It is easily shown from Equations (A2e) and (A2f) for the case of no recombination and an instantaneous sweep of the inversion that:

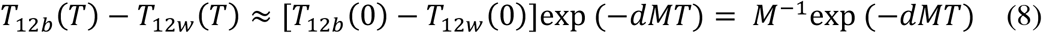

The two measures of *T*_12_ thus converge rapidly when *r* = 0; since it assumed here that migration is a more powerful force than recombination, this is true more generally,

Similarly, Equation (S20e) shows that the coefficient of *T* in the numerator of the expression for *F*_*ATw*_ with no recombination involves *dx*, so that *F*_*ATw*_ also increases very fast initially when the number of demes is large, especially if *x* is close to one. In contrast, there is no contribution from *d* to the expression for *F*_*ATb*_ (Equation S20f); *F*_*ATb*_ starts, however, from a higher initial value than *F*_*ATw*_ because of population subdivision.

When the recombination rate is sufficiently high, Equations (6) and (7) for the initial rates of change in the population statistics can be applied. Δ*T*_12*w*_ in Equation (6e) involves the sum of 1/(2*N*_*T*_) and 1/(2*N*), whereas the term in 1/(2*N*) is absent from the corresponding expressions for Δ*T*_12*b*_ as well as Δ*T*_12_ in the panmictic case. Since 1/(2*N*) is larger than 1/(2*N*_*T*_) by a factor of *d*, this term has a large effect on the initial rate of change of *T*_12*w*_. Similarly, the expression for Δ*F*_*ATw*_ (Equation 7a) involves the positive term (1 – 1/2*d*)/*Ny*, whereas the corresponding expression for Δ*F*_*ATb*_ involves (1 – *F*_*ST*_)/(2*Nd*), which is negative. These results show how it is possible for the two measures to change initially in opposite directions.

## Discussion

### General considerations

The results described here are broadly consistent with previous theory on the patterns of diversity associated with balanced polymorphisms (Kaplan et al. 1988; Hudson and Kaplan 1988; Hudson 1990; Takahata 1990; Charlesworth et al. 1997; Nordborg 1997; Navarro et al. 2000; Charlesworth et al. 2003; Innan and Nordborg 2003; Nordborg and Innan 2003; Guerrero et al. 2012; Kirkpatrick and Guerrero 2014; Zeng et al. 2021), but extend it in several ways. From the purely technical point of view, the recursion relations for mean pairwise coalescence times in a structured population (Nagylaki 1998) provide a simple and computationally efficient method for calculating the expected values of population statistics and their trajectories, on the assumption that the alleles at the target of selection are maintained at constant frequencies. The only previous application of this method to balanced polymorphisms appears to have been that by Zeng et al. (2021) for the case of a single population.

Here this method has been extended to an autosomal balanced polymorphism in a subdivided population, for the simple case of a finite island model of *d* demes of equal size *N*, under a Wright-Fisher model of drift for which the effective population size of a single deme equals *N*. The “migration effective population size” (Nagylaki 1998), which determines the mean coalescent time for a pair of alleles drawn from the same deme at a locus independent of the balanced polymorphism, is then given by *N*_*T*_ = *Nd*, provided that *Nm* is not very close to 0. This result applies more widely to all conservative migration models, where the mean allele frequency across demes is not changed by migration (Nagylaki 1982, 1998). For more general drift models, *N* can be replaced by *N*_*e*_, the effective population size of a deme, given certain restrictions (Charlesworth and Charlesworth 2010, p.327). At recombination-drift equilibrium, the mean coalescent time for this sampling scheme is also equal to the mean within-karyotype coalescent time, if coalescent times for the two arrangements are weighted by the arrangement frequencies (Equation 3) and *N*_*T*_*r* is not very close to 0. For this reason, the various coalescent times used here have been expressed relative to 2*N*_*T*_.

For simplicity, the rest of the discussion will refer only to two arrangements with respect to an inversion polymorphism, but identical results apply to other types of diallelic polymorphisms. In principle, it is possible to extend this approach to more general situations, such as polymorphisms for multiple different arrangements, sex chromosomes, and non-random mating populations, as well as changes in population size. Its main drawback is that the main results are limited to expected pairwise coalescent times, and do not provide information on features such as the site frequency spectra of neutral sites linked to the target of balancing selection. For these properties, either coalescent simulations or more complex analytical approaches, such as the phase-type theory of Zeng et al. (2021), are needed. An extension of Nagylaki’s (1998) recursion equation approach can be used for obtaining the variances and higher moments of pairwise coalescent times, as described in section S8 of the Supplementary Appendix. The statistical properties of mean pairwise diversity and divergence measures over large numbers of nucleotide are discussed in the light of the variances obtained in this way in Section S9.

#### Interpreting population genomic data on inversion polymorphisms

The results on equilibrium expected coalescent times for pairs of alleles (Figures 1, 2, S1 and S2) suggest that the scaled expected coalescent time *T*_12_ for a pair of alleles sampled from the two different arrangements (*In* and *St*) provides the most meaningful basis for interpreting their level of sequence divergence. When the number of demes is large, Equation (A6e) shows that there is essentially no difference in equilibrium *T*_12_ between alleles sampled from the same deme versus alleles from two different demes, at least when *F*_*ST*_ at neutral sites independent of the inversion is moderate, for the reasons described in the section *Numerical results for subdivided populations*.

The numerical results on the time courses of the *T*_*ij*_’s with population subdivision (Tables S4 and S5) show that the lack of dependence of *T*_12_ on the nature of the sample (within or between demes) holds very soon after the equilibrium frequency of the inversion has been reached. In addition, except for the lowest scaled recombination rate considered here (*ρ* = 0.4), the ratio of *T*_12_ to the weighted mean scaled coalescent time within karyotypes and within demes (*T*_*Sw*_) approaches its equilibrium value quite fast, even when both variables deviate considerably from their equilibrium values. For example, with panmixia, *x* = 0.1 and *ρ* = 4, the equilibrium value of *T*_12_ /*T*_*S*_ is 1.5 (*T*_*S*_ = 1), respectively; at time *T* = 0.050, *T*_12_ = 1.039, *T*_*S*_ = 0.916, and *T*_12_/*T*_*S*_ = 1.14; at *T* = 0.5, *T*_12_ = 1.250, *T*_*S*_ = 0.926, and *T*_12_/ *T*_*S*_ = 1.37. With *ρ* = 40, the equilibrium *T*_12_ /*T*_*S*_ is 1.05; at *T* = 0.050, *T*_12_/*T*_*S*_ = 1.039; by *T* = 0.1, *T*_12_/*T*_*S*_ = 1.04. It is, however, not always the case that *T*_12_ /*T*_*S*_ is initially lower than its equilibrium value; for *x* = 0.5 and *ρ* = 4, its initial value is 2 compared with 1.73 at *T* = 0.1, with a final value of 1.55.

These findings are important for the interpretation of population genomic data for several reasons. First, if there is evidence of significant population subdivision at neutral loci independent of the inversion, the measure of sequence divergence between *In* and *St*, estimated by Nei’s *d*_*xy*_ (Nei 1987), is best obtained from the mean of within-population samples, since its expectation is equal to 2*N*_*T*_*T*_12_*u*, and 4*N*_*T*_*u* is equal to the expected within-population neutral *π* in the absence of an inversion polymorphism. Second, *T*_12_/*T*_*Sw*_ (which is equal to *T*_12_ for an equilibrium population) can be estimated by dividing an estimate of *d*_*xy*_ either by an estimate of the mean within-karyotype, within-population diversity (*π*_S*w*_), or by an estimate of the corresponding mean *π* at independent neutral sites (*π*_*nw*_). This is because the expectations of both these statistics are equal to 4*N*_*T*_*u* under the infinite sites model of mutation (for the same mutation rate), unless there is a very low frequency of recombination, or the inversion has arisen very recently. To control for possible differences in mutation rate, it is preferable to use the ratio of an estimate of *d*_*xy*_ to an estimate of *π*_*nw*_ in order to obtain *T*_12_. As discussed above in the section *The model and its analysis*, the theoretical value of the commonly used statistic 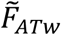, obtained by applying the <*F*_*ST*_> statistic of Hudson, Slatkin and Maddison (1992) to *In* versus *St* haplotypes is equivalent to 1 – *T*_*Sw*_/*T*_12_. However, reciprocals have undesirable statistical properties if means are to be taken over sets of sequences; it is thus preferable to estimate *T*_12_ directly.

This raises the question of whether recombinational exchange in heterokaryotypes occurs at a sufficient rate that the condition *T*_*Sw*_ = 1 is likely to hold. At least for simple inversions (with only a single pair of breakpoints) there is ample experimental evidence for recombination between *In* and *St* at sites within and adjacent to the inversion in species of *Drosophila*, much of which appears to be caused by gene conversion rather than double crossovers(Chovnick 1973; Korunes and Noor 2019; Li et al. 2022; Koury 2023), with *r* = 10^−5^ per basepair per generation in female meioisis being a commonly accepted typical rate for central regions of inversions (Chovnick 1973; Korunes and Noor 2019); for autosomal loci, the lack of recombination in males means that the effective *r* is half of this value. Complex inversions, with three or more breakpoints, might be expected to have much lower rates of exchange than simple inversions, but heterozygotes for multiply inverted chromosomes in *D. melanogaster* have been found to experience non-crossover associated gene conversion events at rates that are even higher than in the absence of inversions (Crown et al. 2018), so that this expectation may not be well founded.

The occurrence of such “gene flux” (Navarro et al. 1997) is consistent with numerous observations of substantial nucleotide site diversity within *In* karyotypes, but at a reduced level compared with genome-wide diversity. In many *Drosophila* examples, *F*_*AT*_ and/or 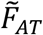 increase and/or within-inversion diversity levels decrease towards the inversion breakpoints (Andolfatto et al. 2001; Nobréga et al. 2008; Corbett-Detig and Hartl 2012; Kapun et al. 2016, 2023), suggesting that flux is lowest near the breakpoints and highest towards the center of the inversion; an exception is the polymorphism in *D. subobscura* for the overlapping pair of inversions O_3+4_ versus the standard arrangement O_st_ (Munté et al. 2005; Papaceit et al. 2012).

Patterns of this kind do not, however, necessarily imply a complete absence of exchange at the breakpoints. Rozas et al. (1999) obtained population genetic evidence for gene conversion events near the breakpoints of arrangements in the O chromosome system of inversions in *D. subobscura*, and Li et al. (2022) found no evidence for an effect of proximity to breakpoints for the *dl-49* inversion of *D. melanogaster* on the rate of gene conversion. There are, however, mechanistic reasons for believing that exchange rates will be lowest at the breakpoints and highest in the center of a simple inversion (Navarro et al. 1997). Population genomic data, which reveal the effects of recombination over very long time spans compared with laboratory crosses, offer an excellent opportunity for exploring the effects of breakpoints on recombination.

#### Estimating the exchange parameter

If equilibrium can be assumed, estimates of between-arrangement divergence and within-arrangement diversities can be compared with the theoretical predictions for the corresponding variables, in order to obtain estimates of the rate of exchange *r* (mutation rate estimates are also needed to estimate *r* rather than *ρ*). Figures 1 and 2 show that, for a low frequency inversion, *T*_11*w*_/*T*_22*w*_, as well as *T*_12_, are strongly affected by the level of population subdivision, unless recombination is rare or absent. Even a modest change from *F*_*ST*_ = 0 to *F*_*ST*_ = 0.05 causes this ratio to shift from close to 0.8 to approximately 0.6 when *r* = 10^−5^ and the inversion frequency is 0.1 (Figure 2), reflecting the effect of population subdivision in multiplying the effective rate of recombination within populations by 1 – *F*_*ST*_ (Wakeley and Lessard 2003). Estimates of *F*_*ST*_ ideally need to be included in any such attempts to estimate *r*. Another difficulty here is that the current model assumes a long-term constancy of the inversion frequency over time, as well as an absence of among-population variation in inversion frequencies between populations. Many studies of *Drosophila* and other groups have revealed major differences among populations in the frequencies of polymorphic inversions, often clinal in nature, which are likely to reflect locally varying selection pressures on genes contained in the inversion, e.g., Krimbas and Powell (1992b), Aulard et al. (2002), Cheng et al. (2012), Kapun and Flatt (2019) and Mérot et al. (2021). Further work is needed to evaluate the consequences of relaxing the assumption of constancy.

This raises the question of whether equilibrium can safely be assumed. Studies of inversion polymorphisms in many *Drosophila* species suggest that they rarely persist beyond species boundaries (Krimbas and Powell 1992a). There are examples of very close relatives with totally different sets of inversion polymorphisms and many fixed differences with respect to gene arrangements, e.g., *D. miranda* versus *D. pseudoobscura* (Bartolomé and Charlesworth 2006). It is thus quite likely *a priori* that the times of origin of many polymorphic inversions are only a small fraction of the mean neutral coalescent time, 2*N*_*T*_, as was proposed by Andolfatto et al. (2001) in their review of early data on the molecular population genetics of *Drosophila* inversions.

Figs. 3-5 show that for the two higher recombination rates 10^−6^ and 10^−5^ corresponding to scaled recombination rates (*ρ*) of 4 and 40, the times for statistics such as *T*_11_, *T*_12_ and *F*_*AT*_ to approach their equilibrium values after the inversion has approached its equilibrium frequency are either commensurate with 2*N*_*T*_ (*r* = 10^−6^) or much smaller (*r* = 10^−5^), consistent with the theoretical prediction that the timescale of approach to equilibrium in terms of generations is of the order of 1/*rx* when *x* < 0.5. both for a panmictic population and a subdivided population with a large number of demes (see section S2 of the Supplementary Appendix). In the absence of recombination, however, diversity within the inversion recovers over a timescale of 2*xN*_*T*_ generations for a panmictic population and 2(1 + *Mx*)*N*_*T*_/*M* generations for a subdivided population (see Equations S5b and S18a), so that the signature of the selective sweep of the inversion on within-karyotype diversity can persist for a long time, as has been noted previously (Navarro et al. 2000; Zeng et al. 2021). In addition, the fact that the ratio *T*_12_/*T*_*Sw*_ converges on its equilibrium value much faster than the absolute *T*_*ij*_’s (see the above section *Interpreting population genomic data on inversion* polymorphisms) means that it is not necessary to assume that the population is very close to equilibrium when using the corresponding divergence to diversity ratio as a statistic.

If recombination in heterokaryotypes is as frequent as is suggested by the data on *Drosophila* (Korunes and Noor 2019), the assumption of recombination-drift equilibrium may thus often be reasonably accurate as a predictor of observed patterns of population genomic statistics for genomic regions covered by an inversion polymorphism, at least for sites that are located well away from inversion breakpoints. The equilibrium assumption can be tested by comparing the within-*In* neutral diversity level with the diversity at neutral sites independent of the inversion (or against the within-*St* diversity). A very recent sweep of the inversion, leaving the system far from equilibrium, would cause diversity across the whole length of the inversions to be much lower than a fraction *x* of the diversity outside the inversion or a fraction *x*/(1 – *x*) of diversity within the standard arrangement – see the curves in Figures 3-5. It would also be expected to leave a signature of an excess of low frequency variants at neutral sites compared with what is seen at comparable sites independent of the inversions (Navarro et al. 2000; Andolfatto et al. 2001; Zeng et al. 2021), although the effects of background selection and selective sweeps within the low recombination environment created by a low-frequency inversion could also cause such a pattern (Charlesworth and Jensen 2021). The inversion *In(1)Be* in African populations of *D. melanogaster* is an example of a very recent spread of an inversion (Corbett-Detig and Hartl 2012).

The study by Kapun et al. of *In(3R)P* in the Zambian population (ZI) of *D. melanogaster* provide an example of how to apply the theoretical results. As mentioned in the section *Numerical results for subdivided populations*, panmixia can be probably be assumed for this case, which implies that Equations (5) can be used for the expected coalescence times. Let *X* = *T*_11_/*T*_22_. Using Equations (5b) and (5c), simple algebra yields following formula for *ρ*, the scaled recombination rate:

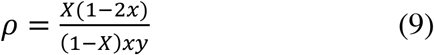

Fig. 2 of Kapun et al. (2023) show that the mean nucleotide site diversity (across all classes of nucleotide sites) is significantly lower (0.00792) for the *In(3R)P* haplotypes than for the standard haplotypes (0.00979), giving an estimate of *T*_11_/*T*_22_ = 0.00792/0.00979 = 0.81 for the region covered by the inversion; *x* ≈ 0.11 for this population (Kapun and Flatt 2019, Table S1). Substituting these numbers into Equation (10) yields an estimate of 34 for *ρ*, corresponding to *r* = 5.3 × 10^−6^ with *N*_*e*_ = 1.6 × 10^6^, the estimate for this population obtained by Johri et al. (2020). close to the estimate of 5 × 10^−6^ from crossing experiments after adjustment for the lack of recombination in males (Korunes and Noor 2019). The estimate of diversity for regions outside the inversion on chromosome 3 is 0.00854, giving an estimate of *T*_11_/*T*_*S*_ = (0.1 × 0.00792 + 0.89 × 0.00979)/0.00854 of 1.12, slightly higher than the theoretical value of 1 for equilibrium; this discrepancy probably reflects different levels of selective constraints among the genome regions being compared and/or the fact that inversions suppress exchange well outside their breakpoints (Li et al. 2022; Koury 2023).

Fig. 4 of Kapun et al. (2023) shows that 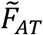 for the central region of *In(3R)P* in the Zambian population approximately 0.1. Equation (3a) implies that:

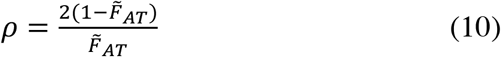

This expression yields an estimate of *ρ* = 18, considerably less than the above value. This may be due to the fact that the estimate of 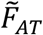 involves taking the mean of a reciprocal over 100kb windows, which biases it toward low values compared with using the mean of estimates of *T*_12_.

The empirical results on *In(3R)P* are thus consistent with this inversion being close to recombination-drift equilibrium, with a mean *r* for central regions of the inversion close to 5 × 10^−6^. The noticeable increase in within-inversion diversity towards the middle of the inversion (Fig. S1 of Kapun et al. 2023), and the increase in 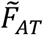 near the breakpoints (their Fig. 4), strongly indicate that flux rates are reduced close to the breakpoints. The analysis of the diversity data near the breakpoints described below suggests, however, that this reduction is not total.

#### Estimating the ages of inversions: divergence between *In* and *St*

If exchange were indeed completely suppressed around inversion breakpoints, divergence between *In* and *St* for sequences close to the breakpoints could be used to estimate the date of origin of an inversion (Hasson and Eanes 1996; Andolfatto et al. 2001; Cáceres et 2001; Corbett-Detig and Hartl 2012). In this case, it can no longer be assumed that *T*_*Sw*_ = 1 for the sequences concerned. Given the reservations about whether such suppression of exchange is absolute, however, caution should be used in making such inferences.

If, as is usually assumed, the new arrangement had a unique origin, *T*_12_ in the absence of exchange is equal to the value for a pair of randomly chosen alleles in the ancestral population. In the case of a panmictic Wright-Fisher population of constant size, *T*_12_ at time *T* since the origin of the inversions is given by 1 + *T* (Equation S5a). With population subdivision, this expression is no longer accurate; if *dMT* >> 1, and *d* is large, *T*_12_ ≈ (1 – *F*_*ST*_)^−1^ + *T* for both within- and between deme measures of *T*_12_ (see the approximation to Equation [S9b]). The first term here corresponds to the mean coalescent time for pairs of alleles sampled from different demes. For species like *D. melanogaster*, with *N*_*T*_ of approximately one million for populations in the ancestral species range (Lack et al. 2015), a *T* value of 0.1 corresponds to 200,000 generations; with *F*_*ST*_ = 0.05, (1 – *F*_*ST*_)^−1^ is approximately 50% of this, so that ignoring this term could create a substantial error in the estimated time of origin. The situation would clearly be worse for a species with a higher level of population subdivision.

Under these conditions, and assuming population size stability, if there is population subdivision, the neutral diversity estimated from pairs of alleles that are independent of the inversion and are sampled randomly across populations (*π*_*nT*_) can be treated as a proxy for the corresponding mean coalescence time, and subtracted from *d*_*xy*_. The ratio of this corrected value to the neutral diversity statistic then provides an estimate of *T* as defined here. For a very recent origin of the inversion, the full Equation (S9b) would usually have to be applied if there is significant population subdivision. The simple approximations of Equations (S9c) and (S9d) could be used when *dMT* << 1, which corresponds to an extremely recent origin when *M* ≥ 1, as is likely to be the case for between-population differences within species of most outcrossing species of animals and plants (Charlesworth 2003; Roux et al. 2016).

In practice, different authors have used different methods for estimating *T* from divergence between arrangements. Taking some of the pioneering studies as examples, Andolfatto et al. (1999) used the number of fixed differences between *In* and *St*, which is strongly affected by sample size. Cáceres et al. (2001), Hasson and Eanes (1996) and Corbett-Detig and Hartl (2012) used *d*_*xy*_ corrected by subtraction of the within-*St* diversity. Somewhat different values will be generated by each of these methods. Difficulties clearly arise if there is evidence for recent strong population expansions or contractions, so that the current population statistics cannot be equated to their values at the time of origin of *In*. There are also likely to be considerable statistical errors, especially as the *d*_*xy*_ values are often very small for *D. melanogaster* inversions (see Table S2 in Corbett-Detig and Hartl [2012]).

#### Estimating the ages of inversions: diversity within *In*

The nucleotide site diversities within arrangements also provide information on the age of an inversion, assuming an absence of exchange. If the inversion frequency is low, however, the within-inversion within-deme diversity (*π*_11*w*_ = 2*N*_*T*_*T*_11*w*_*u*) is only a small fraction of the diversity within the standard sequence or at sites independent of the inversion (see Fig. 1), so that a reduced level of within-inversion diversity does not necessarily imply a recent origin of the inversion. In the case of a subdivided population, provided that *dT* >> *x*(1 + *Mx*), Equation (S18a), which corresponds to the exponential growth model used for simulation-based estimates of inversion age by Corbett-Detig and Hartl (2012), implies that:

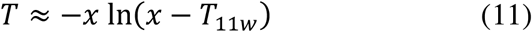

This expression is independent of *M*, and is only meaningful if *T*_11*w*_ < *x*. Given values of *x*, mean diversity within the inversion (*π*_11w_) and *π*_*nw*_ as defined above, it is possible to estimate *T* from this formula, equating *π*_11w_/*π*_*nw*_ to *T*_11*w*_. For example, Table 1 of Andolfatto et al. (1999) provides estimates of diversity values of 0.0009 and 0.0125 for a breakpoint of *In(2L)t* and for independent sites, respectively, in a *D. melanogaster* population with an inversion frequency of 0.23. *T*_11*w*_ is thus estimated as 0.0009/0.0125 = 0.072, giving *T* = 0.42. Andolfatto et al. (1999) estimated the time to the most common recent ancestor of the sequences around the breakpoint from the standard neutral coalescent formula as 0.15 in the present notation; since this method ignores the effect of the recovery from the sweep of the inversion on the gene tree, it overestimates the opportunity for diversity to recover from its complete loss, and hence underestimates the time since the origin of the inversion.

Similar calculations applied to the data in Figs. 2 and S1B of Kapun et al. (2023) on the Zambian population of *D. melanogaster* for sites within 100kb of the distal and proximal breakpoints of *In(3R)P* yield estimates of *T*_11*w*_ of 0.59 and 0.11, respectively. These are highly discordant, reflecting the much lower diversity at the proximal breakpoint, which is also seen in the European and North American samples. Given the estimated frequency of 0.11 for *In(3R)P* in this population (Kapun and Flatt 2019, Table S1), these estimates of *T*_11*w*_ are inconsistent with a total absence of exchange near the breakpoints. The corresponding values of *T*_22*w*_ are 1.28 and 1.13, yielding estimates of *T*_*Sw*_ of 1.20 and 1.02, respectively, so that the data are reasonably consistent with recombination-drift equilibrium for sites close to the breakpoints, although the flux rates must be substantially lower than the mean of 5 × 10^−6^ estimated for the central region of the inversion (see the above section of the Discussion, *Estimating the exchange parameter*).

The numerical results in Supplementary Table S1 for the panmictic case suggest *r* values around 10^−7^ and 10^−6^ for the distal and proximal breakpoints, respectively. Caution should, however, be used in interpreting these conclusions, as they are highly sensitive to the frequency of the inversion in question, and there is considerable continent-wide variation in the frequency of inversions such as *In(3R)P* within Africa (Kapun and Flatt 2019, Table S1); Sprengelmeyer et al. (2020) estimated a frequency of 0.23 for a small Zambian sample. However, the data for Zambian, Bechuanaland and Swaziland samples in Table 1 of Aulard et al. (2002), yield an overall frequency of 0.11 with s.e. of 0.024, so that a frequency around 0.10 for this region of Africa seems reasonable.

An alternative has been to assume that the recent sweep of an inversion results in a star phylogeny, so that the expected pairwise diversity within the inversion (in the absence of exchange) is equal to 2*Tu* (Rogers 1995); Rozas et al. (1999) and Nobréga et al. (2008) used this method to estimate the ages of several inversions of *D. subobscura*. Since a star phylogeny must have the smallest mean divergence between a pair of alleles compared with a gene tree with one or more coalescent events after the origin of the inversion, this method overestimates the age of the inversion.

#### The relevance of *F*_*A*T_ and 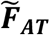

Another point concerns the interpretation of *F*_*AT*_, the analogue of *F*_*ST*_ for differentiation between *In* and *St*), and the related statistic 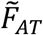; see Equations (4). These statistics are sometimes loosely referred to as measures of the extent of divergence between *In* and *St*, e.g., Guerrero et al. (2012), Kennington and Hoffmann (2013) and Kapun and Flatt (2019). As noted in the section *Theoretical results: Approach to equilibrium, General considerations*, the fact that *F*_*AT*_ and 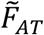 do not necessarily measure the extent of divergence between karyotypes is brought out by their high initial values immediately after an instantaneous sweep to an equilibrium frequency of *x*, for which *F*_*AT*_ = *x*/(1 + *x*), and 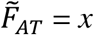. This reflects the fact that *In* and *St* are initially no more divergent on average than a random pair of sequences, but there is no diversity within the inversion. Furthermore, in the presence of recombination *F*_*AT*_ can reach an equilibrium level that is lower than its initial value whereas *T*_12_ increases above 1 (e.g., Figure 4, lower middle panel), so that the two statistics can change in opposite directions over time (see also Zeng et al. 2021, Fig. 8).

As described earlier, *F*_*AT*_ is equivalent to the measure of linkage disequilibrium 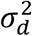 of Ohta and Kimura (1971), which is approximately equal to *R*^2^, the squared correlation coefficient between two loci of Hill and Robertson (1968), so that the use of *F*_*AT*_ is effectively equivalent to estimating linkage disequilibrium (LD) between SNPs and karyotype. This equivalence does not hold for 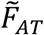, however, which yields much larger values than *F*_*AT*_ for equilibrium situations (Charlesworth 1998; Gammerdinger et al. 2020), and has frequently been used to characterize differentiation between arrangements (e.g., Corbett-Detig and Hartl [2012], Kapun et al. 2023). This difference arises because the theoretical value of 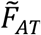is equal to 1 – *T*_*S*_/*T*_12_ whereas *F*_*AT*_ = 1 – *T*_*S*_/*T*_*T*_ with *T*_*T*_ < *T*_12_ at recombination-drift equilibrium (see Equations 4). The difference between them can be considerable; Table 2 of Laayouni et al. (2003) shows both statistics for inversion polymorphisms of *D. buzzatii*, with approximately four-fold larger values of 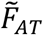 compared with *F*_*AT*_. If data on diversity within inverted and standard haplotypes are available, as well as an estimate of *x*, it is possible to determine *F*_*AT*_ from an estimate of 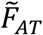 and vice-versa. In the case of *In(3R)P* in the Zambian population of *D. melanogaster*, the mean 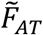 value of 0.1 combined with the diversity values in Figure 2 of Kapun et al. (2023) gives mean *F*_*AT*_ ≈ 0.033, close to a direct estimate (Thomas Flatt and Martin Kapun, personal communication). The magnitude of LD between single nucleotide variants (SNPs) and an inversion polymorphism can therefore be rather small, even when there is noticeable sequence differentiation between arrangements.

The larger expected equilibrium value of 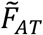 compared with *F*_*AT*_, as well as its closer relationship with the divergence between arrangements, makes it a more powerful measure of the extent of differentiation between arrangements relative to within-arrangement diversity. It is also more convenient for use with population genomic data based on haploid genome sequences, as is the case for much of the *Drosophila* Genome Nexus data (Lack et al. 2015, 2016) and for the data in Kapun et al. (2023), because it does not require the reconstruction of the frequencies of diploid genotypes used for calculating the mean diversity over a random set of individuals. As pointed out above, however, direct estimates of *T*_12_ have better statistical properties.

#### Linkage disequilibrium among neutral variants associated with inversions

As just discussed, estimates of *F*_*AT*_ can be used to estimate LD between neutral variants such as silent site SNPs and an inversion polymorphism. It is also of interest to examine LD between the SNPs themselve, as this has been used to infer the existence of inversion polymorphisms in non-model organisms from evidence for large blocks of LD in specific regions of the genome (e.g., Faria et al. 2019; Mérot et al. 2021). It is difficult to obtain exact analytical results on LD in structured populations (Wakeley and Lessard 2023), but some approximations are easily obtained. From Equation (A9b) of Charlesworth et al. (1997), if LD within arrangements is small compared with LD between neutral sites and the inversion polymorphism, 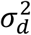 for a pair of SNPS is approximately equal to the product of 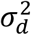 for each SNP and the inversion polymorphism. The standard formula for a partial correlation coefficient implies that this is also true for the *R*^2^ statistic of Hill and Robertson (1968), if there is little or no correlation between SNPs within arrangements. If there is LD within arrangements, these products somewhat underestimate the corresponding statistics for the pairs of SNPs. If there is a major effect of divergence between arrangements on LD between SNPs and karyotype, there should be little dependence of 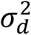 or *R*^2^ on the physical distance between SNPs across the region where crossing over is suppressed by the inversion, in contrast to what is expected for within-karyotype patterns of LD.

Figure S3 of Kapun et al. (2023) shows a pattern of elevated *R*^2^ among SNPs across the entire region of *In*(*3R*)*P* in the Zambian population, contrasted with the rapid decay of LD with physical distance between SNPs within inverted and standard haplotypes. As expected from the product formula, the magnitude of individual *R*^2^ values is modest in this case – the estimate of 0.033 for *F*_*AT*_ given above gives an expected *R*^2^ of approximately 0.001 for a pair of SNPs in genomic regions affected by the inversion polymorphism. Much stronger patterns are, however, seen in the non-African samples studied by Kapun et al. (2023), which probably reflect the effects of population bottlenecks or recent selection. This example illustrates the point that substantial associations between SNPs and an inversion polymorphism may exist, but could be hard to detect simply from LD patterns in samples without prior knowledge of the existence of the inversion polymorphism. Conversely, false signals of an inversion polymorphism could be generated from localized clusters of LD in bottlenecked populations (e.g., Haddrill et al. 2005). Of course, if there are virtually complete associations between SNPs and arrangement, as in the inversion polymorphisms of *Coelopa frigida* (Mérot et al. 2021), there will be a strong pattern of localized LD blocks (see their Fig. 2).

#### Effects of inversions on neutral divergence between populations

For the case of populations with a large number of demes, the extent of population differentiation at a neutral locus associated with an inversion can be measured by *F*_*ST*_ as defined here (which in this case is nearly the same as the < *F*_*ST*_ > statistic of Hudson, Slatkin and Maddison 1992) for standard and inverted haplotypes, denoted here by *F*_*ST, St*_ and *F*_*ST, In*_, respectively – see Equation (S23). Provided that *m* >> *r*, which is likely to be true for sites within the inversion or close to the breakpoints, the equilibrium values of *F*_*ST, In*_ and *F*_*ST, St*_ are equal to 1/(1 + *Mx*) and 1/(1 + *My*), respectively, which are somewhat surprisingly independent of *r* (Equations [A6h] and [A6i]). These expressions are similar in form to that for a subdivided population in the absence of a polymorphism, with *M* replaced by *Mx* and *My*. For *r* = 0, it can be shown that these values are reached almost instantaneously once the inversion has reached its equilibrium frequency (Equation S23). Numerical examples show that this is also true for *r* > 0. The rapid equilibration of *F*_*ST*_ in subdivided populations is well known (Crow and Aoki 1984; Pannell and Charlesworth 2000).

These results imply that within-arrangement *F*_*ST*_ or < *F*_*ST*_ > statistics should be larger for loci within the inversion than for loci that are independent of the inversion, independently of location with respect to the inversion breakpoints. A low-frequency inversion should thus show a considerable inflation in between-populations *F*-statistics, even in the absence of differences in karyotype frequencies between populations, whereas only a small effect should be seen for the standard arrangement; such a pattern was reported by Kennington et al. (2013) for the *D. melanogaster* inversion *In(2L)t*. Caution should therefore be used in interpreting such a pattern as evidence for spatial differences in selection pressures.

In contrast, if inversion frequencies vary considerably between populations because of divergent selection, the LD between neutral sites and karyotype will cause among-population differentiation at neutral sites (Charlesworth et al. 1997; Guerrero et al. 2012), as has been found in some studies of *Drosophila* (Pegueroles et al. 2013) and other taxa (Faria et al. 2019; Mérot et al. 2021). If exchange rates increase with distance from breakpoints, then higher values of *F*_*ST, In*_ and *F*_*ST, St*_ (or the corresponding <*F*_*ST*_> statistics) are expected towards the centers of inversions, as has been observed in some cases, such as Inv2La in *Anopheles gambiae* (Cheng et al. 2012).

## Conclusions

The theoretical results described here provide pointers on how to interpret population genomic data on inversion polymorphisms, with caveats about methods for estimating the ages of inversions. They are subject to several important limitations. In particular, they are based on expectations for pairwise coalescence times. A study of the variances of pairwise diversity and divergence statistics suggests that averages taken over the many thousands of sites in the central regions of several megabase inversions that are several megabases in size have high statistical precision (see sections S8 and S9 of the Supplementary Information), but a rigorous statistical framework such as maximum likelihood inference based on the structured coalescent process (e.g., Lohse et al. 2011) remains to be developed. In addition, many simplifying assumptions have been made, including ignoring the consequences of selection on loci responsible for maintaining the inversion polymorphisms, as well as the effects of Hill-Robertson interference in the low recombination environment characteristic of low frequency inversions. There is, therefore, plenty of scope for further work.

## Data Availability

No new data or reagents were generated by this research. The codes for the computer programs used to produce the results described below will be made available on Figshare on acceptance.

## Acknowledgments

I thank Deborah Charlesworth and Thomas Flatt for helpful discussions, and Thomas Flatt for his comments on a draft of this paper. Aneil Agrawal and two anonymous reviewers made helpful suggestions for improving the paper. Thomas Flatt and Martin Kapun generously shared their results on *In(3R)P*.

## Appendix

### Recursion relations for expected coalescence times

The dynamics of coalescent events are determined jointly by migration, gene conversion and genetic drift. To determine pairwise coalescent times, we need to consider cases where two *In* haplotypes are sampled (a sample of class 1/1), two *St* haplotypes are sampled (a class 2/2 sample), and one of each type are sampled (a class 1/2 sample). The method of Nagylaki (1998) is used for this purpose, which provides a simple way of determining expected coalescent times (see also Charlesworth and Charlesworth 2010, p.316). For 1/1 samples, the probability that the focal neutral site in one of the two haplotypes was associated with *St* in the previous generation is 2*ry* (ignoring second-order terms in *ry*), since the chance that a given *In* haplotype was combined with an *St* haplotype is *y*, and either of the two *In* haplotypes could have experienced a recombination event. There is a probability 2*m* that one of the two haplotypes was present in a different deme from the current one, and a probability of 1/(2*Nx*) that the two haplotypes coalesced in the previous generation if they were both in the same deme, given that haplotypes from different demes cannot coalesce. Similar considerations apply to 2/2 samples, except that *x* and *y* are interchanged. For 1/2 samples, there is a probability *y* that the *In* haplotype was associated with an *St* haplotype in the previous generation, so the probability that the focal neutral site in *In* derives from *St* is *ry*. Similarly, there is a probability *rx* that the focal neutral site in the *St* haplotype derives from *In*; the net probability of the focal site coming from a different karyotype from its current one is *r*. No coalescence of the two haplotypes is possible in this case, but there is a probability 2*m* that one of them has moved from a different deme.

By ignoring second-order terms in *m* and *r*, the following recursion relations for the mean coalescence times defined in the main text can be obtained, with primes indicating their values in the next generation:

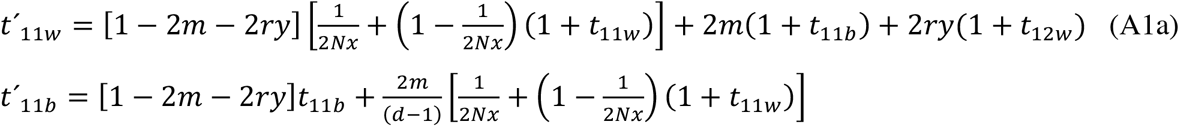

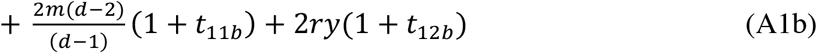

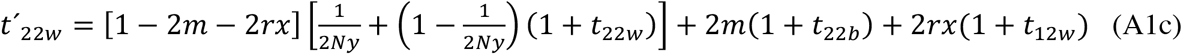

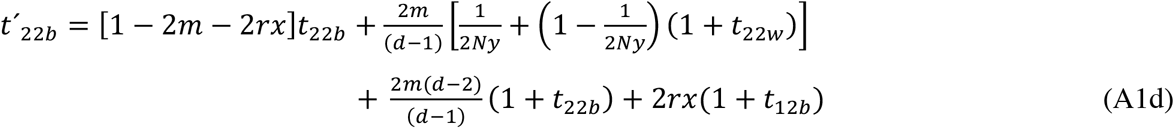

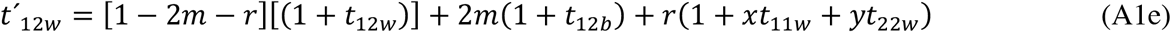

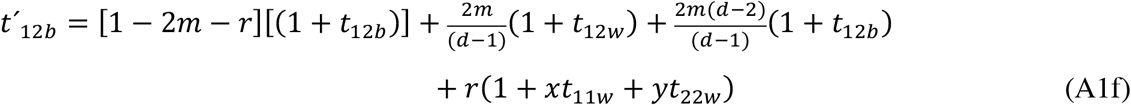

These equations can be simplified by neglecting products of 1*/Nx* and 1/*Ny* with *m* and *r*, and rearranging. This yields the simplified recursion relations:

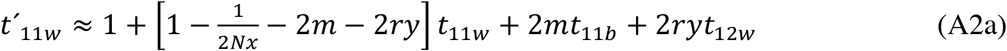

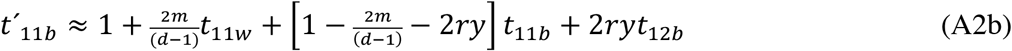

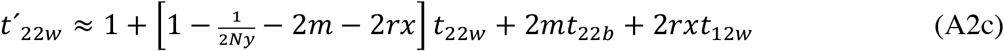

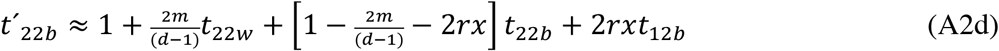

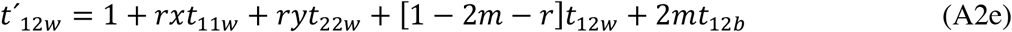

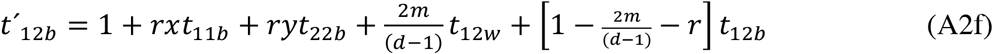

### Expressions for equilibrium mean coalescence times

Writing *M* = 4*Nm* and *R* = 4*Nr*, we obtain the following expressions for equilibrium, equating the *t*′s and *t*s:

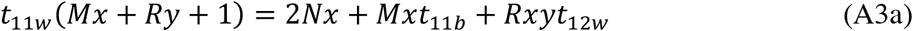

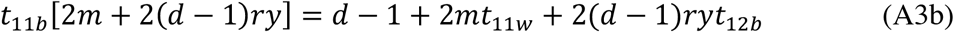

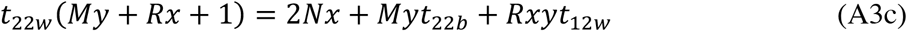

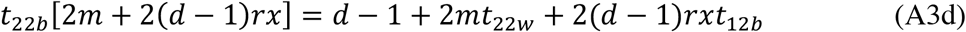

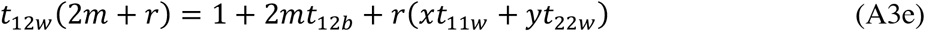

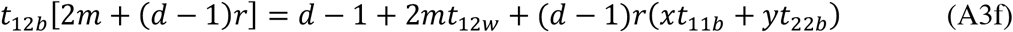

This is a set of 6 linear equations that can be solved exactly (see the Supplementary Appendix, section S1). However, it is possible to obtain informative approximate expressions by assuming that *d* is very large, with 2*m* >> *r* but *dr* >> 2*m*, which are plausible conditions for many real-life cases. This means that the terms in *R* or *r* on the left-hand sides of Equations (A3a), (A3c) and (A3e) can be neglected, as well as the terms in *R* or *r* on the right-hand sides of Equations (A3a), (A3c) and (A3e); in addition, *d* –1 can be replaced by *d* with neglible loss of accuracy.

Using these approximations, we have:

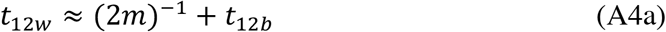

The intuitive interpretation of this expression is that, if recombination is infrequent compared with migration, a class 1/2 pair of alleles will wait an average of 1/(2*m*) generations before one of them migrates to a different deme, after which their expected coalescent time is *t*_12*b*_. The various *t* values are of the order of 2*N*_*T*_ (see below), so that for large *d* and moderate *F*_*ST*_ values, 1/*m* is small compared with the *t*’s, and the term in *m* in this equation can be neglected.

Equations (A3a) and (A3c) can then be approximated as:

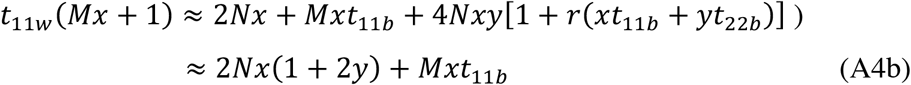

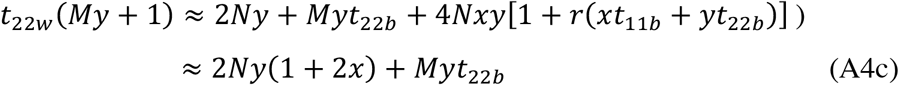

From Equations (A3c) and (A4a) we obtain:

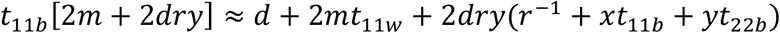

so that:

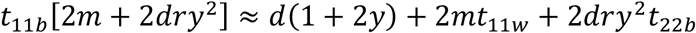

Similarly,

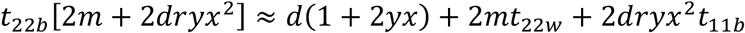

Using Equations (A4b) and (A4c), after some algebra we obtain:

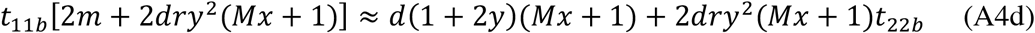

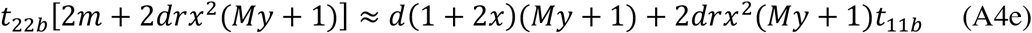

For further work, it is convenient to scale the *t*’s by division by the expected coalescent time for a pair of neutral alleles sampled from the same deme, *T*_*Sn*_ = 2*N*_*T*_ = 2*dN*. Times on this scale are denoted by upper-case *T*’s. The corresponding scaled recombination rate is denoted by *ρ* = 4*N*_*T*_*r*, which replaces 2*r* in Equations (A4d) and (A4e) when using the rescaled *t*’s; 2*m* is also replaced by *dM*, and *d* can be cancelled from both sides of these equations, which then give a pair of linear equations in *T*_11*b*_ and *T*_22*b*_. After some further algebra, their solution can be written in the standard form for a pair of linear equations as:

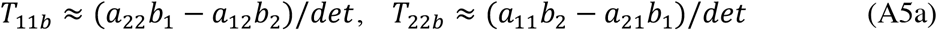

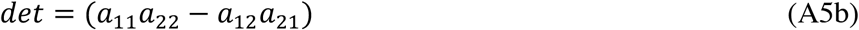

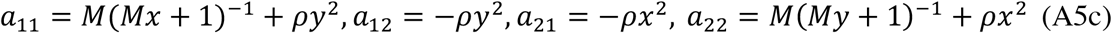

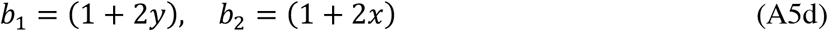

Use of the expressions for the *a*_*ij*_ yields the following formulae:

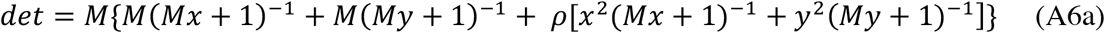

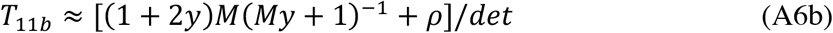

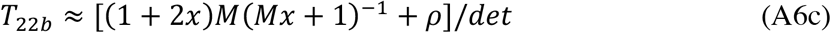

These expressions can be substituted into the scaled versions of Equations (A4d) – (A4e) to obtain the complete approximate solution to the set of equations for the *T*’s:

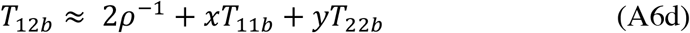

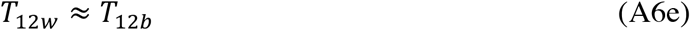

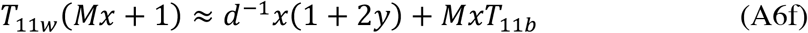

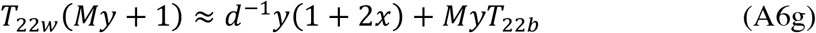

If *d* >> 1, the terms in *d* in Equations (A6f) and (A6g) can be neglected, so that:

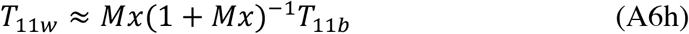

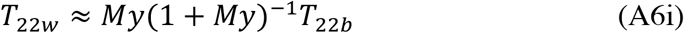

For calculating the approach to the equilibrium described by Equations (A6), it is convenient to eliminate the terms in 1 on the right-hand sides of Equations (A2) by using the vector of deviations of the *T*’s from their equilibrium values, and applying a matrix ***A*** that contains the multipliers of the *T*’s in Equations (A2). Details are given in the Supplementary Appendix.

### Supplementary File S1-Appendix

#### S1 The exact equilibrium solution with population subdivision

The full solution for the equilibrium *T’*s can be found from Equations (A3) as follows, considering each row in turn and after dividing both sides by 2*N*_*T*_.

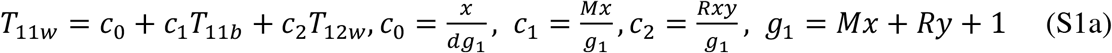

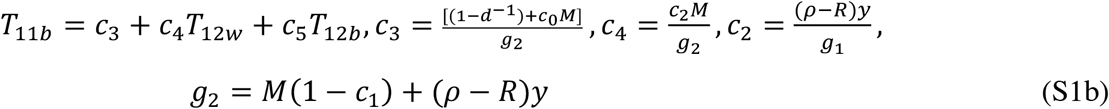

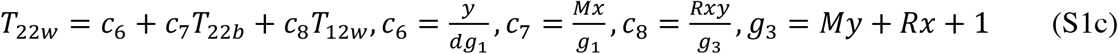

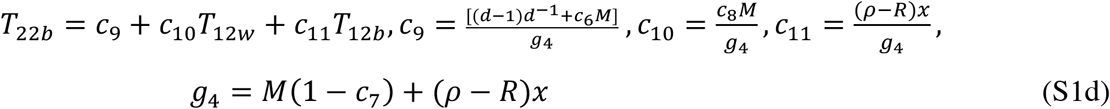

These equations can be used to eliminate all the *T*’s other than *T*_12*w*_ and T_12*b*_ from Equations (A1e) and (A1f), yielding a pair of linear equations:

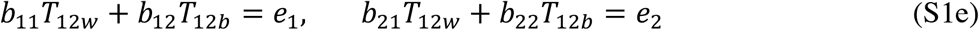

where

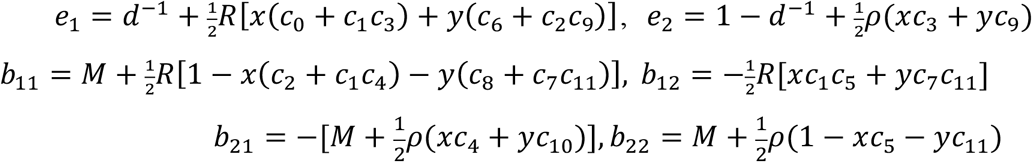

This set of equations has the following standard form solution:

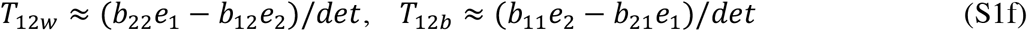

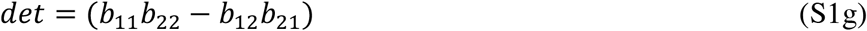

#### S2 The approach to the approximate equilibrium solutions

Let the column vector ***x*** = (*δT*_11*w*_, *δT*_11*b*_, *δT*_22*w*_, *δT*_22*b*_, *δT*_12*w*_, *δT*_12*b*_)^*T*^, where *δ* denotes the deviation from the equilibrium value of the element of the vector in question. Let *N*_1_ = 2*Nx*, and *N*_2_ = 2*Ny*. With large *d*, the 6 × 6 matrix ***A*** that describes the approach of ***x*** to a vector of zero elements, by iterating ***x***_*n*_ = ***Ax***_*n*-1_, consists of the following elements:

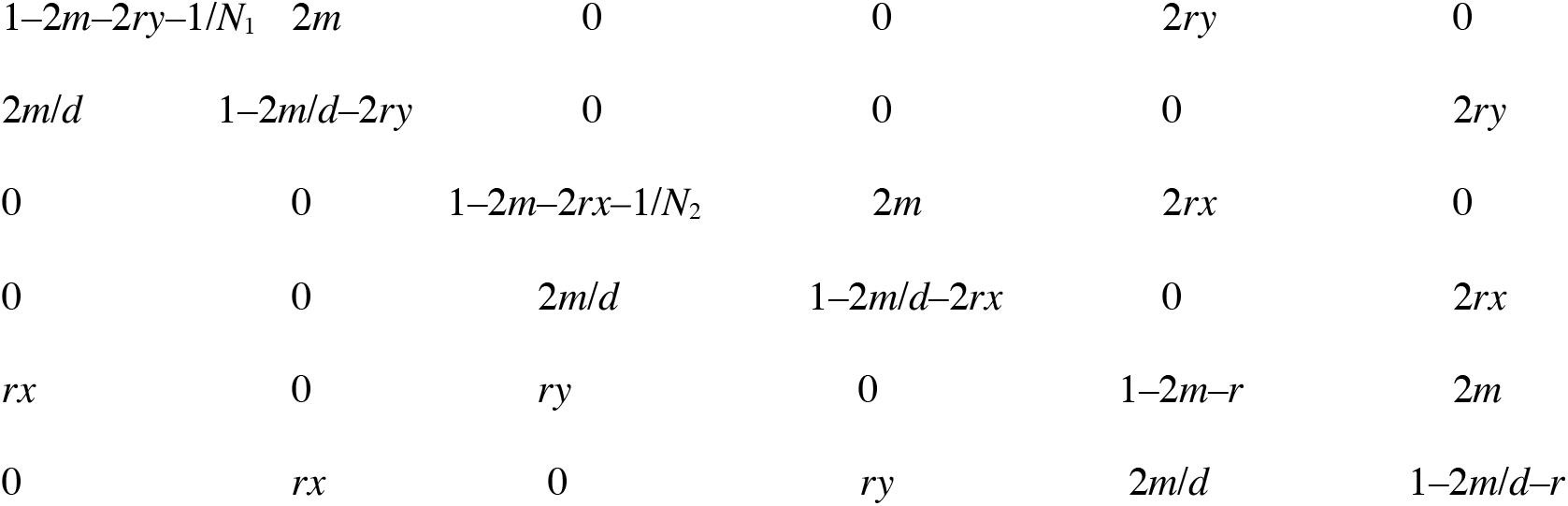

#### S3 Approximations for the eigenvalues and eigenvectors of *A*

An approximation for the set of eigenvalues λ_*i*_ (*i* = 1 – 6) of ***A*** can be obtained as follows, by noting that all its elements *a*_*ij*_ other than the diagonals *a*_*ii*_ are zero or of the order of *m* or *r*, so that second-order terms in non-diagonal elements can be neglected. Ignoring the zero elements in the first row of ***A***, the characteristic equation of ***A***, det(***A*** – λ***I***), can be expanded as (*a*_11_ – λ)*A*_11_ – *a*_12_*A*_12_ + *a*_15_*A*_15_, where *A*_*ij*_ denotes the minor of element *a*_*ij*_.

Similarly, *A*_11_ = (*a*_22_ – λ)*B*_22_ *– a*_34_*B*_34_ *+ a*_35_*B*_35_, where *B*_*ij*_ is the minor of *a*_*ij*_ in the submatrix corresponding to the determinant *A*_11_. Writing *O*(*ε*) for terms of order *m* or *r*, and repeating this procedure on the *B*_*ij*_, it is easily seen that *B*_22_ = (*a*_22_ – λ)(*a*_33_ – λ)(*a*_44_ – λ)(*a*_55_ – λ)(*a*_66_ – λ), *B*_34_ = *O*(*ε*), and *B*_35_ = *O*(*ε*^2^). Since *a*_34_ = *O*(*ε*), the only significant term in (*a*_11_ – λ)*A*_11_ is thus (*a*_11_ – λ)(*a*_22_ – λ)(*a*_33_ – λ)(*a*_44_ – λ)(*a*_55_ – λ)(*a*_66_ – λ).

We also have *A*_12_ = *a*_21_*C*_21_ – *a*_26_*C*_26_, where *C*_*ij*_ is the minor of *a*_*ij*_ in the submatrix corresponding to the determinant *A*_34_. Since both *a*_21_ and *a*_26_ are *O*(*ε*), their product with *a*_12_ can be neglected. Similarly, *A*_15_ can be expanded as *a*_21_*D*_21_ – *a*_22_*D*_22_ + *a*_26_*D*_26_; the products of *a*_21_*D*_21_ and *a*_26_*D*_22_ with *a*_15_ can thus be neglected. *D*_22_ can in turn be expanded as *a*_33_*E*_33_ + *a*_34_*E*_34_ where *E*_33_ = 0 and *E*_34_ = *O*(*ε*).

The final result, therefore, is that det(***A*** – λ***I***) = Π_i_ (*a*_*ii*_ – λ) + *O*(*ε*^2^). This means that the six eigenvalues of ***A*** can be equated to the corresponding diagonal elements, which are of order 1 – *O*(*ε*).

This in turn allows approximations for the right and left eigenvectors of ***A***, and hence its spectral expansion, to be determined, giving a complete (but approximate) analytic solution to the trajectory of deviations of the *t*’s from the equilibrium values. The relevant components of these vectors can be obtained from the equations ***Az***_*i*_ = λ_*i*_***z***_*i*_ and ***y***_*i*_***A***= λ_*i*_***y***_*i*_, where ***z*** and ***y*** refer to right and left eigenvectors, respectively, by successive elimination of elements of the vectors.

Using a similar procedure to that leading to Equation (S1), by considering only the non-zero elements of ***A***, examining each row ***A*** of in succession and writing λ = 1 – *α* for a given λ, we have:

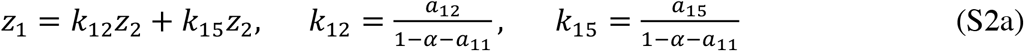

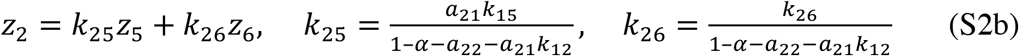

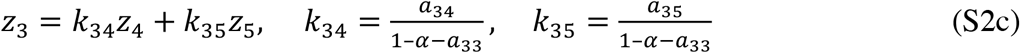

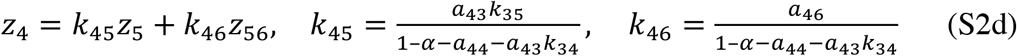

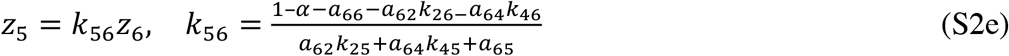

All elements of ***z*** can thus be evaluated explicitly as ratios with respect to *z*_6_. The same procedure can be carried out for the components of the left eigenvector, ***y***; the coefficients corresponding to the *k*_*ij*_ above are identical in form, except that the elements of the transpose of ***A*** are used. Constructing the matrices ***Z*** and ***Y***, whose columns and rows respectively are given by the sets of column vectors ***z***_*i*_ and row vectors ***y***_*i*_ corresponding to each eigenvalue, we have the standard result ***A***^*n*^ = ***ZΛ***^*n*^***Y***, where **λ** is the diagonal matrix whose non-zero elements are equal to the eigenvalues of ***A***, and the eigenvectors are normalized such that the product ***y***_*i*_***z***_*i*_ = 1.

Similar results can be obtained for the panmictic case, using rows and columns 1, 3 and 5 of ***A***, with *m* = 0.

#### S4 Approximations for the initial per generation changes in the *F*_*AT*_’s

The first-order approximation for Δ*F*_*ATw*_ is as follows:

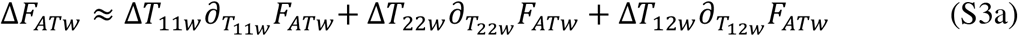

Here, the derivatives are evaluated at the initial conditions: *T*_22*w*_ = *T*_22*b*_ = 0, *T*_12*w*_ = *T*_22*w*_ = 1, and *T*_22*b*_ = *T*_12*b*_ ≈ 1/(1 – *F*_*ST*_). Using the relation *F*_*ATw*_ = 1 – *T*_*Sw*_*/T*_*Tw*_, and the definitions of *T*_*Sw*_ and *T*_*Tw*_ given in the main text (Equations 1 - 4), we obtain the following expressions:

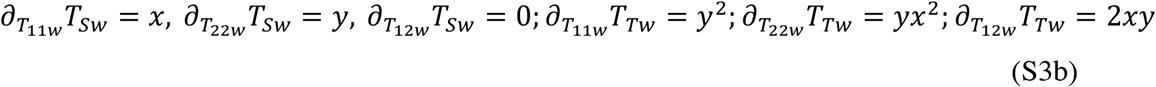

Substituting these relations into Equations (6) and (S3), and using the fact that the derivative of *F*_*ATw*_ = 1 – *T*_*Sw*_/*T*_*Tw*_ with respect to an arbitrary variable *u* is 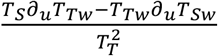, yields Equation (7a), after some simplification.

A similar procedure can be used for *F*_*ATb*_ noting that terms involving the derivative of *T*_22*b*_ can be ignored, because Δ*T*_22*b*_ ≈ 0 by Equation (6d). It is useful to note that the initial value of *T*_*Tb*_ can be written as (1 − *F*_*ST*_)^−1^ ***y***[1 + ***x***(1 − *F*_*ST*_)]. We have:

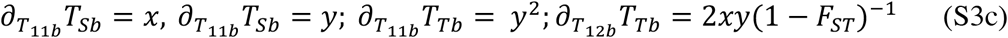

This yields Equation (7b).

#### S5 Inaccuracy in the Approximate Expressions for Equilibrium

Denote the true equilibrium vector of *T’s* by 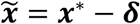, where ***x***\* is the approximate value. Let the true value of ***x*** in generation *n* be ***x***_*n*_, where *n* = 0 for the initial generation under consideration. Let ***A*** be the true value of the recursion matrix discussed above. The true equilibrium satisfies 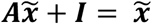, where ***I*** is the unit matrix. For generation 1, we can write:

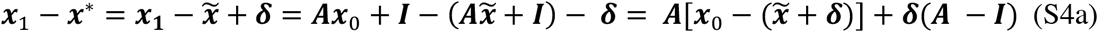

In the next generation,

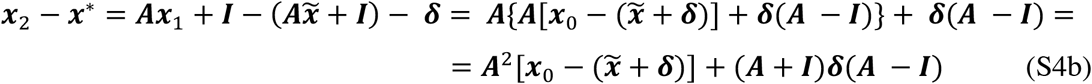

Continuing with the iteration, the general expression is:

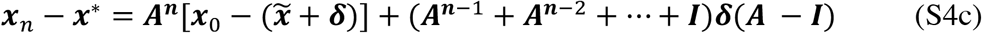

From the analysis in section S3, the term in ***A***^*n*^ must tend to zero as *n* increases, at least as far as first-order terms are concerned, so only the series in ***A*** need be considered. If we write ***P***_*i*_ for the matrix formed by the vector product ***z***_***i***_***y***_***i***_, such that 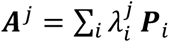, for large *n* this term approaches 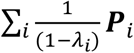, provided that |***λ***_S_| < 1, as must be the case from the results in section 3. This implies that the iteration of ***x*** – ***x***\* by the matrix ***A***, as used in the numerical studies of the approach to equilibrium will yield a final value whose magnitude is of the same order as the error ***δ*** in the approximate equilibrium. If this value is small, the error involved in the approximation must also be small.

#### S6 Time courses of changes in coalescent times in the absence of recombination in a panmictic population

In this case, there is no distinction between within- and between-deme coalescence times, and the subscripts *w* and *b* can be dropped. The mean between-karyotype coalescent time after a time *T* has elapsed (where both quantities are scaled relative to the neutral coalescence time 2*N*_*T*_), is simply a linear function of *T*:

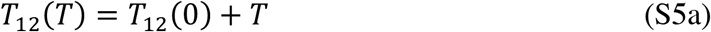

The mean coalescence times within-karyotypes obey the rule for the approach to their equilibrium values described in section S2 above. In this case, the relevant expressions simplify to:

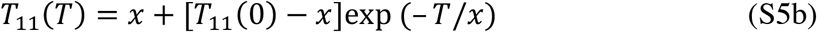

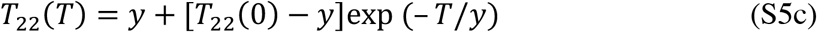

If the sweep of the inversion to its equilibrium frequency is assumed to be instantaneous, we have *T*_11_(0) = 0 and *T*_12_(0) = *T*_22_(0) = 1; we also have *T*_*T*_ (0) = 2*xy* + *y*^2^ = *y*(1 + *x*), so that *F*_*AT*_(0) = *x*/(1 + *x*). For arbitrary *T, T*_*S*_(*T*) = ***x***^2^[1 − exp(−*T*/***x***)] + ***y***[***y*** + ***x*** exp(−*T*/***y***)] and *T*_*T*_(*T*) = 2***xy***(1 + *T*) + ***x***^2^[1 − exp(−*T*/***x***)]+***y***^2^[***y*** + ***x*** exp(−*T*/***y***)].

These relations can be used to determine the time-course of *F*_*AT*_ following an instantaneous sweep, using the relation:

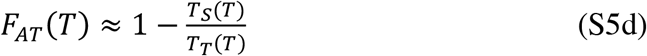

For sufficiently large *T*, the exponential terms in the above equations can be neglected, and *T*_*S*_*(T)/ T*_*T*_*(T*) ≈ (1 – 2*xy*)/(2*xyT*). This implies that, as expected, *F*_*AT*_ always increases towards 1 once a large *T* value has been reached. For small *T*, the first-order approximations exp(–*z*) ≈ 1 – *z* and 1/(1 + *T*) ≈ 1 – *T* yield the following formulae:

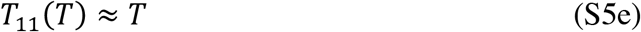

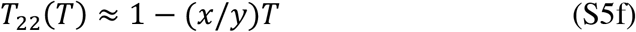

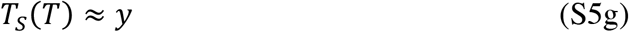

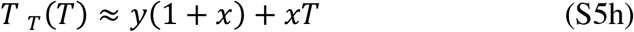

Substituting these approximations into Equation (S5d) and simplifying, we have:

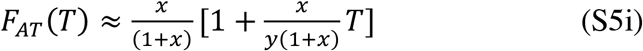

The multiplicand of *T* is always positive for 0 < *x* < 0, implying that *F*_*AT*_ always increases initially with time.

#### S7 Time courses of changes in coalescent times in the absence of recombination in a subdivided population

With no recombination, the recursion relations for *T*_12*w*_ and *T*_12*b*_, which can be obtained from Equations (A3) reduce to:

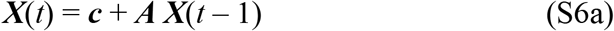

where ***X***(*t*) is the column vector with elements *T*_12*w*_(*t*) and *T*_12*b*_(*t*); ***c*** is the column vector whose elements are both equal to 1/(2*N*_*T*_); ***A*** is the 2 × 2 matrix with elements *a*_11_= 1 – 2*m, a*_12_ = 1 – 2*m, a*_21_ = 2*m/*(*d* – 1) and *a*_22_ = 1 – 2*m/*(*d* – 1).

Iteration of this equation yields the following expression:

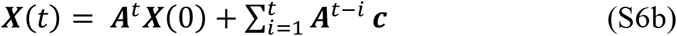

The powers of ***A*** can be evaluated using the spectral expansion of ***A*** in terms of its eigenvalues and eigenvectors. The eigenvalues of ***A*** satisfy the following characteristic equation:

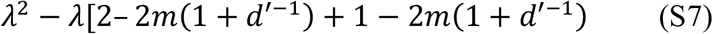

where for convenience *d* – 1 is denoted by *d′*.

This equation has roots λ_0_ = 1 and λ_1_ = 1 – 2*m*(1 + *d′*^−1^), with corresponding right and left eigenvectors ***u***_*i*_ and ***v***_*i*_, respectively (*i* = 0, 1). Use of the identities ***Au***_*i*_ = *λ*_*i*_***u***_*i*_ and ***v***_*i*_***A*** = *λ*_*i*_***v***_*i*_ give the result *u*_01_ = *u*_02_ for the 1^st^ and 2^nd^ elements of ***u***_0_, respectively. Similarly, *u*_11_ = (–*d′*) *u*_12_, *v*_01_ = *v*_02_*/d* and *v*_11_ = –*v*_12_. In order to use these in the spectral expansion of ***A***, we arbitrarily set *u*_01_ = *u*_02_ = 1 and *u*_11_ = 1, with *u*_12_ = – *d′*^−1^, and use the normalization requirement 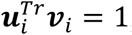, where the superscript *Tr* denotes transposing a column vector into a row vector. This gives *v*_01_ = (1 + *d′*)^−1^, *v*_02_ = *v*_11_ = (1 + *d′*^−1^)^−1^, *v*_12_ = – (1 + *d′*^−1^)^−1^. Using the formula for the spectral decomposition of a matrix, ***A*** = ∑_*i*_ ***λ***_*i*_***u***_*i*_ ***υ***_*i*_, we obtain:

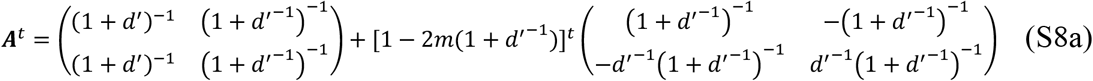

For large *d*, this expression reduces to:

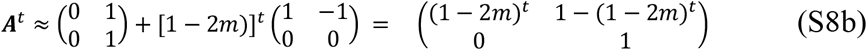

This equation implies that:

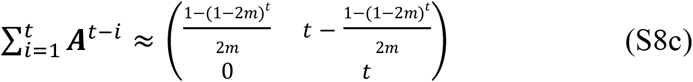

If time is transformed to the coalescent timescale, *T* = *t*/(2*N*_*T*_), (1 – 2*m*)^*t*^ can be approximated by exp(– *MdT*). On the assumption of an instantaneous approach of the inversion to its equilibrium frequency, so that the initial vector ***X***(0) = (1, (1 – *F*_*ST*_)^−1^)^*Tr*^ = (1, (1+*M*)/*M*)^*Tr*^ Equations (S6b), (S8b) and (S8c) yield the following expression:

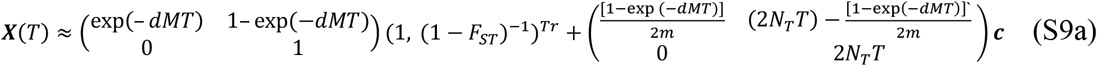

which reduces to:

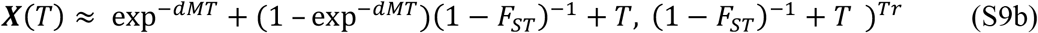

When *dMT* >> 1, this expression implies that both *T*_12*w*_ and *T*_12b_ ≈ 1/(1 – *F*_*ST*_) + *T* ≈ (1+*M*)/*M* + *T*. For *dMT* << 1, *T*_12*w*_ increases faster with *T* than does *T*_12*b*_, which always increases in direct proportion to *T*, especially when *M* is small. This can be seen by using the first-order approximation exp(– *z*) ≈ 1 – *z* in Equation (S9b), which yields:

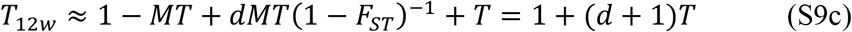

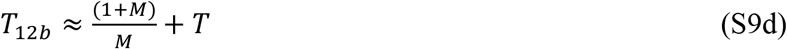

The deviations of *T*_11*w*_(*t*) and *T*_11*b*_(*t*) from their equilibrium values, *T*_11*w*_ = *x* and *T*_11 *b*_ = *x*/(1 – *F*_*ST, In*_), where *F*_*ST, In*_ = is the value of *F*_*ST*_ for the inversion subpopulation are described by the following matrix equation:

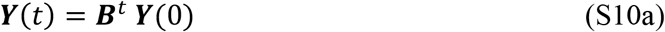

where ***Y***(*t*) is the column vector (*δT*_11*w*_(*t*), *δT*_11b_(*t*))^*Tr*^. We have:

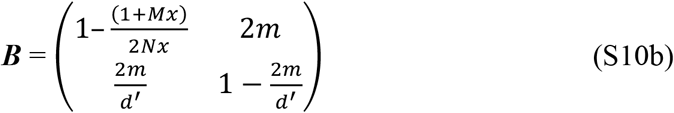

The characteristic equation of ***B*** can be written as:

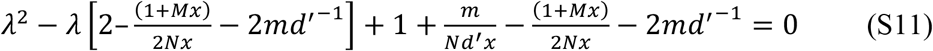

This equation has the following roots:

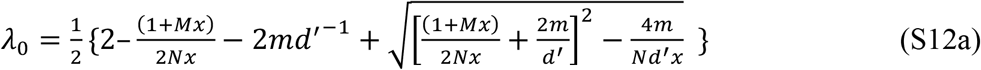

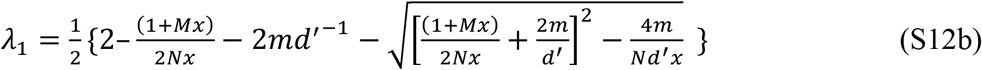

The quantity inside the square term can be written as:

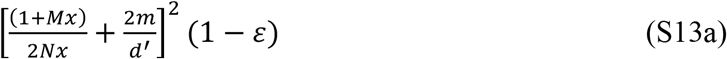

where

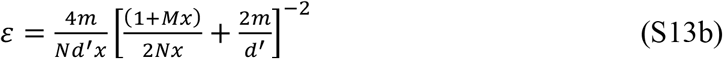

Unless *Nx* is close to 1, *ε* << 1, and 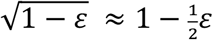, so that the eigenvalues can be approximated by:

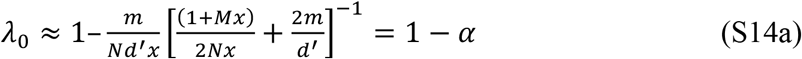

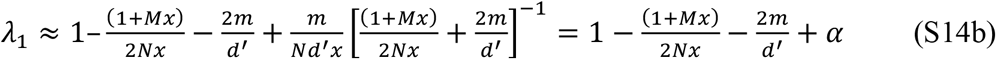

where

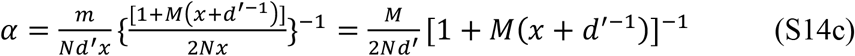

For large *d*, we have:

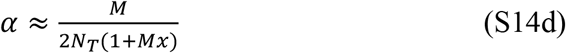

This expression implies that the asymptotic rate of approach to equilibrium on the coalescent timescale of 2*N*_*T*_ generations is approximately *M*/(1 + *Mx*).

The elements of the corresponding right and left eigenvectors, ***u***′_*i*_ and ***v***′_*i*_, are given by the following expressions:

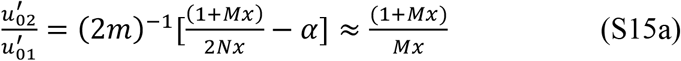

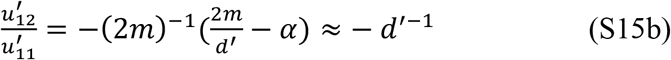

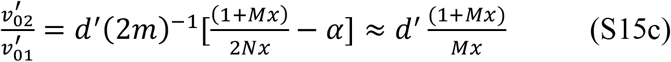

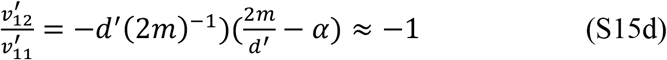

As before, we can arbitrarily set 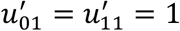, and use the normalizations 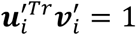 to obtain the following relations:

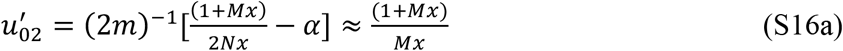

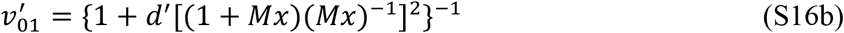

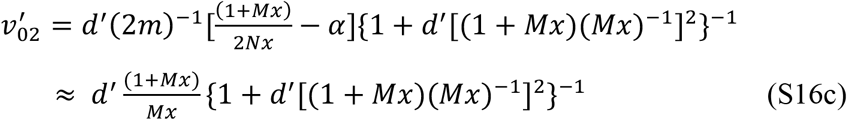

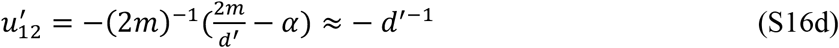

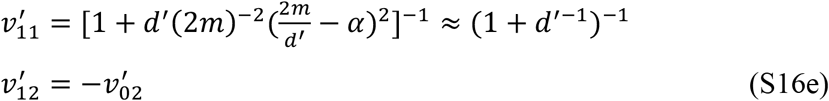

The spectral decomposition of the power of a matrix can be used to write 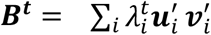. This can be substituted into Equation (S10a) to obtain *T*_11*w*_ (*t*) and *T*_11*b*_ (*t*). When *d* is large, Equations (S16) yield:

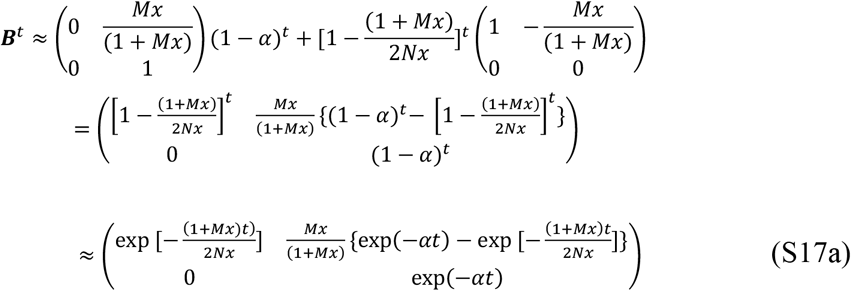

If time is measured in units of coalescent time (*T*), the last expression can be written as:

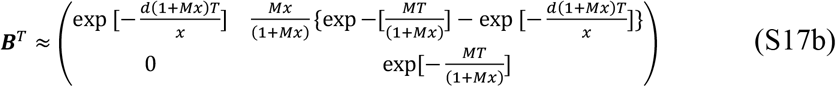

The terms involving exp [– *d*(1 + *Mx*)*T*/*x*] decay much faster than those involving exp [– *MT*/(1 + *Mx*)], unless *d*/*x* is of order l or less, and can therefore be neglected when *d* is large, except when *T* is of order 1/*d*, i.e., for early stages in the process. For large *d*, the equilibrium values of *T*_11*w*_ and *T*_11b_ are *x* and approximately (1 + *Mx*)/*M*, respectively; Equation (S17b) implies that, for *dT* >> *x*/(1 + *Mx*), we have:

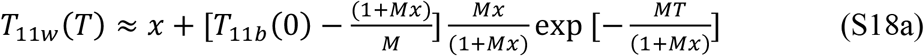

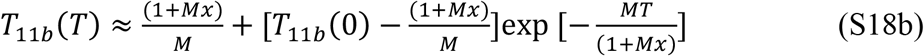

where *T*_11*w*_(0) = *T*_11*b*_(0) = 0 in the case of an instantaneous approach of the inversion to its equilibrium frequency. In this case, Equation (S18a) has the same form as Equation (S5b) for the panmictic case, except that (1 + *Mx*)/*M* replaces *x* in the exponential function. This corresponds to the replacement of the migration effective population size, *N*_*T*_, which determines the expected coalescent time for pairs of allele sampled within a deme, by the “total effective population size”, *N*_*T*_/(1 – *F*_*ST*_), which determines the expected coalescence time for pairs of alleles sample randomly from the whole population {Charlesworth, 2010 #3373}, p.318. Since *x* < (1 + *Mx*)/*M*, this corresponds to a slower asymptotic rate of approach to equilibrium in the case of a subdivided population, with the rate being an increasing function of *M*.

A similar treatment of changes in mean coalescent times can be applied to the *St* subpopulation by replacing *T*_11*w*_(*t*) and *T*_11*b*_(*T*) with *T*_22*w*_(*T*) and *T*_22*b*_(*T*), and *x* with *y*, setting *T*_22*w*_(0) = 1 and *T*_11*b*_(0) = (1 + *My*)/*M* in the case of an instantaneous approach of the inversion to its equilibrium frequency.

The relevance of these asymptotic rates of change is not, however, clear, since the rates of change caused by the smaller eigenvalue are substantial when *d* is large, as assumed here, so that the equilibrium values of the *T*_*iiw*_ and *T*_*iib*_ (*i* = 1 or 2) may be approached quite early on. The properties of the early stages of the process can be examined by approximating the exponentials in Equation (S17), and its equivalent for *T*_22*w*_ and *T*_22*b*_, by the first-order terms in the relevant exponential functions, as was done to obtain Equations (S5) for the panmictic case in section S5. But it is important to note that this approximation requires *dT*(1+*Mx*)/*x* << 1in Equation (S17b), so that it will be accurate only for *T* << *x*/*d*(1+*Mx*), a brief period of time.

Applying the initial conditions *δT*_11*w*_(0) = – *x, δT*_11*b*_(0) = – (1 + *Mx*)/*M, δT*_22*w*_(0) = *x* and *δT*_22*b*_(0) = (1+*M*)*M*^−1^ – (1+*My*)*M*^−1^ = *x* to these equations, some simple algebra yields the following expressions for the case of an instantaneous sweep:

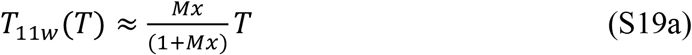

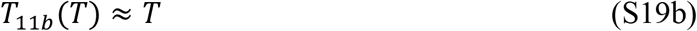

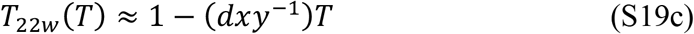

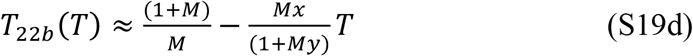

Equation (S19b) shows that *T*_11*b*_ is initially proportional to *T*, and Equation (S19a) shows that *T*_11*w*_ increases with time at a rate that is a fraction *Mx*/(1 + *Mx*) of *T*. Equation (S19c) shows that *T*_22*w*_ decreases rapidly with time when *d* is large, due to the dominance of the relevant term in *d* in the relevant recursion equation; this means that the decline in *T*_22*w*_ dominates the expressions for *T*_*Sw*_ and *T*_*Tw*_. These expressions yield the following results for the mean within-karyotype and overall coalescent times, assuming that *d* is large and *dMT* << 1:

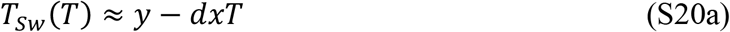

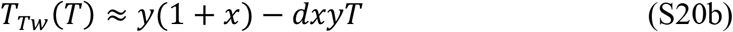

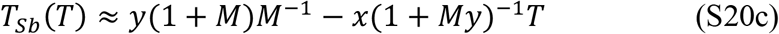

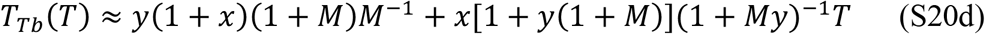

Combined with Equations (S9c) and (S9d), these expressions can be used to obtain approximations for *F*_*ATw*_ and *F*_*ATb*_ as functions of *T* when *dT* is small and *d* is large. After some algebra, the following expressions are obtained:

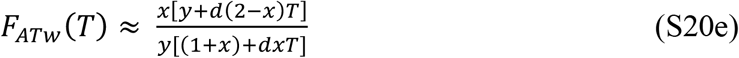

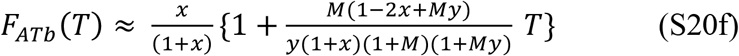

These equations imply that *F*_*ATw*_ always increases initially when there is no recombination, for the case of an instantaneous sweep; *F*_*ATb*_ also increases initially, provided that *y* > (2*x*–1)/*M*, a fairly light condition

It is also of interest to ask how the between-population measure of differentiation for the *In* subpopulation behaves in the absence of recombination. This can be measured by considering the within- and between-population mean coalescence times for pairs of randomly sampled haploid genomes that both carry the inversion, giving *F*_*ST,In*_ = 1 – *T*_11*w*_/*T*_11*b*_, given that *d* is assumed here to be very large. The general equations for coalescence times then imply that:

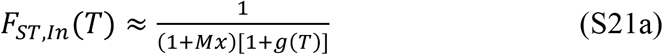

where:

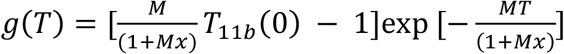

Similarly,

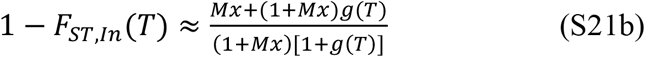

and

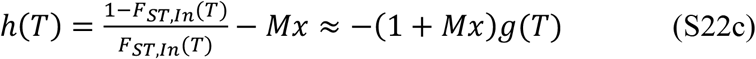

Equation (S22c) implies that:

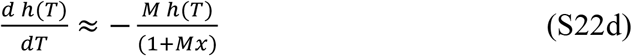

i.e. the asymptotic proportional rate of change of the natural logarithm of the deviation of the ratio of 1 – *F*_*ST,In*_ to *F*_*ST,In*_ from its equilibrium value of 1/(1 + *Mx*) is equal to – *M*/(1 + *Mx*), which is the same as the asymptotic proportional rates of change of *δT*_11w_(*T*) and *δT*_11b_(*T*).

As in the case of *F*_*AT*_, the relevance of these asymptotic rates of change is unclear. For small *T*, following the same approach as used above for *F*_*AT*_, we have:

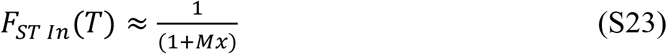

with the somewhat surprising result that *F*_*ST,In*_ is constant and equal to its equilibrium value, to the order of the approximations used here, despite the fact that it is undefined when *T*_11*b*_ = 0. This arises from the fact that Equation (S19a) shows that *T*_11*w*_ for small *T* ≈ *MxT*/(1+*Mx*) whereas Equation (S19b) shows that *T*_12*b*_ is proportional to *T*, so that 1 – *T*_11*w*_ / *T*_11*b*_ ≈ 1/(1 + *Mx*).

Another way of obtaining this result is to note that, if *α* in Equation (S17a) is neglected (as is reasonable for small *t*), the rate of change per generation in *T*_11*w*_, given by ***B***, is as follows:

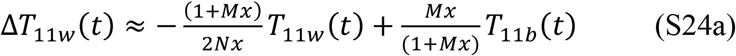

This expression is equal to zero when:

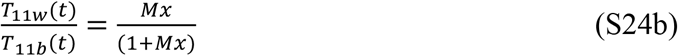

which is equivalent to Equation (S23). The rate of change per generation in *T*_11*w*_ is initially very small, so that *T*_11*w*_ reaches a quasi-equilibrium such that its ratio with respect to *T*_11*b*_, and hence *F*_*ST,In*_, is approximately constant over time.

A similar treatment of *F*_*ST*_ can be applied to the *St* subpopulation by replacing *T*_11*w*_(*t*) and *T*_11*b*_(*T*) with *T*_22*w*_(*T*) and *T*_22*b*_(*T*), and *x* with *y*.

#### S8 Recursion equations for the second moments of the *T*_*ij*_ in a single population

The method for obtaining recursion relations for the expected coalescent times (Equations A1) can be extended to their higher moments in a conceptually straightforward way by replacing the (1 + *t*_*ij*._) in Equations (A1) by (1 + *t*_*ij*._)^*k*^, where the. subscript is either *w* or *b*., and expanding by the binomial theorem. Given expressions for *k* = 1 and 2, recursions for *k* = 3, 4, etc. can be developed.

For simplicity, only the case of *k* = 2 for a single randomly mating population of size *N* will be considered in detail here, so that the subscripts *w* and *b* can be dropped, allowing determination of the variances of the *t*_*ij*_. This yields the following recursion relations, where *t*_*ij*_ and 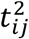 denotes the expectations of the relevant coalescent time and its square, respectively:

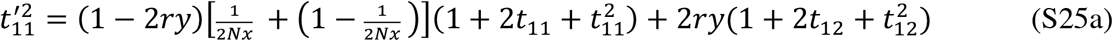

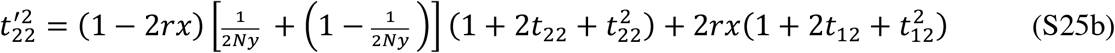

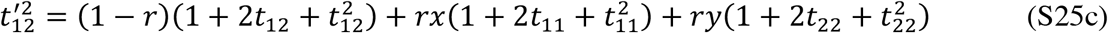

These equations can be iterated from a specified set of initial conditions in conjunction with the corresponding recursion equations for the expectations of the *t*_*ij*_ (see Equations A1 and A2) providing exact expressions for the first and second moments of the *t*_*ij*_. Useful approximations for the equilibrium values of the second moments can be obtained by noting that the expected squares of the equilibrium coalescent times are much greater than than the expected values of the equilibrium coalescent times themselves, so that many of the terms in Equations (S25) can then be neglected. At equilibrium, this procedure yields:

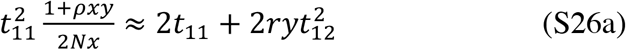

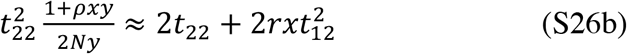

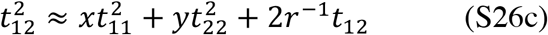

Transforming to the coalescent time-scale of 2*N* generations, these equations become:

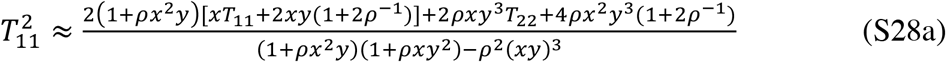

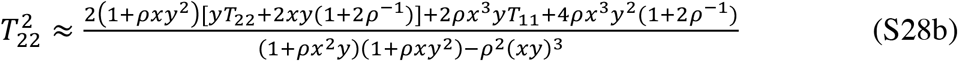

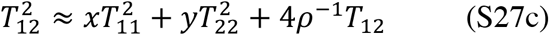

where the *T*_*ij*_ are given by Equations (5)

After some algebra, we obtain the following expressions:

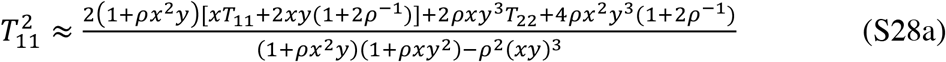

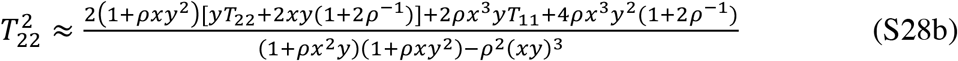

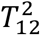 can be obtained by substituting Equations (S28) into Equation (S27c). The variances and standard deviations of the three coalescent times are then easily found from the corresponding first and second moments.

This method of determining the variances of the pairwise coalescence times can be validated by comparing the standard deviations generated by Equations (5) and (S28) with the results of coalescent simulations when the time of sampling is sufficiently large that the process is close to equilibrium. Results for a wide range of *ρ* values with *x* = 0.1 and 0.5 are shown in Table S2, and display excellent agreement between the analytical and simulation results.

#### S9 Statistical error in population genomic estimates of diversity and divergence for polymorphic inversions

Tajima (1983, Equation 30) showed that, for a Wright-Fisher population of size *N* and a sample size of *n*, the stochasticity of coalescent times generates a variance in *π* at a single nucleotide site of:

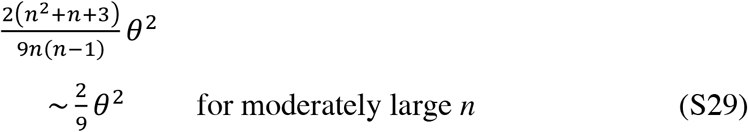

where *θ* is the scaled mutation rate 4*Nu*. In other words, the coefficient of variation (CV) of *π* due to the coalescent process is only somewhat smaller than 1, as would be expected from the exponential distribution of coalescent times in the Kingman coalescent process, and the strong, but not complete, correlations between the coalescence times of different pairs of alleles in the same sample. There is a also a variance of (*n* +1)*θ*/3(*n* – 1) arising from the mutational process itself. For *n* = 2, the variance is simply *θ* + *θ* ^2^.

In general, the variances of the means of pairwise diversity and divergence statistics over large numbers of nucleotide sites, which are discussed in the main text, depend on the extent to which different sites have correlated genealogies; a mean over *m* totally independent sites has a variance of 1/*m* times the variance for a single site. Most parts of the genome of eukaryote taxa such as *Drosophila* experience significant rates of recombination, so that different sites do not have totally correlated gene genealogies (McVean 2002). In normally recombining regions of the *Drosophila* genome, linkage disequilibrium (LD), as measured by the squared correlation coefficient between pairs of sites (*R*^2^), approaches the baseline level generated by finite sample size once they are a few hundred basepairs apart (e.g., Charlesworth & Charlesworth 2010, p.384). This suggests that the majority of pairs of sites in a window of 100kb, as in the Kapun et al. (2023) dataset on *In(3R)P* of *D. melanogaster*, can be treated as having independent gene trees, given that *R*^2^ is closely related to the covariances of the lengths of gene trees (McVean 2002). Thus, the stochastic coalescence variances of mean pairwise diversity and divergenc statistics for a sequence of *m* basepairs over a single window in most regions of the *Drosophila* genome are likely to be multiplied by a factor of the order of 1/*m* compared to the single nucleotide site values.

To apply this reasoning to population genomic data on inversion polymorphisms, it is first necessary to determine the variances of the diversity and divergence statistics for a single site when the division of the population into *In* and *St* haplotypes plus any population structuring is taken into account. Each class of event in a structured coalescent process (e.g., coalescence, migration to a new deme) has an exponentially distributed waiting time. However, the distribution along a chain of such events involves convolutions of the distributions of the individual events, and hence is not itself exponential.

In the case of *In(3R)P*, population subdivision can safely be neglected, so that the results in section S8 can be applied. The results in Table S2 show that *T*_11_ has the largest coefficient of variation among the *T*_*ij*_, but this is less than 5 for the lowest scaled rate of recombination (*ρ* = 0.4) between *In* and *St* with *x* = 0.1 and less than 2.5 with *x* = 0.5. For *ρ* = 4 or 40, which the analyses in the main text show are consistent with the *Drosophila* data, the standard deviations of all of the *T*_*ij*_ are close to their means.

It is also necessary to determine whether LD decays sufficiently fast within inversion haplotypes for the argument about the variance of mean diversity and divergence statistics to be valid. Kapun et al. (2023, Fig. 3) give detailed information for the *In(3R)P* inversion of *D. melanogaster*. For out-of-Africa populations, bottleneck effects have created substantial LD within both standard and inverted haplotypes, so only their panel A for the Zambian population (where the inversion is present at a frequency of approximately 0.1) provides useful information. This panel shows that *St* haplotypes display the same pattern as for normal genomic regions, as might be expected from its high frequency. For *In* haplotypes, LD starts off at a very high value, but approaches the baseline value for sites about 20kb apart.

There are three reasons for expecting the relatively rare *In* haplotypes to experience more LD than commoner *St* haplotypes. First, an inversion with frequency *x* << 0.5 experiences a higher sampling variance for the drift process than *St*, since its effective population size is *Nx*. Second, the effective recombination rate between a pair of *In* sites (ignoring the low frequency of exchange between *In* and *St*) is *rx*, where *r* is the recombination rate between the sites, so that their net scaled recombination rate is 4*Nx*^2^*r*. This low effective recombination rate may intensify Hill-Robertson interference effects (Charlesworth and Jensen 2021), reducing *N*_*e*_ even further. Third, exchange between *In* and *St* in heterokarotypes is analogous to migration between demes in a subdivided population, inducing some LD (Wakeley & Lessard 2003). However, this effect is small for a pair of populations (Wakeley & Lessard 2003) and can be neglected as a first approximation, relative to the much bigger effect of the low frequency of the inversion.

Remarkably, LD within inversion haplotypes of *In(3R)P* seems to be close to the predictions of equilibrium neutral theory. The expected value of *R*^2^ for the normal autosomal genome is given by the equation of Ohta & Kimura (1971):

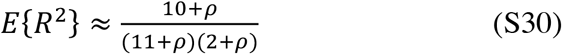

For the inversion, *ρ* = 4*Nr* is replaced by *ρ*_1_ = 4*Nx*^2^*r*.

In *Drosophila*, gene conversion plays an important role in determining *r*. If we have two sites that are separated by substantially more than the mean tract length of around 440 basepairs, the chance that they recombine is approximately twice the probability that one of them experiences a conversion event, which is typically 10^−5^ in female meiosis (Korunes and Noor 2019). This is offset by the lack of exchange in male meiosis, so the net contribution to *r* for autosomal loci is 10^−5^. For crossovers, the typical rate of recombination in female meiosis for a pair of sites separated by *l* basepairs is approximately 2*l* × 10^−8^, so the net contribution to *r* is *l* × 10^−8^ (Campos and Charlesworth 2019).

For the normal part of the genome, an *N*_*e*_ of approximately 1.6 × 10^6^ is a widely accepted estimate for the Zambian population (Johri et al. 2020), so that *ρ* ≈ 4 × 10^6^ x *r* = 64 + 0.064*l*, which can substituted into Equation (S30). For sites 1kb apart, *ρ* = 128, E{*R*^2^} ≈ 0.007; for sites 10kb apart, *ρ* = 704, E{*R*^2^} ≈ 0.001. For an inversion with *x* = 0.1, we have *r*_1_ = 10^−6^ + *l* × 10^−9^, and *ρ*_1_ = 0.64 + 0.00064*l*. For sites 1kb apart, *ρ*_1_= 1.28 and E{*R*^2^} ≈ 0.28; for sites 10kb apart, *ρ*_1_ = 7.04, E{*R*^2^} ≈ 0.10; for sites 20kb apart, *ρ*_1_ = 13.44, E{*R*^2^} ≈ 0.06; for sites 40kb apart, *ρ*_1_ = 26.24, E{*R*^2^} ≈ 0.03. These values are very similar to what is seen in panel A of Kapun et al. (2023), suggesting that complications due to Hill-Robertson interference do not play a major role in affecting LD within the *In(3R)P* inversion in Zambia.

It follows that, if the means of diversity and divergence statistics for the inversion over a large number of 100kb windows are used, as was done for the central regions of *In(3R)P* by Kapun et al. (2023), most pairs of sites will be approximately independent of each other (i.e., there is strong statistical mixing: Billingsley [1995]), and their variances will be close to 1/*m* times the variances for individual sites. To allow for the presence of the high degree of correlation between a substantial fraction of pairs within the same window in the *In* sample, one could (probably quite conservatively) elevate the multiplier to 2/*m* (essentially zero correlation is expected between non-adjacent windows).

While exact calculations of the variances of the mean coalescence time for a sample of *n* alleles are complicated, even for the case of a single population (Tajima 1983), a very conservative procedure is to assume that, for a sample of size *n*, all combinations of pairwise coalescence times are completely correlated. The variance of the mean of the pairwise coalescent times over all *n*(*n* – 1)/2 values is then equal to the variance for a single pair. It follows that upper bounds to the variances of the mean diversity and divergence statistics over *m* sites from the stochasticity of the coalescent process are given by the squares of their expectations, unless *ρ* is of order 1 or less, giving an upper bound to their coefficients of variation of approximately 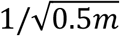.

In addition, we need to consider the variance associated with the mutational process itself; for a pair of alleles of type *ij* under the infinite sites model, this equal to E{*T*_*ij*_}*θ*, where *θ* is the population scaled mutation rate (Tajima 1983). For the mean of a pairwise diversity statistic for a sample of *n* alleles taken over *m* sites, the argument used above shows that the variance ≈ 2E{*T*_*ij*_}*θ*/*m*, corresponding to a coefficient of variation of 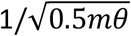. For *θ* ≈ 0.01, this dominates over the coalescent process stochasticity term, but is still very small for *m* = 10^4^. By the central limit theorem for strongly mixed correlated sites (Billingsley 1995), normal deviate tests on pairwise diversity and divergence statistics should be quite robust for data taken over a large number of nucleotide sites, as would be the case for the central region of a several megabase inversion such as *In(3R)P*, with ample power to detect small differences between theoretical predictions and observations.

Difficulties are, however, likely to arise for comparisons involving the relatively small regions associated with inversion breakpoints, where the arguments based on large numbers of sites are likely to fail; simulation-based methods will be probably be needed for inference and hypothesis testing with these regions. As noted in the text, however, patterns such as an increase in 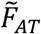 towards inversion breakpoints appear to be repeatable across several independently-derived inversions in *D. melanogaster*, suggesting that conclusions based on such patterns are statistically robust.

---

**Figure S1.**
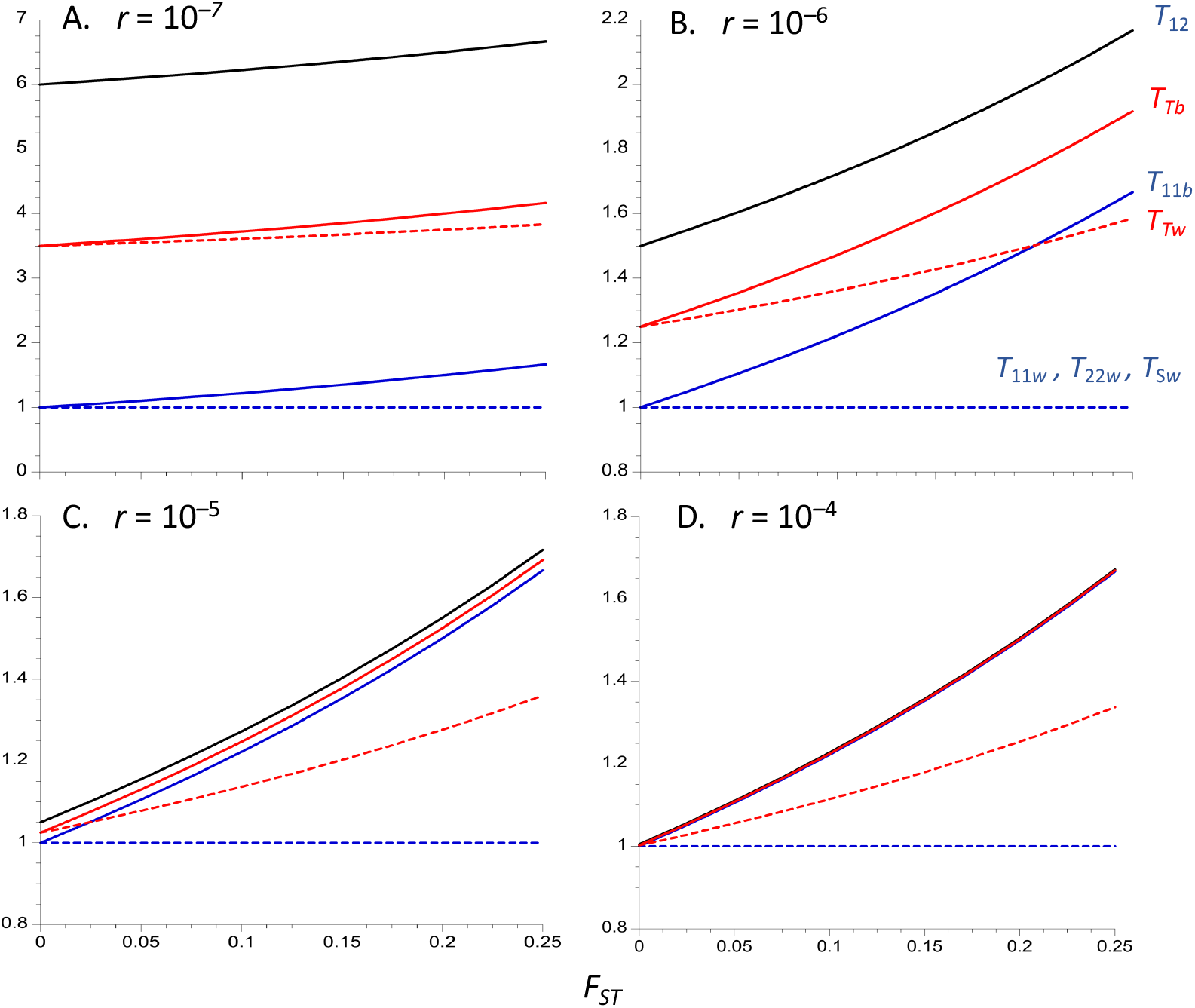
The equilibrium population statistics for an inversion polymorphism where the inversion is maintained at a constant frequency of 0.5, under the same assumptions as in Figure 1. The X axis is the equilibrium *F*_*ST*_ for neutral sites unlinked to the inversion. Subscripts 1 and 2 denote alleles sampled from the inversion and standard arrangement, respectively; subscripts *w* and *b* denote alleles sampled from the same and from separate demes, respectively; subscript *T* denotes pairs of alleles sampled without regard to karyotype. In this case, *T*_11*w*_ =*T*_22*w*_ = *T*_*Sw*_ = 1 (blue dashed line) for all values of *F*_*ST*_. The solid black curve is *T*_12_, the sold blue curve is *T*_11*b*_ and the solid and dashed red curves are *T*_*Tb*_ and *T*_*Tw*_, respectively.

**Figure S2.**
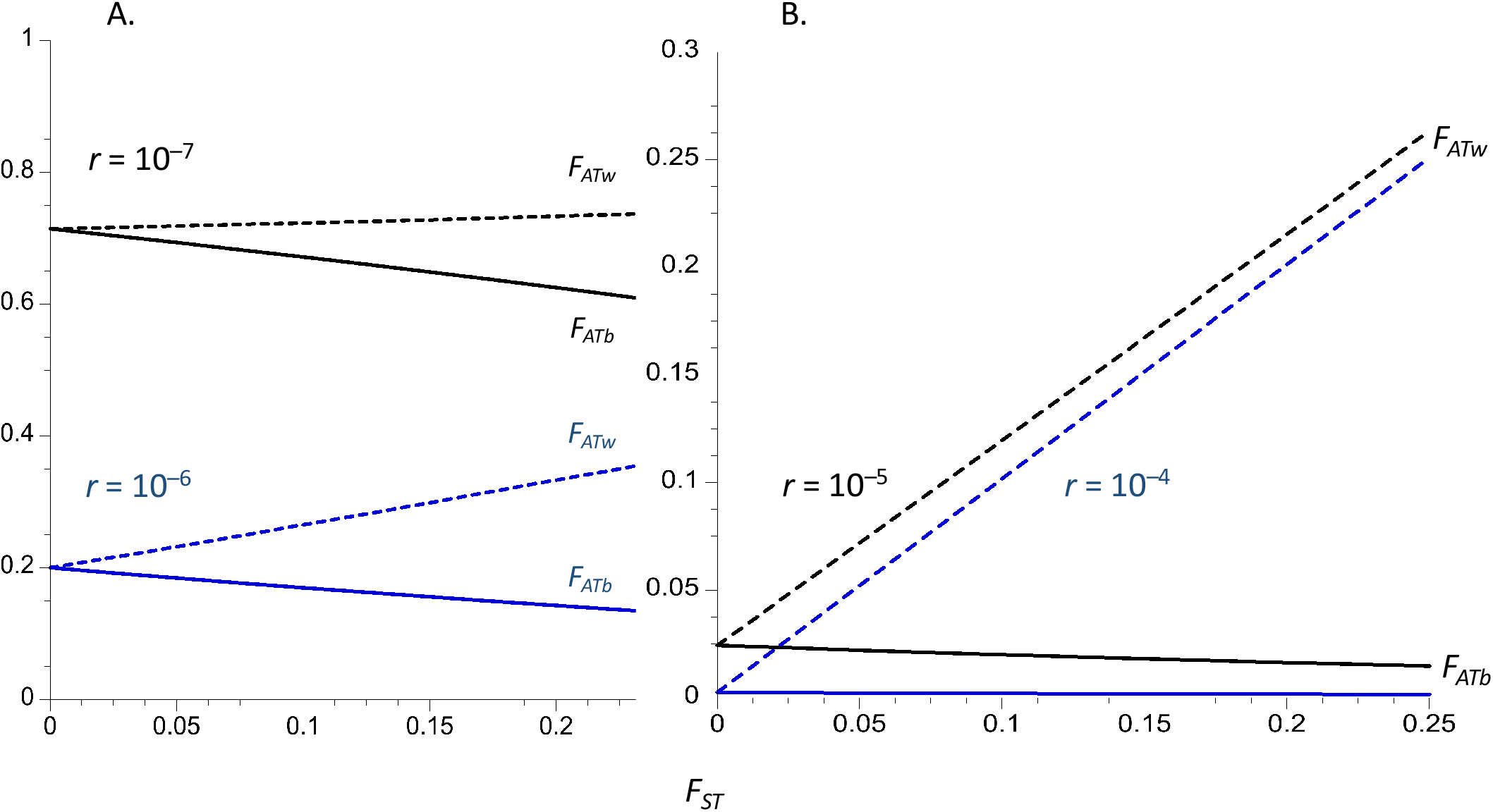
Equilibrium values of *F*_*ATw*_ (dashed curves) and *F*_*ATb*_ (blue solid curves), *T*_11*w*_/*T*_22*w*_ (black dashed curves) and *T*_11*w*_/*T*_22*w*_ (black solid curves), for an inversion polymorphism where the inversion is maintained at a constant frequency of 0.5. Four different recombination rates are modeled, with the higher rate for the panel corresponding to the black curves and the lower rate to the blue curves. The dashed curves are for within-deme samples and the solid curve for samples from separate demes. The population and recombination parameters in Figure 1 are used.

## Literature cited

Andolfatto P, Wall JD, Kreitman M. 1999. Unusual haplotype structure at the proximal breakpoint of In(2L)t in a natural population of Drosophila melanogaster. Genetics 153:1297–1311.

Andolfatto P, Depaulis F, Navarro A. 2001. Inversion polymorphisms and nucleotide variability in Drosophila. Genet. Res. 77:1–8.

Assaf ZJ, Tilk S, Siegal ML, Petrov DA. 2017. Deep sequencing of natural and experimental populations of Drosophila melanogaster reveals biases in the spectrum of new mutations. Genome Res. 27:1988–2000.

Aulard S, David, J.R., Lemeunier F. 2002. Chromosomal inversion polymorphism in Afrotropical populations of Drosophila melanogaster. Genet. Res. 79:49–63.

Bartolomé C, Charlesworth B. 2006. Rates and patterns of chromosomal evolution in Drosophila pseudoobscura and D. miranda. Genetics 173:779–791.

Barton NH, Etheridge AM, Kelleher J, Véber A. 2013. Genetic hitchhiking in spatially extended populations. Theor. Pop. Biol. 87:75–89.

Bergero R, Gardner J, Bader B, Yong L, Charlesworth d. 2019. Exaggerated heterochiasmy in a fish with sex-linked male coloration polymorphisms. Proc. Natl. Acad. Sci. USA 116:6924–6931.

Buffalo V. 2021. Quantifying the relationship between genetic diversity and population size suggests natural selection cannot explain Lewontin’s paradox. Elife 10:e67509.

Cáceres M, Puig M, Ruiz A. 2001. Molecular characterization of two natural hotspots in the Drosophila buzzatii genome induced by transposon insertions. Genome Res. 11:1353–1364.

Charlesworth B. 2020. How good are predictions of the effects of selective sweeps on levels of neutral diversity? Genetics 216:1217–1238.

Charlesworth B. 1998. Measures of divergence between populations and the effect of forces that reduce variability. Mol. Biol. Evol. 15:538–543.

Charlesworth B, Charlesworth D. 2010. Elements of Evolutionary Genetics. Greenwood Village, CO: Roberts and Company.

Charlesworth B, Jensen JD. 2021. The effects of selection at linked sites on patterns of genetic variability. Ann. Rev. Ecol. Evol. Syst. 52:177–197.

Charlesworth B, Charlesworth D, Barton NH. 2003. The effects of genetic and geographic structure on neutral variation. Ann. Rev. Ecol. Evol. Syst. 34:99–125.

Charlesworth B, Nordborg M, Charlesworth D. 1997. The effects of local selection, balanced polymorphism and background selection on equilibrium patterns of genetic diversity in subdivided populations. Genet. Res. 70:155–174.

Charlesworth D. 2003. Effects of inbreeding on the genetic diversity of plant populations. Phil. Trans. Roy. Soc. B 358:1051–1070.

Chovnick A. 1973. Gene conversion and transfer of genetic information within the inverted region of inversion heterozygotes. Genetics 74:123–131.

Corbett-Detig RB, Hartl DL. 2012. Population genomics of inversion polymorphisms in Drosophila melanogaster. PloS Genet. 8:e1003056.

Crow JF, Aoki K. 1984. Group selection for a polygenic behavioral trait - estimating the degree of population subdivision. Proc. Natl. Acad. Sci. USA 81:6073–6077.

Crown KN, Miller DE, Sekelsky J, Hawley RS. 2018. Local inversion heterozygosity alters recombination throughout the genome. Curr. Biol. 28:2984–2990.

Dobzhansky T. 1947. Adaptive changes induced by natural selection in wild populations of Drosophila. Evolution 1:1–16.

Dobzhansky T. 1950. Genetics of natural populations. XIX. Origin of heterosis through naturak selection in populations of Drosophila pseudoobscura. Proc. Natl. Acad. Sci.USA 35:288–302.

Faria R, Chaube P, Morales HE, Larsson T, Lemmon AR, Lemmon EM, Rafajlovic M, Panova M, Ravinet M, Johannesson K, et al. 2019. Multiple chromosomal rearrangements in a hybrid zone between Littorina saxatilis ecotypes. Mol. Ecol. 28:1375–1393.

Gammerdinger W, Toups M, Vicoso B. 2020. Disagreement in FST estimators: A case study from sex chromosomes. Mol. Ecol. Res. 20:1517–1525.

Gong WJ, McKim KS, Hawley RS. 2005. All paired up with no place to go: Pairing, synapsis, and DSB formation in a balancer heterozygote. PloS Genet. 1:e67.

Guerrero RF, Rousset F, Kirkpatrick M. 2012. Coalescent patterns for chromosomal inversions in divergent populations. Phil. Trans. R. Soc. B 367:430–438.

Haddrill PR, Thornton KR, Charlesworth B, Andolfatto P. 2005. Multilocus patterns of nucleotide variability and the demographic and selection history of Drosophila melanogaster populations. Genome Research 15:790–799.

Hasson E, Eanes WF. 1996. Contrasting histories of three gene regions associated with In (3L)Payne of Drosophila melanogaster. Genetics 144.

Hill WG, Robertson A. 1968. Linkage disequilibrium in finite populations. Theor. Appl. Genet. 38:226–231.

Hudson RR. 1990. Gene genealogies and the coalescent process. Oxf. Surv. Evol. Biol. 7:1–45.

Hudson RR, Boos DD, Kaplan NL. 1992. A statistical test for detecting geographic subdivision. Mol. Biol. Evol. 9:138–151.

Hudson RR, Kaplan NL. 1988. The coalescent process in models with selection and recombination. Genetics 120:831–840.

Hudson RR, Slatkin M, Maddison WP. 1992. Estimation of levels of gene flow from population data. Genetics 132:583–589.

Innan H, Nordborg M. 2003. The extent of linkage disequilibrium and haplotype sharing around a polymorphic site. Genetics 165:437–444.

Ishii K, Charlesworth B. 1977. Associations between allozyme loci and gene arrangements due to hitch-hiking effects of new inversions. Genet. Res. 30:93–106.

Ives PT, Band HT. 1986. Continuing studies on the South Amherst Drosophila melanogaster natural population during the 1970’s and 1980’s. Evolution 40:1289–1302.

Johri P, Charlesworth B, Jensen, JD. 2020. Toward an evolutionarily appropriate null model: Jointly inferrring demography and purifying selection. Genetics 215:173–192.

Kaplan NL, Dardern T, Hudson RR. 1988. The coalescent process in models with selection and recombination. Genetics 120:831–840.

Kapun M, Flatt T. 2019. The adaptive significance of chromosomal inversion polymorphisms in Drosophila melanogaster. Mol. Ecol. 28:1263–1282.

Kapun M, Fabian DK, Goudet J, Flatt T. 2016. Genomic evidence for adaptive inversion clines in Drosophila melanogaster. Mol. Biol. Evol. 33:1317–1336.

Kapun M, Durmaz Mitchell E, Kawecki T, Schmidt P, Flatt T. 2023. An ancestral balanced inversion polymorphism confers global adaptation. Mol. Biol. Evol., in press.

Kennington WJ, Hoffmann AA. 2013. Patterns of genetic variation across inversions, geographic variation in the In(2L)t inversion in populations of Drosophila melanogaster from eastern Australia. BMC Evol. Biol. 13:100.

Kimura M. 1971. Theoretical foundations of population genetics at the molecular level. Theor. Pop. Biol. 2:174–208.

Kirkpatrick M, Guerrero RF. 2014. Signatures of sex-antagonistic selection on recombining sex chromosomes. Genetics 197:531–541.

Korunes K, Noor MAF. 2019. Pervasive gene conversion in chromosomal inversion heterozygotes. Mol. Ecol. 28:1302–1315.

Koury SA. 2023. Predicting recombination suppression outside chromosomal inversions in Drosophila melanogaster using crossover interference theory. Heredity: in press.

Krimbas CB, Powell JR. 1992a. Drosophila Inversion Polymorphism. In. Boca Raton, FL: CRC Press.

Krimbas CB, Powell JR. 1992b. Introduction. In: Krimbas CB, Powell JR, editors. Drosophila Inversion Polymorphism. Boca Raton, FL: CRC Press. p. 1–52.

Laayouni H, Hasson E, Santos M, Fontdevila A. 2003. The evolutionary history of Drosophila buzzatii. XXXV. Inversion polymorphism and nucleotide variability in different regions of the second chromosome. Mol. Biol. Evol. 20:931–944.

Lack JB, Cardeno CM, Crepeau MW, Taylor W, Corbett-Detig RB, Stevens KA, Langley CH, Pool JE. 2015. The Drosophila Genome Nexus: A population genomic resource of 623 Drosophila melanogaster genomes, including 197 from a single ancestral range population. Genetics 199:1229–1241.

Lack JB, Lange JD, Tang AD, Corbett-Detig RB, Pool JE. 2016. A thousand fly genomes: An expanded Drosophila Genome Nexus. Mol. Biol. Evol. 33:3308–3313.

Langley CH, Stevens K, Cardeno C, Lee YCG, Schrider DR, Cardeno C., et al. 2012. Genomic variation in natural populations of Drosophila melanogaster. Genetics 192:533–598.

Li L, Brent E, Hadjipanteli S, Galey M, Miller DE, Crown N. 2022. Heterozygous Inversion breakpoints suppress meiotic crossovers by altering recombination repair outcomes. BioRxiv 2022.11.09.515852.

Lohse K, Harrison RJ, Barton NH. 2011. A general method for calculating likelihoods under the coalescent process. Genetics 189:977–987.

Lange JD, Bastide H, Lack JB, Pool JE. 2022. A population genomic assessment of three decades of evolution in a natural Drosophila population. Mol. Biol. Evol. 39:msab368.

Maruyama T. 1974. A simple proof that certain quantitities are independent of the geographic structure of populations. Theor. Pop. Biol. 5:148–154.

Maynard Smith J, Haigh J. 1974. The hitch-hiking effect of a favourable gene. Genet. Res. 23: 23–35.

Mérot C, Llaurens V, Normandeau E, Bernatchez L, Wellenreuther M. 2018. Balancing selection via life-history trade-offs maintains an inversion polymorphism in a seaweed fly. Nat. Commun. 11:1–11.

Mérot C, Berdan EL, Cayuela H, Djambazian H, Ferchaud A, Laporte M, Normandeau E, Ragoussis J, Wellenreuther M, Bernatchez L. 2021. Locally adaptive inversions modulate genetic variation at different geographic scales in a seaweed fly. Mol. Biol. Evol. 38:3953–3971.

Mukai T, Watanabe TK, O. Y. 1974. The genetic structure of natural populations of Drosophila melanogaster. XII. Linkage disequilibrium in a local population. Genetics 77:771–793.

Mukai T, Yamaguchi O. 1974. The genetic structure of natural populations of Drosophila melanogaster. XI. Genetic variability in a local population. Genetics 76:339–366.

Munté A, Rozas J, Aguadé M, Segarra C. 2005. Chromosomal inversion oolymorphism leads to extensive genetic structure: A multilocus survey in Drosophila subobscura. Genetics 169:1573–1581.

Nagylaki T. 1998. The expected number of heterozygous sites in a subdivided population. Genetics 149:1599–1604.

Nagylaki T. 1982. Geographical invariance in population genetics. J. Theor. Biol. 99:159–172.

Navarro A, Barbadilla A, Ruiz A. 2000. Effect of inversion polymorphism on the neutral nucleotide variability of linked chromosomal regions in Drosophila. Genetics 155:685–698.

Navarro A, Betrán E, Barbadilla A, Ruiz A. 1997. Recombination and gene flux caused by gene conversion and crossing over in inversion heterokaryotypes. Genetics 146:695–709.

Nei M. 1982. Evolution of human races at the gene level. In: Alan LR, editor. Human genetics, part A: The unfolding genome. New York: Liss. p. 167–181.

Nei M. 1987. Molecular Evolutionary Genetics. New York: Columbia University Press.

Nei M, Li W-H. 1975. Probability of identical monomorphism in related species. Genet. Res. 26:31–43.

Nobréga C, Khadem M, Aguadé M, Segarra C. 2008. Genetic exchange versus genetic differentiation in a medium-sized inversionof Drosophila: The A2/Ast arrangements of Drosophila subobscura. Mol. Biol. Evol. 25:1534–1543.

Nordborg M. 1997. Structured coalescent processes on different time scales. Genetics 146:1501–1514.

Nordborg M, Innan H. 2003. The genealogy of sequences containing multiple sites subject to strong selection in a subdivided population. Genetics 163:1201–1213.

Ohta T, Kimura M. 1970. Development of associative overdominance through linkage disequilibrium in finite populations. Genet. Res. 18:277–286.

Ohta T, Kimura M. 1971. Linkage disequilibrium between two segregating nucleotide sites under steady flux of mutations in a finite population. Genetics 68:571–580.

Ohta T, Kimura M. 1969. Linkage disequilibrium due to random genetic drift. Genet. Res. 13:47–55.

Orengo DJ, Puerma E, Cereijo U, Aguadé M. 2019. The molecular genealogy of sequential overlapping inversions implies both homologous chromosomes of a heterokaryotype in an inversion origin. Scientific Reports 9:17009.

Pannell JR, Charlesworth B. 2000. Effects of metapopulation processes on measures of genetic diversity. Phil. Trans. R. Soc. B 355:1851–1864.

Pegueroles C, Aquadro CF, Mestres F, Pascual M. 2013. Gene flow and gene flux shape evolutionary patterns of variation in Drosophila subobscura. Heredity 110:520–529.

Prakash S, Lewontin RC. 1968. A molecular approach to the study of genic heterozygosity in natural populations. III. Direct evidence of coadaptation in gene arrangements of Drosophila. Proc. Natl. Acad. Sci. USA 59:398–405.

Prakash S, Lewontin RC. 1971. A molecular approach to the study of genic heterozygosity in natural populations. V. Further direct evidence of coadaptation in inversions of Drosophila. Genetics:405–408.

Rogers AR. 1995. Genetic evidence for a pleistocene explosion. Evolution 49:608–615.

Rousset F, Kirkpatrick M, Guerrero RF. 2014. Matrix inversions for chromosomal inversions: A method to construct summary statistics in complex coalescent models. Theor. Pop. Biol. 97:1–10.

Roux C, Fraisse C J. R, Anciaux Y, Galtier N, Bierne N. 2016. Shedding light on the grey zone of speciation along a continuum of genomic divergence. PloS Biol. 14:e2000234.

Rozas J, Segarra C, Ribó G, Aguadé M. 1999. Molecular population genetics of the rp49 gene region in different chromosomal inversions of Drosophila subobscura. Genetics 151:189–202.

Schaeffer SW. 2002. Molecular population genetics of sequence length diversity in the Adh region of Drosophila pseudoobscura. Genet. Res. 80:163–175.

Sheppard PM. 1975. Natural Selection and Heredity. 4th Edition. London: Hutchinson.

Singh RS, Rhomberg LR. 1987. A comprehensive study of genic variation in natural populations of Drosophila melanogaster. II. Estimates of heterozygosity and patterns of geographic differentiation. Genetics 117:255–271.

Sprengelmeyer QD, Mansourian S, Lange JD, Matute DR, Cooper BS, Jirle EV, Stensmyr MC, Pool JE. 2020. Recurrent collection of Drosophila melanogaster from wild African environments and genomic insights into species history. Mol. Biol. Evol. 37:627–638.

Strobeck C. 1983. Expected linkage disequilibrium for a neutral locus linked to a chromosomal rearrangement. Genetics 103:545–555.

Sturtevant AH. 1926. A crossover reducer in Drosophila melanogaster due to inversion of a section of chromosomes. Biol. Zentralbl. 46:697–702.

Sturtevant AH. 1917. Genetic factors affecting the strength of linkage in Drosophila. Proc. Natl. Acad. Sci. USA 3:555–558.

Sturtevant AH, Beadle GW. 1936. The relations of inversions in the X chromosome of Drosophila melanogaster to crossing over and disjunction. Genetics 21:554–604.

Takahata N. 1990. A simple genealogical structure of strongly balanced allelic lines and trans-species evolution of polymorphism. Proc. Natl. Acad. Sci. USA 87:2419–2423.

Toups M, Rodrigues N, Perrin N, Kirkpatrick M. 2019. A reciprocal translocation radically reshapes sex-linked inheritance in the common frog. Mol. Ecol. 28:1877–1889.

Villoutreix R, Ayala D, Joron M, Gompert Z, Feder JL, Nosil P. 2021. Inversion breakpoints and the evolution of supergenes. Mol. Ecol. 30:2738–2755.

Wakeley J, Aliacar N. 2001. Gene genealogies in a metapopulation. Genetics 159:893–905.

Wakeley J, Lessard S. 2003. Theory of the effects of population structure and sampling on patterns of linkage disequilibrium applied to genomic data from humans. Genetics 164:1043–1053.

Weir BS, Cockerham CC. 1984. Estimating F-statistics for the analysis of population structure. Evolution 38:1358–1370.

Wellenreuther M, Bernatchez L. 2018. Eco-evolutionary genomics of chromosomal inversions. Trnds. Ecol. Evol 33:6.

Wellenreuther M, Mérot C, Berdan EL, Bernatchez L. 2019. Going beyond SNPs: The role of structural genomic variants in adaptive evolution and species diversification. Mol. Ecol. 28:1203–1209.

White BJ, Hahn MW, Pombi M, Cassone BJ, Lobo NF, Simard F, Besansky NJ. 2007. Localization of candidate regions maintaining a common polymorphic inversion (2La) in Anopheles gambiae. PloS Genet. 3:e217.

Wright S. 1951. The genetical structure of populations. Ann. Eugen. 15:323–354.

Wright S, Dobzhansky T, Hovanitz W. 1942. Genetics of natural populations. VII. The allelism of lethals in the third chromosome of Drosophila pseudoobscura. Genetics 27:363–394.

Zeng K, Charlesworth B, Hobolth A. 2021. Studying models of balancing selection using phase-type theory. Genetics 218:iyab055.

## Literature cited

Billingsley P. 1995. Probability and Measure. 3rd edition. Chichester, UK: John Wiley.

Campos JL, Charlesworth B. 2019. The effects on neutral variability of recurrent selective sweeps and background selection. Genetics 212:287–303.

Kapun M, Durmaz Mitchell E, Kawecki T, Schmidt P, Flatt T. 2023. An ancestral balanced inversion polymorphism confers global adaptation. Mol. Biol. Evol. In press.

McVean GAT. 2002. A genealogical interpretation of linkage disequilibrium. Genetics 62:987–991.

Tajima F. 1983. Evolutionary relationship of DNA sequences in a finite population. Genetics 105:437–460.

