## Supplementary Tables for "The effects of inversion polymorphisms on patterns of neutral genetic diversity"

**Table S1**

**Equilibrium coalescent times for inversion and standard arrangements in a subdivided population**

**Inversion frequency=	0.1**

**Local Ne=	5000**

**Total Ne=	1000000**

**Number of populations =	200**

**Initial neutral Fst=	2.50E-002**

**Steps of FST=	2.5E-002**

**Initial gene conversion rate=	9.9999999999999995E-008**

**Steps of gene conversion rates (ratios)=	10.000000000000000**

**Number of increments=	4**

**Gene conversion rate=	9.9999999999999995E-008**

**Scaled value=	0.4**

Results for equivalent panmictic population

Times are scaled by 2Nd

T11=	0.305019289	T12=	6.00000000	T22=	1.07722008

TT=	1.95559847	TS=	1.00000000

FAT=	0.488647580	T11/T22=	0.283154100

Results for subdivided population

Fst=	2.5000000000000001E-002

Scaled migration rate=	39.000000000000000	Migration rate=	1.9499999999999999E-003

Alleles sampled from different populations

T11b=	0.38137720374659301	T12b=	6.0354084750480110	T22b=	1.1080786163037240

TTb=	1.9877309767521245	TSb=	1.0354084750480108

FATb=	0.47910029719422687	T11/T22=	0.344178826

Alleles sampled from the same population

T11w=	0.30383083563017776	T12w=	6.0355366801762163	T22w=	1.0775335022846340

TTw=	1.9622370476385744	TSw=	1.0001632356191883

FATw=	0.49029438781475454	T11/T22=	0.281968802

Fst=	5.0000000000000003E-002

Scaled migration rate=	18.999999999999996	Migration rate=	9.4999999999999978E-004

Alleles sampled from different populations

T11b=	0.46097604162330830	T12b=	6.0726091063843928	T22b=	1.1405683358022909

TTb=	2.0215397515652795	TSb=	1.0726091063843928

FATb=	0.46940983695528582	T11/T22=	0.404163450

Alleles sampled from the same population

T11w=	0.30250154450360134	T12w=	6.0728722642791295	T22w=	1.0778518531731356

TTw=	1.9692020240855193	TSw=	1.0003168223061822

FATw=	0.49201919860369792	T11/T22=	0.280652255

Fst=	7.5000000000000011E-002

Scaled migration rate=	12.333333333333332	Migration rate=	6.1666666666666662E-004

Alleles sampled from different populations

T11b=	0.54403543337370186	T12b=	6.1117432832507905	T22b=	1.1748219332371332

TTb=	2.0571599112409578	TSb=	1.1117432832507901

FATb=	0.45957371754335619	T11/T22=	0.463079065

Alleles sampled from the same population

T11w=	0.30106434379271213	T12w=	6.1121486886561955	T22w=	1.0781754924914673

TTw=	1.9765195563141309	TSw=	1.0004643776215918

FATw=	0.49382520682604025	T11/T22=	0.279235005

Fst=	0.10000000000000001

Scaled migration rate=	9.0000000000000000	Migration rate=	4.4999999999999999E-004

Alleles sampled from different populations

T11b=	0.63079614468311151	T12b=	6.1529677637253410	T22b=	1.2109868325078110

TTb=	2.0947414932487196	TSb=	1.1529677637253413

FATb=	0.44958947562679352	T11/T22=	0.520894289

Alleles sampled from the same population

T11w=	0.29953501588997822	T12w=	6.1535233192808967	T22w=	1.0785047630250379

TTw=	1.9842184056797421	TSw=	1.0006077883115321

FATw=	0.49571691027190645	T11/T22=	0.277731746

Fst=	0.12500000000000000

Scaled migration rate=	7.0000000000000000	Migration rate=	3.5000000000000000E-004

Alleles sampled from different populations

T11b=	0.72152279781838535	T12b=	6.1964568851710764	T22b=	1.2492273393213753

TTb=	2.1344516121592916	TSb=	1.1964568851710764

FATb=	0.43945466912660736	T11/T22=	0.577575266

Alleles sampled from the same population

T11w=	0.29792115202883995	T12w=	6.1971711708853618	T22w=	1.0788400325944165

TTw=	1.9923304486811313	TSw=	1.0007481445378590

FATw=	0.49769971883914788	T11/T22=	0.276149511

Fst=	0.14999999999999999

Scaled migration rate=	5.6666666666666670	Migration rate=	2.8333333333333335E-004

Alleles sampled from different populations

T11b=	0.81650702138672060	T12b=	6.2424051165106240	T22b=	1.2897271270799462

TTb=	2.1764769641205364	TSb=	1.2424051165106238

FATb=	0.42916688897157673	T11/T22=	0.633085072

Alleles sampled from the same population

T11w=	0.29622594389061691	T12w=	6.2432874694518006	T22w=	1.0791816964462790

TTw=	2.0008911780617167	TSw=	1.0008861211907130

FATw=	0.49977983202450749	T11/T22=	0.274491251

Fst=	0.17500000000000002

Scaled migration rate=	4.7142857142857135	Migration rate=	2.3571428571428569E-004

Alleles sampled from different populations

T11b=	0.91607112976378779	T12b=	6.2910300706023730	T22b=	1.3326921751399938

TTb=	2.2210267858694603	TSb=	1.2910300706023732

FATb=	0.41872377279908557	T11/T22=	0.687383890

Alleles sampled from the same population

T11w=	0.29444997359743752	T12w=	6.2920906766629789	T22w=	1.0795301799226118

TTw=	2.0099402672726265	TSw=	1.0010221592900943

FATw=	0.50196422471379021	T11/T22=	0.272757530

Fst=	0.20000000000000001

Scaled migration rate=	4.0000000000000000	Migration rate=	2.0000000000000001E-004

Alleles sampled from different populations

T11b=	1.0205724419113942	T12b=	6.3425760768012198	T22b=	1.3783542584556452

TTb=	2.2683363675924064	TSb=	1.3425760768012200

FATb=	0.40812302091412511	T11/T22=	0.740428269

Alleles sampled from the same population

T11w=	0.29259212624336850	T12w=	6.3438260768012196	T22w=	1.0798859414467172

TTw=	2.0195222276584945	TSw=	1.0011565599263825

FATw=	0.50426068789192835	T11/T22=	0.270947248

Fst=	0.22500000000000001

Scaled migration rate=	3.4444444444444446	Migration rate=	1.7222222222222224E-004

Alleles sampled from different populations

T11b=	1.1304083765953425	T12b=	6.3973184395208680	T22b=	1.4269751131792596

TTb=	2.3186712445549102	TSb=	1.3973184395208680

FATb=	0.39736241487348256	T11/T22=	0.792171061

Alleles sampled from the same population

T11w=	0.29065007993644509	T12w=	6.3987700524240934	T22w=	1.0802494758708006

TTw=	2.0296871856910501	TSw=	1.0012895362773651

FATw=	0.50667790419317504	T11/T22=	0.269058287

Fst=	0.25000000000000000

Scaled migration rate=	3.0000000000000000	Migration rate=	1.4999999999999999E-004

Alleles sampled from different populations

T11b=	1.2460224962953286	T12b=	6.4555685420506608	T22b=	1.4788514360234750

TTb=	2.3723322257110873	TSb=	1.4555685420506606

FATb=	0.38643983912735247	T11/T22=	0.842560947

Alleles sampled from the same population

T11w=	0.28862057604981289	T12w=	6.4572352087173277	T22w=	1.0806213182372864

TTw=	2.0404918111018193	TSw=	1.0014212440185390

FATw=	0.50922555112936507	T11/T22=	0.267087609

**Gene conversion rate=	9.9999999999999995E-007**

**Scaled value=	4**

Results for equivalent panmictic population

Times are scaled by 2Nd

T11=	0.470588237	T12=	1.50000000	T22=	1.05882359

TT=	1.13235295	TS=	1.00000000

FAT=	0.116883099	T11/T22=	0.444444418

Results for subdivided population

Fst=	2.5000000000000001E-002

Scaled migration rate=	39.000000000000000	Migration rate=	1.9499999999999999E-003

Alleles sampled from different populations

T11b=	0.56574155825635286	T12b=	1.5387529531343556	T22b=	1.0913097747874669

TTb=	1.1665938647245957	TSb=	1.0387529531343556

FATb=	0.10958476249180360	T11/T22=	0.518406034

Alleles sampled from the same population

T11w=	0.45056981166855803	T12w=	1.5388811582625608	T22w=	1.0612291716137028

TTw=	1.1410999356110458	TSw=	1.0001632356191883

FATw=	0.12350951533126464	T11/T22=	0.424573541

Fst=	5.0000000000000003E-002

Scaled migration rate=	18.999999999999996	Migration rate=	9.4999999999999978E-004

Alleles sampled from different populations

T11b=	0.65832852331905301	T12b=	1.5786582245972889	T22b=	1.1253615247393149

TTb=	1.2022846006995476	TSb=	1.0786582245972889

FATb=	0.10282621604762043	T11/T22=	0.584992945

Alleles sampled from the same population

T11w=	0.43180144630426170	T12w=	1.5789213824920256	T22w=	1.0634851974175064

TTw=	1.1499468732197875	TSw=	1.0003168223061818

FATw=	0.13011909888902073	T11/T22=	0.406024873

Fst=	7.5000000000000011E-002

Scaled migration rate=	12.333333333333332	Migration rate=	6.1666666666666662E-004

Alleles sampled from different populations

T11b=	0.74888586530068824	T12b=	1.6198965342727596	T22b=	1.1611199419363230

TTb=	1.2395773877905252	TSb=	1.1198965342727596

FATb=	9.6549723072223648E-002 T11/T22=	0.644968569

Alleles sampled from the same population

T11w=	0.41419070172254047	T12w=	1.6203019396781650	T22w=	1.0656058971659306

TTw=	1.1589370328636990	TSw=	1.0004643776215916

FATw=	0.13673965948825206	T11/T22=	0.388690323

Fst=	0.10000000000000001

Scaled migration rate=	9.0000000000000000	Migration rate=	4.4999999999999999E-004

Alleles sampled from different populations

T11b=	0.83791524787265947	T12b=	1.6626575463306998	T22b=	1.1987400239371486

TTb=	1.2786369302073430	TSb=	1.1626575463306998

FATb=	9.0705485769002503E-002 T11/T22=	0.698996663

Alleles sampled from the same population

T11w=	0.39764406476923775	T12w=	1.6632131018862555	T22w=	1.0676037575940089

TTw=	1.1681138426383657	TSw=	1.0006077883115319

FATw=	0.14339874095532978	T11/T22=	0.372464091

Fst=	0.12500000000000000

Scaled migration rate=	7.0000000000000000	Migration rate=	3.5000000000000000E-004

Alleles sampled from different populations

T11b=	0.92589287194396774	T12b=	1.7071429021188189	T22b=	1.2383929054715801

TTb=	1.3196429045328071	TSb=	1.2071429021188189

FATb=	8.5250337062825698E-002 T11/T22=	0.747656822

Alleles sampled from the same population

T11w=	0.38207353549231504	T12w=	1.7078571878331046	T22w=	1.0694897677651412

TTw=	1.1775217410546464	TSw=	1.0007481445378585

FATw=	0.15012342477724505	T11/T22=	0.357248425

Fst=	0.14999999999999999

Scaled migration rate=	5.6666666666666670	Migration rate=	2.8333333333333335E-004

Alleles sampled from different populations

T11b=	1.0132781434076163	T12b=	1.7535694354905325	T22b=	1.2802684679441898

TTb=	1.3627927388571659	TSb=	1.2535694354905325

FATb=	8.0146672529402996E-002 T11/T22=	0.791457534

Alleles sampled from the same population

T11w=	0.36739847738753667	T12w=	1.7544517884317090	T22w=	1.0712736371688434

TTw=	1.1872069527983462	TSw=	1.0008861211907127

FATw=	0.15694048216990275	T11/T22=	0.342954844

Fst=	0.17500000000000002

Scaled migration rate=	4.7142857142857135	Migration rate=	2.3571428571428569E-004

Alleles sampled from different populations

T11b=	1.1005217452258673	T12b=	1.8021727183541276	T22b=	1.3245783820350454

TTb=	1.4083047962043886	TSb=	1.3021727183541276

FATb=	7.5361582333813182E-002 T11/T22=	0.830846846

Alleles sampled from the same population

T11w=	0.35354580185227857	T12w=	1.8032333244147336	T22w=	1.0729639767831849

TTw=	1.1972182776075546	TSw=	1.0010221592900943

FATw=	0.16387664804911450	T11/T22=	0.329503894

Fst=	0.20000000000000001

Scaled migration rate=	4.0000000000000000	Migration rate=	2.0000000000000001E-004

Alleles sampled from different populations

T11b=	1.1880734150125345	T12b=	1.8532110592203399	T22b=	1.3715596863545405

TTb=	1.4564220707569646	TSb=	1.3532110592203399

FATb=	7.0866140804211719E-002 T11/T22=	0.866220713

Alleles sampled from the same population

T11w=	0.34044954712940856	T12w=	1.8544610592203399	T22w=	1.0745684502371573

TTw=	1.2076079308230527	TSw=	1.0011565599263825

FATw=	0.17095893926099170	T11/T22=	0.316824436

Fst=	0.22500000000000001

Scaled migration rate=	3.4444444444444446	Migration rate=	1.7222222222222224E-004

Alleles sampled from different populations

T11b=	1.2763897115508014	T12b=	1.9069700980303197	T22b=	1.4214790298613773

TTb=	1.5074165289486812	TSb=	1.4069700980303197

FATb=	6.6634821225169838E-002 T11/T22=	0.897930741

Alleles sampled from the same population

T11w=	0.32805025666057086	T12w=	1.9084217109335455	T22w=	1.0760939006792309

TTw=	1.2184324700848210	TSw=	1.0012895362773651

FATw=	0.17821499273762753	T11/T22=	0.304852813

Fst=	0.25000000000000000

Scaled migration rate=	3.0000000000000000	Migration rate=	1.4999999999999999E-004

Alleles sampled from different populations

T11b=	1.3659420538034992	T12b=	1.9637681699144667	T22b=	1.4746377383712408

TTb=	1.5615942592033440	TSb=	1.4637681699144667

FATb=	6.2645010835774895E-002 T11/T22=	0.926289916

Alleles sampled from the same population

T11w=	0.31629432009015995	T12w=	1.9654348365811334	T22w=	1.0775464577883584

TTw=	1.2297538445940761	TSw=	1.0014212440185386

FATw=	0.18567341877341870	T11/T22=	0.293531954

**Gene conversion rate=	9.9999999999999991E-006**

**Scaled value=	40**

Results for equivalent panmictic population

Times are scaled by 2Nd

T11=	0.843478262	T12=	1.04999995	T22=	1.01739132

TT=	1.02152169	TS=	1.00000000

FAT=	2.10682750E-02 T11/T22=	0.829059839

Results for subdivided population

Fst=	2.5000000000000001E-002

Scaled migration rate=	39.000000000000000	Migration rate=	1.9499999999999999E-003

Alleles sampled from different populations

T11b=	0.91963076207742589	T12b=	1.0951727126788890	T22b=	1.0591218174734314

TTb=	1.0642160680564536	TSb=	1.0451727119338310

FATb=	1.7894257279352122E-002 T11/T22=	0.868295550

Alleles sampled from the same population

T11w=	0.73223672899553449	T12w=	1.0953009178070943	T22w=	1.0299328474662610

TTw=	1.0387221389429038	TSw=	1.0001632356191883

FATw=	3.7121480209285296E-002 T11/T22=	0.710955799

Fst=	5.0000000000000003E-002

Scaled migration rate=	18.999999999999996	Migration rate=	9.4999999999999978E-004

Alleles sampled from different populations

T11b=	0.98689906185568399	T12b=	1.1387293529603277	T22b=	1.1000438289218903

TTb=	1.1058757755781472	TSb=	1.0887293522152697

FATb=	1.5504836747069040E-002 T11/T22=	0.897145212

Alleles sampled from the same population

T11w=	0.64707179913860624	T12w=	1.1389925108550645	T22w=	1.0395662693248011

TTw=	1.0535380480983867	TSw=	1.0003168223061816

FATw=	5.0516662296409942E-002 T11/T22=	0.622444034

Fst=	7.5000000000000011E-002

Scaled migration rate=	12.333333333333332	Migration rate=	6.1666666666666662E-004

Alleles sampled from different populations

T11b=	1.0487332071490028	T12b=	1.1818307575789446	T22b=	1.1410638179099848

TTb=	1.1474785609427878	TSb=	1.1318307568338866

FATb=	1.3636685373925150E-002 T11/T22=	0.919083774

Alleles sampled from the same population

T11w=	0.57977803975817688	T12w=	1.1822361629843501	T22w=	1.0472073040508603

TTw=	1.0668382060159616	TSw=	1.0004643776215920

FATw=	6.2215458745369046E-002 T11/T22=	0.553642094

Fst=	0.10000000000000001

Scaled migration rate=	9.0000000000000000	Migration rate=	4.4999999999999999E-004

Alleles sampled from different populations

T11b=	1.1073375644909476	T12b=	1.2252620999000052	T22b=	1.1828092696731694

TTb=	1.1896960620621777	TSb=	1.1752620991549472

FATb=	1.2132479351248060E-002 T11/T22=	0.936192811

Alleles sampled from the same population

T11w=	0.52526516211474261	T12w=	1.2258176554555609	T22w=	1.0534236356667306

TTw=	1.0791729744932002	TSw=	1.0006077883115319

FATw=	7.2801291394981371E-002 T11/T22=	0.498626709

Fst=	0.12500000000000000

Scaled migration rate=	7.0000000000000000	Migration rate=	3.5000000000000000E-004

Alleles sampled from different populations

T11b=	1.1642198205905956	T12b=	1.2696044426623936	T22b=	1.2257582887314178

TTb=	1.2330352117575853	TSb=	1.2196044419173355

FATb=	1.0892446308248926E-002 T11/T22=	0.949795604

Alleles sampled from the same population

T11w=	0.48020816140563233	T12w=	1.2703187283766793	T22w=	1.0585859204414396

TTw=	1.0909140482794248	TSw=	1.0007481445378590

FATw=	8.2651702839260599E-002 T11/T22=	0.453631729

Fst=	0.14999999999999999

Scaled migration rate=	5.6666666666666670	Migration rate=	2.8333333333333335E-004

Alleles sampled from different populations

T11b=	1.2204783583475174	T12b=	1.3153254767995568	T22b=	1.2703084891330523

TTb=	1.2779132456051678	TSb=	1.2653254760544987

FATb=	9.8502536020808051E-003 T11/T22=	0.960773230

Alleles sampled from the same population

T11w=	0.44234323598281999	T12w=	1.3162078297407334	T22w=	1.0629464417693679

TTw=	1.1023274595463484	TSw=	1.0008861211907132

FATw=	9.2024686019689939E-002 T11/T22=	0.416148186

Fst=	0.17500000000000002

Scaled migration rate=	4.7142857142857135	Migration rate=	2.3571428571428569E-004

Alleles sampled from different populations

T11b=	1.2769637031502592	T12b=	1.3628315644538418	T22b=	1.3168168815486196

TTb=	1.3247009926875761	TSb=	1.3128315637087837

FATb=	8.9600815914778753E-003 T11/T22=	0.969735205

Alleles sampled from the same population

T11w=	0.41007574953679243	T12w=	1.3638921705144478	T22w=	1.0666828714849057

TTw=	1.1136144740907423	TSw=	1.0010221592900945

FATw=	0.10110529040364580	T11/T22=	0.384440184

Fst=	0.20000000000000001

Scaled migration rate=	4.0000000000000000	Migration rate=	2.0000000000000001E-004

Alleles sampled from different populations

T11b=	1.3343750466778874	T12b=	1.4125000524520872	T22b=	1.3656250522658226

TTb=	1.3737500522434709	TSb=	1.3625000517070291

FATb=	8.1892630453912485E-003 T11/T22=	0.977116704

Alleles sampled from the same population

T11w=	0.38225001331950942	T12w=	1.4137500524520872	T22w=	1.0699239539938128

TTw=	1.1249359123095592	TSw=	1.0011565599263825

FATw=	0.11003235920262378	T11/T22=	0.357268393

Fst=	0.22500000000000001

Scaled migration rate=	3.4444444444444446	Migration rate=	1.7222222222222224E-004

Alleles sampled from different populations

T11b=	1.3933214948204005	T12b=	1.4647011257684095	T22b=	1.4170766394903458

TTb=	1.4254114955736981	TSb=	1.4147011250233514

FATb=	7.5138797348031083E-003 T11/T22=	0.983236492

Alleles sampled from the same population

T11w=	0.35800798625856733	T12w=	1.4661527386716353	T22w=	1.0727652640572314

TTw=	1.1364274367098375	TSw=	1.0012895362773651

FATw=	0.11891467599877825	T11/T22=	0.333724439

Fst=	0.25000000000000000

Scaled migration rate=	3.0000000000000000	Migration rate=	1.4999999999999999E-004

Alleles sampled from different populations

T11b=	1.4543634274213952	T12b=	1.5198140765479278	T22b=	1.4715308145119226

TTb=	1.4800501278074985	TSb=	1.4698140758028697

FATb=	6.9160171080098198E-003 T11/T22=	0.988333642

Alleles sampled from the same population

T11w=	0.33669925246352062	T12w=	1.5214807432145945	T22w=	1.0752792430802076

TTw=	1.1482097131982305	TSw=	1.0014212440185390

FATw=	0.12784116655033861	T11/T22=	0.313127279

**Gene conversion rate=	9.9999999999999991E-005**

**Scaled value=	400**

Results for equivalent panmictic population

Times are scaled by 2Nd

T11=	0.980540514	T12=	1.00500000	T22=	1.00216222

TT=	1.00245678	TS=	1.00000000

FAT=	2.45076418E-03 T11/T22=	0.978424966

Results for subdivided population

Fst=	2.5000000000000001E-002

Scaled migration rate=	39.000000000000000	Migration rate=	1.9499999999999999E-003

Alleles sampled from different populations

T11b=	1.0321719250317052	T12b=	1.0522142748688621	T22b=	1.0488856471971668

TTb=	1.0493176629564174	TSb=	1.0472142749806208

FATb=	2.0045292765494827E-003 T11/T22=	0.984065235

Alleles sampled from the same population

T11w=	0.82181030767343033	T12w=	1.0523424799970673	T22w=	1.0199802276131615

TTw=	1.0238237338428673	TSw=	1.0001632356191885

FATw=	2.3109933323063658E-002 T11/T22=	0.805711985

Fst=	5.0000000000000003E-002

Scaled migration rate=	18.999999999999996	Migration rate=	9.4999999999999978E-004

Alleles sampled from different populations

T11b=	1.0797019142532269	T12b=	1.0965738839777650	T22b=	1.0928929918491124

TTb=	1.0934236416563112	TSb=	1.0915738840895237

FATb=	1.6917116992143377E-003 T11/T22=	0.987930119

Alleles sampled from the same population

T11w=	0.70787366795078943	T12w=	1.0968370418725018	T22w=	1.0328105061234474

TTw=	1.0410859141765507	TSw=	1.0003168223061816

FATw=	3.9160160862050986E-002 T11/T22=	0.685385823

Fst=	7.5000000000000011E-002

Scaled migration rate=	12.333333333333332	Migration rate=	6.1666666666666662E-004

Alleles sampled from different populations

T11b=	1.1253962589364843	T12b=	1.1398820224649124	T22b=	1.1359359963144697

TTb=	1.1365408836477697	TSb=	1.1348820225766711

FATb=	1.4595700823136770E-003 T11/T22=	0.990721524

Alleles sampled from the same population

T11w=	0.62211435193932341	T12w=	1.1402874278703179	T22w=	1.0425032693640652

TTw=	1.0559005287209433	TSw=	1.0004643776215911

FATw=	5.2501300635301673E-002 T11/T22=	0.596750498

Fst=	0.10000000000000001

Scaled migration rate=	9.0000000000000000	Migration rate=	4.4999999999999999E-004

Alleles sampled from different populations

T11b=	1.1705967831583575	T12b=	1.1832215946533595	T22b=	1.1790687960547583

TTb=	1.1797315796735426	TSb=	1.1782215947651182

FATb=	1.2799393815008386E-003 T11/T22=	0.992814660

Alleles sampled from the same population

T11w=	0.55523005516772628	T12w=	1.1837771502089152	T22w=	1.0500942031052878

TTw=	1.0692084921045653	TSw=	1.0006077883115316

FATw=	6.4160268366372675E-002 T11/T22=	0.528743088

Fst=	0.12500000000000000

Scaled migration rate=	7.0000000000000000	Migration rate=	3.5000000000000000E-004

Alleles sampled from different populations

T11b=	1.2161888996821939	T12b=	1.2273217792744182	T22b=	1.2230032104643973

TTb=	1.2237124097423790	TSb=	1.2223217793861769

FATb=	1.1364029204335990E-003 T11/T22=	0.994428217

Alleles sampled from the same population

T11w=	0.50160719397276110	T12w=	1.2280360649887039	T22w=	1.0562082501562027

TTw=	1.0815912462642185	TSw=	1.0007481445378585

FATw=	7.4744596912733385E-002 T11/T22=	0.474913150

Fst=	0.14999999999999999

Scaled migration rate=	5.6666666666666670	Migration rate=	2.8333333333333335E-004

Alleles sampled from different populations

T11b=	1.2628175963166377	T12b=	1.2727277022104917	T22b=	1.2682732696562071

TTb=	1.2690205107825827	TSb=	1.2677277023222504

FATb=	1.0187451261406411E-003 T11/T22=	0.995698333

Alleles sampled from the same population

T11w=	0.45765742843973589	T12w=	1.2736100551516683	T22w=	1.0612448648297104

TTw=	1.0934347247237632	TSw=	1.0008861211907130

FATw=	8.4640263785687786E-002 T11/T22=	0.431245834

Fst=	0.17500000000000002

Scaled migration rate=	4.7142857142857135	Migration rate=	2.3571428571428569E-004

Alleles sampled from different populations

T11b=	1.3109978869022936	T12b=	1.3198875660674814	T22b=	1.3153197527655673

TTb=	1.3160987405012794	TSb=	1.3148875661792401

FATb=	9.2027618047718640E-004 T11/T22=	0.996714234

Alleles sampled from the same population

T11w=	0.42097990549618208	T12w=	1.3209481721280874	T22w=	1.0654712986005281

TTw=	1.1050122219044454	TSw=	1.0010221592900936

FATw=	9.4107613067961338E-002 T11/T22=	0.395111442

Fst=	0.20000000000000001

Scaled migration rate=	4.0000000000000000	Migration rate=	2.0000000000000001E-004

Alleles sampled from different populations

T11b=	1.3611765220585990	T12b=	1.3692017325718235	T22b=	1.3645378671974693

TTb=	1.3653437495134644	TSb=	1.3642017326835822

FATb=	8.3643172665426313E-004 T11/T22=	0.997536659

Alleles sampled from the same population

T11w=	0.38990757771399837	T12w=	1.3704517325718235	T22w=	1.0690731135055362

TTw=	1.1165296095795527	TSw=	1.0011565599263825

FATw=	0.10333183165345317	T11/T22=	0.364715546

Fst=	0.22500000000000001

Scaled migration rate=	3.4444444444444446	Migration rate=	1.7222222222222224E-004

Alleles sampled from different populations

T11b=	1.4137695802674959	T12b=	1.4210530644452839	T22b=	1.4163067850336588

TTb=	1.4171357432800900	TSb=	1.4160530645570426

FATb=	7.6399083727951567E-004 T11/T22=	0.998208582

Alleles sampled from the same population

T11w=	0.36324675195162487	T12w=	1.4225046773485097	T22w=	1.0721831789802243

TTw=	1.1281516844162298	TSw=	1.0012895362773644

FATw=	0.11245132183134676	T11/T22=	0.338791698

Fst=	0.25000000000000000

Scaled migration rate=	3.0000000000000000	Migration rate=	1.4999999999999999E-004

Alleles sampled from different populations

T11b=	1.4691876030438937	T12b=	1.4758276945370936	T22b=	1.4710099270494033

TTb=	1.4718589019571326	TSb=	1.4708276946488523

FATb=	7.0061560038747750E-004 T11/T22=	0.998761177

Alleles sampled from the same population

T11w=	0.34012021606871257	T12w=	1.4774943612037603	T22w=	1.0748991360129636

TTw=	1.1400184873478647	TSw=	1.0014212440185386

FATw=	0.12157455766507641	T11/T22=	0.316420585

**Inversion frequency=	0.5**

**Gene conversion rate=	9.9999999999999995E-008**

**Scaled value=	0.4**

Results for equivalent panmictic population

Times are scaled by 2Nd

T11=	1.00000000	T12=	6.00000000	T22=	1.00000000

TT=	3.50000000	TS=	1.00000000

FAT=	0.714285731	T11/T22=	1.00000000

Results for subdivided population

Fst=	2.5000000000000001E-002

Scaled migration rate=	39.000000000000000	Migration rate=	1.9499999999999999E-003

Alleles sampled from different populations

T11b=	1.0512820512820513	T12b=	6.0512820512820511	T22b=	1.0512820512820513

TTb=	3.5512820512820511	TSb=	1.0512820512820513

FATb=	0.70397111913357402	T11/T22=	1.00000000

Alleles sampled from the same population

T11w=	1.0002439024390244	T12w=	6.0514102564102563	T22w=	1.0002439024390244

TTw=	3.5258270794246402	TSw=	1.0002439024390244

FATw=	0.71630942757344518	T11/T22=	1.00000000

Fst=	5.0000000000000003E-002

Scaled migration rate=	18.999999999999996	Migration rate=	9.4999999999999978E-004

Alleles sampled from different populations

T11b=	1.1052631578947367	T12b=	6.1052631578947363	T22b=	1.1052631578947367

TTb=	3.6052631578947363	TSb=	1.1052631578947367

FATb=	0.69343065693430650	T11/T22=	1.00000000

Alleles sampled from the same population

T11w=	1.0004761904761903	T12w=	6.1055263157894730	T22w=	1.0004761904761903

TTw=	3.5530012531328317	TSw=	1.0004761904761903

FATw=	0.71841378057662431	T11/T22=	1.00000000

Fst=	7.5000000000000011E-002

Scaled migration rate=	12.333333333333332	Migration rate=	6.1666666666666662E-004

Alleles sampled from different populations

T11b=	1.1621621621621623	T12b=	6.1621621621621623	T22b=	1.1621621621621623

TTb=	3.6621621621621623	TSb=	1.1621621621621623

FATb=	0.68265682656826565	T11/T22=	1.00000000

Alleles sampled from the same population

T11w=	1.0006976744186047	T12w=	6.1625675675675673	T22w=	1.0006976744186047

TTw=	3.5816326209930862	TSw=	1.0006976744186047

FATw=	0.72060292600832432	T11/T22=	1.00000000

Fst=	0.10000000000000001

Scaled migration rate=	9.0000000000000000	Migration rate=	4.4999999999999999E-004

Alleles sampled from different populations

T11b=	1.2222222222222221	T12b=	6.2222222222222223	T22b=	1.2222222222222221

TTb=	3.7222222222222219	TSb=	1.2222222222222221

FATb=	0.67164179104477606	T11/T22=	1.00000000

Alleles sampled from the same population

T11w=	1.0009090909090907	T12w=	6.2227777777777780	T22w=	1.0009090909090907

TTw=	3.6118434343434340	TSw=	1.0009090909090907

FATw=	0.72288137370743000	T11/T22=	1.00000000

Fst=	0.12500000000000000

Scaled migration rate=	7.0000000000000000	Migration rate=	3.5000000000000000E-004

Alleles sampled from different populations

T11b=	1.2857142857142858	T12b=	6.2857142857142856	T22b=	1.2857142857142858

TTb=	3.7857142857142860	TSb=	1.2857142857142858

FATb=	0.66037735849056611	T11/T22=	1.00000000

Alleles sampled from the same population

T11w=	1.0011111111111111	T12w=	6.2864285714285710	T22w=	1.0011111111111111

TTw=	3.6437698412698412	TSw=	1.0011111111111111

FATw=	0.72525402132363359	T11/T22=	1.00000000

Fst=	0.14999999999999999

Scaled migration rate=	5.6666666666666670	Migration rate=	2.8333333333333335E-004

Alleles sampled from different populations

T11b=	1.3529411764705881	T12b=	6.3529411764705879	T22b=	1.3529411764705881

TTb=	3.8529411764705883	TSb=	1.3529411764705881

FATb=	0.64885496183206115	T11/T22=	1.00000000

Alleles sampled from the same population

T11w=	1.0013043478260868	T12w=	6.3538235294117644	T22w=	1.0013043478260868

TTw=	3.6775639386189258	TSw=	1.0013043478260868

FATw=	0.72772618925502153	T11/T22=	1.00000000

Fst=	0.17500000000000002

Scaled migration rate=	4.7142857142857135	Migration rate=	2.3571428571428569E-004

Alleles sampled from different populations

T11b=	1.4242424242424241	T12b=	6.4242424242424239	T22b=	1.4242424242424241

TTb=	3.9242424242424239	TSb=	1.4242424242424241

FATb=	0.63706563706563712	T11/T22=	1.00000000

Alleles sampled from the same population

T11w=	1.0014893617021274	T12w=	6.4253030303030298	T22w=	1.0014893617021274

TTw=	3.7133961960025785	TSw=	1.0014893617021274

FATw=	0.73030366035807937	T11/T22=	1.00000000

Fst=	0.20000000000000001

Scaled migration rate=	4.0000000000000000	Migration rate=	2.0000000000000001E-004

Alleles sampled from different populations

T11b=	1.5000000000000000	T12b=	6.5000000000000000	T22b=	1.5000000000000000

TTb=	4.0000000000000000	TSb=	1.5000000000000000

FATb=	0.62500000000000000	T11/T22=	1.00000000

Alleles sampled from the same population

T11w=	1.0016666666666667	T12w=	6.5012499999999998	T22w=	1.0016666666666667

TTw=	3.7514583333333333	TSw=	1.0016666666666667

FATw=	0.73299272505136892	T11/T22=	1.00000000

Fst=	0.22500000000000001

Scaled migration rate=	3.4444444444444446	Migration rate=	1.7222222222222224E-004

Alleles sampled from different populations

T11b=	1.5806451612903225	T12b=	6.5806451612903221	T22b=	1.5806451612903225

TTb=	4.0806451612903221	TSb=	1.5806451612903225

FATb=	0.61264822134387353	T11/T22=	1.00000000

Alleles sampled from the same population

T11w=	1.0018367346938775	T12w=	6.5820967741935474	T22w=	1.0018367346938775

TTw=	3.7919667544437128	TSw=	1.0018367346938775

FATw=	0.73580023255218430	T11/T22=	1.00000000

Fst=	0.25000000000000000

Scaled migration rate=	3.0000000000000000	Migration rate=	1.4999999999999999E-004

Alleles sampled from different populations

T11b=	1.6666666666666667	T12b=	6.6666666666666670	T22b=	1.6666666666666667

TTb=	4.1666666666666670	TSb=	1.6666666666666667

FATb=	0.60000000000000009	T11/T22=	1.00000000

Alleles sampled from the same population

T11w=	1.0020000000000000	T12w=	6.6683333333333339	T22w=	1.0020000000000000

TTw=	3.8351666666666673	TSw=	1.0020000000000000

FATw=	0.73873364912433193	T11/T22=	1.00000000

**Gene conversion rate=	9.9999999999999995E-007**

**Scaled value=	4**

Results for equivalent panmictic population

Times are scaled by 2Nd

T11=	1.00000000	T12=	1.50000000	T22=	1.00000000

TT=	1.25000000	TS=	1.00000000

FAT=	0.199999988	T11/T22=	1.00000000

Results for subdivided population

Fst=	2.5000000000000001E-002

Scaled migration rate=	39.000000000000000	Migration rate=	1.9499999999999999E-003

Alleles sampled from different populations

T11b=	1.0512820512820513	T12b=	1.5512820512820513	T22b=	1.0512820512820513

TTb=	1.3012820512820513	TSb=	1.0512820512820513

FATb=	0.19211822660098521	T11/T22=	1.00000000

Alleles sampled from the same population

T11w=	1.0002439024390244	T12w=	1.5514102564102565	T22w=	1.0002439024390244

TTw=	1.2758270794246405	TSw=	1.0002439024390244

FATw=	0.21600354893697327	T11/T22=	1.00000000

Fst=	5.0000000000000003E-002

Scaled migration rate=	18.999999999999996	Migration rate=	9.4999999999999978E-004

Alleles sampled from different populations

T11b=	1.1052631578947369	T12b=	1.6052631578947369	T22b=	1.1052631578947369

TTb=	1.3552631578947369	TSb=	1.1052631578947369

FATb=	0.18446601941747576	T11/T22=	1.00000000

Alleles sampled from the same population

T11w=	1.0004761904761905	T12w=	1.6055263157894737	T22w=	1.0004761904761905

TTw=	1.3030012531328321	TSw=	1.0004761904761905

FATw=	0.23217557306969161	T11/T22=	1.00000000

Fst=	7.5000000000000011E-002

Scaled migration rate=	12.333333333333332	Migration rate=	6.1666666666666662E-004

Alleles sampled from different populations

T11b=	1.1621621621621621	T12b=	1.6621621621621621	T22b=	1.1621621621621621

TTb=	1.4121621621621621	TSb=	1.1621621621621621

FATb=	0.17703349282296654	T11/T22=	1.00000000

Alleles sampled from the same population

T11w=	1.0006976744186045	T12w=	1.6625675675675675	T22w=	1.0006976744186045

TTw=	1.3316326209930860	TSw=	1.0006976744186045

FATw=	0.24851820341235076	T11/T22=	1.00000000

Fst=	0.10000000000000001

Scaled migration rate=	9.0000000000000000	Migration rate=	4.4999999999999999E-004

Alleles sampled from different populations

T11b=	1.2222222222222221	T12b=	1.7222222222222221	T22b=	1.2222222222222221

TTb=	1.4722222222222221	TSb=	1.2222222222222221

FATb=	0.16981132075471694	T11/T22=	1.00000000

Alleles sampled from the same population

T11w=	1.0009090909090907	T12w=	1.7227777777777777	T22w=	1.0009090909090907

TTw=	1.3618434343434342	TSw=	1.0009090909090907

FATw=	0.26503365536167933	T11/T22=	1.00000000

Fst=	0.12500000000000000

Scaled migration rate=	7.0000000000000000	Migration rate=	3.5000000000000000E-004

Alleles sampled from different populations

T11b=	1.2857142857142860	T12b=	1.7857142857142860	T22b=	1.2857142857142860

TTb=	1.5357142857142860	TSb=	1.2857142857142860

FATb=	0.16279069767441856	T11/T22=	1.00000000

Alleles sampled from the same population

T11w=	1.0011111111111113	T12w=	1.7864285714285717	T22w=	1.0011111111111113

TTw=	1.3937698412698416	TSw=	1.0011111111111113

FATw=	0.28172422629046501	T11/T22=	1.00000000

Fst=	0.14999999999999999

Scaled migration rate=	5.6666666666666670	Migration rate=	2.8333333333333335E-004

Alleles sampled from different populations

T11b=	1.3529411764705879	T12b=	1.8529411764705879	T22b=	1.3529411764705879

TTb=	1.6029411764705879	TSb=	1.3529411764705879

FATb=	0.15596330275229364	T11/T22=	1.00000000

Alleles sampled from the same population

T11w=	1.0013043478260868	T12w=	1.8538235294117644	T22w=	1.0013043478260868

TTw=	1.4275639386189258	TSw=	1.0013043478260868

FATw=	0.29859229367002449	T11/T22=	1.00000000

Fst=	0.17500000000000002

Scaled migration rate=	4.7142857142857135	Migration rate=	2.3571428571428569E-004

Alleles sampled from different populations

T11b=	1.4242424242424243	T12b=	1.9242424242424243	T22b=	1.4242424242424243

TTb=	1.6742424242424243	TSb=	1.4242424242424243

FATb=	0.14932126696832582	T11/T22=	1.00000000

Alleles sampled from the same population

T11w=	1.0014893617021277	T12w=	1.9253030303030303	T22w=	1.0014893617021277

TTw=	1.4633961960025790	TSw=	1.0014893617021277

FATw=	0.31564031364998657	T11/T22=	1.00000000

Fst=	0.20000000000000001

Scaled migration rate=	4.0000000000000000	Migration rate=	2.0000000000000001E-004

Alleles sampled from different populations

T11b=	1.5000000000000004	T12b=	2.0000000000000004	T22b=	1.5000000000000004

TTb=	1.7500000000000004	TSb=	1.5000000000000004

FATb=	0.14285714285714279	T11/T22=	1.00000000

Alleles sampled from the same population

T11w=	1.0016666666666669	T12w=	2.0012500000000006	T22w=	1.0016666666666669

TTw=	1.5014583333333338	TSw=	1.0016666666666669

FATw=	0.33287082003607604	T11/T22=	1.00000000

Fst=	0.22500000000000001

Scaled migration rate=	3.4444444444444446	Migration rate=	1.7222222222222224E-004

Alleles sampled from different populations

T11b=	1.5806451612903225	T12b=	2.0806451612903225	T22b=	1.5806451612903225

TTb=	1.8306451612903225	TSb=	1.5806451612903225

FATb=	0.13656387665198233	T11/T22=	1.00000000

Alleles sampled from the same population

T11w=	1.0018367346938775	T12w=	2.0820967741935483	T22w=	1.0018367346938775

TTw=	1.5419667544437128	TSw=	1.0018367346938775

FATw=	0.35028642361663320	T11/T22=	1.00000000

Fst=	0.25000000000000000

Scaled migration rate=	3.0000000000000000	Migration rate=	1.4999999999999999E-004

Alleles sampled from different populations

T11b=	1.6666666666666665	T12b=	2.1666666666666665	T22b=	1.6666666666666665

TTb=	1.9166666666666665	TSb=	1.6666666666666665

FATb=	0.13043478260869568	T11/T22=	1.00000000

Alleles sampled from the same population

T11w=	1.0020000000000000	T12w=	2.1683333333333330	T22w=	1.0020000000000000

TTw=	1.5851666666666664	TSw=	1.0020000000000000

FATw=	0.36788981179686664	T11/T22=	1.00000000

**Gene conversion rate=	9.9999999999999991E-006**

**Scaled value=	40**

Results for equivalent panmictic population

Times are scaled by 2Nd

T11=	1.00000000	T12=	1.04999995	T22=	1.00000000

TT=	1.02499998	TS=	1.00000000

FAT=	2.43902206E-02 T11/T22=	1.00000000

Results for subdivided population

Fst=	2.5000000000000001E-002

Scaled migration rate=	39.000000000000000	Migration rate=	1.9499999999999999E-003

Alleles sampled from different populations

T11b=	1.0512820512820511	T12b=	1.1012820520271092	T22b=	1.0512820512820511

TTb=	1.0762820516545801	TSb=	1.0512820512820511

FATb=	2.3228112309497506E-002 T11/T22=	1.00000000

Alleles sampled from the same population

T11w=	1.0002439024390242	T12w=	1.1014102571553144	T22w=	1.0002439024390242

TTw=	1.0508270797971693	TSw=	1.0002439024390242

FATw=	4.8136537714567385E-002 T11/T22=	1.00000000

Fst=	5.0000000000000003E-002

Scaled migration rate=	18.999999999999996	Migration rate=	9.4999999999999978E-004

Alleles sampled from different populations

T11b=	1.1052631578947367	T12b=	1.1552631586397948	T22b=	1.1052631578947367

TTb=	1.1302631582672658	TSb=	1.1052631578947367

FATb=	2.2118743046402489E-002 T11/T22=	1.00000000

Alleles sampled from the same population

T11w=	1.0004761904761903	T12w=	1.1555263165345315	T22w=	1.0004761904761903

TTw=	1.0780012535053609	TSw=	1.0004761904761903

FATw=	7.1915559260326112E-002 T11/T22=	1.00000000

Fst=	7.5000000000000011E-002

Scaled migration rate=	12.333333333333332	Migration rate=	6.1666666666666662E-004

Alleles sampled from different populations

T11b=	1.1621621621621629	T12b=	1.2121621629072210	T22b=	1.1621621621621629

TTb=	1.1871621625346920	TSb=	1.1621621621621629

FATb=	2.1058622959437945E-002 T11/T22=	1.00000000

Alleles sampled from the same population

T11w=	1.0006976744186054	T12w=	1.2125675683126265	T22w=	1.0006976744186054

TTw=	1.1066326213656159	TSw=	1.0006976744186054

FATw=	9.5727294588771383E-002 T11/T22=	1.00000000

Fst=	0.10000000000000001

Scaled migration rate=	9.0000000000000000	Migration rate=	4.4999999999999999E-004

Alleles sampled from different populations

T11b=	1.2222222222222221	T12b=	1.2722222229672802	T22b=	1.2222222222222221

TTb=	1.2472222225947511	TSb=	1.2222222222222221

FATb=	2.0044543722544050E-002 T11/T22=	1.00000000

Alleles sampled from the same population

T11w=	1.0009090909090907	T12w=	1.2727777785228358	T22w=	1.0009090909090907

TTw=	1.1368434347159633	TSw=	1.0009090909090907

FATw=	0.11957173666647891	T11/T22=	1.00000000

Fst=	0.12500000000000000

Scaled migration rate=	7.0000000000000000	Migration rate=	3.5000000000000000E-004

Alleles sampled from different populations

T11b=	1.2857142857142860	T12b=	1.3357142864593441	T22b=	1.2857142857142860

TTb=	1.3107142860868151	TSb=	1.2857142857142860

FATb=	1.9073569761086095E-002 T11/T22=	1.00000000

Alleles sampled from the same population

T11w=	1.0011111111111113	T12w=	1.3364285721736298	T22w=	1.0011111111111113

TTw=	1.1687698416423706	TSw=	1.0011111111111113

FATw=	0.14344888493671526	T11/T22=	1.00000000

Fst=	0.14999999999999999

Scaled migration rate=	5.6666666666666670	Migration rate=	2.8333333333333335E-004

Alleles sampled from different populations

T11b=	1.3529411764705885	T12b=	1.4029411772156466	T22b=	1.3529411764705885

TTb=	1.3779411768431176	TSb=	1.3529411764705885

FATb=	1.8143009870569604E-002 T11/T22=	1.00000000

Alleles sampled from the same population

T11w=	1.0013043478260872	T12w=	1.4038235301568232	T22w=	1.0013043478260872

TTw=	1.2025639389914553	TSw=	1.0013043478260872

FATw=	0.16735874462871125	T11/T22=	1.00000000

Fst=	0.17500000000000002

Scaled migration rate=	4.7142857142857135	Migration rate=	2.3571428571428569E-004

Alleles sampled from different populations

T11b=	1.4242424242424243	T12b=	1.4742424249874824	T22b=	1.4242424242424243

TTb=	1.4492424246149533	TSb=	1.4242424242424243

FATb=	1.7250392306981555E-002 T11/T22=	1.00000000

Alleles sampled from the same population

T11w=	1.0014893617021277	T12w=	1.4753030310480884	T22w=	1.0014893617021277

TTw=	1.2383961963751080	TSw=	1.0014893617021277

FATw=	0.19130132615589990	T11/T22=	1.00000000

Fst=	0.20000000000000001

Scaled migration rate=	4.0000000000000000	Migration rate=	2.0000000000000001E-004

Alleles sampled from different populations

T11b=	1.4999999999999996	T12b=	1.5500000007450576	T22b=	1.4999999999999996

TTb=	1.5250000003725286	TSb=	1.4999999999999996

FATb=	1.6393442863227525E-002 T11/T22=	1.00000000

Alleles sampled from the same population

T11w=	1.0016666666666663	T12w=	1.5512500007450576	T22w=	1.0016666666666663

TTw=	1.2764583337058619	TSw=	1.0016666666666663

FATw=	0.21527664459004325	T11/T22=	1.00000000

Fst=	0.22500000000000001

Scaled migration rate=	3.4444444444444446	Migration rate=	1.7222222222222224E-004

Alleles sampled from different populations

T11b=	1.5806451612903223	T12b=	1.6306451620353803	T22b=	1.5806451612903223

TTb=	1.6056451616628513	TSb=	1.5806451612903223

FATb=	1.5570065522221843E-002 T11/T22=	1.00000000

Alleles sampled from the same population

T11w=	1.0018367346938772	T12w=	1.6320967749386062	T22w=	1.0018367346938772

TTw=	1.3169667548162416	TSw=	1.0018367346938772

FATw=	0.23928471920032246	T11/T22=	1.00000000

Fst=	0.25000000000000000

Scaled migration rate=	3.0000000000000000	Migration rate=	1.4999999999999999E-004

Alleles sampled from different populations

T11b=	1.6666666666666676	T12b=	1.7166666674117257	T22b=	1.6666666666666676

TTb=	1.6916666670391967	TSb=	1.6666666666666676

FATb=	1.4778325340112475E-002 T11/T22=	1.00000000

Alleles sampled from the same population

T11w=	1.0020000000000004	T12w=	1.7183333340783924	T22w=	1.0020000000000004

TTw=	1.3601666670391965	TSw=	1.0020000000000004

FATw=	0.26332557304822901	T11/T22=	1.00000000

**Gene conversion rate=	9.9999999999999991E-005**

**Scaled value=	400**

Results for equivalent panmictic population

Times are scaled by 2Nd

T11=	1.00000000	T12=	1.00500000	T22=	1.00000000

TT=	1.00250006	TS=	1.00000000

FAT=	2.49379873E-03 T11/T22=	1.00000000

Results for subdivided population

Fst=	2.5000000000000001E-002

Scaled migration rate=	39.000000000000000	Migration rate=	1.9499999999999999E-003

Alleles sampled from different populations

T11b=	1.0512820512820491	T12b=	1.0562820511702904	T22b=	1.0512820512820491

TTb=	1.0537820512261697	TSb=	1.0512820512820491

FATb=	2.3724070278210441E-003 T11/T22=	1.00000000

Alleles sampled from the same population

T11w=	1.0002439024390222	T12w=	1.0564102562984956	T22w=	1.0002439024390222

TTw=	1.0283270793687589	TSw=	1.0002439024390222

FATw=	2.7309576391760126E-002 T11/T22=	1.00000000

Fst=	5.0000000000000003E-002

Scaled migration rate=	18.999999999999996	Migration rate=	9.4999999999999978E-004

Alleles sampled from different populations

T11b=	1.1052631578947376	T12b=	1.1102631577829789	T22b=	1.1052631578947376

TTb=	1.1077631578388583	TSb=	1.1052631578947376

FATb=	2.2568000446936409E-003 T11/T22=	1.00000000

Alleles sampled from the same population

T11w=	1.0004761904761912	T12w=	1.1105263156777156	T22w=	1.0004761904761912

TTw=	1.0555012530769534	TSw=	1.0004761904761912

FATw=	5.2131688560629774E-002 T11/T22=	1.00000000

Fst=	7.5000000000000011E-002

Scaled migration rate=	12.333333333333332	Migration rate=	6.1666666666666662E-004

Alleles sampled from different populations

T11b=	1.1621621621621583	T12b=	1.1671621620503996	T22b=	1.1621621621621583

TTb=	1.1646621621062789	TSb=	1.1621621621621583

FATb=	2.1465451746104369E-003 T11/T22=	1.00000000

Alleles sampled from the same population

T11w=	1.0006976744186014	T12w=	1.1675675674558050	T22w=	1.0006976744186014

TTw=	1.0841326209372033	TSw=	1.0006976744186014

FATw=	7.6960092250037304E-002 T11/T22=	1.00000000

Fst=	0.10000000000000001

Scaled migration rate=	9.0000000000000000	Migration rate=	4.4999999999999999E-004

Alleles sampled from different populations

T11b=	1.2222222222222190	T12b=	1.2272222221104603	T22b=	1.2222222222222190

TTb=	1.2247222221663396	TSb=	1.2222222222222190

FATb=	2.0412791560999732E-003 T11/T22=	1.00000000

Alleles sampled from the same population

T11w=	1.0009090909090883	T12w=	1.2277777776660159	T22w=	1.0009090909090883

TTw=	1.1143434342875520	TSw=	1.0009090909090883

FATw=	0.10179477878019461	T11/T22=	1.00000000

Fst=	0.12500000000000000

Scaled migration rate=	7.0000000000000000	Migration rate=	3.5000000000000000E-004

Alleles sampled from different populations

T11b=	1.2857142857142825	T12b=	1.2907142856025238	T22b=	1.2857142857142825

TTb=	1.2882142856584031	TSb=	1.2857142857142825

FATb=	1.9406708743668899E-003 T11/T22=	1.00000000

Alleles sampled from the same population

T11w=	1.0011111111111086	T12w=	1.2914285713168094	T22w=	1.0011111111111086

TTw=	1.1462698412139591	TSw=	1.0011111111111086

FATw=	0.12663574045455117	T11/T22=	1.00000000

Fst=	0.14999999999999999

Scaled migration rate=	5.6666666666666670	Migration rate=	2.8333333333333335E-004

Alleles sampled from different populations

T11b=	1.3529411764705910	T12b=	1.3579411763588323	T22b=	1.3529411764705910

TTb=	1.3554411764147116	TSb=	1.3529411764705910

FATb=	1.8444178822525359E-003 T11/T22=	1.00000000

Alleles sampled from the same population

T11w=	1.0013043478260890	T12w=	1.3588235293000088	T22w=	1.0013043478260890

TTw=	1.1800639385630489	TSw=	1.0013043478260890

FATw=	0.15148297045212100	T11/T22=	1.00000000

Fst=	0.17500000000000002

Scaled migration rate=	4.7142857142857135	Migration rate=	2.3571428571428569E-004

Alleles sampled from different populations

T11b=	1.4242424242424292	T12b=	1.4292424241306705	T22b=	1.4242424242424292

TTb=	1.4267424241865498	TSb=	1.4242424242424292

FATb=	1.7522433634410417E-003 T11/T22=	1.00000000

Alleles sampled from the same population

T11w=	1.0014893617021310	T12w=	1.4303030301912765	T22w=	1.0014893617021310

TTw=	1.2158961959467036	TSw=	1.0014893617021310

FATw=	0.17633646273367465	T11/T22=	1.00000000

Fst=	0.20000000000000001

Scaled migration rate=	4.0000000000000000	Migration rate=	2.0000000000000001E-004

Alleles sampled from different populations

T11b=	1.5000000000000056	T12b=	1.5049999998882468	T22b=	1.5000000000000056

TTb=	1.5024999999441262	TSb=	1.5000000000000056

FATb=	1.6638934736862288E-003 T11/T22=	1.00000000

Alleles sampled from the same population

T11w=	1.0016666666666703	T12w=	1.5062499998882468	T22w=	1.0016666666666703

TTw=	1.2539583332774586	TSw=	1.0016666666666703

FATw=	0.20119621195975157	T11/T22=	1.00000000

Fst=	0.22500000000000001

Scaled migration rate=	3.4444444444444446	Migration rate=	1.7222222222222224E-004

Alleles sampled from different populations

T11b=	1.5806451612903316	T12b=	1.5856451611785729	T22b=	1.5806451612903316

TTb=	1.5831451612344523	TSb=	1.5806451612903316

FATb=	1.5791350062752674E-003 T11/T22=	1.00000000

Alleles sampled from the same population

T11w=	1.0018367346938832	T12w=	1.5870967740817987	T22w=	1.0018367346938832

TTw=	1.2944667543878410	TSw=	1.0018367346938832

FATw=	0.22606221341879407	T11/T22=	1.00000000

Fst=	0.25000000000000000

Scaled migration rate=	3.0000000000000000	Migration rate=	1.4999999999999999E-004

Alleles sampled from different populations

T11b=	1.6666666666666630	T12b=	1.6716666665549043	T22b=	1.6666666666666630

TTb=	1.6691666666107836	TSb=	1.6666666666666630

FATb=	1.4977533365178042E-003 T11/T22=	1.00000000

Alleles sampled from the same population

T11w=	1.0019999999999978	T12w=	1.6733333332215710	T22w=	1.0019999999999978

TTw=	1.3376666666107844	TSw=	1.0019999999999978

FATw=	0.25093446296397415	T11/T22=	1.00000000

**Table S2**

**Neutral coalescent times for inversion and standard arrangements in a coalescent simulation of approach to recombination-drift equilibrium in a single population.**

**Times scaled by 2 x population size**

The italic numbers in brackets are the *T_ij_* values from the iterations of Equations (A3)

The standard deviations for the equilibrium results were obtained by the procedure described in section 8 of the Supplementary Information. These can be compared with the first set of results from the coalescent simulations, where the time from the initial state is chosen to be close to equilibrium.

For an inversion frequency of 0.5, only the equilibrium results are displayed.

**Inversion frequency=	0.1**

**Scaled recombination rate=	40**

**Number of replicates=	1000000**

Initial state of population

T11=	0.00000000	T12=	1.00000000	T22b=	1.00000000

Equilibrium results for panmictic population

T11=	0.843478322	T12=	1.04999995	T22=	1.01739132

s.d.= 1.00314	s.d.= 1.02252	s.d.= 1.02181

Time to origin of inversion=	4

Mean of T11=	0.842840254 (*0.84298*)	s.d.=	1.00316882

s.e.=	1.00316887E-03

Mean of T12=	1.04935372 (*1.04937*)	s.d.=	1.01939905

s.e.=	1.01939903E-03

Mean of T22=	1.01615655 (*1.01679*)	s.d.=	1.01933658

s.e.=	1.01933663E-03

Time to origin of inversion=	2

Mean of T11=	0.839714766 (*0.83988*)	s.d.=	0.992495000

s.e.=	9.92494985E-04

Mean of T12=	1.04427683 (*1.04550*)	s.d.=	1.00812614

s.e.=	1.00812619E-03

Mean of T22=	1.01287293 (*1.01304*)	s.d.=	1.00943804

s.e.=	1.00943807E-03

Time to origin of inversion=	1

Mean of T11=	0.832606018 (*0.83413*)	s.d.=	0.983978033

s.e.=	9.83978040E-04

Mean of T12=	1.03788888 (*1.03832*)	s.d.=	1.00350165

s.e.=	1.00350170E-03

Mean of T22=	1.00672770 (*1.00608*)	s.d.=	1.00149477

s.e.=	1.00149482E-03

Time to origin of inversion=	0.5

Mean of T11=	0.827529371 (*0.82824*)	s.d.=	0.982901752

s.e.=	9.82901780E-04

Mean of T12=	1.03096688 (*1.03094*)	s.d.=	1.00062251

s.e.=	1.00062252E-03

Mean of T22=	0.998158276 (*0.99895*)	s.d.=	0.997037828

s.e.=	9.97037860E-04

Time to origin of inversion=	0.25

Mean of T11=	0.822732747 (*0.82352*)	s.d.=	0.980459869

s.e.=	9.80459852E-04

Mean of T12=	1.02539289 (*1.02532*)	s.d.=	0.998387218

s.e.=	9.98387230E-04

Mean of T22=	0.993745387 (*0.99392*)	s.d.=	0.998511076

s.e.=	9.98511096E-04

Time to origin of inversion=	0.125

Mean of T11=	0.811701477 (*0.81268*)	s.d.=	0.976616263

s.e.=	9.76616284E-04

Mean of T12=	1.01626980 (*1.01772*)	s.d.=	0.994884372

s.e.=	9.94884409E-04

Mean of T22=	0.991116047 (*0.99480*)	s.d.=	0.998467326

s.e.=	9.98467323E-04

Time to origin of inversion=	6.25E-02

Mean of T11=	0.758589447 (*0.758545*)	s.d.=	0.961394668

s.e.=	9.61394690E-04

Mean of T12=	1.00485098 (*1.00452*)	s.d.=	0.997913659

s.e.=	9.97913652E-04

Mean of T22=	0.993107736 (*0.99345*)	s.d.=	0.998382032

s.e.=	9.98381991E-04

**Inversion frequency=	0.1**

**Scaled recombination rate=	4.0**

**Number of replicates=	1000000**

Initial state of population

T11=	0.00000000	T12=	1.00000000	T22b=	1.00000000

Equilibrium results for panmictic population

T11=	0.470588297	T12=	1.50000000	T22=	1.05882359

s.d.= 0.91885	s.d.= 1.23073	s.d.= 1.12996

Time to origin of inversion=	20

Mean of T11=	0.469264179	s.d.=	0.915292263

s.e.=	9.15292243E-04

Mean of T12=	1.50086200	s.d.=	1.23265827

s.e.=	1.23265828E-03

Mean of T22=	1.05968940	s.d.=	1.13081312

Time to origin of inversion=	1

Mean of T11=	0.434916764 (*0.43413*)	s.d.=	0.809637189

s.e.=	8.09637189E-04

Mean of T12=	1.37598729 (*1.37391*)	s.d.=	1.05607760

s.e.=	1.05607766E-03

Mean of T22=	1.00097787 (*1.00080*)	s.d.=	1.01011753

Time to origin of inversion=	0.5

Mean of T11=	0.401043385 (*0.40122*)	s.d.=	0.756564558

s.e.=	7.56564550E-04

Mean of T12=	1.26687658 (*1.26656*)	s.d.=	1.01439202

s.e.=	1.01439201E-03

Mean of T22=	0.984599292 (*0.98489*)	s.d.=	0.996770203

s.e.=	9.96770221E-04

Time to origin of inversion=	0.25

Mean of T11=	0.359776527 (*0.35102*)	s.d.=	0.706095636

s.e.=	7.06095656E-04

Mean of T12=	1.16544998 (*1.15114*)	s.d.=	1.00426865

s.e.=	1.00426865E-03

Mean of T22=	0.983902514 (*0.985288*)	s.d.=	0.997553110

s.e.=	9.97553114E-04

Time to origin of inversion=	0.125

Mean of T11=	0.288712919 (*0.28931*)	s.d.=	0.632238030

s.e.=	6.32238050E-04

Mean of T12=	1.09267497 (*1.09138*)	s.d.=	1.00096273

s.e.=	1.00096269E-03

Mean of T22=	0.988179803 (*0.98956*)	s.d.=	0.996701419

s.e.=	9.96701419E-04

Time to origin of inversion=	6.25E-02

Mean of T11=	0.198425397 (*0.19780*)	s.d.=	0.533404410

s.e.=	5.33404411E-04

Mean of T12=	1.04694235 (*1.04796*)	s.d.=	0.999398291

s.e.=	9.99398297E-04

Mean of T22=	0.993636668 (*0.99397*)	s.d.=	0.999168098

s.e.=	9.99168144E-04

**Inversion frequency=	0.1**

**Scaled recombination rate=	0.4**

**Number of replicates=	1000000**

Initial state of population

T11=	0.00000000	T12=	1.00000000	T22b=	1.00000000

Equilibrium results for panmictic population

T11=	0.305019408	T12=	6.00000000	T22=	1.07722008

s.d.= 1.47875	s.d.= 5.28310	s.d.= 1.71233

Time to origin of inversion=	100

Mean of T11=	0.305183679	s.d.=	1.48238468

s.e.=	1.48238463E-03

Mean of T12=	6.00524044	s.d.=	5.29317570

s.e.=	5.29317558E-03

Mean of T22=	1.07521260	s.d.=	1.70390737

s.e.=	1.70390739E-03

Time to origin of inversion=	4

Mean of T11=	0.221483573 (*0.22321*)	s.d.=	0.742023706

s.e.=	7.42023694E-04

Mean of T12=	3.68989038 (*3.68930*)	s.d.=	1.76187861

s.e.=	1.76187861E-03

Mean of T22=	0.981181800 (*0.98215*)	s.d.=	1.08416986

s.e.=	1.08416984E-03

Time to origin of inversion=	2

Mean of T11=	0.184408382 (*0.18488*)	s.d.=	0.520189285

s.e.=	5.20189293E-04

Mean of T12=	2.60776067 (*2.60694*)	s.d.=	1.18764746

s.e.=	1.18764746E-03

Mean of T22=	0.947988510 (*0.94868*)	s.d.=	0.982948661

s.e.=	9.82948695E-04

Time to origin of inversion=	1

Mean of T11=	0.159522906 (*0.15929*)	s.d.=	0.393521845

s.e.=	3.93521856E-04

Mean of T12=	1.88445938 (*1.88484*)	s.d.=	1.03488922

s.e.=	1.03488925E-03

Mean of T22=	0.946950376 (*0.94684*)	s.d.=	0.978458345

s.e.=	9.78458324E-04

Time to origin of inversion=	0.5

Mean of T11=	0.143433765 (*0.143705*)	s.d.=	0.326370001

s.e.=	3.26370005E-04

Mean of T12=	1.46712434 (*1.46544*)	s.d.=	1.00711167

s.e.=	1.00711163E-03

Mean of T22=	0.961967230 (*0.96170*)	s.d.=	0.992745221

s.e.=	9.92745277E-04

Time to origin of inversion=	0.25

Mean of T11=	0.126929134 (*0.12679*)	s.d.=	0.280853540

s.e.=	2.80853536E-04

Mean of T12=	1.24095392 (*1.23881*)	s.d.=	1.00302839

s.e.=	1.00302836E-03

Mean of T22=	0.976458669 (*0.97696*)	s.d.=	0.997133672

s.e.=	9.97133669E-04

Time to origin of inversion=	0.125

Mean of T11=	9.67181697E-02 (*0.097163*) s.d.=	0.231224895

s.e.=	2.31224898E-04

Mean of T12=	1.12050974 (*1.12093*)	s.d.=	1.00053322

s.e.=	1.00053323E-03

Mean of T22=	0.986687541 (*0.98736*)	s.d.=	0.996315181

s.e.=	9.96315153E-04

Time to origin of inversion=	6.25E-02

Mean of T11=	6.32091165E-02 (0.06311) s.d.=	0.183888689

s.e.=	1.83888682E-04

Mean of T12=	1.06051600 (*1.06083*)	s.d.=	0.997987986

s.e.=	9.97988041E-04

Mean of T22=	0.992533922 (*0.99337*)	s.d.=	0.998949587

s.e.=	9.98949632E-04

**Inversion frequency=	0.5**

**Scaled recombination rate=	40**

**Number of replicates=	1000000**

Initial state of population

T11=	0.00000000	T12=	1.00000000	T22b=	1.00000000

Equilibrium results

T11=	1.00000000	T12=	1.04999995	T22=	1.00000000

s.d. = 1.02470	s.d. = 1.02590	s.d. = 1.02470

Time to origin of inversion=	10

Mean of T11=	1.00014043	s.d.=	1.02391338

s.e.=	1.02391338E-03

Mean of T12=	1.05008972	s.d.=	1.02473772

s.e.=	1.02473772E-03

Mean of T22=	1.00084615	s.d.=	1.02373850

s.e.=	1.02373853E-03

**Scaled recombination rate=	4**

**Number of replicates=	1000000**

Initial state of population

T11=	0.00000000	T12=	1.00000000	T22b=	1.00000000

Equilibrium results

T11=	1.00000000	T12=	1.50000000	T22=	1.00000000

s.d. = 1.22474	s.d. = 1.32288	s.d. = 1.22474

Time to origin of inversion=	20.0000000

Mean of T11=	1.00003147	s.d.=	1.22361922

s.e.=	1.22361921E-03

Mean of T12=	1.49854517	s.d.=	1.32191551

s.e.=	1.32191554E-03

Mean of T22=	0.998929322	s.d.=	1.22275007

**Scaled recombination rate=	0.4**

**Number of replicates=	1000000**

Initial state of population

T11=	0.00000000	T12=	1.00000000	T22b=	1.00000000

Equilibrium results

T11=	1.00000000	T12=	6.00000000	T22=	1.00000000

s.d. =	2.44949	s.d. =	5.56776	s.d. = 2.44949

Time to origin of inversion=	100.000000

Mean of T11=	0.999278069	s.d.=	2.44362330

s.e.=	2.44362326E-03

Mean of T12=	6.01205254	s.d.=	5.57754183

s.e.=	5.57754189E-03

Mean of T22=	1.00279987	s.d.=	2.45711279

s.e.=	2.45711277E-03

**Table S3**

**Coalescent times for inversion and standard arrangements in a panmictic population**

**Time course of approach to equilibrium: instantaneous sweep to equilibrium frequency**

**Times scaled by 2x population size**

**Inversion frequency = 0.1**

**Initial state of population**

T11=	0	T12=	1	T22=	1

TT=	0.99	TS=	0.9

FAT=	0.0909 T11/T22=	0

**Scaled recombination rate=	40**

Equilibrium results for panmictic population

T11=	0.84347826086956512	T12=	1.0500000000000000	T22=	1.0173913043478262

TT=	1.0215217391304350	TS=	1.0000000000000002

FAT=	2.1068312406895107E-002 T11/T22=	0.829059839

Time (relative to 2Ne) =	2.49999994E-03

T11=	8.5726814840451304E-002 T12=	0.99780845886065672	T22=	0.99971854494364520

TT=	0.99023481214767539	TS=	0.90831937193332579

FAT=	8.2723248273494754E-002 T11/T22=	8.57509524E-02

Time (relative to 2Ne) =	4.99999989E-03

T11=	0.16361435090221210	T12=	0.99606660101951916	T22=	0.99941541048695215

TT=	0.99045461418696679	TS=	0.91583530452847817

FAT=	7.5338444174689712E-002 T11/T22=	0.163710058

Time (relative to 2Ne) =	7.49999983E-03

T11=	0.23290197718569094	T12=	0.99475505471399739	T22=	0.99910096911829749

TT=	0.99065671460619742	TS=	0.92248106992503687

FAT=	6.8818636845621617E-002 T11/T22=	0.233111545

Time (relative to 2Ne) =	9.99999978E-03

T11=	0.29455892399112360	T12=	0.99381284030198780	T22=	0.99877933641844285

TT=	0.99084316299320774	TS=	0.92835729517571097

FAT=	6.3063328437101163E-002 T11/T22=	0.294918925

Time (relative to 2Ne) =	1.25000002E-02

T11=	0.34944415596201400	T12=	0.99318657855016379	T22=	0.99845400759923375

TT=	0.99101577185402900	TS=	0.93355302243551186

FAT=	5.7983688101162834E-002 T11/T22=	0.349985242

Time (relative to 2Ne) =	1.49999997E-02

T11=	0.39831897135329936	T12=	0.99282958725101933	T22=	0.99812793641015540

TT=	0.99117614391094233	TS=	0.93814703990446990

FAT=	5.3501190814815525E-002 T11/T22=	0.399066061

Time (relative to 2Ne) =	1.75000001E-02

T11=	0.44185815878531814	T12=	0.99270108310330007	T22=	0.99780360458980444

TT=	0.99132569626418887	TS=	0.94220906000935578

FAT=	4.9546416924255210E-002 T11/T22=	0.442830801

Time (relative to 2Ne) =	1.99999996E-02

T11=	0.48065987679121175	T12=	0.99276547669695592	T22=	0.99748308296784505

TT=	0.99146568177731864	TS=	0.94580076235018173

FAT=	4.6057992996063390E-002 T11/T22=	0.481872708

Time (relative to 2Ne) =	2.25000009E-02

T11=	0.51525440251372479	T12=	0.99299174984272298	T22=	0.99716808519511302

TT=	0.99159720800486906	TS=	0.94897671692697427

FAT=	4.2981657001282647E-002 T11/T22=	0.516717732

Time (relative to 2Ne) =	2.50000004E-02

T11=	0.54611187912881620	T12=	0.99335290572473900	T22=	0.99686001496636045

TT=	0.99172125394449318	TS=	0.95178520138260614

FAT=	4.0269432971255292E-002 T11/T22=	0.547832072

Time (relative to 2Ne) =	2.74999999E-02

T11=	0.57364917671887905	T12=	0.99382548345065569	T22=	0.99656000749999019

TT=	0.99183868486329896	TS=	0.95426892442187905

FAT=	3.7878902098477885E-002 T11/T22=	0.575629354

Time (relative to 2Ne) =	2.99999993E-02

T11=	0.59823596816702584	T12=	0.99438912954385117	T22=	0.99626896595052827

TT=	0.99195026541949149	TS=	0.95646566617217799

FAT=	3.5772558851332348E-002 T11/T22=	0.600476384

Time (relative to 2Ne) =	3.24999988E-02

T11=	0.62020011000053987	T12=	0.99502621978102890	T22=	0.99598759335118836

TT=	0.99205667127505326	TS=	0.95840884501612345

FAT=	3.3917242062072472E-002 T11/T22=	0.622698605

Time (relative to 2Ne) =	3.50000001E-02

T11=	0.63983240780325579	T12=	0.99572152553850002	T22=	0.99571642061452781

TT=	0.99215849937273015	TS=	0.96012801933340064

FAT=	3.2283632161171916E-002 T11/T22=	0.642584980

Time (relative to 2Ne) =	3.75000015E-02

T11=	0.65739083669026221	T12=	0.99646191948311369	T22=	0.99545583105783708

TT=	0.99225627703071129	TS=	0.96164933162107968

FAT=	3.0845806792194508E-002 T11/T22=	0.660391748

Time (relative to 2Ne) =	3.99999991E-02

T11=	0.67310427925833771	T12=	0.99723611603915108	T22=	0.99520608186562654

TT=	0.99235046999078813	TS=	0.96299590160489767

FAT=	2.9580847970135937E-002 T11/T22=	0.676346600

Time (relative to 2Ne) =	4.25000004E-02

T11=	0.68717583627197942	T12=	0.99803444258941043	T22=	0.99496732285356138

TT=	0.99244148954019851	TS=	0.96418817419540315

FAT=	2.8468494760215290E-002 T11/T22=	0.690651655

Time (relative to 2Ne) =	4.50000018E-02

T11=	0.69978575901149753	T12=	0.99884863783503786	T22=	0.99473961285572399

TT=	0.99252969881355835	TS=	0.96524422747130134

FAT=	2.7490836168301391E-002 T11/T22=	0.703486383

Time (relative to 2Ne) =	4.74999994E-02

T11=	0.71109404660240794	T12=	0.99967167415135738	T22=	0.99452293401952330

TT=	0.99261541836908229	TS=	0.96618004527781176

FAT=	2.6632039561409582E-002 T11/T22=	0.715010226

Time (relative to 2Ne) =	5.00000007E-02

T11=	0.72124274668096389	T12=	1.0004976011421889	T22=	0.99431720425934633

TT=	0.99269893112247432	TS=	0.96700975850150805

FAT=	2.5878110488059769E-002 T11/T22=	0.725364864

Time (relative to 2Ne) =	7.50000030E-02

T11=	0.78083186370440705	T12=	1.0081449104967588	T22=	0.99281607100865399

TT=	0.99345542004347043	TS=	0.97161765027822922

FAT=	2.1981630302329691E-002 T11/T22=	0.786481917

Time (relative to 2Ne) =	0.100000001

T11=	0.80316560291563410	T12=	1.0138366644900647	T22b=	0.99212304923408579

TT=	0.99414192551697755	TS=	0.97322730460224061

FAT=	2.1037862278930453E-002 T11/T22=	0.809542298

Time (relative to 2Ne) =	0.200000003

T11=	0.82156346127234769	T12=	1.0234661375495657	T22=	0.99289221368984448

TT=	0.99668223246041943	TS=	0.97575933844809482

FAT=	2.0992542388032942E-002 T11/T22=	0.827444732

Time (relative to 2Ne) =	0.300000012

T11=	0.82477839311595191	T12=	1.0267098071118019	T22=	0.99498708188995566

TT=	0.99899508554214800	TS=	0.97796621301255526

FAT=	2.1050026005063338E-002 T11/T22=	0.828933775

Time (relative to 2Ne) =	0.400000006

T11=	0.82665421811487083	T12=	1.0289710259826508	T22=	0.99705342140387965

TT=	1.0010945981951684	TS=	0.98001350107497875

FAT=	2.1058047019927861E-002 T11/T22=	0.829097211

Time (relative to 2Ne) =	0.500000000

T11=	0.82823793111655786	T12=	1.0309417748340661	T22=	0.99894663940391648

TT=	1.0029986766984700	TS=	0.98187576857518066

FAT=	2.1059756721533041E-002 T11/T22=	0.829111278

Time (relative to 2Ne) =	1.00000000

T11=	0.83413601959982364	T12=	1.0383166122559258	T22=	1.0060828945181461

TT=	1.0101654949617633	TS=	0.98888820702631397

FAT=	2.1063170383041840E-002 T11/T22=	0.829092741

Time (relative to 2Ne) =	2.00000000

T11=	0.83996674456235199	T12=	1.0456085049078498	T22=	1.0131407541192006

TT=	1.0172532091655890	TS=	0.99582335316351567

FAT=	2.1066393115290682E-002 T11/T22=	0.829072118

Time (relative to 2Ne) =	3.00000000

T11=	0.84215836915566711	T12=	1.0483493461295206	T22=	1.0157936281488731

TT=	1.0199173047954577	TS=	0.99843010224955253

FAT=	2.1067592877262120E-002 T11/T22=	0.829064429

Time (relative to 2Ne) =	4.00000000

T11=	0.84298214637542535	T12=	1.0493795602310934	T22=	1.0167907775690785

TT=	1.0209186721363048	TS=	0.99940991444971317

FAT=	2.1068042218861494E-002 T11/T22=	0.829061568

Time (relative to 2Ne) =	5.00000000

T11=	0.84329178376034175	T12=	1.0497667921096450	T22=	1.0171655812556069

TT=	1.0212950612343812	TS=	0.99977820150608032

FAT=	2.1068210887355776E-002 T11/T22=	0.829060495

**Scaled recombination rate=	4**

**Equilibrium results for panmictic population**

T11=	0.47058823529411764	T12=	1.5000000000000000	T22=	1.0588235294117647

TT=	1.1323529411764706	TS=	1.0000000000000000

FAT=	0.11688311688311681	T11/T22=	0.444444448

Time (relative to 2Ne) =	2.49999994E-03

T11=	1.1096388746625274E-002 T12=	1.0019573916219786	T22=	0.99972920969789181

TT=	0.99024395423471490	TS=	0.90086592760276518

FAT=	9.0258593601839521E-002 T11/T22=	1.10993944E-02

Time (relative to 2Ne) =	4.99999989E-03

T11=	2.2061826857566136E-002 T12=	1.0039491496460526	T22=	0.99945589562227066

TT=	0.99049074065890441	TS=	0.90171648874580013

FAT=	8.9626533867493752E-002 T11/T22=	2.20738370E-02

Time (relative to 2Ne) =	9.99999978E-03

T11=	4.2957098298150698E-002 T12=	1.0079152229510406	T22=	0.99891829789310815

TT=	0.99097813240758648	TS=	0.90332217793361247

FAT=	8.8453974520117273E-002 T11/T22=	4.30036150E-02

Time (relative to 2Ne) =	1.49999997E-02

T11=	6.2547103243070057E-002 T12=	1.0118572106048804	T22=	0.99839262578598165

TT=	0.99145779582795435	TS=	0.90480807353169057

FAT=	8.7396279156697343E-002 T11/T22=	6.26478046E-02

Time (relative to 2Ne) =	1.99999996E-02

T11=	8.0917250165951804E-002 T12=	1.0157742022575573	T22=	0.99787874062625104

TT=	0.99193030881528321	TS=	0.90618259158022108

FAT=	8.6445304143872082E-002 T11/T22=	8.10892582E-02

Time (relative to 2Ne) =	2.50000004E-02

T11=	9.8147315928966861E-002 T12=	1.0196653775491908	T22=	0.99737650305918069

TT=	0.99239620859608046	TS=	0.90745358434615930

FAT=	8.5593459058139199E-002 T11/T22=	9.84054804E-02

Time (relative to 2Ne) =	2.99999993E-02

T11=	0.11431181763479226	T12=	1.0235299997910292	T22=	0.99688577322805361

TT=	0.99285599445345663	TS=	0.90862837766872750

FAT=	8.4833668986502353E-002 T11/T22=	0.114668921

Time (relative to 2Ne) =	3.99999991E-02

T11=	0.14371795930429515	T12=	1.0311770217392162	T22=	0.99593827581189653

TT=	0.99375904691373806	TS=	0.91071624416113639

FAT=	8.3564323776978977E-002 T11/T22=	0.144304082

Time (relative to 2Ne) =	5.00000007E-02

T11=	0.16963925249512640	T12=	1.0387108335837751	T22=	0.99503512543134032

TT=	0.99464279416941659	TS=	0.91249553813771889

FAT=	8.2589706086691472E-002 T11/T22=	0.170485690

Time (relative to 2Ne) =	5.99999987E-02

T11=	0.19251508886647206	T12=	1.0461280009197025	T22=	0.99417520040601193

TT=	0.99551010338308088	TS=	0.91400918925205787

FAT=	8.1868495210701830E-002 T11/T22=	0.193643019

Time (relative to 2Ne) =	7.00000003E-02

T11=	0.21272886993363155	T12=	1.0534259508902317	T22=	0.99335738399240858

TT=	0.99636344089342910	TS=	0.91529453258653093

FAT=	8.1364796197464395E-002 T11/T22=	0.214151397

Time (relative to 2Ne) =	7.99999982E-02

T11=	0.23061515578359398	T12=	1.0606028515965615	T22=	0.99258056748593937

TT=	0.99720492450882792	TS=	0.91638402631570492

FAT=	8.1047431883603305E-002 T11/T22=	0.232338980

Time (relative to 2Ne) =	9.00000036E-02

T11=	0.24646590070432836	T12=	1.0676575071793162	T22=	0.99184365282943165

TT=	0.99803636909115989	TS=	0.91730587761692128

FAT=	8.0889328259404092E-002 T11/T22=	0.248492688

Time (relative to 2Ne) =	0.100000001

T11=	0.26053589236279495	T12=	1.0745892665629801	T22=	0.99114555479365429

TT=	0.99885932628782448	TS=	0.91808458855056840

FAT=	8.0866980576182357E-002 T11/T22=	0.262863398

Time (relative to 2Ne) =	0.125000000

T11=	0.28931064339766055	T12=	1.0913807367104837	T22=	0.98956314665074274

TT=	1.0008877878289653	TS=	0.91953789632543459

FAT=	8.1277734120412726E-002 T11/T22=	0.292361975

Time (relative to 2Ne) =	0.150000006

T11=	0.31104005589704598	T12=	1.1074107960196478	T22=	0.98820043141187752

TT=	1.0028866932861278	TS=	0.92048439386039438

FAT=	8.2165113942960333E-002 T11/T22=	0.314754009

Time (relative to 2Ne) =	0.200000003

T11=	0.34069054043478508	T12=	1.1372620828375040	T22=	0.98607270202473729

TT=	1.0068329689551359	TS=	0.92153448586574205

FAT=	8.4719596715147616E-002 T11/T22=	0.345502466

Time (relative to 2Ne) =	0.250000000

T11=	0.35940459663122504	T12=	1.1643454646626152	T22=	0.98464752588842397

TT=	1.0107407255752063	TS=	0.92212323296270415

FAT=	8.7675791001763148E-002 T11/T22=	0.365008384

Time (relative to 2Ne) =	0.300000012

T11=	0.37223517319045663	T12=	1.1889030065057464	T22=	0.98382278648362020

TT=	1.0146213499546715	TS=	0.92266402515430390

FAT=	9.0632160267942075E-002 T11/T22=	0.378355920

Time (relative to 2Ne) =	0.400000006

T11=	0.38936411401406135	T12=	1.2314107142395252	T22=	0.98362599538920847

TT=	1.0222846259685141	TS=	0.92419980725169382

FAT=	9.5946682778188630E-002 T11/T22=	0.395845681

Time (relative to 2Ne) =	0.500000000

T11=	0.40121561499171421	T12=	1.2665651575900454	T22=	0.98489003475322467

TT=	1.0297548126662375	TS=	0.92652259277707372

FAT=	0.10024932014823540	T11/T22=	0.407370985

Time (relative to 2Ne) =	0.600000024

T11=	0.41044710066362017	T12=	1.2958318227580001	T22=	0.98716589404185329

TT=	1.0369585732769775	TS=	0.92949401470403004

FAT=	0.10363437975476675	T11/T22=	0.415783316

Time (relative to 2Ne) =	0.699999988

T11=	0.41799191972638633	T12=	1.3203758383637625	T22=	0.99011608719144228

TT=	1.0438416007278095	TS=	0.93290367044493672

FAT=	0.10627851026968294	T11/T22=	0.422164559

Time (relative to 2Ne) =	0.800000012

T11=	0.42428825490456684	T12=	1.3411150229725710	T22=	0.99348884295676865

TT=	1.0503695494790912	TS=	0.93656878415154843

FAT=	0.10834354954785186	T11/T22=	0.427068979

Time (relative to 2Ne) =	1.00000000

T11=	0.43413407272776527	T12=	1.3739150447213688	T22=	1.0008069464758820

TT=	1.0622996754225884	TS=	0.94413965910107034

FAT=	0.11123039859210482	T11/T22=	0.433784038

Time (relative to 2Ne) =	1.50000000

T11=	0.44913323565582680	T12=	1.4248435669841952	T22=	1.0182733227874903

TT=	1.0857645658715804	TS=	0.96135931407432396

FAT=	0.11457847834386836	T11/T22=	0.441073358

Time (relative to 2Ne) =	2.00000000

T11=	0.45721668779238622	T12=	1.4528724214711399	T22=	1.0316116156145867

TT=	1.1016946113905444	TS=	0.97417212283236676

FAT=	0.11575121384792875	T11/T22=	0.443206221

Time (relative to 2Ne) =	3.00000000

T11=	0.46504524975310890	T12=	1.4803878342590027	T22=	1.0470295291892042

TT=	1.1192141813074070	TS=	0.98883110124559470

FAT=	0.11649520014972081	T11/T22=	0.444156766

Time (relative to 2Ne) =	4.00000000

T11=	0.46823519454943191	T12=	1.4916686502297680	T22=	1.0537775155125058

TT=	1.1267424965519823	TS=	0.99522328341619848

FAT=	0.11672517326563459	T11/T22=	0.444339722

Time (relative to 2Ne) =	5.00000000

T11=	0.46958511488444071	T12=	1.4964478413701185	T22=	1.0566694263506828

TT=	1.1299586979395189	TS=	0.99796099520405868

FAT=	0.11681639601178184	T11/T22=	0.444401145

**Scaled recombination rate=	0.4**

Equilibrium results for panmictic population

T11=	0.30501930501930496	T12=	6.0000000000000000	T22=	1.0772200772200773

TT=	1.9555984555984556	TS=	1.0000000000000000

FAT=	0.48864758144126352	T11/T22=	0.283154130

Time (relative to 2Ne) =	4.99999989E-03

T11=	6.5681621938282198E-003 T12=	1.0048487063379463	T22=	0.99945202261081834

TT=	0.99049458707753157	TS=	0.90016363656911935

FAT=	9.1197823478202933E-002 T11/T22=	6.57176320E-03

Time (relative to 2Ne) =	9.99999978E-03

T11=	1.2877860134889951E-002 T12=	1.0097416687766518	T22=	0.99890264236831872

TT=	0.99099341929948448	TS=	0.90030016414497582

FAT=	9.1517515039220010E-002 T11/T22=	1.28920069E-02

Time (relative to 2Ne) =	1.49999997E-02

T11=	1.8877537330213467E-002 T12=	1.0146298634998638	T22=	0.99835739046173688

TT=	0.99149163707728460	TS=	0.90040940514858447

FAT=	9.1863842843084353E-002 T11/T22=	1.89085975E-02

Time (relative to 2Ne) =	1.99999996E-02

T11=	2.4582839580652660E-002 T12=	1.0195132687698507	T22=	0.99781624224476839

TT=	0.99198937299264212	TS=	0.90049290197835685

FAT=	9.2235333870823499E-002 T11/T22=	2.46366393E-02

Time (relative to 2Ne) =	2.99999993E-02

T11=	3.5168992067134253E-002 T12=	1.0292656315407873	T22=	0.99674615898029162

TT=	0.99298389237204931	TS=	0.90058844228897583

FAT=	9.3048286878408826E-002 T11/T22=	3.52838002E-02

Time (relative to 2Ne) =	3.99999991E-02

T11=	4.4746387007337740E-002 T12=	1.0389986122841179	T22=	0.99569219814084375

TT=	0.99397789457529817	TS=	0.90059761702749308

FAT=	9.3946030447391582E-002 T11/T22=	4.49399799E-02

Time (relative to 2Ne) =	5.00000007E-02

T11=	5.3414258304859463E-002 T12=	1.0487120870320421	T22=	0.99465416746437540

TT=	0.99497219389496028	TS=	0.90053017654842382

FAT=	9.4919252945984134E-002 T11/T22=	5.37013374E-02

Time (relative to 2Ne) =	7.50000030E-02

T11=	7.1670426643324681E-002 T12=	1.0729097856881555	T22=	0.99212754291431959

TT=	0.99746377545090015	TS=	0.90008183128722008

FAT=	9.7629554636867732E-002 T11/T22=	7.22391233E-02

Time (relative to 2Ne) =	0.100000001

T11=	8.5951407166925314E-002 T12=	1.0969837432950174	T22=	0.98969638720620945

TT=	0.99997066150180203	TS=	0.89932188920228107

FAT=	0.10065172526998145	T11/T22=	8.68462399E-02

Time (relative to 2Ne) =	0.150000006

T11=	0.10600603519374022	T12=	1.1447579933086063	T22=	0.98510923944092610

TT=	1.0050549830946367	TS=	0.89719891901620752

FAT=	0.10731359566651033	T11/T22=	0.107608408

Time (relative to 2Ne) =	0.200000003

T11=	0.11861868257528732	T12=	1.1920333118979096	T22=	0.98087102487541622

TT=	1.0102577131164638	TS=	0.89464579064540339

FAT=	0.11443804978674044	T11/T22=	0.120931990

Time (relative to 2Ne) =	0.300000012

T11=	0.13231328454328592	T12=	1.2850981444780887	T22=	0.97336060939407909

TT=	1.0210628924606930	TS=	0.88925587690899977

FAT=	0.12908804788121053	T11/T22=	0.135934502

Time (relative to 2Ne) =	0.400000006

T11=	0.13923279693312712	T12=	1.3762161961623480	T22=	0.96701472625515317

TT=	1.0323931715452281	TS=	0.88423653332295060

FAT=	0.14350796024786272	T11/T22=	0.143982083

Time (relative to 2Ne) =	0.500000000

T11=	0.14370566326046796	T12=	1.4654371585132902	T22=	0.96169948723753129

TT=	1.0441923298273974	TS=	0.87990010483982506

FAT=	0.15733904597319681	T11/T22=	0.149428859

Time (relative to 2Ne) =	1.00000000

T11=	0.15929185847124055	T12=	1.8848428657086771	T22=	0.94684427984976427

TT=	1.1078085010905836	TS=	0.86808903771191193

FAT=	0.21639070574262564	T11/T22=	0.168234482

Time (relative to 2Ne) =	2.00000000

T11=	0.18488077151046656	T12=	2.6069355457021461	T22=	0.94867900991965426

TT=	1.2395272039764109	TS=	0.87229918607873547

FAT=	0.29626458920756693	T11/T22=	0.194882318

Time (relative to 2Ne) =	3.00000000

T11=	0.20589528178091823	T12=	3.2003175836660005	T22=	0.96454529302112002

TT=	1.3593978052247966	TS=	0.88868029189709985

FAT=	0.34626914323276892	T11/T22=	0.213463575

Time (relative to 2Ne) =	4.00000000

T11=	0.22320931647323239	T12=	3.6892986617781132	T22=	0.98214778235533862

TT=	1.4618455559926171	TS=	0.90625393576712798

FAT=	0.38006177735255575	T11/T22=	0.227266535

Time (relative to 2Ne) =	5.00000000

T11=	0.23749136337677768	T12=	4.0926785184864967	T22=	0.99809253689438038

TT=	1.5475120018457855	TS=	0.92203241954262005

FAT=	0.40418399440981945	T11/T22=	0.237945229

**Inversion frequency=	0.5**

**Scaled gene conversion rate=	40**

**Initial state of population**

T11=	0	T12=	1	T22=	1

TT=	0.75	TS=	0.5

FAT=	0.3333	T11/T22=	0

**Scaled recombination rate=	40**

**Equilibrium results for panmictic population**

T11=	1	T12=	1.05	T22=	1

TT=	1.025	TS=	1

FAT=	0.02439 T11/T22=	1

Time (relative to 2Ne) =	2.49999994E-03

T11=	4.9611521938714209E-002 T12=	0.97903210528902196	T22=	0.99711026687333160

TT=	0.75119649984752246	TS=	0.52336089440602285

FAT=	0.30329694758661097	T11/T22=	4.97553013E-02

Time (relative to 2Ne) =	4.99999989E-03

T11=	9.6572137562790483E-002 T12=	0.95976708345804351	T22=	0.99333847995891533

TT=	0.75236119610944818	TS=	0.54495530876085296

FAT=	0.27567329152688425	T11/T22=	9.72197726E-02

Time (relative to 2Ne) =	7.49999983E-03

T11=	0.14012817344260553	T12=	0.94244557507855931	T22=	0.98887850388237519

TT=	0.75347445687052483	TS=	0.56450333866249036

FAT=	0.25079963426086893	T11/T22=	0.141704142

Time (relative to 2Ne) =	9.99999978E-03

T11=	0.18055187689970686	T12=	0.92687778991963188	T22=	0.98385714089612197

TT=	0.75454114940877315	TS=	0.58220450889791442

FAT=	0.22839926045901471	T11/T22=	0.183514327

Time (relative to 2Ne) =	1.25000002E-02

T11=	0.21809213350672840	T12=	0.91289245658478158	T22=	0.97838561914224664

TT=	0.75556566645463452	TS=	0.59823887632448747

FAT=	0.20822384752919854	T11/T22=	0.222910196

Time (relative to 2Ne) =	1.49999997E-02

T11=	0.25297657512510996	T12=	0.90033501556753237	T22=	0.97256128371278527

TT=	0.75655197249323991	TS=	0.61276892941894756

FAT=	0.19005045033515799	T11/T22=	0.260113776

Time (relative to 2Ne) =	1.75000001E-02

T11=	0.28541349143436479	T12=	0.88906598861646868	T22=	0.96646911352796483

TT=	0.75750364554881677	TS=	0.62594130248116486

FAT=	0.17367882496767928	T11/T22=	0.295315683

Time (relative to 2Ne) =	1.99999996E-02

T11=	0.31559356363535618	T12=	0.87895950720670302	T22=	0.96018308151945642

TT=	0.75842391489205463	TS=	0.63788832257740635

FAT=	0.15892905003108182	T11/T22=	0.328680605

Time (relative to 2Ne) =	2.25000009E-02

T11=	0.34369143721345397	T12=	0.86990198459322310	T22=	0.95376737387307897

TT=	0.75931569506824470	TS=	0.64872940554326641

FAT=	0.14563940959371213	T11/T22=	0.360351443

Time (relative to 2Ne) =	2.50000004E-02

T11=	0.36986714902473705	T12=	0.86179091743635650	T22=	0.94727748252327282

TT=	0.76018161660518069	TS=	0.65857231577400488

FAT=	0.13366450675950658	T11/T22=	0.390452802

Time (relative to 2Ne) =	2.74999999E-02

T11=	0.39426742250270796	T12=	0.85453380435658211	T22=	0.94076118368329931

TT=	0.76102405372479287	TS=	0.66751430309300364

FAT=	0.12287358089946121	T11/T22=	0.419094056

Time (relative to 2Ne) =	2.99999993E-02

T11=	0.41702684345903562	T12=	0.84804717000950269	T22=	0.93425941392543599

TT=	0.76184514935086933	TS=	0.67564312869223575

FAT=	0.11314900505973169	T11/T22=	0.446371585

Time (relative to 2Ne) =	3.24999988E-02

T11=	0.43826892775586890	T12=	0.84225568438503284	T22=	0.92780705418083576

TT=	0.76264683767669261	TS=	0.68303799096835238

FAT=	0.10438494303714485	T11/T22=	0.472370774

Time (relative to 2Ne) =	3.50000001E-02

T11=	0.45810709104687752	T12=	0.83709136803946027	T22=	0.92143363099692588

TT=	0.76343086453068099	TS=	0.68977036102190170

FAT=	9.6486148164918739E-002 T11/T22=	0.497167766

Time (relative to 2Ne) =	3.99999991E-02

T11=	0.49398002199795343	T12=	0.82840484490462296	T22=	0.90901862335383699

TT=	0.76495208379025914	TS=	0.70149932267589521

FAT=	8.2949981389634275E-002	T11/T22=	0.543421209

Time (relative to 2Ne) =	4.25000004E-02

T11=	0.51019865488866589	T12=	0.82477732016061178	T22=	0.90301463536450544

TT=	0.76569198264359883	TS=	0.70660664512658566

FAT=	7.7165934679134796E-002	T11/T22=	0.564994872

Time (relative to 2Ne) =	4.50000018E-02

T11=	0.52538248689218547	T12=	0.82156521760681867	T22=	0.89716572347090440

TT=	0.76641966139418172	TS=	0.71127410518154499

FAT=	7.1952167970641967E-002	T11/T22=	0.585602522

Time (relative to 2Ne) =	5.00000007E-02

T11=	0.55293839528680711	T12=	0.81622851399767482	T22=	0.88597434555598653

TT=	0.76784244220953579	TS=	0.71945637042139676

FAT=	6.3015625508930917E-002	T11/T22=	0.624102056

Time (relative to 2Ne) =	5.49999997E-02

T11=	0.57717719667685663	T12=	0.81211461765706416	T22=	0.87550410671281054

TT=	0.76922763467594879	TS=	0.72634065169483364

FAT=	5.5753305065778136E-002	T11/T22=	0.659251273

Time (relative to 2Ne) =	5.99999987E-02

T11=	0.59854918977114802	T12=	0.80899541458105406	T22=	0.86578443923795811

TT=	0.77058111454280354	TS=	0.73216681450455301

FAT=	4.9851079027601441E-002 T11/T22=	0.691337407

Time (relative to 2Ne) =	6.49999976E-02

T11=	0.61743822185223451	T12=	0.80668514015320314	T22=	0.85682218674156108

TT=	0.77190767222505041	TS=	0.73713020429689780

FAT=	4.5053921834855459E-002	T11/T22=	0.720614195

Time (relative to 2Ne) =	7.00000003E-02

T11=	0.63417223852172322	T12=	0.80503251796934172	T22=	0.84860758251346513

TT=	0.77321121424346795	TS=	0.74138991051759418

FAT=	4.1154736428659633E-002	T11/T22=	0.747309208

Time (relative to 2Ne) =	7.50000030E-02

T11=	0.64903206120709434	T12=	0.80391435790120536	T22=	0.84111893208048638

TT=	0.77449492727249791	TS=	0.74507549664379036

FAT=	3.7985310933297600E-002	T11/T22=	0.771629333

Time (relative to 2Ne) =	7.99999982E-02

T11=	0.66225870201151038	T12=	0.80323034252619996	T22=	0.83432625988271703

TT=	0.77576141173665680	TS=	0.74829248094711365

FAT=	3.5408993504910113E-002	T11/T22=	0.793764651

Time (relative to 2Ne) =	8.50000009E-02

T11=	0.67405947004277467	T12=	0.80289878133162185	T22=	0.82819412972639128

TT=	0.77701279060810235	TS=	0.75112679988458297

FAT=	3.3314754964664850E-002	T11/T22=	0.813890636

Time (relative to 2Ne) =	9.00000036E-02

T11=	0.68461307799372673	T12=	0.80285315304920568	T22=	0.82268380794252316

TT=	0.77825079800866526	TS=	0.75364844296812494

FAT=	3.1612373676315086E-002	T11/T22=	0.832170367

Time (relative to 2Ne) =	0.100000001

T11=	0.70257566630899737	T12=	0.80341308406616352	T22=	0.81336660624238433

TT=	0.78069211017092721	TS=	0.75797113627569090

FAT=	2.9103629458048119E-002	T11/T22=	0.863787174

Time (relative to 2Ne) =	0.125000000

T11=	0.73426899701534187	T12=	0.80704897664278041	T22=	0.79817046157412397

TT=	0.78663435296875672	TS=	0.76621972929473292

FAT=	2.5951858823580531E-002	T11/T22=	0.919940054

Time (relative to 2Ne) =	0.150000006

T11=	0.75428798276719267	T12=	0.81207490036992369	T22=	0.79114476162665881

TT=	0.79239563628342469	TS=	0.77271637219692568

FAT=	2.4835149495270659E-002	T11/T22=	0.953413367

Time (relative to 2Ne) =	0.174999997

T11=	0.76787114092980913	T12=	0.81750905794664763	T22=	0.78912921626931909

TT=	0.79800461827310576	TS=	0.77850017859956411

FAT=	2.4441512275642618E-002	T11/T22=	0.973061323

Time (relative to 2Ne) =	0.200000003

T11=	0.77781484264791478	T12=	0.82300193388889120	T22b=	0.79007597292366094

TT=	0.80347367083733956	TS=	0.78394540778578792

FAT=	2.4304795241392618E-002	T11/T22=	0.984481096

Time (relative to 2Ne) =	0.300000012

T11=	0.80339612411134487	T12=	0.84403568995348521	T22=	0.80475305199488112

TT=	0.82405513900329908	TS=	0.80407458805311305

FAT=	2.4246618951193843E-002	T11/T22=	0.998313844

Time (relative to 2Ne) =	0.400000006

T11=	0.82220216002195046	T12=	0.86316912770957788	T22=	0.82235232996874208

TT=	0.84272318635246213	TS=	0.82227724499534627

FAT=	2.4261752480801535E-002	T11/T22=	0.999817371

Time (relative to 2Ne) =	0.500000000

T11=	0.83877961311918359	T12=	0.88052593032160587	T22=	0.83879623228752243

TT=	0.85965692651247949	TS=	0.83878792270335301

FAT=	2.4275967732603987E-002	T11/T22=	0.999980211

Time (relative to 2Ne) =	0.600000024

T11=	0.85376380325673085	T12=	0.89627027922494917	T22b=0.85376564248463349

TT=	0.87501750104781573	TS=	0.85376472287068217

FAT=	2.4288403548139104E-002	T11/T22=	0.999997854

Time (relative to 2Ne) =	0.699999988

T11=	0.86735005788964803	T12=	0.91055195902459340	T22=	0.86735026143528948

TT=	0.88895105934353102	TS=	0.86735015966246876

FAT=	2.4299312604468937E-002	T11/T22=	0.999999762

Time (relative to 2Ne) =	0.800000012

T11=	0.87967348142935020	T12=	0.92350685324028003	T22=	0.87967350395555721

TT=	0.90159017296636690	TS=	0.87967349269245365

FAT=	2.4308916546643466E-002	T11/T22=	1.00000000

Time (relative to 2Ne) =	0.899999976

T11=	0.89085197170727604	T12=	0.93525822187785557	T22=	0.89085197420023032

TT=	0.91305509741580437	TS=	0.89085197295375318

FAT=	2.4317398287236003E-002	T11/T22=	1.00000000

Time (relative to 2Ne) =	1.00000000

T11=	0.90099195853630643	T12=	0.94591787394110649	T22=	0.90099195881219929

TT=	0.92345491630767962	TS=	0.90099195867425286

FAT=	2.4324909897325719E-002	T11/T22=	1.00000000

Time (relative to 2Ne) =	1.50000000

T11=	0.93919442991471080	T12=	0.98607819197354518	T22=	0.93919442991471536

TT=	0.96263631094412916	TS=	0.93919442991471302

FAT=	2.4351752331495669E-002	T11/T22=	1.00000000

Time (relative to 2Ne) =	2.00000000

T11=	0.96265639332029473	T12=	1.0107425626657656	T22=	0.96265639332029473

TT=	0.98669947799303004	TS=	0.96265639332029473

FAT=	2.4367180898523966E-002	T11/T22=	1.00000000

Time (relative to 2Ne) =	3.00000000

T11=	0.98591483134930114	T12=	1.0351929774649367	T22=	0.98591483134930114

TT=	1.0105539044071188	TS=	0.98591483134930114

FAT=	2.4381750394871893E-002	T11/T22=	1.00000000

Time (relative to 2Ne) =	4.00000000

T11=	0.99468739113443883	T12=	1.0444151240824200	T22=	0.99468739113443883

TT=	1.0195512576084294	TS=	0.99468739113443883

FAT=	2.4387068613219043E-002	T11/T22=	1.00000000

Time (relative to 2Ne) =	5.00000000

T11=	0.99799620340669204	T12=	1.0478935103974547	T22=	0.99799620340669204

TT=	1.0229448569020734	TS=	0.99799620340669204

FAT=	2.4389050227924147E-002	T11/T22=	1.00000000

**Scaled recombination rate=	4**

**Equilibrium results for panmictic population**

T11=	1	T12=	1.5	T22=	1

TT=	1.25	TS=	1

FAT=	0.2	T11/T22=	1

Time (relative to 2Ne) =	2.49999994E-03

T11=	7.3148393266577738E-003	T12=	1.0000058524484028	T22=	0.99756173239565249

TT=	0.75122206915477907	TS=	0.50243828586115513

FAT=	0.33117209079538557	T11/T22=	7.33271847E-03

Time (relative to 2Ne) =	4.99999989E-03

T11=	1.4705483837823330E-002	T12=	1.0000240212241656	T22=	0.99509827529125083

TT=	0.75246295039435129	TS=	0.50490187956453703

FAT=	0.32900101021594785	T11/T22=	1.47779211E-02

Time (relative to 2Ne) =	7.49999983E-03

T11=	2.2022703007873012E-002	T12=	1.0000543274159619	T22=	0.99265945255117649

TT=	0.75369770259774338	TS=	0.50734107777952475

FAT=	0.32686397207940254	T11/T22=	2.21855566E-02

Time (relative to 2Ne) =	9.99999978E-03

T11=	2.9267287450317481E-002	T12=	1.0000965895481704	T22=	0.99024507898148262

TT=	0.75492638638203524	TS=	0.50975618321590010

FAT=	0.32476041053632643	T11/T22=	2.95556001E-02

Time (relative to 2Ne) =	1.49999997E-02

T11=	4.3541670882651484E-002	T12=	1.0002162677270774	T22=	0.98548894330910641

TT=	0.75736578741147820	TS=	0.51451530709587900

FAT=	0.32065150598578340	T11/T22=	4.41828109E-02

Time (relative to 2Ne) =	1.99999996E-02

T11=	5.7534785527347032E-002	T12=	1.0003816515025010	T22=	0.98082840568263108

TT=	0.75978162355374512	TS=	0.51918159560498900

FAT=	0.31666997527972807	T11/T22=	5.86593784E-02

Time (relative to 2Ne) =	2.99999993E-02

T11=	8.4701139372310053E-002	T12=	1.0008441028681534	T22=	0.97178835047338397

TT=	0.76454442389550015	TS=	0.52824474492284701

FAT=	0.30907252945311015	T11/T22=	8.71600658E-02

Time (relative to 2Ne) =	3.99999991E-02

T11=	0.11081290367374597	T12=	1.0014734325597010	T22=	0.96311351847354609

TT=	0.76921832181667360	TS=	0.53696321107364597

FAT=	0.30193652979365793	T11/T22=	0.115056947

Time (relative to 2Ne) =	5.00000007E-02

T11=	0.13591460683518541	T12=	1.0022596768833969	T22=	0.95479276097571364

TT=	0.77380668039442324	TS=	0.54535368390544958

FAT=	0.29523264954565531	T11/T22=	0.142349854

Time (relative to 2Ne) =	5.99999987E-02

T11=	0.16004884089004545	T12=	1.0031933915246636	T22=	0.94681517624709610

TT=	0.77831270004661712	TS=	0.55343200856857078

FAT=	0.28893360144910529	T11/T22=	0.169039160

Time (relative to 2Ne) =	7.00000003E-02

T11=	0.18325634735549423	T12=	1.0042656249654214	T22=	0.93917010976480930

TT=	0.78273942676278652	TS=	0.56121322856015177

FAT=	0.28301397710194742	T11/T22=	0.195125833

Time (relative to 2Ne) =	7.99999982E-02

T11=	0.20557609921181830	T12=	1.0054678932583054	T22=	0.93184715393417294

TT=	0.78708975991565044	TS=	0.56871162657299568

FAT=	0.27745010094662848	T11/T22=	0.220611393

Time (relative to 2Ne) =	9.00000036E-02

T11=	0.22704537918518408	T12=	1.0067921560885336	T22=	0.92483614733620034

TT=	0.79136645967461283	TS=	0.57594076326069221

FAT=	0.27221989734376340	T11/T22=	0.245497957

Time (relative to 2Ne) =	0.100000001

T11=	0.24769985450331711	T12=	1.0082307940577298	T22=	0.91812717354734341

TT=	0.79557215404153003	TS=	0.58291351402533031

FAT=	0.26730276937910347	T11/T22=	0.269788176

Time (relative to 2Ne) =	0.109999999

T11=	0.26757364828577734	T12=	1.0097765871273459	T22=	0.91171055957160563

TT=	0.79970934552801876	TS=	0.58964210392869143

FAT=	0.26267948820909126	T11/T22=	0.293485314

Time (relative to 2Ne) =	0.119999997

T11=	0.28669940772294167	T12=	1.0114226941625246	T22=	0.90557687392237540

TT=	0.80378041749259155	TS=	0.59613814082265848

FAT=	0.25833209188857464	T11/T22=	0.316593111

Time (relative to 2Ne) =	0.129999995

T11=	0.30510836919063833	T12=	1.0131626335202448	T22=	0.89971692438873518

TT=	0.80778764015496574	TS=	0.60241264678968676

FAT=	0.25424379274468711	T11/T22=	0.339115947

Time (relative to 2Ne) =	0.140000001

T11=	0.32283042044056143	T12=	1.0149902646284741	T22=	0.89412175551856854

TT=	0.81173317630401964	TS=	0.60847608797956498

FAT=	0.25039889246602443	T11/T22=	0.361058682

Time (relative to 2Ne) =	0.150000006

T11=	0.33989416000005912	T12=	1.0168997705057534	T22=	0.88878264584849831

TT=	0.81561908671501604	TS=	0.61433840292427866

FAT=	0.24678270416820991	T11/T22=	0.382426649

Time (relative to 2Ne) =	0.159999996

T11=	0.35632695390868718	T12=	1.0188856411732352	T22=	0.88369110490855185

TT=	0.81944733529092739	TS=	0.62000902940861957

FAT=	0.24338148077753097	T11/T22=	0.403225690

Time (relative to 2Ne) =	0.170000002

T11=	0.37215498991302232	T12=	1.0209426579136378	T22=	0.87883887002742911

TT=	0.82321979394193179	TS=	0.62549692997022577

FAT=	0.24018234914508507	T11/T22=	0.423462152

Time (relative to 2Ne) =	0.180000007

T11=	0.38740332923558485	T12=	1.0230658783338973	T22=	0.87421790296237589

TT=	0.82693824721643883	TS=	0.63081061609898037

FAT=	0.23717324936613426	T11/T22=	0.443142742

Time (relative to 2Ne) =	0.189999998

T11=	0.40209595602833803	T12=	1.0252506221905076	T22=	0.86982038637588854

TT=	0.83060439669631037	TS=	0.63595817120211329

FAT=	0.23434287883424798	T11/T22=	0.462274700

Time (relative to 2Ne) =	0.200000003

T11=	0.41625582461615329	T12=	1.0274924579386415	T22=	0.86563872017982679

TT=	0.83421986516831570	TS=	0.64094727239799010

FAT=	0.23168064060825277	T11/T22=	0.480865538

Time (relative to 2Ne) =	0.224999994

T11=	0.44946654767128347	T12=	1.0333198386147062	T22=	0.85608094040985117

TT=	0.84304679132763682	TS=	0.65277374404056732

FAT=	0.22569689991634512	T11/T22=	0.525028110

Time (relative to 2Ne) =	0.250000000

T11=	0.47980062892884623	T12=	1.0394187651899907	T22=	0.84771686698408089

TT=	0.85158875657322719	TS=	0.66375874795646350

FAT=	0.22056421854673980	T11/T22=	0.565991580

Time (relative to 2Ne) =	0.275000006

T11=	0.50754348128857996	T12=	1.0457372514787169	T22=	0.84044453113347950

TT=	0.85986562884487328	TS=	0.67399400621102967

FAT=	0.21616356835141781	T11/T22=	0.603898823

Time (relative to 2Ne) =	0.300000012

T11=	0.53295104668747173	T12=	1.0522300436580150	T22=	0.83416936087125204

TT=	0.86789512371868849	TS=	0.68356020377936189

FAT=	0.21239308172340332	T11/T22=	0.638900280

Time (relative to 2Ne) =	0.349999994

T11=	0.57765502471512131	T12=	1.0655862976455972	T22=	0.82426678960845656

TT=	0.88327360240369313	TS=	0.70096090716178894

FAT=	0.20640568759868771	T11/T22=	0.700810730

Time (relative to 2Ne) =	0.400000006

T11=	0.61548062625603861	T12=	1.0792309559937487	T22=	0.81738522408437497

TT=	0.89783194058197768	TS=	0.71643292517020685

FAT=	0.20204116963603169	T11/T22=	0.752987206

Time (relative to 2Ne) =	0.449999988

T11=	0.64769151381139123	T12=	1.0929714628684057	T22=	0.81299371114108576

TT=	0.91165703767232209	TS=	0.73034261247623844

FAT=	0.19888446828537920	T11/T22=	0.796674669

Time (relative to 2Ne) =	0.500000000

T11=	0.67530720707725755	T12=	1.1066643468978119	T22=	0.81064249283066858

TT=	0.92481959842588757	TS=	0.74297484995396301

FAT=	0.19662726523252527	T11/T22=	0.833051801

Time (relative to 2Ne) =	0.600000024

T11=	0.71988962743390617	T12=	1.1335125288946917	T22=	0.81060395373929151

TT=	0.94937965974064520	TS=	0.76524679058659884

FAT=	0.19395072062566454	T11/T22=	0.888090432

Time (relative to 2Ne) =	0.699999988

T11=	0.75410509409336812	T12=	1.1592345688057293	T22=	0.81491029320481090

TT=	0.97187113122740942	TS=	0.78450769364908957

FAT=	0.19278629805753378	T11/T22=	0.925384164

Time (relative to 2Ne) =	0.800000012

T11=	0.78116005916124909	T12=	1.1835673823631789	T22=	0.82191737269224951

TT=	0.99255304914496412	TS=	0.80153871592674930

FAT=	0.19244748014503033	T11/T22=	0.950411916

Time (relative to 2Ne) =	0.899999976

T11=	0.80317571862184756	T12=	1.2064082494550084	T22=	0.83049507010065349

TT=	1.0116218219081294	TS=	0.81683539436125052

FAT=	0.19254866129664061	T11/T22=	0.967104733

Time (relative to 2Ne) =	1.00000000

T11=	0.82156567935667058	T12=	1.2277455334031240	T22=	0.83987765579875973

TT=	1.0292336004904197	TS=	0.83072166757771515

FAT=	0.19287354476001906	T11/T22=	0.978196859

Time (relative to 2Ne) =	1.50000000

T11=	0.88367449202415815	T12=	1.3139057821840123	T22=	0.88615225293735145

TT=	1.0994095773323835	TS=	0.88491337248075475

FAT=	0.19510127005813804	T11/T22=	0.997203887

Time (relative to 2Ne) =	2.00000000

T11=	0.92131689269515360	T12=	1.3729680316253119	T22=	0.92165215411067403

TT=	1.1472262775141129	TS=	0.92148452340291387

FAT=	0.19677177775281740	T11/T22=	0.999636233

Time (relative to 2Ne) =	3.00000000

T11=	0.96342430829046877	T12=	1.4408243005991705	T22=	0.96343044636175212

TT=	1.2021258389626404	TS=	0.96342737732611039

FAT=	0.19856362279219608	T11/T22=	0.999993622

Time (relative to 2Ne) =	4.00000000

T11=	0.98296353437066308	T12=	1.4724345110376140	T22=	0.98296364674839976

TT=	1.2276990507985728	TS=	0.98296359055953142

FAT=	0.19934483135736725	T11/T22=	0.999999881

Time (relative to 2Ne) =	5.00000000

T11=	0.99206402428306961	T12=	1.4871593233670048	T22=	0.99206402634051649

TT=	1.2396116743393990	TS=	0.99206402531179305

FAT=	0.19969773934206170	T11/T22=	1.00000000

**Scaked recombination rate=	0.4**

Equilibrium results for panmictic population

Times are scaled by 2Ne

T11=	1	T12=	6	T22=	1

TT=	3.5	TS=	1

FAT=	0.71428571428571430	T11/T22=	1.00000000

Time (relative to 2Ne) =	4.99999989E-03

T11=	5.9102713051500944E-003 T12=	1.0044530600706141	T22=	0.99507876043654164

TT=	0.75247378797072995	TS=	0.50049451587084581

FAT=	0.33486784008705661	T11/T22=	5.93950087E-03

Time (relative to 2Ne) =	9.99999978E-03

T11=	1.1819703700804651E-002 T12=	1.0089471234833187	T22=	0.99016637362076798

TT=	0.75497008107205255	TS=	0.50099303866078637

FAT=	0.33640676469009267	T11/T22=	1.19370883E-02

Time (relative to 2Ne) =	1.99999996E-02

T11=	2.3458672844973494E-002 T12=	1.0179232711620969	T22=	0.98051559154815349

TT=	0.75995520167933017	TS=	0.50198713219656343

FAT=	0.33945167940520082	T11/T22=	2.39248332E-02

Time (relative to 2Ne) =	2.99999993E-02

T11=	3.4862108448943774E-002 T12=	1.0268834667285063	T22=	0.97109256110899889

TT=	0.76493040075373875	TS=	0.50297733477897133

FAT=	0.34245346467684767	T11/T22=	3.58998813E-02

Time (relative to 2Ne) =	3.99999991E-02

T11=	4.6035104412177841E-002 T12=	1.0358277343339104	T22=	0.96189229467809623

TT=	0.76989571693952374	TS=	0.50396369954513709

FAT=	0.34541303652332944	T11/T22=	4.78588976E-02

Time (relative to 2Ne) =	5.00000007E-02

T11=	5.6982643833890956E-002 T12=	1.0447560981864727	T22=	0.95290991321352625

TT=	0.77485118835509059	TS=	0.50494627852370866

FAT=	0.34833128462299368	T11/T22=	5.97985648E-02

Time (relative to 2Ne) =	5.99999987E-02

T11=	6.7709601424279953E-002 T12=	1.0536685825488510	T22=	0.94414064389389241

TT=	0.77979685260396858	TS=	0.50592512265908618

FAT=	0.35120907327382123	T11/T22=	7.17155859E-02

Time (relative to 2Ne) =	7.00000003E-02

T11=	7.8220745863273433E-002 T12=	1.0625652117359810	T22=	0.93557981780696742

TT=	0.78473274678555072	TS=	0.50690028183512048

FAT=	0.35404724231083406	T11/T22=	8.36067051E-02

Time (relative to 2Ne) =	7.99999982E-02

T11=	8.8520742107957018E-002 T12=	1.0714460101128154	T22=	0.92722286768863249

TT=	0.78965890750555512	TS=	0.50787180489829475

FAT=	0.35684660798343248	T11/T22=	9.54686776E-02

Time (relative to 2Ne) =	9.00000036E-02

T11=	9.8614153649770553E-002 T12=	1.0803110020922304	T22b=	0.91906532571101485

TT=	0.79457537088631158	TS=	0.50883973968039276

FAT=	0.35960796379479287	T11/T22=	0.107298307

Time (relative to 2Ne) =	0.100000001

T11=	0.10850544472259804	T12=	1.0891602121329038	T22b=	0.91110282131875364

TT=	0.79948217257678977	TS=	0.50980413302067584

FAT=	0.36233208130515326	T11/T22=	0.119092427

Time (relative to 2Ne) =	0.125000000

T11=	0.13237782144380705	T12=	1.1112143521519604	T22=	0.89202202104242545

TT=	0.81170713669753836	TS=	0.51219992124311631

FAT=	0.36898433180343670	T11/T22=	0.148401961

Time (relative to 2Ne) =	0.150000006

T11=	0.15507966145998475	T12=	1.1331703922599221	T22=	0.87406943991048935

TT=	0.82387247147257958	TS=	0.51457455068523705

FAT=	0.37541965716430259	T11/T22=	0.177422583

Time (relative to 2Ne) =	0.159999996

T11=	0.16384668653258172	T12=	1.1419254266122492	T22=	0.86719056564980301

TT=	0.82872202635172076	TS=	0.51551862609119237

FAT=	0.37793541175602907	T11/T22=	0.188939661

Time (relative to 2Ne) =	0.170000002

T11=	0.17244023720798007	T12=	1.1506648513522988	T22=	0.86047868659702731

TT=	0.83356215662740130	TS=	0.51645946190250369

FAT=	0.38041877525714152	T11/T22=	0.200400352

Time (relative to 2Ne) =	0.180000007

T11=	0.18086405759800284	T12=	1.1593886912350566	T22=	0.85393013792254258

TT=	0.83839289449766463	TS=	0.51739709776027265

FAT=	0.38287036882596825	T11/T22=	0.211801931

Time (relative to 2Ne) =	0.189999998

T11=	0.18912181038805276	T12=	1.1680969710444433	T22=	0.84754133459589687

TT=	0.84321427176820907	TS=	0.51833157249197481

FAT=	0.38529079755132656	T11/T22=	0.223141700

Time (relative to 2Ne) =	0.200000003

T11=	0.19721707860908033	T12=	1.1767897155916476	T22b=	0.84130876964935752

TT=	0.84802631986043320	TS=	0.51926292412921893

FAT=	0.38768065097946680	T11/T22=	0.234417006

Time (relative to 2Ne) =	0.250000000

T11=	0.23537619672069165	T12=	1.2200212794929985	T22=	0.81237203018187798

TT=	0.87194769647214176	TS=	0.52387411345128476

FAT=	0.39919089691864074	T11/T22=	0.289739400

Time (relative to 2Ne) =	0.300000012

T11=	0.26996778316429049	T12=	1.2628681964507713	T22=	0.78685722058873742

TT=	0.89564034916364266	TS=	0.52841250187651401

FAT=	0.41001708736107012	T11/T22=	0.343096286

Time (relative to 2Ne) =	0.349999994

T11=	0.30135984467516452	T12=	1.3053335894307123	T22=	0.76440424643154536

TT=	0.91910781749203363	TS=	0.53288204555335494

FAT=	0.42021813392097096	T11/T22=	0.394241452

Time (relative to 2Ne) =	0.400000006

T11=	0.32988208186347712	T12=	1.3474205877496646	T22=	0.74469055437993481

TT=	0.94235345293568529	TS=	0.53728631812170602

FAT=	0.42984628915200118	T11/T22=	0.442978740

Time (relative to 2Ne) =	0.449999988

T11=	0.35582987965460733	T12=	1.3891323234086217	T22=	0.72742722154832218

TT=	0.96538043700504317	TS=	0.54162855060146475

FAT=	0.43894807700703875	T11/T22=	0.489162177

Time (relative to 2Ne) =	0.500000000

T11=	0.37946788200746995	T12=	1.4304719278378544	T22=	0.71235545217877794

TT=	0.98819179746548913	TS=	0.54591166709312389

FAT=	0.44756506935872553	T11/T22=	0.532694578

Time (relative to 2Ne) =	0.550000012

T11=	0.40103319422084960	T12=	1.4714425290107025	T22=	0.69924343923991084

TT=	1.0107904228705413	TS=	0.55013831673038016

FAT=	0.45573453776100914	T11/T22=	0.573524415

Time (relative to 2Ne) =	0.600000024

T11=	0.42073825162563405	T12=	1.5120472488870469	T22=	0.68788355293130965

TT=	1.0331790755827595	TS=	0.55431090227847180

FAT=	0.46349000344803193	T11/T22=	0.611641705

Time (relative to 2Ne) =	0.649999976

T11=	0.43877338941895938	T12=	1.5522892011514289	T22=	0.67808982204231993

TT=	1.0553604034410342	TS=	0.55843160573063966

FAT=	0.47086170382188242	T11/T22=	0.647072673

Time (relative to 2Ne) =	0.699999988

T11=	0.45530914477529483	T12=	1.5921714892148628	T22=	0.66969567766077964

TT=	1.0773369502164500	TS=	0.56250241121803723

FAT=	0.47787699001224859	T11/T22=	0.679874718

Time (relative to 2Ne) =	0.750000000

T11=	0.47049831912590445	T12=	1.6316972044522702	T22=	0.66255193190473705

TT=	1.0991111649837955	TS=	0.56652512551532075

FAT=	0.48456066723362490	T11/T22=	0.710130453

Time (relative to 2Ne) =	0.800000012

T11=	0.48447782559233676	T12=	1.6708694246505305	T22=	0.65652496719663045

TT=	1.1206854105225070	TS=	0.57050139639448361

FAT=	0.49093528742513592	T11/T22=	0.737942755

Time (relative to 2Ne) =	0.850000024

T11=	0.49737034395665736	T12=	1.7096912126449322	T22=	0.65149511414931660

TT=	1.1420619708489597	TS=	0.57443272905298692

FAT=	0.49702140188944532	T11/T22=	0.763429105

Time (relative to 2Ne) =	0.899999976

T11=	0.50928580321918138	T12=	1.7481656151238099	T22=	0.64735519841770817

TT=	1.1632430579711273	TS=	0.57832050081844477

FAT=	0.50283778024248549	T11/T22=	0.786717713

Time (relative to 2Ne) =	1.00000000

T11=	0.53056933681491292	T12=	1.8240843634181525	T22=	0.64137128163496604

TT=	1.2050273363215460	TS=	0.58597030922493953

FAT=	0.51372861713356111	T11/T22=	0.827242136

Time (relative to 2Ne) =	1.25000000

T11=	0.57244691572414341	T12=	2.0080117438276197	T22=	0.63637214367621664

TT=	1.3062106367638997	TS=	0.60440952970018003

FAT=	0.53728019609640620	T11/T22=	0.899547398

Time (relative to 2Ne) =	1.50000000

T11=	0.60351046952524623	T12=	2.1838456940744582	T22=	0.64039100240182201

TT=	1.4028982150189961	TS=	0.62195073596353412

FAT=	0.55666724121171363	T11/T22=	0.942409337

Time (relative to 2Ne) =	1.75000000

T11=	0.62803123142323258	T12=	2.3519391088851380	T22=	0.64930880443838013

TT=	1.4953045634079722	TS=	0.63867001793080636

FAT=	0.57288298747968547	T11/T22=	0.967230439

Time (relative to 2Ne) =	2.00000000

T11=	0.64848669360297251	T12=	2.5126308788596776	T22=	0.66076241410110159

TT=	1.5836277163558574	TS=	0.65462455385203699

FAT=	0.58662976967944402	T11/T22=	0.981421888

Time (relative to 2Ne) =	2.50000000

T11=	0.68237216009670154	T12=	2.8130950711955633	T22=	0.68645814535761052

TT=	1.7487551119613596	TS=	0.68441515272715603

FAT=	0.60862721827327027	T11/T22=	0.994047701

Time (relative to 2Ne) =	3.00000000

T11=	0.71093657960949841	T12=	3.0876738538576416	T22=	0.71229660369565195

TT=	1.8996452227551084	TS=	0.71161659165257518

FAT=	0.62539500369416468	T11/T22=	0.998090625

Time (relative to 2Ne) =	3.50000000

T11=	0.73624064993205518	T12=	3.3385959928835738	T22=	0.73669333524460756

TT=	2.0375314927359525	TS=	0.73646699258833137

FAT=	0.63854939410069211	T11/T22=	0.999385536

Time (relative to 2Ne) =	4.00000000

T11=	0.75909853751511691	T12=	3.5678992251670483	T22=	0.75924921425260028

TT=	2.1635365505254534	TS=	0.75917387588385865

FAT=	0.64910513034804995	T11/T22=	0.999801517

Time (relative to 2Ne) =	4.50000000

T11=	0.77989848191275324	T12=	3.7774460477787204	T22=	0.77994863480835053

TT=	2.2786848030696363	TS=	0.77992355836055194

FAT=	0.65773082906863201	T11/T22=	0.999935687

Time (relative to 2Ne) =	5.00000000

T11=	0.79887686185175388	T12=	3.9689385957635070	T22=	0.79889355529082762

TT=	2.3839119021673989	TS=	0.79888520857129075

FAT=	0.66488476027786003	T11/T22=	0.999979079

**Table S4**

**Coalescence times for inversion and standard arrangements in a subdivided population**

**Time course of approach to equilibrium**

**Times scaled by 2x population size**

**Inversion frequency= 0.1**

**Local N=	500**

**Total N=	100000**

**Number of populations =	200**

**Neutral Fst= 0.05**

**Scaled migration rate=	19	Migration rate=	0.0095**

Initial state of subdivided population

Alleles sampled from different populations

T11b=	0	T12b=	1.0523684210526316	T22b=	1.0523684210526316

TTb=	1.0418447368421053	TSb=	0.94713157894736844

FATb=	0.0909	T11/T22= 0

Alleles sampled from the same population

T11w=	0	T12w=	1	T22w=	1

TTw=	0.99	TSw=	0.9

FATw=	0.0909 T11/T22=	0

**Scaled gene recombination rate=	40**

Equilibrium results for subdivided population

Alleles sampled from different populations

T11b=	0.98689902793589634	T12b=	1.1387293107977932	T22b=	1.1000437866713373

TTb=	1.1058757334267451	TSb=	1.0887293107977931

FATb=	1.5504836674389066E-002	T11/T22=	0.897145212

Alleles sampled from the same population

T11w=	0.64707177692351836	T12w=	1.1389924686925299	T22w=	1.0395662293966779

TTw=	1.0535380079451997	TSw=	1.0003167841493619

FATw=	5.0516662326819617E-002	T11/T22=	0.622444034

Time (units of 2NT generations) =	2.49999994E-03

Alleles sampled from different populations

T11b=	9.0470778058974188E-002	T12b=	1.0496957230670723	T22b=	1.0521244381570143

TTb=	1.0420707328398444	TSb=	0.95595907214721032

FATb=	8.2635139802807012E-002	T11/T22=	8.59886706E-02

Alleles sampled from the same population

T11w=	5.8903433078786316E-002	T12w=	1.0501319171837042	T22w=	0.99427794860199548

TTw=	0.99497791779147104	TSw=	0.90074049704967463

FATw=	9.4713077603745255E-002	T11/T22=	5.92424199E-02

Time (units of 2NT generations) =	4.99999989E-03

Alleles sampled from different populations

T11b=	0.17336087601959593	T12b=	1.0477666110871331	T22b=	1.0518308263196747

TTb=	1.0423145680748165	TSb=	0.96398383128966691

FATb=	7.5150764638959755E-002	T11/T22=	0.164818212

Alleles sampled from the same population

T11w=	0.11326810589247005	T12w=	1.0481893820694277	T22w=	0.99400029174213245

TTw=	0.99494700614254894	TSw=	0.90592707315716625

FATw=	8.9472034626765340E-002	T11/T22=	0.113951780

Time (units of 2NT generations) =	7.49999983E-03

Alleles sampled from different populations

T11b=	0.24772926823943830	T12b=	1.0463013012228561	T22b=	1.0515240394185894

TTb=	1.0425459988315660	TSb=	0.97114456230067436

FATb=	6.8487564683874624E-002	T11/T22=	0.235590681

Alleles sampled from the same population

T11w=	0.16204423441218752	T12w=	1.0467106660487946	T22w=	0.99371028322323540

TTw=	0.99493369164372569	TSw=	0.91054367834213068

FATw=	8.4819736240085097E-002	T11/T22=	0.163069904

Time (units of 2NT generations) =	9.99999978E-03

Alleles sampled from different populations

T11b=	0.31447412533363805	T12b=	1.0452373701969842	T22b=	1.0512085426045419

TTb=	1.0427663873984725	TSb=	0.97753510087745155

FATb=	6.2555992702988950E-002	T11/T22=	0.299154848

Alleles sampled from the same population

T11w=	0.20582083564037623	T12w=	1.0456347143175924	T22w=	0.99341206739270616

TTw=	0.99493623152166244	TSw=	0.91465294421747312

FATw=	8.0691892365205686E-002	T11/T22=	0.207185760

Time (units of 2NT generations) =	1.25000002E-02

Alleles sampled from different populations

T11b=	0.37439782880788863	T12b=	1.0445197697530839	T22b=	1.0508881608740943

TTb=	1.0429769471516503	TSb=	0.98323912766747379

FATb=	5.7276260656881628E-002	T11/T22=	0.356268018

Alleles sampled from the same population

T11w=	0.24512408549924869	T12w=	1.0449063320381333	T22w=	0.99310925568562980

TTw=	0.99495287772721674	TSw=	0.91831073866699175

FATw=	7.7030923550168162E-002	T11/T22=	0.246824890

Time (units of 2NT generations) =	1.49999997E-02

Alleles sampled from different populations

T11b=	0.42821724585284993	T12b=	1.0440999917447931	T22b=	1.0505661565586388

TTb=	1.0431787577850888	TSb=	0.98833126548805994

FATb=	5.2577271045551988E-002	T11/T22=	0.407606155

Alleles sampled from the same population

T11w=	0.28042408381189321	T12w=	1.0444768801148983	T22w=	0.99280492819580513

TTw=	0.99498207109740289	TSw=	0.92156684375741393

FATw=	7.3785477620733952E-002	T11/T22=	0.282456368

Time (units of 2NT generations) =	1.75000001E-02

Alleles sampled from different populations

T11b=	0.47657289899592614	T12b=	1.0439353317837168	T22b=	1.0502452977350358

TTb=	1.0433727798764074	TSb=	0.99287805786112493

FATb=	4.8395667386745389E-002	T11/T22=	0.453772932

Alleles sampled from the same population

T11w=	0.31214086723677315	T12w=	1.0443035371961642	T22w=	0.99250169828090617

TTw=	0.99502242097521143	TSw=	0.92446561517649284

FATw=	7.0909764756422855E-002	T11/T22=	0.314499080

Time (units of 2NT generations) =	1.99999996E-02

Alleles sampled from different populations

T11b=	0.52003714958527680	T12b=	1.0439882338945441	T22b=	1.0499279188608608

TTb=	1.0435598678741680	TSb=	0.99693884193330251

FATb=	4.4674989309273605E-002	T11/T22=	0.495307475

Alleles sampled from the same population

T11w=	0.34064977553292020	T12w=	1.0443486428278757	T22w=	0.99220176982510910

TTw=	0.99507268702268525	TSw=	0.92704657039589033

FATw=	6.8362962338292088E-002	T11/T22=	0.343327105

Time (units of 2NT generations) =	2.25000009E-02

Alleles sampled from different populations

T11b=	0.55912150131590010	T12b=	1.0442257073591183	T22b=	1.0496159744932438

TTb=	1.0437407816773279	TSb=	1.0005665271755095

FATb=	4.1364920543236594E-002	T11/T22=	0.532691479

Alleles sampled from the same population

T11w=	0.36628624080226435	T12w=	1.0445791129579991	T22w=	0.99190698796986132

TTw=	0.99513176299605022	TSw=	0.92934491325310153

FATw=	6.6108682477266867E-002	T11/T22=	0.369274795

Time (units of 2NT generations) =	2.50000004E-02

Alleles sampled from different populations

T11b=	0.59428311856136840	T12b=	1.0446188078865632	T22b=	1.0493110868565909

TTb=	1.0439161969590338	TSb=	1.0038082900270688

FATb=	3.8420619441293047E-002	T11/T22=	0.566355526

Alleles sampled from the same population

T11w=	0.38935006184904197	T12w=	1.0449659199110757	T22w=	0.99161888403553999

TTw=	0.99519866227127163	TSw=	0.93139200181689019

FATw=	6.4114495802034099E-002	T11/T22=	0.392640829

Time (units of 2NT generations) =	2.74999999E-02

Alleles sampled from different populations

T11b=	0.62593064406202548	T12b=	1.0451421760984978	T22b=	1.0490145879409567

TTb=	1.0440867143705248	TSb=	1.0067061935530637

FATb=	3.5802122853366369E-002	T11/T22=	0.596684396

Alleles sampled from the same population

T11w=	0.41010921909944570	T12w=	1.0454836298064578	T22w=	0.99133871527786899

TTw=	0.99527250493123087	TSw=	0.93321576566002673

FATw=	6.2351505706964128E-002	T11/T22=	0.413692325

Time (units of 2NT generations) =	2.99999993E-02

Alleles sampled from different populations

T11b=	0.65442939140902323	T12b=	1.0457736270782663	T22b=	1.0487275567383629

TTb=	1.0442528677462521	TSb=	1.0092977402054291

FATb=	3.3473815222805792E-002	T11/T22=	0.624022305

Alleles sampled from the same population

T11w=	0.42880327954991498	T12w=	1.0461099911553280	T22w=	0.99106750005262712

TTw=	0.99535250624608629	TSw=	0.93484107800235594

FATw=	6.0793967829493534E-002	T11/T22=	0.432668090

Time (units of 2NT generations) =	3.24999988E-02

Alleles sampled from different populations

T11b=	0.68010597963426012	T12b=	1.0464937854108252	T22b=	1.0484508521579663

TTb=	1.0444151314182439	TSb=	1.0116163649055956

FATb=	3.1403955693470209E-002	T11/T22=	0.648677051

Alleles sampled from the same population

T11w=	0.44564643588182268	T12w=	1.0468255690507660	T22w=	0.99080604889948309

TTw=	0.99543796639653748	TSw=	0.93629008759771704

FATw=	5.9418950045610930E-002	T11/T22=	0.449781716

Time (units of 2NT generations) =	3.50000001E-02

Alleles sampled from different populations

T11b=	0.70325246996328195	T12b=	1.0472857607444015	T22b=	1.0481851421017929

TTb=	1.0445739267360774	TSb=	1.0136918748879418

FATb=	2.9564256830180602E-002	T11/T22=	0.670923889

Alleles sampled from the same population

T11w=	0.46083021912421834	T12w=	1.0476134199709528	T22w=	0.99055499199989561

TTw=	0.99552826130592920	TSw=	0.93758251471232790

FATw=	5.8206028744566596E-002	T11/T22=	0.465224266

Time (units of 2NT generations) =	3.75000015E-02

Alleles sampled from different populations

T11b=	0.72413005831715671	T12b=	1.0481348594441098	T22b=	1.0479309291300074

TTb=	1.0447296278784175	TSb=	1.0155508420487223

FATb=	2.7929509273083464E-002	T11/T22=	0.691009343

Alleles sampled from the same population

T11w=	0.47452592000302135	T12w=	1.0484588027558692	T22w=	0.99031480341419575

TTw=	0.99562283446158528	TSw=	0.93873591507307841

FATw=	5.7137017572794324E-002	T11/T22=	0.479166746

Time (units of 2NT generations)=	3.99999991E-02

Alleles sampled from different populations

T11b=	0.74297237137581906	T12b=	1.0490283283885478	T22b=	1.0476885730976506

TTb=	1.0448825670327939	TSb=	1.0172169529254675

FATb=	2.6477247281376126E-002	T11/T22=	0.709153831

Alleles sampled from the same population

T11w=	0.48688675032911910	T12w=	1.0493489217997185	T22w=	0.99008582245855636

TTw=	0.99572118961867129	TSw=	0.93976591524561259

FATw=	5.6195725225540061E-002	T11/T22=	0.491762161

Time (units of 2NT generations) =	4.25000004E-02

Alleles sampled from different populations

T11b=	0.75998840886416563	T12b=	1.0499551273891234	T22b=	1.0474583111028573

TTb=	1.0450330390119984	TSb=	1.0187113208789882

FATb=	2.5187450683755830E-002	T11/T22=	0.725554824

Alleles sampled from the same population

T11w=	0.49804977240001075	T12w=	1.0502726989310018	T22w=	0.98986827254295362

TTw=	0.99582288429137300	TSw=	0.94068642252865942

FATw=	5.5367739215943623E-002	T11/T22=	0.503147542

Time (units of 2NT generations) =	4.50000018E-02

Alleles sampled from different populations

T11b=	0.77536517012624073	T12b=	1.0509057270943245	T22b=	1.0472402750492256

TTb=	1.0451813053681136	TSb=	1.0200527645569271

FATb=	2.4042279250618859E-002	T11/T22=	0.740388989

Alleles sampled from the same population

T11w=	0.50813762137589591	T12w=	1.0512205708354649	T22w=	0.98966227775596716

TTw=	0.99592752394647621	TSw=	0.94150981211796003

FATw=	5.4640232868431782E-002	T11/T22=	0.513445497

Time (units of 2NT generations) =	4.74999994E-02

Alleles sampled from different populations

T11b=	0.78926999895241490	T12b=	1.0518719295820933	T22b=	1.0470345070916971

TTb=	1.0453275980585757	TSb=	1.0212580562777689

FATb=	2.3025835944166850E-002	T11/T22=	0.753814697

Alleles sampled from the same population

T11w=	0.51726004290230443	T12w=	1.0521843092188430	T22w=	0.98946787745080267

TTw=	0.99603475682356502	TSw=	0.94224709399595286

FATw=	5.4001793069094695E-002	T11/T22=	0.522765875

Time (units of 2NT generations) =	5.00000007E-02

Alleles sampled from different populations

T11b=	0.80185267696573104	T12b=	1.0528467091476230	T22b=	1.0468409732056425

TTb=	1.0454721227128001	TSb=	1.0223421435816513

FATb=	2.2123955893850966E-002	T11/T22=	0.765973747

Alleles sampled from the same population

T11w=	0.52551526585220232	T12w=	1.0531568612111590	T22w=	0.98928503905890475

TTw=	0.99614426931424349	TSw=	0.94290806173823449

FATw=	5.3442266563112795E-002	T11/T22=	0.531207144

Time (units of 2NT generations) =	5.24999984E-02

Alleles sampled from different populations

T11b=	0.81324729260946271	T12b=	1.0538240710650513	T22b=	1.0466595750924210

TTb=	1.0456150615426649	TSb=	1.0233183468441251

FATb=	2.1324018291821401E-002	T11/T22=	0.776993096

Alleles sampled from the same population

T11w=	0.53299122792008424	T12w=	1.0541322077860842	T22w=	0.98911366933256895

TTw=	0.99625578184007690	TSw=	0.94350142519132052

FATw=	5.2952622820737316E-002	T11/T22=	0.538857400

Time (units of 2NT generations) =	5.49999997E-02

Alleles sampled from different populations

T11b=	0.82357390986548062	T12b=	1.0547989263433333	T22b=	1.0464901606111014

TTb=	1.0457565759354470	TSb=	1.0241985355365395

FATb=	2.0614778711406601E-002	T11/T22=	0.786986768

Alleles sampled from the same population

T11w=	0.54591211270177054	T12w=	1.0560716387612392	T22w=	0.98880471736205167

TTw=	0.99648383716730271	TSw=	0.94451545689602356

FATw=	5.2151754331519640E-002	T11/T22=	0.552092969

Time (units of 2NT generations) =	5.99999987E-02

Alleles sampled from different populations

T11b=	0.84144206319015646	T12b=	1.0567246362665428	T22b=	1.0461864583735252

TTb=	1.0460358864424348	TSb=	1.0257120188551885

FATb=	1.9429417145876093E-002	T11/T22=	0.804294527

Alleles sampled from the same population

T11w=	0.55149072989715953	T12w=	1.0570277941186503	T22w=	0.98866672786262200

TTw=	0.99659995980905258	TSw=	0.94494912806607578

FATw=	5.1827045781612346E-002	T11/T22=	0.557812572

Time (units of 2NT generations) =	6.49999976E-02

Alleles sampled from different populations

T11b=	0.85618992597643406	T12b=	1.0585973285591748	T22b=	1.0459278915051708

TTb=	1.0463110105196043	TSb=	1.0269540949522973

FATb=	1.8500154708010030E-002 T11/T22=	0.818593621

Alleles sampled from the same population

T11w=	0.56116803584393304	T12w=	1.0588978861663725	T22w=	0.98842248209867178

TTw=	0.99683551036831064	TSw=	0.94569703747319789

FATw=	5.1300813788443533E-002	T11/T22=	0.567741096

Time (units of 2NT generations) =	7.00000003E-02

Alleles sampled from different populations

T11b=	0.86840609889184406	T12b=	1.0603990790511684	T22b=	1.0457120986573969

TTb=	1.0465826951306203	TSb=	1.0279814986808415

FATb=	1.7773269648278633E-002	T11/T22=	0.830444753

Alleles sampled from the same population

T11w=	0.56918462084199861	T12w=	1.0606974840403942	T22w=	0.98821865433222922

TTw=	0.99707450334479664	TSw=	0.94631525098320612

FATw=	5.0908184083850339E-002	T11/T22=	0.575970352

Time (units of 2NT generations) =	7.50000030E-02

Alleles sampled from different populations

T11b=	0.87856388075845115	T12b=	1.0621184792786358	T22b=	1.0455364902233419

TTb=	1.0468515221586461	TSb=	1.0288392292768529

FATb=	1.7206158180532727E-002	T11/T22=	0.840299606

Alleles sampled from the same population

T11w=	0.57585090380272164	T12w=	1.0624150941140629	T22w=	0.98805279666848411

TTw=	0.99731599128003068	TSw=	0.94683260738190789

FATw=	5.0619246396850226E-002	T11/T22=	0.582813919

Time (units of 2NT generations) =	7.99999982E-02

Alleles sampled from different populations

T11b=	0.88704431443783394	T12b=	1.0637490204213309	T22b=	1.0453983662301403

TTb=	1.0471179434666316	TSb=	1.0295629610509096

FATb=	1.6765047839409242E-002	T11/T22=	0.848522782

Alleles sampled from the same population

T11w=	0.58141680211189017	T12w=	1.0640441391722182	T22w=	0.98792235720413768

TTw=	0.99755922240746986	TSw=	0.94727180169491298

FATw=	5.0410461437262022E-002	T11/T22=	0.588524818

Time (units of 2NT generations) =	8.50000009E-02

Alleles sampled from different populations

T11b=	0.89415457848012214	T12b=	1.0652878357322400	T22b=	1.0452950030228554

TTb=	1.0473823086651173	TSb=	1.0301809605685821

FATb=	1.6423179916470243E-002	T11/T22=	0.855408847

Alleles sampled from the same population

T11w=	0.58608379228913454	T12w=	1.0655816974803878	T22w=	0.98782476187092083

TTw=	0.99780360058480710	TSw=	0.94765066491274230

FATw=	5.0263334029532869E-002	T11/T22=	0.593307436

Time (units of 2NT generations) =	9.00000036E-02

Alleles sampled from different populations

T11b=	0.90014266939354570	T12b=	1.0667347254272788	T22b=	1.0452237158254498

TTb=	1.0476448870894601	TSb=	1.0307156111822595

FATb=	1.6159364796055176E-002	T11/T22=	0.861196160

Alleles sampled from the same population

T11w=	0.59001453821537964	T12w=	1.0670275249936578	T22w=	0.98775747349918996

TTw=	0.99804865341535609	TSw=	0.94798317997080894

FATw=	5.0163359544869279E-002	T11/T22=	0.597327352

Time (units of 2NT generations) =	9.49999988E-02

Alleles sampled from different populations

T11b=	0.90520912707825385	T12b=	1.0680914038064127	T22b=	1.0451819027464406

TTb=	1.0479058851805538	TSb=	1.0311846251796219

FATb=	1.5956833755209621E-002	T11/T22=	0.866078079

Alleles sampled from the same population

T11w=	0.59334058040397053	T12w=	1.0683833003384744	T22w=	0.98771803335955632

TTw=	0.99829400688620584	TSw=	0.94828028806399778

FATw=	5.0099187691416303E-002	T11/T22=	0.600718558

Time (units of 2NT generations) =	0.100000001

Alleles sampled from different populations

T11b=	0.90951640284813451	T12b=	1.0693609209272172	T22b=	1.0451670745960733

TTb=	1.0481654602181998	TSb=	1.0316020074212795

FATb=	1.5802326469975703E-002	T11/T22=	0.870211482

Alleles sampled from the same population

T11w=	0.59616847938428275	T12w=	1.0696520447784534	T22w=	0.98770408930647280

TTw=	0.99853936519220743	TSw=	0.94855052831425379

FATw=	5.0061959117987609E-002	T11/T22=	0.603590190

Time (units of 2NT generations) =	0.150000006

Alleles sampled from different populations

T11b=	0.93070648605327999	T12b=	1.0782581247651568	T22b=	1.0460556270549819

TTb=	1.0506985852327964	TSb=	1.0345207129548117

FATb=	1.5397253318277015E-002	T11/T22=	0.889729440

Alleles sampled from the same population

T11w=	0.61008624508874321	T12w=	1.0785452079202469	T22w=	0.98854425132880464

TTw=	1.0009598434528637	TSw=	0.95069845070479853

FATw=	5.0213195940694177E-002	T11/T22=	0.617156208

Time (units of 2NT generations) =	0.200000003

Alleles sampled from different populations

T11b=	0.93765203199979130	T12b=	1.0831027966114239	T22b=	1.0478946750258054

TTb=	1.0531297104809567	TSb=	1.0368704107232039

FATb=	1.5439028636204166E-002	T11/T22=	0.894796073

Alleles sampled from the same population

T11w=	0.61465220450263358	T12w=	1.0833881620863208	T22w=	0.99028241076125667

TTw=	1.0032851439371822	TSw=	0.95271939013539442

FATw=	5.0400181949623035E-002	T11/T22=	0.620683730

Time (units of 2NT generations) =	0.250000000

Alleles sampled from different populations

T11b=	0.94120551823829013	T12b=	1.0863322410405130	T22b=	1.0500091539073417

TTb=	1.0554592732346220	TSb=	1.0391287903404365

FATb=	1.5472395106386427E-002	T11/T22=	0.896378398

Alleles sampled from the same population

T11w=	0.61698983075015590	T12w=	1.0866164271600707	T22w=	0.99228076414768929

TTw=	1.0055082741559427	TSw=	0.95475167080793599

FATw=	5.0478553635586398E-002 T11/T22=	0.621789575

Time (units of 2NT generations) =	0.500000000

Alleles sampled from different populations

T11b=	0.95105296840551912	T12b=	1.0972904745945269	T22b=	1.0600514079534005

TTb=	1.0656644555533246	TSb=	1.0491515639986124

FATb=	1.5495394885942981E-002	T11/T22=	0.897176266

Alleles sampled from the same population

T11w=	0.62347211492932830	T12w=	1.0975703190785335	T22w=	1.0017712676274020

TTw=	1.0152321053616249	TSw=	0.96394135235759471

FATw=	5.0521208631163717E-002	T11/T22=	0.622369707

Time (units of 2NT generations) =	0.750000000

Alleles sampled from different populations

T11b=	0.95831303085108877	T12b=	1.1056815884754432	T22b=	1.0681474613468760

TTb=	1.0738052599250603	TSb=	1.0571640182972972

FATb=	1.5497448419021898E-002	T11/T22=	0.897172987

Alleles sampled from the same population

T11w=	0.62825185627997349	T12w=	1.1059580543209853	T22w=	1.0094224762618225

TTw=	1.0229871741126535	TSw=	0.97130541426363759

FATw=	5.0520437750204539E-002	T11/T22=	0.622387409

Time (units of 2NT generations) =	1.00000000

Alleles sampled from different populations

T11b=	0.96410030939513913	T12b=	1.1123721415570265	T22b=	1.0746049016869017

TTb=	1.0802979589406068	TSb=	1.0635544424577255

FATb=	1.5498980021494080E-002	T11/T22=	0.897167265

Alleles sampled from the same population

T11w=	0.63206197779174278	T12w=	1.1126459131926336	T22w=	1.0155251067539692

TTw=	1.0291722206233067	TSw=	0.97717879385774653

FATw=	5.0519656208822816E-002	T11/T22=	0.622399151

Time (units of 2NT generations) =	2.00000000

Alleles sampled from different populations

T11b=	0.97767458900944271	T12b=	1.1280651106146165	T22b=	1.0897511273883405

TTb=	1.0955268789852812	TSb=	1.0785434735504507

FATb=	1.5502499993940155E-002	T11/T22=	0.897154033

Alleles sampled from the same population

T11w=	0.64099876123437161	T12w=	1.1283325628649572	T22w=	1.0298391095054762

TTw=	1.0436795276274720	TSw=	0.99095507467836574

FATw=	5.0517856826186147E-002	T11/T22=	0.622426093

Time (units of 2NT generations) =	3.00000000

Alleles sampled from different populations

T11b=	0.98316678861207518	T12b=	1.1344145391788265	T22b=	1.0958793417626578

TTb=	1.1016885517660624	TSb=	1.0846080864475995

FATb=	1.5503896533264339E-002	T11/T22=	0.897148728

Alleles sampled from the same population

T11w=	0.64461461414647225	T12w=	1.1346794345844280	T22w=	1.0356306036469798

TTw=	1.0495492333207155	TSw=	0.99652900469692907

FATw=	5.0517142922427150E-002	T11/T22=	0.622436821

Time (units of 2NT generations) =	4.00000000

Alleles sampled from different populations

T11b=	0.98538895122756309	T12b=	1.1369835395440602	T22b=	1.0983588381152261

TTb=	1.1041815855035397	TSb=	1.0870618494264599

FATb=	1.5504457148932338E-002	T11/T22=	0.897146642

Alleles sampled from the same population

T11w=	0.64607760051727392	T12w=	1.1372474004415949	T22w=	1.0379738618495584

TTw=	1.0519241361828022	TSw=	0.99878423571633002

FATw=	5.0516856338428617E-002	T11/T22=	0.622441113

Time (units of 2NT generations) =	5.00000000

Alleles sampled from different populations

T11b=	0.98628804577017737	T12b=	1.1380229658183827	T22b=	1.0993620507619912

TTb=	1.1051902754222236	TSb=	1.0880546502628099

FATb=	1.5504683257249297E-002	T11/T22=	0.897145808

Alleles sampled from the same population

T11w=	0.64666952977246839	T12w=	1.1382864081504396	T22w=	1.0389219520780550

TTw=	1.0528850299480286	TSw=	0.99969670984749648

FATw=	5.0516740752936218E-002	T11/T22=	0.622442842

**Scaled gene conversion rate=	4**

Equilibrium results for subdivided population

Alleles sampled from different populations

T11b=	0.65832851359167155	T12b=	1.5786581839213418	T22b=	1.1253614806246386

TTb=	1.2022845575477157	TSb=	1.0786581839213418

FATb=	0.10282621767889377	T11/T22=	0.584992945

Alleles sampled from the same population

T11w=	0.43180143993937109	T12w=	1.5789213418160786	T22w=	1.0634851557282499

TTw=	1.1499468320661703	TSw=	1.0003167841493621

FATw=	0.13011910093961476	T11/T22=	0.406024873

Time (units of 2NT generations) =	2.49999994E-03

Alleles sampled from different populations

T11b=	1.1620979840086809E-002	T12b=	1.0540391898923172	T22b=	1.0521388899114541

TTb=	1.0420757648072958	TSb=	0.94808709890431742

FATb=	9.0193697116024096E-002	T11/T22=	1.10451002E-02

Alleles sampled from the same population

T11w=	7.5042152747256674E-003	T12w=	1.0543142873459301	T22w=	0.99428717262944577

TTw=	0.99522422370486585	TSw=	0.89560887689397384

FATw=	0.10009337035633992	T11/T22=	7.54733197E-03

Time (units of 2NT generations) =	4.99999989E-03

Alleles sampled from different populations

T11b=	2.3201186595337431E-002	T12b=	1.0560075183332016	T22b=	1.0518810434850940

TTb=	1.0423370103888558	TSb=	0.94901305779611844

FATb=	8.9533377077267717E-002	T11/T22=	2.20568534E-02

Alleles sampled from the same population

T11w=	1.5100273978622680E-002	T12w=	1.0562840322698954	T22w=	0.99404343381983085

TTw=	0.99545730994243042	TSw=	0.89614911783570994

FATw=	9.9761377122705275E-002	T11/T22=	1.51907587E-02

Time (units of 2NT generations) =	7.49999983E-03

Alleles sampled from different populations

T11b=	3.4508084892729274E-002	T12b=	1.0579705854448260	T22b=	1.0516260926588619

TTb=	1.0425969212826742	TSb=	0.94991429188224863

FATb=	8.8895936203610804E-002	T11/T22=	3.28140259E-02

Alleles sampled from the same population

T11w=	2.2517061416104101E-002	T12w=	1.0582469019180243	T22w=	0.99380251643402095

TTw=	0.99568965127096254	TSw=	0.89667397093222934

FATw=	9.9444320037215594E-002	T11/T22=	2.26574801E-02

Time (units of 2NT generations) =	9.99999978E-03

Alleles sampled from different populations

T11b=	4.5548490717030043E-002	T12b=	1.0599282879645253	T22b=	1.0513740220454082

TTb=	1.0428555345975656	TSb=	0.95079146891257038

FATb=	8.8280747074447330E-002	T11/T22=	4.33228239E-02

Alleles sampled from the same population

T11w=	2.9759051182332574E-002	T12w=	1.0602044116713807	T22w=	0.99356432081627455

TTw=	0.99592148447385431	TSw=	0.89718379385288038

FATw=	9.9142042982572121E-002	T11/T22=	2.99518108E-02

Time (units of 2NT generations) =	1.25000002E-02

Alleles sampled from different populations

T11b=	5.6329048066894316E-002	T12b=	1.0618805344785984	T22b=	1.0511248158189339

TTb=	1.0431128875001532	TSb=	0.95164523904373000

FATb=	8.7687200064824999E-002	T11/T22=	5.35893030E-02

Alleles sampled from the same population

T11w=	3.6830601159208998E-002	T12w=	1.0621564700369781	T22w=	0.99332883200969535

TTw=	0.99615282454610155	TSw=	0.89767900892464669

FATw=	9.8854124783839814E-002	T11/T22=	3.70779559E-02

Time (units of 2NT generations) =	1.49999997E-02

Alleles sampled from different populations

T11b=	6.6856233299915635E-002	T12b=	1.0638272372254585	T22b=	1.0508784581214323

TTb=	1.0433690161119420	TSb=	0.95247623563928063

FATb=	8.7114701576407105E-002	T11/T22=	6.36193752E-02

Alleles sampled from the same population

T11w=	4.3735959270986835E-002	T12w=	1.0641029891366873	T22w=	0.99309603502800603

TTw=	0.99638368600999849	TSw=	0.89816002745230406

FATw=	9.8580155352632648E-002	T11/T22=	4.40400094E-02

Time (units of 2NT generations) =	1.75000001E-02

Alleles sampled from different populations

T11b=	7.7136359374530006E-002	T12b=	1.0657683119946213	T22b=	1.0506349330664089

TTb=	1.0436239555365685	TSb=	0.95328507569722098

FATb=	8.6562673614463681E-002	T11/T22=	7.34188035E-02

Alleles sampled from the same population

T11w=	5.0479266266642564E-002	T12w=	1.0660438846464351	T22w=	0.99286591485802633

TTw=	0.99661408293402609	TSw=	0.89862724999888788

FATw=	9.8319735405168585E-002	T11/T22=	5.08419760E-02

Time (units of 2NT generations) =	1.99999996E-02

Alleles sampled from different populations

T11b=	8.7175579984404283E-002	T12b=	1.0677036780285485	T22b=	1.0503942247424818

TTb=	1.0438777398863932	TSb=	0.95407236026667408

FATb=	8.6030553376387542E-002	T11/T22=	8.29932019E-02

Alleles sampled from the same population

T11w=	5.7064558431666468E-002	T12w=	1.0679790756979703	T22w=	0.99263845646307858

TTw=	0.99684402894504498	TSw=	0.89908106665993737

FATw=	9.8072476181223278E-002	T11/T22=	5.74877560E-02

Time (units of 2NT generations)=	1.25000002E-02

Alleles sampled from different populations

T11b=	5.6329048066894316E-002 T12b=	1.0618805344785984	T22b=	1.0511248158189339

TTb=	1.0431128875001532	TSb=	0.95164523904373000

FATb=	8.7687200064824999E-002 T11/T22=	5.35893030E-02

Alleles sampled from the same population

T11w=	3.6830601159208998E-002 T12w=	1.0621564700369781	T22w=	0.99332883200969535

TTw=	0.99615282454610155	TSw=	0.89767900892464669

FATw=	9.8854124783839814E-002 T11/T22=	3.70779559E-02

Time (units of 2NT generations) =	1.49999997E-02

Alleles sampled from different populations

T11b=	6.6856233299915635E-002	T12b=	1.0638272372254585	T22b=	1.0508784581214323

TTb=	1.0433690161119420	TSb=	0.95247623563928063

FATb=	8.7114701576407105E-002	T11/T22=	6.36193752E-02

Alleles sampled from the same population

T11w=	4.3735959270986835E-002	T12w=	1.0641029891366873	T22w=	0.99309603502800603

TTw=	0.99638368600999849	TSw=	0.89816002745230406

FATw=	9.8580155352632648E-002	T11/T22=	4.40400094E-02

Time (units of 2NT generations) =	1.75000001E-02

Alleles sampled from different populations

T11b=	7.7136359374530006E-002	T12b=	1.0657683119946213	T22b=	1.0506349330664089

TTb=	1.0436239555365685	TSb=	0.95328507569722098

FATb=	8.6562673614463681E-002	T11/T22=	7.34188035E-02

Alleles sampled from the same population

T11w=	5.0479266266642564E-002	T12w=	1.0660438846464351	T22w=	0.99286591485802633

TTw=	0.99661408293402609	TSw=	0.89862724999888788

FATw=	9.8319735405168585E-002	T11/T22=	5.08419760E-02

Time (units of 2NT generations) =	1.99999996E-02

Alleles sampled from different populations

T11b=	8.7175579984404283E-002 T12b=	1.0677036780285485	T22b=	1.0503942247424818

TTb=	1.0438777398863932	TSb=	0.95407236026667408

FATb=	8.6030553376387542E-002 T11/T22=	8.29932019E-02

Alleles sampled from the same population

T11w=	5.7064558431666468E-002	T12w=	1.0679790756979703	T22w=	0.99263845646307858

TTw=	0.99684402894504498	TSw=	0.89908106665993737

FATw=	9.8072476181223278E-002	T11/T22=	5.74877560E-02

Time (units of 2NT generations) =	2.50000004E-02

Alleles sampled from different populations

T11b=	0.10655514733736160	T12b=	1.0715569775539389	T22b=	1.0499211945387903

TTb=	1.0443819750095027	TSb=	0.95558458981864747

FATb=	8.5023858430769317E-002	T11/T22=	0.101488709

Alleles sampled from the same population

T11w=	6.9776736886457136E-002	T12w=	1.0718320376607440	T22w=	0.99219146475375330

TTw=	0.99730262059833874	TSw=	0.89994999196702374

FATw=	9.7615935845939750E-002	T11/T22=	7.03258812E-02

Time (units of 2NT generations) =	2.99999993E-02

Alleles sampled from different populations

T11b=	0.12504113021347130	T12b=	1.0753865552272543	T22b=	1.0494592398515576

TTb=	1.0448819755228023	TSb=	0.95701742888774899

FATb=	8.4090403216201204E-002	T11/T22=	0.119148150

Alleles sampled from the same population

T11w=	8.1902794430157022E-002	T12w=	1.0756612936197443	T22w=	0.99175493925944380

TTw=	0.99775956159600510	TSw=	0.90076972477651518

FATw=	9.7207624514613622E-002	T11/T22=	8.25837031E-02

Time (units of 2NT generation s)=	3.50000001E-02

Alleles sampled from different populations

T11b=	0.14267742274858830	T12b=	1.0791918798521030	T22b=	1.0490082328363208

TTb=	1.0453779811982844	TSb=	0.95837515182754762

FATb=	8.3226192760448403E-002	T11/T22=	0.136011735

Alleles sampled from the same population

T11w=	9.3471521791083645E-002	T12w=	1.0794663115892464	T22w=	0.99132875916438690

TTw=	0.99821494622712870	TSw=	0.90154303542705660

FATw=	9.6844783947039659E-002 T11/T22=	9.42891240E-02

Time (units of 2NT generations) =	3.99999991E-02

Alleles sampled from different populations

T11b=	0.15950573330363205	T12b=	1.0829724668891898	T22b=	1.0485680455600177

TTb=	1.0458702182767050	TSb=	0.95966181433437914

FATb=	8.2427439309222028E-002	T11/T22=	0.152117670

Alleles sampled from the same population

T11w=	0.10451027602396135	T12w=	1.0832466062800936	T22w=	0.99091280356935196

TTw=	0.99866886278183165	TSw=	0.90227255081481295

FATw=	9.6524799720402732E-002	T11/T22=	0.105468690

Time (units of 2NT generations) =	5.00000007E-02

Alleles sampled from different populations

T11b=	0.19089496276331441	T12b=	1.0904577079707887	T22b=	1.0477196185896986

TTb=	1.0468442281200310	TSb=	0.96203715300706016

FATb=	8.1012124664689655E-002	T11/T22=	0.182200432

Alleles sampled from the same population

T11w=	0.12510055111516871	T12w=	1.0907313027827117	T22w=	0.99011108237599210

TTw=	0.99957261673659348	TSw=	0.90361002924990985

FATw=	9.6003617826168952E-002	T11/T22=	0.126350015

Time (units of 2NT generations) =	5.99999987E-02

Alleles sampled from different populations

T11b=	0.21950278513828025	T12b=	1.0978392404501756	T22b=	1.0469129370253161

TTb=	1.0478055701229205	TSb=	0.96417192183661249

FATb=	7.9817907702568136E-002	T11/T22=	0.209666699

Alleles sampled from the same population

T11w=	0.14386643625311613	T12w=	1.0981123398237154	T22w=	0.98934880995417074

TTw=	1.0004714215936783	TSw=	0.90480057258406532

FATw=	9.5625768957214419E-002	T11/T22=	0.145415276

Time (units of 2NT generations) =	7.00000003E-02

Alleles sampled from different populations

T11b=	0.24559463884190724	T12b=	1.1051146207743714	T22b=	1.0461469827982290

TTb=	1.0487556341943716	TSb=	0.96609174840259693

FATb=	7.8820921763414376E-002	T11/T22=	0.234761119

Alleles sampled from the same population

T11w=	0.16098203607125994	T12w=	1.1053872690831996	T22w=	0.98862502421706222

TTw=	1.0013657984115090	TSw=	0.90586072540248197

FATw=	9.5374810244696850E-002	T11/T22=	0.162834272

Time (units of 2NT generations) =	7.99999982E-02

Alleles sampled from different populations

T11b=	0.26941021282061883	T12b=	1.1122819345696935	T22b=	1.0454207438608383

TTb=	1.0496956528780301	TSb=	0.96781969075681640

FATb=	7.7999715342945586E-002	T11/T22=	0.257705063

Alleles sampled from the same population

T11w=	0.17660456584282613	T12w=	1.1125541718853955	T22w=	0.98793876877064901

TTw=	1.0022561993020251	TSw=	0.90680534847786676

FATw=	9.5235979473742072E-002	T11/T22=	0.178760633

Time (units of 2NT generations) =	9.00000036E-02

Alleles sampled from different populations

T11b=	0.29116595432493453	T12b=	1.1193397379022878	T22b=	1.0447332160835483

TTb=	1.0506267173933352	TSb=	0.96937648990768699

FATb=	7.7335009799898047E-002	T11/T22=	0.278698862

Alleles sampled from the same population

T11w=	0.19087599635404545	T12w=	1.1196116004138625	T22w=	0.98728909470602455

TTw=	1.0031430147499156	TSw=	0.90764778487082665

FATw=	9.5196027360960112E-002	T11/T22=	0.193333432

Time (units of 2NT generations) =	0.100000001

Alleles sampled from different populations

T11b=	0.31105733220211762	T12b=	1.1262870044454369	T22b=	1.0440834049029857

TTb=	1.0515497920936183	TSb=	0.97078079763289882

FATb=	7.6809481650802036E-002	T11/T22=	0.297923833

Alleles sampled from the same population

T11w=	0.20392453842150021	T12w=	1.1265585248378938	T22w=	0.98667506215666134

TTw=	1.0040265802019317	TSw=	0.90840000978314528

FATw=	9.5243066572554103E-002	T11/T22=	0.206678510

Time (units of 2NT generations) =	0.125000000

Alleles sampled from different populations

T11b=	0.35374603445557318	T12b=	1.1431680620428089	T22b=	1.0426174626994884

TTb=	1.0538278562988470	TSb=	0.97373031987509695

FATb=	7.6006281239386575E-002	T11/T22=	0.339286506

Alleles sampled from the same population

T11w=	0.23192831403130748	T12w=	1.1434388501329549	T22w=	0.98528984138025655

TTw=	1.0062230476822527	TSw=	0.90995368864536164

FATw=	9.5673975326483696E-002	T11/T22=	0.235390946

Time (units of 2NT generations) =	0.150000006

Alleles sampled from different populations

T11b=	0.38816301880345405	T12b=	1.1593529973666525	T22b=	1.0413661214366221

TTb=	1.0560717280776959	TSb=	0.97604581117330524

FATb=	7.5776971181736874E-002	T11/T22=	0.372744054

Alleles sampled from the same population

T11w=	0.25450639381602164	T12w=	1.1596231969378503	T22w=	0.98410741431744697

TTw=	1.0084042449841055	TSw=	0.91114731226730450

FATw=	9.6446373763860893E-002 T11/T22=	0.258616477

Time (units of 2NT generations)=	0.174999997

Alleles sampled from different populations

T11b=	0.41611064897149830	T12b=	1.1748491855035095	T22b=	1.0403149306839665

TTb=	1.0582890537343597	TSb=	0.97789450251271970

FATb=	7.5966533848151996E-002	T11/T22=	0.399985284

Alleles sampled from the same population

T11w=	0.27284097754614128	T12w=	1.1751189081671960	T22w=	0.98311412537489351

TTw=	1.0105722547992206	TSw=	0.91208681059201835

FATw=	9.7455123806826971E-002	T11/T22=	0.277527273

Time (units of 2NT generations) =	0.200000003

Alleles sampled from different populations

T11b=	0.43898743007837221	T12b=	1.1896707655138969	T22b=	1.0394500622869491

TTb=	1.0604851625457141	TSb=	0.97940379906609154

FATb=	7.6456857995999994E-002	T11/T22=	0.422326624

Alleles sampled from the same population

T11w=	0.28784936933681754	T12w=	1.1899400979864758	T22w=	0.98229690704645378

TTw=	1.0127282060385614	TSw=	0.91285215327549019

FATw=	9.8620787065614945E-002	T11/T22=	0.293037027

Time (units of 2NT generations) =	0.250000000

Alleles sampled from different populations

T11b=	0.47362941257297730	T12b=	1.2173687565726818	T22b=	1.0382272754220103

TTb=	1.0648267634006410	TSb=	0.98176748913710710

FATb=	7.8002617062586710E-002	T11/T22=	0.456190497

Alleles sampled from the same population

T11w=	0.31057748174470962	T12w=	1.2176374961086953	T22w=	0.98114152655370068

TTw=	1.0170051606255099	TSw=	0.91408512207280157

FATw=	0.10119913107363909	T11/T22=	0.316547066

Time (units of 2NT generations) =	0.300000012

Alleles sampled from different populations

T11b=	0.49818425443773601	T12b=	1.2426294260357365	T22b=	1.0376004367284797

TTb=	1.0691114929808785	TSb=	0.98365881849940529

FATb=	7.9928683811279155E-002	T11/T22=	0.480131119

Alleles sampled from the same population

T11w=	0.32668876353533605	T12w=	1.2428977392584493	T22w=	0.98054930271495944

TTw=	1.0212334159009915	TSw=	0.91516324879699718

FATw=	0.10386476338557038	T11/T22=	0.333169132

Time (units of 2NT generations) =	0.349999994

Alleles sampled from different populations

T11b=	0.51634469383483950	T12b=	1.2656553447589578	T22b=	1.0374827871846182

TTb=	1.0733424666145015	TSb=	0.98536897784964028

FATb=	8.1962180293065345E-002	T11/T22=	0.497689903

Alleles sampled from the same population

T11w=	0.33860538185393557	T12w=	1.2659233338644418	T22w=	0.98043824951237990

TTw=	1.0254072360191666	TSw=	0.91625496274653551

FATw=	0.10644773065614577	T11/T22=	0.345361263

Time (units of 2NT generations) =	0.400000006

Alleles sampled from different populations

T11b=	0.53036006990414419	T12b=	1.2866503570330543	T22b=	1.0377975799415187

TTb=	1.0775167047176215	TSb=	0.98705382893778126

FATb=	8.3954963652788428E-002	T11/T22=	0.511043847

Alleles sampled from the same population

T11w=	0.34780275640993019	T12w=	1.2869180855953346	T22w=	0.98073584242267897

TTw=	1.0295193153336295	TSw=	0.91744253382140417

FATw=	0.10886321397078924	T11/T22=	0.354634494

Time (units of 2NT generations) =	0.449999988

Alleles sampled from different populations

T11b=	0.54161113484223533	T12b=	1.3058095757741963	T22b=	1.0384772962738673

TTb=	1.0816284449696103	TSb=	0.98879068013070404

FATb=	8.5831475004815250E-002	T11/T22=	0.521543562

Alleles sampled from the same population

T11w=	0.35518658546379989	T12w=	1.3060770841905369	T22w=	0.98137827743638895

TTw=	1.0335621457324098	TSw=	0.91875910823913010

FATw=	0.11107511818936366	T11/T22=	0.361926287

Time (units of 2NT generations) =	0.500000000

Alleles sampled from different populations

T11b=	0.55095470401794799	T12b=	1.3233145158376374	T22b=	1.0394627779416987

TTb=	1.0856710100237303	TSb=	0.99061197054932371

FATb=	8.7557868448867260E-002	T11/T22=	0.530037940

Alleles sampled from the same population

T11w=	0.36131890612281881	T12w=	1.3235818305725298	T22w=	0.98230965111410640

TTw=	1.0375287359667098	TSw=	0.92021057661497763

FATw=	0.11307461209006597	T11/T22=	0.367825866

Time (units of 2NT generations) =	0.750000000

Alleles sampled from different populations

T11b=	0.58256972088072767	T12b=	1.3912969587428829	T22b=	1.0473859104492429

TTb=	1.1046417372464130	TSb=	1.0009042914923914

FATb=	9.3910489035669009E-002	T11/T22=	0.556213081

Time (units of 2NT generations) =	1.00000000

Alleles sampled from different populations

T11b=	0.60189249423715685	T12b=	1.4367940467145277	T22b=	1.0575228752203918

TTb=	1.1212353822795040	TSb=	1.0119598371220684

FATb=	9.7459950769012704E-002	T11/T22=	0.569153190

Alleles sampled from the same population

T11w=	0.39475396469454521	T12w=	1.4370599773150619	T22w=	0.99937702744046231

TTw=	1.0721137277904311	TSw=	0.93891472116587060

FATw=	0.12423962418527790	T11/T22=	0.395000041

Time (units of 2NT generations) =	1.50000000

Alleles sampled from different populations

T11b=	0.62442098276893632	T12b=	1.4918743671977910	T22b=	1.0771167233215408

TTb=	1.1472461418137399	TSb=	1.0318471492662804

FATb=	0.10058782360777374	T11/T22=	0.579715252

Alleles sampled from the same population

T11w=	0.40954248149085998	T12w=	1.4921393849640723	T22w=	1.0178934773608050

TTw=	1.0971742307706938	TSw=	0.95705837777381053

FATw=	0.12770610999354304	T11/T22=	0.402343154

Time (units of 2NT generations) =	2.00000000

Alleles sampled from different populations

T11b=	0.63672529980558479	T12b=	1.5228931523552991	T22b=	1.0923081172320868

TTb=	1.1652575953800000	TSb=	1.0467498354894365

FATb=	0.10170091176442164	T11/T22=	0.582917273

Alleles sampled from the same population

T11w=	0.41761968036106217	T12w=	1.5231575552173371	T22w=	1.0322494852262798

TTw=	1.1144666397760181	TSw=	0.97078650473975803

FATw=	0.12892277786362139	T11/T22=	0.404572427

Time (units of 2NT generations) =	3.00000000

Alleles sampled from different populations

T11b=	0.64895780512021262	T12b=	1.5543384222054311	T22b=	1.1103835399313426

TTb=	1.1856811613925673	TSb=	1.0642409664502297

FATb=	0.10242230280500342	T11/T22=	0.584444702

Alleles sampled from the same population

T11w=	0.42564986744045480	T12w=	1.5546021367215765	T22w=	1.0493308972420983

TTw=	1.1340429100503879	TSw=	0.98696279426193401

FATw=	0.12969537085851435	T11/T22=	0.405639321

Time (units of 2NT generations) =	4.00000000

Alleles sampled from different populations

T11b=	0.65416437923053483	T12b=	1.5678405690398736	T22b=	1.1186544478542344

TTb=	1.1948630489814125	TSb=	1.0722054409918644

FATb=	0.10265411428875493	T11/T22=	0.584777892

Alleles sampled from the same population

T11w=	0.42906781576069375	T12w=	1.5681039756107116	T22w=	1.0571469633474504

TTw=	1.1428384340789699	TSw=	0.99433904858877475

FATw=	0.12993908943032084	T11/T22=	0.405873388

Time (units of 2NT generations) =	5.00000000

Alleles sampled from different populations

T11b=	0.65647012046141662	T12b=	1.5738296232334745	T22b=	1.1223642387448927

TTb=	1.1989690667700028	TSb=	1.0757748269165450

FATb=	0.10275014032291963	T11/T22=	0.584899366

Alleles sampled from the same population

T11w=	0.43058146270627307	T12w=	1.5740928922121580	T22w=	1.0606527414415412

TTw=	1.1467712557928997	TSw=	0.99764561356801440

FATw=	0.13003957107538155	T11/T22=	0.405958951

**Scaled gene conversion rate= 0.4**

Equilibrium results for subdivided population

Alleles sampled from different populations

T11b=	0.46097604595137653	T12b=	6.0726090661392638	T22b=	1.1405682906045846

TTb=	2.0215397077542949	TSb=	1.0726090661392638

FATb=	0.46940984536444608	T11/T22=	0.404163480

Alleles sampled from the same population

T11w=	0.30250154734745360	T12w=	6.0728722240340005	T22w=	1.0778518104606849

TTw=	1.9692019822727496	TSw=	1.0003167841493619

FATw=	0.49201920719435355	T11/T22=	0.280652255

Time (units of 2NT generations) =	2.49999994E-03

Alleles sampled from different populations

T11b=	3.3669678394421254E-003	T12b=	1.0545033399372157	T22b=	1.0521382639147969

TTb=	1.0420762646380788	TSb=	0.94726113430726144

FATb=	9.0986747849733818E-002	T11/T22=	3.20011913E-03

Alleles sampled from the same population

T11w=	2.2417407886499152E-003	T12w=	1.0547659235233589	T22w=	0.99428295363745867

TTw=	0.99524947608843262	TSw=	0.89507883235257779

FATw=	0.10064877816318907	T11/T22=	2.25463067E-03

Time (units of 2NT generations) =	4.99999989E-03

Alleles sampled from different populations

T11b=	6.7464766976160773E-003	T12b=	1.0569491871585575	T22b=	1.0518778264635178

TTb=	1.0423393578909661	TSb=	0.94736469148692759

FATb=	9.1116837990466992E-002	T11/T22=	6.41374569E-03

Alleles sampled from the same population

T11w=	4.4584653106743977E-003	T12w=	1.0572133932301888	T22w=	0.99403675549843551

TTw=	0.99551276738827355	TSw=	0.89507892647965948

FATw=	0.10088654229126781	T11/T22=	4.48521180E-03

Time (units of 2NT generations) =	7.49999983E-03

Alleles sampled from different populations

T11b=	1.0070110694841428E-002	T12b=	1.0593938702377566	T22b=	1.0516183418008593

TTb=	1.0426024546084407	TSb=	0.94746351869025758

FATb=	9.1251402198082698E-002	T11/T22=	9.57582239E-03

Alleles sampled from the same population

T11w=	6.6385403280067035E-003	T12w=	1.0596580710242822	T22w=	0.99379154430497163

TTw=	0.99577598907467801	TSw=	0.89507624390727514

FATw=	0.10112690632456189	T11/T22=	6.68001268E-03

Time (units of 2NT generations) =	9.99999978E-03

Alleles sampled from different populations

T11b=	1.3338828704771988E-002	T12b=	1.0618373792850706	T22b=	1.0513598076241786

TTb=	1.0428655607339452	TSb=	0.94755770973223796

FATb=	9.1390352304501987E-002	T11/T22=	1.26872156E-02

Alleles sampled from the same population

T11w=	8.7825950752806747E-003	T12w=	1.0621015748424458	T22w=	0.99354723131943679

TTw=	0.99603936679113692	TSw=	0.89507076769502114

FATw=	0.10137008883634646	T11/T22=	8.83963518E-03

Time (units of 2NT generations) =	1.49999997E-02

Alleles sampled from different populations

T11b=	1.9715270168932930E-002	T12b=	1.0667208685157332	T22b=	1.0508455799173255

TTb=	1.0433918287675552	TSb=	0.94773254894248626

FATb=	9.1681070512178331E-002	T11/T22=	1.87613387E-02

Alleles sampled from the same population

T11w=	1.2965105676667199E-002	T12w=	1.0669850538979944	T22w=	0.99306128975211483

TTw=	0.99656660545761877	TSw=	0.89505167134456998

FATw=	0.10186467573477798	T11/T22=	1.30556952E-02

Time (units of 2NT generations) =	1.99999996E-02

Alleles sampled from different populations

T11b=	2.5883147648977922E-002	T12b=	1.0715996417725355	T22b=	1.0503351217919308

TTb=	1.0439182156470102	TSb=	0.94788992437763553

FATb=	9.1988328041442635E-002	T 11/T22=	2.46427525E-02

Alleles sampled from the same population

T11w=	1.7010815836103754E-002	T12w=	1.0718638173463848	T22w=	0.99257891042969593

TTw=	0.99709451272876415	TSw=	0.89502210097033674

FATw=	0.10236984604306387	T11/T22=	1.71379987E-02

Time (units of 2NT generations) =	2.99999993E-02

Alleles sampled from different populations

T11b=	3.7621354019372610E-002	T12b=	1.0813429915999757	T22b=	1.0493254286540543

TTb=	1.0449715492379734	TSb=	0.94815502119058614

FATb=	9.2649917711146279E-002	T11/T22=	3.58528942E-02

Alleles sampled from the same population

T11w=	2.4710294494734719E-002	T12w=	1.0816071486065439	T22w=	0.99162475759796798

TTw=	0.99815244334847941	TSw=	0.89493331128764475

FATw=	0.10341018824195614	T11/T22=	2.49189977E-02

Time (units of 2NT generations) =	3.99999991E-02

Alleles sampled from different populations

T11b=	4.8607366148727404E-002	T12b=	1.0910673379847209	T22b=	1.0483305580861702

TTb=	1.0460259465485349	TSb=	0.94835823889242588

FATb=	9.3370253365485989E-002	T11/T22=	4.63664494E-02

Alleles sampled from the same population

T11w=	3.1916396917245904E-002	T12w=	1.0913314777433349	T22w=	0.99068461205698122

TTw=	0.99921336572912767	TSw=	0.89480779054300763

FATw=	0.10448776884598132	T11/T22=	3.22165079E-02

Time (units of 2NT generations) =	5.00000007E-02

Alleles sampled from different populations

T11b=	5.8891491227946458E-002	T12b=	1.1007726004132197	T22b=	1.0473503417746532

TTb=	1.0470817598241282	TSb=	0.94850445671998251

FATb=	9.4144800230980152E-002	T11/T22=	5.62290289E-02

Alleles sampled from the same population

T11w=	3.8662120366150798E-002	T12w=	1.1010367241528698	T22w=	0.98975831475062659

TTw=	1.0002774664991856	TSw=	0.89464869531217905

FATw=	0.10559947087150801	T11/T22=	3.90621834E-02

Time (units of 2NT generations) =	7.50000030E-02

Alleles sampled from different populations

T11b=	8.1831394055967288E-002	T12b=	1.1249518202394944	T22b=	1.0449628249641314

TTb=	1.0497295298046152	TSb=	0.94864968187331500

FATb=	9.6291325585662113E-002	T11/T22=	7.83103406E-02

Alleles sampled from the same population

T11w=	5.3709272537144637E-002	T12w=	1.1252159087691318	T22w=	0.98750212897188350

TTw=	1.0029526807710409	TSw=	0.89412284332840963

FATw=	0.10850944369475746	T11/T22=	5.43890186E-02

Time (units of 2NT generations) =	0.100000001

Alleles sampled from different populations

T11b=	0.10132093562612965	T12b=	1.1490104681087905	T22b=	1.0426633078531093

TTb=	1.0523923729768623	TSb=	0.94852907063041147

FATb=	9.8692564687310136E-002	T11/T22=	9.71751213E-02

Alleles sampled from the same population

T11w=	6.6493276849236282E-002	T12w=	1.1492745274284513	T22w=	0.98532910267990503

TTw=	1.0056509208763367	TSw=	0.89344552009683820

FATw=	0.11157489984866842	T11/T22=	6.74833208E-02

Time (units of 2NT generations) =	0.125000000

Alleles sampled from different populations

T11b=	0.11790809410286873	T12b=	1.1729479569662500	T22b=	1.0404493048602468

TTb=	1.0550736501317537	TSb=	0.94819518378450895

FATb=	0.10129953139659786	T11/T22=	0.113324210

Alleles sampled from the same population

T11w=	7.7373560639361905E-002	T12w=	1.1732119920932362	T22w=	0.98323688700461576

TTw=	1.0083737726569151	TSw=	0.89265055436809038

FATw=	0.11476222550286208	T11/T22=	7.86926970E-02

Time (units of 2NT generations) =	0.150000006

Alleles sampled from different populations

T11b=	0.13205365639991251	T12b=	1.1967639422087446	T22b=	1.0383183967851957

TTb=	1.0577759475575816	TSb=	0.94769192274666736

FATb=	0.10407121192829139	T11/T22=	0.127180308

Alleles sampled from the same population

T11w=	8.6652361160363400E-002	T12w=	1.1970279573348943	T22w=	0.98122319580599004

TTw=	1.0111223445347366	TSw=	0.89176611234142733

FATw=	0.11804331378734445	T11/T22=	8.83105546E-02

Time (units of 2NT generations) =	0.174999997

Alleles sampled from different populations

T11b=	0.14414509279300491	T12b=	1.2204582809069704	T22b=	1.0362682292421215

TTb=	1.0605012071773032	TSb=	0.94705591559720981

FATb=	0.10697327906117760	T11/T22=	0.139100179

Alleles sampled from the same population

T11w=	9.4583826153097905E-002	T12w=	1.2207222795307473	T22w=	0.97928580419370781

TTw=	1.0138973499739690	TSw=	0.89081560638964685

FATw=	0.12139467924191949	T11/T22=	9.65844989E-02

Time (units of 2NT generations) =	0.200000003

Alleles sampled from different populations

T11b=	0.15450822366840700	T12b=	1.2440309975679549	T22b=	1.0342965110984244

TTb=	1.0632508357886399	TSb=	0.94631768235542268

FATb=	0.10997701529807402	T11/T22=	0.149384841

Alleles sampled from the same population

T11w=	0.10138166624641232	T12w=	1.2442949826057017	T22w=	0.97742254705172904

TTw=	1.0166991766433910	TSw=	0.88981845897119749

FATw=	0.12479671527923075	T11/T22=	0.103723481

Time (units of 2NT generations) =	0.300000012

Alleles sampled from different populations

T11b=	0.18354396307121601	T12b=	1.3371105084476804	T22b=	1.0271504373820397

TTb=	1.0745071854307469	TSb=	0.94278978995095741

FATb=	0.12258400619907139	T11/T22=	0.178692386

Alleles sampled from the same population

T11w=	0.12042864015872651	T12w=	1.3373744596334927	T22w=	0.97066957136775711

TTw=	1.0281740419434993	TSw=	0.88564547824685402

FATw=	0.13862299365896413	T11/T22=	0.124067597

Time (units of 2NT generations) =	0.400000006

Alleles sampled from different populations

T11b=	0.20045399473187009	T12b=	1.4282735112757523	T22b=	1.0210916712834137

TTb=	1.0861780257165194	TSb=	0.93902790362825939

FATb=	0.13547514183155152	T11/T22=	0.196313411

Alleles sampled from the same population

T11w=	0.13152213406493557	T12w=	1.4285374476777815	T22w=	0.96494409831682160

TTw=	1.0400566815592756	TSw=	0.88160190189163301

FATw=	0.15235206164925907	T11/T22=	0.136300266

Time (units of 2NT generations) =	0.500000000

Alleles sampled from different populations

T11b=	0.21125199748814771	T12b=	1.5175589124637296	T22b=	1.0160007033715208

TTb=	1.0982336939492847	TSb=	0.93552583278318358

FATb=	0.14815413337119376	T11/T22=	0.207925051

Alleles sampled from the same population

T11w=	0.13860661371155519	T12w=	1.5178228426047138	T22w=	0.96013319172994960

TTw=	1.0523020631072233	TSw=	0.87798053392811015

FATw=	0.16565731009248319	T11/T22=	0.144361854

Time (units of 2NT generations) =	1.00000000

Alleles sampled from different populations

T11b=	0.23957773776084629	T12b=	1.9374073079670113	T22b=	1.0016871093641289

TTb=	1.1624956513966149	TSb=	0.92547617220380063

FATb=	0.20388848672944337	T11/T22=	0.239174232

Alleles sampled from the same population

T11w=	0.15719485178812359	T12w=	1.9376712242319751	T22w=	0.94660710064699960

TTw=	1.1171045204037064	TSw=	0.86766587576111209

FATw=	0.22329033683656618	T11/T22=	0.166061342

Time (units of 2NT generations) =	1.50000000

Alleles sampled from different populations

T11b=	0.26008355029049385	T12b=	2.3168754098339157	T22b=	0.99951774776806368

TTb=	1.2292477849651415	TSb=	0.92557432802030670

FATb=	0.24704006845409632	T11/T22=	0.260209024

Alleles sampled from the same population

T11w=	0.17065301783823927	T12w=	2.3171392948359184	T22w=	0.94455726461575307

TTw=	1.1838829875876078	TSw=	0.86716683993800170

FATw=	0.26752318512066542	T11/T22=	0.180669844

Time (units of 2NT generations) =	2.00000000

Alleles sampled from different populations

T11b=	0.27848545533982638	T12b=	2.6605415644119130	T22b=	1.0038059248915985

TTb=	1.2947651353097376	TSb=	0.93127387793642125

FATb=	0.28073914523992871	T11/T22=	0.277429581

Alleles sampled from the same population

T11w=	0.18273043548912446	T12w=	2.6608054051927867	T22w=	0.94860977116758594

TTw=	1.2491461919353375	TSw=	0.87202183759973984

FATw=	0.30190569908499520	T11/T22=	0.192629725

Time (units of 2NT generations) =	2.50000000

Alleles sampled from different populations

T11b=	0.29516222184379493	T12b=	2.9721942242999573	T22b=	1.0113184217935225

TTb=	1.3571145042451835	TSb=	0.93970280179854981

FATb=	0.30757294328586871	T11/T22=	0.291858852

Alleles sampled from the same population

T11w=	0.19367562587669515	T12w=	2.9724580156026139	T22w=	0.95570923647972339

TTw=	1.3111036806158136	TSw=	0.87950587541942049

FATw=	0.32918663228347855	T11/T22=	0.202651203

Time (units of 2NT generations) =	3.00000000

Alleles sampled from different populations

T11b=	0.31029463010489505	T12b=	3.2550537354883620	T22b=	1.0202341977960945

TTb=	1.4154023189037908	TSb=	0.94924024102697457

FATb=	0.32934952250032501	T11/T22=	0.304140598

Alleles sampled from the same population

T11w=	0.20360723422434598	T12w=	3.2553174764522543	T22w=	0.96413478225241689

TTw=	1.3689423917281069	TSw=	0.88808202744960973

FATw=	0.35126413440340265	T11/T22=	0.211181298

Time (units of 2NT generations) =	4.00000000

Alleles sampled from different populations

T11b=	0.33651447494142350	T12b=	3.7452561041275851	T22b=	1.0387011122194600

TTb=	1.5188591443901422	TSb=	0.96848244849165643

FATb=	0.36236190691630921	T11/T22=	0.323976249

Alleles sampled from the same population

T11w=	0.22081568217043684	T12w=	3.7455197500428197	T22w=	0.98158624897813351

TTw=	1.4714865735017002	TSw=	0.90550919229736382

FATw=	0.38462966050548364	T11/T22=	0.224958003

Time (units of 2NT generations) =	5.00000000

Alleles sampled from different populations

T11b=	0.35815693729852965	T12b=	4.1499311136255574	T22b=	1.0555901447046452

TTb=	1.6055971870363481	TSb=	0.98584682396403367

FATb=	0.38599367766473569	T11/T22=	0.339295447

Alleles sampled from the same population

T11w=	0.23501993156700443	T12w=	4.1501946768174482	T22w=	0.99754657363251187

TTw=	1.5573979657851456	TSw=	0.92129390942596112

FATw=	0.40844027688099582	T11/T22=	0.235597953

**Table S5**

**Coalescence times for inversion and standard arrangements in a subdivided population**

**Time course of approach to equilibrium**

**Times scaled by 2x total population size**

**Inversion frequency= 0.5**

**Local N=	500**

**Total N=	10000**

**Number of populations =	200**

**Neutral Fst=	0.05**

**Scaled migration rate=	19	Migration rate=	0.0095**

Initial state of subdivided population

Alleles sampled from different populations

T11b=	0	T12b=	1.0524 T22b=	1.0524

TTb=	0.7893	TSb=	0.5263

FATb=	0.3333	T11/T22=	0

Alleles sampled from the same population

T11w=	0	T12w=	1	T22w=	1

TTw=	0.75	TSw=	0.5

FATw=	0.3333	T11/T22=	0

**Scaled recombination rate=	40**

Equilibrium results for subdivided population

Alleles sampled from different populations

T11b=	1.1052631578947374	T12b=	1.1552631578947374	T22b=	1.1052631578947374

TTb=	1.1302631578947373	TSb=	1.1052631578947374

FATb=	2.2118742724097751E-002	T11/T22=	1.00000000

Alleles sampled from the same population

T11w=	1.0004761904761910	T12w=	1.1555263157894742	T22w=	1.0004761904761910

TTw=	1.0780012531328327	TSw=	1.0004761904761910

FATw=	7.1915558939604440E-002 T11/T22=	1.00000000

Time (units of 2NT generations) =	2.49999994E-03

Alleles sampled from different populations

T11b=	5.2083751635259024E-002	T12b=	1.0299456493807646	T22b=	1.0498671724787803

TTb=	0.79046055571889218	TSb=	0.55097546205701964

FATb=	0.30296906269291379	T11/T22=	4.96098474E-02

Alleles sampled from the same population

T11w=	4.7134560465934561E-002	T12w=	1.0304821410766314	T22w=	0.95028085894582726

TTw=	0.76459492539125617	TSw=	0.49870770970588091

FATw=	0.34774912421674298	T11/T22=	4.96006645E-02

Time (units of 2NT generations) =	4.99999989E-03

Alleles sampled from different populations

T11b=	0.10140042620929490	T12b=	1.0095864870186666	T22b=	1.0462054464220933

TTb=	0.79169471166718042	TSb=	0.57380293631569412

FATb=	0.27522196642269037	T11/T22=	9.69220996E-02

Alleles sampled from the same population

T11w=	9.1764451539401160E-002	T12w=	1.0101130517181780	T22w=	0.94695682533440628

TTw=	0.76473684507754092	TSw=	0.51936063843690372

FATw=	0.32086358623894617	T11/T22=	9.69045758E-02

Time (units of 2NT generations) =	7.49999983E-03

Alleles sampled from different populations

T11b=	0.14715725029783910	T12b=	0.99128126930575478	T22b=	1.0417968142776135

TTb=	0.79287915079674054	TSb=	0.59447703228772630

FATb=	0.25022996040398571	T11/T22=	0.141253307

Alleles sampled from the same population

T11w=	0.13317301559050976	T12w=	0.99179746847834616	T22w=	0.94295801465413320

TTw=	0.76493149180033382	TSw=	0.53806551512232148

FATw=	0.29658339225132857	T11/T22=	0.141229004

Time (units of 2NT generations) =	9.99999978E-03

Alleles sampled from different populations

T11b=	0.18963837799664263	T12b=	0.97482988146787342	T22b=	1.0367760755267068

TTb=	0.79401855411477407	TSb=	0.61320722676167472

FATb=	0.22771675348906695	T11/T22=	0.182911605

Alleles sampled from the same population

T11w=	0.17161741841353551	T12w=	0.97533669036768877	T22w=	0.93840599698868987

TTw=	0.76517419903440065	TSw=	0.55501170770111274

FATw=	0.27465966782269857	T11/T22=	0.182881847

Time (units of 2NT generations)=	1.25000002E-02

Alleles sampled from different populations

T11b=	0.22910360664525675	T12b=	0.96005169683633140	T22b=	1.0312616012952232

TTb=	0.79511715040328568	TSb=	0.63018260397023995

FATb=	0.20743427097427147	T11/T22=	0.222158581

Alleles sampled from the same population

T11w=	0.20733273404807906	T12w=	0.96054999588477374	T22w=	0.93340783412177486

TTw=	0.76546013998485041	TSw=	0.57037028408492696

FATw=	0.25486612000957298	T11/T22=	0.222124487

Time (units of 2NT generations)=	1.49999997E-02

Alleles sampled from different populations

T11b=	0.26579057223879654	T12b=	0.94678367285228759	T22b=	1.0253571105441193

TTb=	0.79617875712187280	TSb=	0.64557384139145790

FATb=	0.18915967599391936	T11/T22=	0.259217560

Alleles sampled from the same population

T11w=	0.24053398513454671	T12w=	0.94727425683748967	T22w=	0.92805733339130059

TTw=	0.76578495805020663	TSw=	0.58429565926292359

FATw=	0.23699773269166802	T11/T22=	0.259180099

Time (units of 2NT generations) =	1.75000001E-02

Alleles sampled from different populations

T11b=	0.29991674065355078	T12b=	0.93487863927684900	T22b=	1.0191532625869797

TTb=	0.79720682044855717	TSb=	0.65953500162026524

FATb=	0.17269272577325590	T11/T22=	0.294280320

Alleles sampled from the same population

T11w=	0.27141794450004308	T12w=	0.93536222568978833	T22w=	0.92243648838097914

TTw=	0.76614472106514964	TSw=	0.59692721644051105

FATw=	0.22086885150024949	T11/T22=	0.294240266

Time (units of 2NT generations) =	1.99999996E-02

Alleles sampled from different populations

T11b=	0.33168121385146909	T12b=	0.92420375304173008	T22b=	1.0127290857562110

TTb=	0.79820445142278507	TSb=	0.67220514980384005

FATb=	0.15785341887576987	T11/T22=	0.327512294

Alleles sampled from the same population

T11w=	0.30016476934603409	T12w=	0.92468098960083722	T22w=	0.91661677137836739

TTw=	0.76653587998151895	TSw=	0.60839077036220068

FATw=	0.20631142487828635	T11/T22=	0.327470303

Time (units of 2NT generations)=	2.50000004E-02

Alleles sampled from different populations

T11b=	0.38883934361364947	T12b=	0.90607645356045308	T22b=	0.99948525866518301

TTb=	0.80011937734993466	TSb=	0.69416230113941624

FATb=	0.13242658434477306	T11/T22=	0.389039606

Alleles sampled from the same population

T11w=	0.35189332495910053	T12w=	0.90654268777629077	T22w=	0.90462083853840403

TTw=	0.76739988476252152	TSw=	0.62825708174875228

FATw=	0.18131720603115309	T11/T22=	0.388995379

Time (units of 2NT generations)=	2.74999999E-02

Alleles sampled from different populations

T11b=	0.41455338931886132	T12b=	0.89841810310850301	T22b=	0.99277639142006358

TTb=	0.80104149673898273	TSb=	0.70366489036946245

FATb=	0.12156249927867369	T11/T22=	0.417569757

Alleles sampled from the same population

T11w=	0.37516496306349789	T12w=	0.89887957666985896	T22w=	0.89854480458554353

TTw=	0.76786723024718984	TSw=	0.63685488382452071

FATw=	0.17061848879850483	T11/T22=	0.417525053

Time (units of 2NT generations)=	2.99999993E-02

Alleles sampled from different populations

T11b=	0.43854909343857607	T12b=	0.89157586341537520	T22b=	0.98607071262554291

TTb=	0.80194288322371732	TSb=	0.71230990303205943

FATb=	0.11176978069977217	T11/T22=	0.444744051

Alleles sampled from the same population

T11w=	0.39688161142349410	T12w=	0.89203300657502838	T22w=	0.89247202764463229

TTw=	0.76835491305454573	TSw=	0.64467681953406319

FATw=	0.16096479819307485	T11/T22=	0.444699228

Time (units of 2NT generations)=	3.24999988E-02

Alleles sampled from different populations

T11b=	0.46095549389125301	T12b=	0.88547013113366146	T22b=	0.97940585367935684

TTb=	0.80282540245948320	TSb=	0.72018067378530493

FATb=	0.10294234390316159	T11/T22=	0.470648080

Alleles sampled from the same population

T11w=	0.41716003281531910	T12w=	0.88592333233327014	T22w=	0.88643653367554298

TTw=	0.76886080778935062	TSw=	0.65179828324543099

FATw=	0.15225450869384405	T11/T22=	0.470603377

Time (units of 2NT generations)=	3.50000001E-02

Alleles sampled from different populations

T11b=	0.48189108974619921	T12b=	0.88002905253776009	T22b=	0.97281376045927215

TTb=	0.80369073882024789	TSb=	0.72735242510273568

FATb=	9.4984687554780844E-002 T11/T22=	0.495358020

Alleles sampled from the same population

T11w=	0.43610745427493547	T12w=	0.88047866247795947	T22w=	0.88046720589965233

TTw=	0.76938299628262663	TSw=	0.65828733008729390

FATw=	0.14439579082473331	T11/T22=	0.495313674

Time (units of 2NT generations)=	3.75000015E-02

Alleles sampled from different populations

T11b=	0.50146475977923988	T12b=	0.87518776910807994	T22b=	0.96632135418599308

TTb=	0.80454041304534818	TSb=	0.73389305698261653

FATb=	8.7810823318765441E-002 T11/T22=	0.518941998

Alleles sampled from the same population

T11w=	0.45382239818786640	T12w=	0.87563410442235001	T22w=	0.87458838263845284

TTw=	0.76991974741775482	TSw=	0.66420539041315962

FATw=	0.13730568329900894	T11/T22=	0.518898249

Time (units of 2NT generations) =	3.99999991E-02

Alleles sampled from different populations

T11b=	0.51977659731008319	T12b=	0.87088773655656027	T22b=	0.95995112225016888

TTb=	0.80537579816834315	TSb=	0.73986385978012603

FATb=	8.1343316420992684E-002	T11/T22=	0.541461527

Alleles sampled from the same population

T11w=	0.47039543763455760	T12w=	0.87133108312746033	T22w=	0.86882039179240467

TTw=	0.77046949892047067	TSw=	0.66960791471348113

FATw=	0.13090924994215858	T11/T22=	0.541418493

Time (units of 2NT generations) =	4.25000004E-02

Alleles sampled from different populations

T11b=	0.53691866916846798	T12b=	0.86707611014413755	T22b=	0.95372164612418819

TTb=	0.80619813389523276	TSb=	0.74532015764632809

FATb=	7.5512425158771102E-002	T11/T22=	0.562972069

Alleles sampled from the same population

T11w=	0.48590988309086258	T12w=	0.86751672609670172	T22w=	0.86318002840206876

TTw=	0.77103084092158380	TSw=	0.67454495574646567

FATw=	0.12513881423963824	T11/T22=	0.562929928

Time (units of 2NT generations) =	4.50000018E-02

Alleles sampled from different populations

T11b=	0.55297570588835443	T12b=	0.86370518983671696	T22b=	0.94764807277009899

TTb=	0.80700853958297181	TSb=	0.75031188932922666

FATb=	7.0255328751593682E-002	T11/T22=	0.583524346

Alleles sampled from the same population

T11w=	0.50044240690666930	T12w=	0.86414330824045449	T22w=	0.85768098109183333

TTw=	0.77160250111985296	TSw=	0.67906169399925131

FATw=	0.11993326484335398	T11/T22=	0.583483160

Time (units of 2NT generations) =	4.74999994E-02

Alleles sampled from different populations

T11b=	0.56802572955357955	T12b=	0.86073191947443994	T22b=	0.94174253531570751

TTb=	0.80780802595454171	TSb=	0.75488413243464358

FATb=	6.5515434137165096E-002	T11/T22=	0.603164554

Alleles sampled from the same population

T11w=	0.51406361137412648	T12w=	0.86116775078228924	T22w=	0.85233421261816811

TTw=	0.77218333138921835	TSw=	0.68319891199614724

FATw=	0.11523742584935259	T11/T22=	0.603124440

Time (units of 2NT generations) =	5.00000007E-02

Alleles sampled from different populations

T11b=	0.58214062510749010	T12b=	0.85811743469614998	T22b=	0.93601452819453790

TTb=	0.80859750567358202	TSb=	0.75907757665101405

FATb=	6.1241753375576691E-002	T11/T22=	0.621935487

Alleles sampled from the same population

T11w=	0.52683854564435750	T12w=	0.85855116894601979	T22w=	0.84714829922291091

TTw=	0.77277229568982708	TSw=	0.68699342243363426

FATw=	0.11100148612292182	T11/T22=	0.621896505

Time (units of 2NT generations) =	5.99999987E-02

Alleles sampled from different populations

T11b=	0.63049967872365331	T12b=	0.85059500715590475	T22b=	0.91499306075910258

TTb=	0.81167068844864132	TSb=	0.77274636974137789

FATb=	4.7955801855627089E-002	T11/T22=	0.689075887

Alleles sampled from the same population

T11w=	0.57060740565926249	T12w=	0.85102191087224521	T22w=	0.82811722568147683

TTw=	0.77519211327130744	TSw=	0.69936231567036966

FATw=	9.7820651555569071E-002	T11/T22=	0.689041853

Time (units of 2NT generations) =	7.00000003E-02

Alleles sampled from different populations

T11b=	0.66836822165132370	T12b=	0.84654804035709896	T22b=	0.89708382830079714

TTb=	0.81463703266657972	TSb=	0.78272602497606036

FATb=	3.9172056278934386E-002	T11/T22=	0.745045424

Alleles sampled from the same population

T11w=	0.60488233650300816	T12w=	0.84696996423376270	T22w=	0.81190477858300869

TTw=	0.77768176088838548	TSw=	0.70839355754300848

FATw=	8.9095831778574275E-002	T11/T22=	0.745016336

Time (units of 2NT generations) =	7.99999982E-02

Alleles sampled from different populations

T11b=	0.69831179461861681	T12b=	0.84479996409521241	T22b=	0.88218541709366360

TTb=	0.81752428497567631	TSb=	0.79024860585614021

FATb=	3.3363753983586708E-002	T11/T22=	0.791570306

Alleles sampled from the same population

T11w=	0.63198500828123338	T12w=	0.84521814067005230	T22w=	0.79841862825374244

TTw=	0.78020997946877002	TSw=	0.71520181826748797

FATw=	8.3321365929649893E-002	T 11/T22=	0.791545928

Time (units of 2NT generations) =	9.00000036E-02

Alleles sampled from different populations

T11b=	0.72222042828719690	T12b=	0.84456976744984302	T22b=	0.87004378646281588

TTb=	0.82035093741242471	TSb=	0.79613210737500639

FATb=	2.9522523755272423E-002	T11/T22=	0.830096662

Alleles sampled from the same population

T11w=	0.65362583145642972	T12w=	0.84498501885358202	T22w=	0.78742847419818540

TTw=	0.78275608584044476	TSw=	0.72052715282730762

FATw=	7.9499775394683780E-002	T11/T22=	0.830076456

Time (units of 2NT generations) =	0.125000000

Alleles sampled from different populations

T11b=	0.77573769028189077	T12b=	0.84965176325765934	T22b=	0.84460700730281646

TTb=	0.82991205602500651	TSb=	0.81017234879235356

FATb=	2.3785300007809651E-002	T11/T22=	0.918459952

Alleles sampled from the same population

T11w=	0.70206932373204189	T12w=	0.85006006203488382	T22w=	0.76440653981850115

TTw=	0.79164899690507773	TSw=	0.73323793177527152

FATw=	7.3784044896364587E-002	T11/T22=	0.918450177

Time (units of 2NT generations) =	0.150000006

Alleles sampled from different populations

T11b=	0.79760803920473855	T12b=	0.85545518701349788	T22b=	0.83751832668300197

TTb=	0.83650918497868409	TSb=	0.81756318294387031

FATb=	2.2648887035587761E-002	T11/T22=	0.952346981

Alleles sampled from the same population

T11w=	0.72186789637115056	T12w=	0.85585984141753524	T22w=	0.75799278220164623

TTw=	0.79789509035196682	TSw=	0.73993033928639840

FATw=	7.2647083265043100E-002	T11/T22=	0.952341378

Time (units of 2NT generations) =	0.174999997

Alleles sampled from different populations

T11b=	0.81263220654527069	T12b=	0.86170074113403039	T22b=	0.83576051847717747

TTb=	0.84294855182262718	TSb=	0.82419636251122408

FATb=	2.2245947597699822E-002	T11/T22=	0.972326636

Alleles sampled from the same population

T11w=	0.73546983475853367	T12w=	0.86210212990285939	T22w=	0.75640447794914678

TTw=	0.80401964312834984	TSw=	0.74593715635384017

FATw=	7.2240134020255087E-002	T11/T22=	0.972323477

Time (units of 2NT generations) =	0.200000003

Alleles sampled from different populations

T11b=	0.82376883497614184	T12b=	0.86801393789540426	T22b=	0.83717186578896241

TTb=	0.84924214413897825	TSb=	0.83047035038255212

FATb=	2.2104171214275214E-002	T11/T22=	0.983990133

Alleles sampled from the same population

T11w=	0.74555299066319913	T12w=	0.86841223987845018	T22w=	0.75768477393234912

TTw=	0.81001556108811212	TSw=	0.75161888229777407

FATw=	7.2093280173398777E-002	T11/T22=	0.983988345

Time (units of 2NT generations) =	0.250000000

Alleles sampled from different populations

T11b=	0.84017909075097374	T12b=	0.88040020298959842	T22b=	0.84468021680945116

TTb=	0.86141492838490541	TSb=	0.84242965378021251

FATb=	2.2039639642987163E-002	T11/T22=	0.994671226

Alleles sampled from the same population

T11w=	0.76041228856877052	T12w=	0.88079262277321735	T22w=	0.76448649287160131

TTw=	0.82162100674670158	TSw=	0.76244939072018592

FATw=	7.2018139191465158E-002	T11/T22=	0.994670689

Time (units of 2NT generations) =	0.300000012

Alleles sampled from different populations

T11b=	0.85306503752469620	T12b=	0.89229789475776300	T22b=	0.85457664606138661

TTb=	0.87305936827540220	TSb=	0.85382084179304141

FATb=	2.2035759744911343E-002	T11/T22=	0.998231173

Alleles sampled from the same population

T11w=	0.77208133307913562	T12w=	0.89268471226535440	T22w=	0.77344956892153749

TTw=	0.83272508163284553	TSw=	0.77276545100033656

FATw=	7.2004112707806711E-002	T11/T22=	0.998230994

Time (units of 2NT generations) =	0.400000006

Alleles sampled from different populations

T11b=	0.87504574748456021	T12b=	0.91458255031192248	T22b=	0.87521622835565283

TTb=	0.89485676911601453	TSb=	0.87513098792010657

FATb=	2.2043506711575844E-002	T11/T22=	0.999805212

Alleles sampled from the same population

T11w=	0.79198715082130255	T12w=	0.91495888745545528	T22w=	0.79214146196245516

TTw=	0.85351159692366707	TSw=	0.79206430639187886

FATw=	7.1993503958545091E-002	T11/T22=	0.999805212

Time (units of 2NT generations) =	0.500000000

Alleles sampled from different populations

T11b=	0.89462451727924086	T12b=	0.93497961309833078	T22b=	0.89464374429913796

TTb=	0.91480687194376009	TSb=	0.89463413078918941

FATb=	2.2051366002212136E-002	T11/T22=	0.999978483

Alleles sampled from the same population

T11w=	0.80971800604186162	T12w=	0.93534635857504567	T22w=	0.80973540942185551

TTw=	0.87253653315345203	TSw=	0.80972670773185862

FATw=	7.1985324436320330E-002	T11/T22=	0.999978483

Time (units of 2NT generations) =	0.750000000

Alleles sampled from different populations

T11b=	0.93646385887186046	T12b=	0.97872671597655581	T22b=	0.93646394100204344

TTb=	0.95759530795675385	TSb=	0.93646389993695189

FATb=	2.2067159105959466E-002	T11/T22=	0.999999940

Alleles sampled from the same population

T11w=	0.84760847127002992	T12w=	0.97907288952782967	T22w=	0.84760854561034871

TTw=	0.91334069898400949	TSw=	0.84760850844018931

FATw=	7.1968971290713335E-002	T11/T22=	0.999999940

Time (units of 2NT generations) =	1.00000000

Alleles sampled from different populations

T11b=	0.96998650605415071	T12b=	1.0137858845435095	T22b=	0.96998650640497985

TTb=	0.99188619538653733	TSb=	0.96998650622956528

FATb=	2.2078832489888400E-002	T11/T22=	1.00000000

Alleles sampled from the same population

T11w=	0.87796719102537768	T12w=	1.0141155716409691	T22w=	0.87796719134293200

TTw=	0.94604138141256200	TSw=	0.87796719118415489

FATw=	7.1956884303267521E-002	T11/T22=	1.00000000

Time (units of 2NT generations) =	1.50000000

Alleles sampled from different populations

T11b=	1.0183816261088150	T12b=	1.0643992721439750	T22b=	1.0183816261088197

TTb=	1.0413904491263961	TSb=	1.0183816261088174

FATb=	2.2094328824390885E-002	T11/T22=	1.00000000

Alleles sampled from the same population

T11w=	0.92179469307782758	T12w=	1.0647051584694491	T22w=	0.92179469307783191

TTw=	0.99324992577363935	TSw=	0.92179469307782980

FATw=	7.1940838696920362E-002	T11/T22=	1.00000000

Time (units of 2NT generations) =	2.00000000

Alleles sampled from different populations

T11b=	1.0494634284325481	T12b=	1.0969057585884152	T22b=	1.0494634284325481

TTb=	1.0731845935104818	TSb=	1.0494634284325481

FATb=	2.2103527409333745E-002	T11/T22=	1.00000000

Alleles sampled from the same population

T11w=	0.94994293929771645	T12w=	1.0971963588500910	T22w=	0.94994293929771656

TTw=	1.0235696490739037	TSw=	0.94994293929771656

FATw=	7.1931313948984776E-002	T11/T22=	1.00000000

Time (units of 2NT generations) =	3.00000000

Alleles sampled from different populations

T11b=	1.0822465488219428	T12b=	1.1311915457330914	T22b=	1.0822465488219428

TTb=	1.1067190472775172	TSb=	1.0822465488219428

FATb=	2.2112656790154395E-002	T11/T22=	1.00000000

Alleles sampled from the same population

T11w=	0.97963193014083538	T12w=	1.1314660232209151	T22w=	0.97963193014083538

TTw=	1.0555489766808752	TSw=	0.97963193014083538

FATw=	7.1921860773109247E-002	T11/T22=	1.00000000

Time (units of 2NT generations) =	4.00000000

Alleles sampled from different populations

T11b=	1.0957691281150981	T12b=	1.1453339540757126	T22b=	1.0957691281150981

TTb=	1.1205515410954052	TSb=	1.0957691281150981

FATb=	2.2116263350172027E-002	T11/T22=	1.00000000

Alleles sampled from the same population

T11w=	0.99187822421050276	T12w=	1.1456017811445021	T22w=	0.99187822421050276

TTw=	1.0687400026775025	TSw=	0.99187822421050276

FATw=	7.1918126274340577E-002	T11/T22=	1.00000000

Time (units of 2NT generations) =	5.00000000

Alleles sampled from different populations

T11b=	1.1013470032095882	T12b=	1.1511674999962267	T22b=	1.1013470032095880

TTb=	1.1262572516029075	TSb=	1.1013470032095882

FATb=	2.2117725198098848E-002	T11/T22=	1.00000000

Alleles sampled from the same population

T11w=	0.99692964947562024	T12w=	1.1514325838600479	T22w=	0.99692964947562024

TTw=	1.0741811166678341	TSw=	0.99692964947562024

FATw=	7.1916612565162130E-002	T11/T22=	1.00000000

**Scaled recombination rate=	4**

Equilibrium results for subdivided population

Alleles sampled from different populations

T11b=	1.1052631578947369	T12b=	1.6052631578947369	T22b=	1.1052631578947369

TTb=	1.3552631578947369	TSb=	1.1052631578947369

FATb=	0.18446601941747576	T11/T22=	1.00000000

Alleles sampled from the same population

T11w=	1.0004761904761905	T12w=	1.6055263157894737	T22w=	1.0004761904761905

TTw=	1.3030012531328321	TSw=	1.0004761904761905

FATw=	0.23217557306969161	T11/T22=	1.00000000

Time (units of 2NT generations) =	2.49999994E-03

Alleles sampled from different populations

T11b=	7.5735331463877564E-003	T12b=	1.0519867248969148	T22b=	1.0503733529514205

TTb=	0.79048008397290948	TSb=	0.52897344304890415

FATb=	0.33082002472432614	T11/T22=	7.21032498E-03

Alleles sampled from the same population

T11w=	6.8751087957386048E-003	T12w=	1.0522768759780301	T22w=	0.95076812472941996

TTw=	0.76554924637030464	TSw=	0.47882161676257928

FATw=	0.37453845192479229	T11/T22=	7.23110978E-03

Time (units of 2NT generations) =	4.99999989E-03

Alleles sampled from different populations

T11b=	1.5228585495341562E-002	T12b=	1.0518799318867653	T22b=	1.0481389514391710

TTb=	0.79178185017701086	TSb=	0.53168376846725629

FATb=	0.32849715063764973	T11/T22=	1.45291667E-02

Alleles sampled from the same population

T11w=	1.3804097506943114E-002	T12w=	1.0521715656921351	T22w=	0.94874522172968123

TTw=	0.76672311265522364	TSw=	0.48127465961831217

FATw=	0.37229665876170159	T11/T22=	1.45498468E-02

Time (units of 2NT generations) =	7.49999983E-03

Alleles sampled from different populations

T11b=	2.2810543286337914E-002	T12b=	1.0517871274496247	T22b=	1.0459252406773387

TTb=	0.79307750971573143	TSb=	0.53436789198183832

FATb=	0.32620975196563606	T11/T22=	2.18089614E-02

Alleles sampled from the same population

T11w=	2.0666919237205539E-002	T12w=	1.0520786279060736	T22w=	0.94674147219257332

TTw=	0.76789141181048159	TSw=	0.48370419571488943

FATw=	0.37008776465614468	T11/T22=	2.18295269E-02

Time (units of 2NT generations) =	9.99999978E-03

Alleles sampled from different populations

T11b=	3.0320168808246617E-002	T12b=	1.0517081039232865	T22b=	1.0437320954674920

TTb=	0.79436711803057791	TSb=	0.53702613213786932

FATb=	0.32395724854613461	T11/T22=	2.90497616E-02

Alleles sampled from the same population

T11w=	2.7464269873437308E-002	T12w=	1.0519994722775141	T22w=	0.94475633823469862

TTw=	0.76905488816579104	TSw=	0.48611030405406797

FATw=	0.36791208074439352	T11/T22=	2.90702153E-02

Time (units of 2NT generations) =	1.25000002E-02

Alleles sampled from different populations

T11b=	3.7758216159404778E-002	T12b=	1.0516426643009911	T22b=	1.0415593889058608

TTb=	0.79565073341681192	TSb=	0.53965880253263276

FATb=	0.32173907486373732	T11/T22=	3.62516195E-02

Alleles sampled from the same population

T11w=	3.4196831980739528E-002	T12w=	1.0519339018243292	T22w=	0.94278970497705505

TTw=	0.77021358515161320	TSw=	0.48849326847889729

FATw=	0.36576908289310484	T11/T22=	3.62719633E-02

Time (units of 2NT generations) =	1.49999997E-02

Alleles sampled from different populations

T11b=	4.5125431312036390E-002	T12b=	1.0515906141125151	T22b=	1.0394069943175013

TTb=	0.79692841346364196	TSb=	0.54226621281476883

FATb=	0.31955467561013440	T11/T22=	4.34145927E-02

Alleles sampled from the same population

T11w=	4.0865280768770385E-002	T12w=	1.0518817220608991	T22w=	0.94084145775077366

TTw=	0.77136754566033561	TSw=	0.49085336925977202

FATw=	0.36365825601390467	T11/T22=	4.34348211E-02

Time (units of 2NT generations) =	1.75000001E-02

Alleles sampled from different populations

T11b=	5.2422552201683015E-002	T12b=	1.0515517613917700	T22b=	1.0372747852666919

TTb=	0.79820021506297878	TSb=	0.54484866873418747

FATb=	0.31740350547111995	T11/T22=	5.05387336E-02

Alleles sampled from the same population

T11w=	4.7470284172901334E-002	T12w=	1.0518427410059374	T22w=	0.93891148210362962

TTw=	0.77251681207210143	TSw=	0.49319088313826548

FATw=	0.36157909390296294	T11/T22=	5.05588502E-02

Time (units of 2NT generations) =	1.99999996E-02

Alleles sampled from different populations

T11b=	5.9650308816008302E-002	T12b=	1.0515259166450239	T22b=	1.0351626355670520

TTb=	0.79946619441827704	TSb=	0.54740647219153016

FATb=	0.31528502891877175	T11/T22=	5.76240942E-02

Alleles sampled from the same population

T11w=	5.4012502934535078E-002	T12w=	1.0518167691506954	T22w=	0.93699966380920152

TTw=	0.77366142626128187	TSw=	0.49550608337186830

FATw=	0.35953109906693814	T11/T22=	5.76440990E-02

Time (units of 2NT generations) =	2.50000004E-02

Alleles sampled from different populations

T11b=	7.3900609953931884E-002	T12b=	1.0515125052722250	T22b=	1.0309980107814232

TTb=	0.80198090781995124	TSb=	0.55244931036767753

FATb=	0.31114406218295509	T11/T22=	7.16787130E-02

Alleles sampled from the same population

T11w=	6.6911194000691676E-002	T12w=	1.0518031071789649	T22w=	0.93323004355518369

TTw=	0.77593686297845132	TSw=	0.50007061877793768

FATw=	0.35552666378240461	T11/T22=	7.16985017E-02

Time (units of 2NT generations) =	2.99999993E-02

Alleles sampled from different populations

T11b=	8.7882018971749964E-002	T12b=	1.0515489123075537	T22b=	1.0269121158251859

TTb=	0.80447298985301074	TSb=	0.55739706739846795

FATb=	0.30712767932666984	T11/T22=	8.55789110E-02

Alleles sampled from the same population

T11w=	7.9566499009029457E-002	T12w=	1.0518392683427837	T22w=	0.92953168802717223

TTw=	0.77819418093044224	TSw=	0.50454909351810084

FATw=	0.35164113805780406	T11/T22=	8.55984762E-02

Time (units of 2NT generations) =	3.99999991E-02

Alleles sampled from different populations

T11b=	0.11506029198399270	T12b=	1.0517654978504229	T22b=	1.0189725221177615

TTb=	0.80939095245064996	TSb=	0.56701640705087708

FATb=	0.29945299569499562	T11/T22=	0.112917952

Alleles sampled from the same population

T11w=	0.10416698282521342	T12w=	1.0520553758739439	T22w=	0.92234515636396341

TTw=	0.78265572273426620	TSw=	0.51325606959458847

FATw=	0.34421220635621230	T11/T22=	0.112937093

Time (units of 2NT generations) =	5.00000007E-02

Alleles sampled from different populations

T11b=	0.14122821611901482	T12b=	1.0521646666395377	T22b=	1.0113359182291699

TTb=	0.81422336690681507	TSb=	0.57628206717409236

FATb=	0.29223098894428345	T11/T22=	0.139645204

Alleles sampled from the same population

T11w=	0.12785295532765195	T12w=	1.0524540842361221	T22w=	0.91543288484042273

TTw=	0.78704850216007971	TSw=	0.52164292008403734

FATw=	0.33721629778549644	T11/T22=	0.139663935

Time (units of 2NT generations) =	5.99999987E-02

Alleles sampled from different populations

T11b=	0.16642705764749022	T12b=	1.0527359692305047	T22b=	1.0039944541627310

TTb=	0.81897336256780773	TSb=	0.58521075590511062

FATb=	0.28543371170187781	T11/T22=	0.165764913

Alleles sampled from the same population

T11w=	0.15066176830262012	T12w=	1.0530249431541052	T22w=	0.90878776754375312

TTw=	0.79137485553864595	TSw=	0.52972476792318668

FATw=	0.33062724419942335	T11/T22=	0.165783226

Time (units of 2NT generations) =	7.00000003E-02

Alleles sampled from different populations

T11b=	0.19069633964784560	T12b=	1.0534694855866376	T22b=	0.99694037358068166

TTb=	0.82364392110045048	TSb=	0.59381835661426363

FATb=	0.27903510072547189	T11/T22=	0.191281587

Alleles sampled from the same population

T11w=	0.17262919570588786	T12w=	1.0537580318014164	T22w=	0.90240278336058732

TTw=	0.79563701066732695	TSw=	0.53751598953323754

FATw=	0.32442058083446224	T11/T22=	0.191299498

Time (units of 2NT generations) =	7.99999982E-02

Alleles sampled from different populations

T11b=	0.21407391733083481	T12b=	1.0543557986792071	T22b=	0.99016602043417401

TTb=	0.82823788378085572	TSb=	0.60211996888250441

FATb=	0.27301083339261989	T11/T22=	0.216200024

Alleles sampled from the same population

T11w=	0.19378950184174015	T12w=	1.0546439323984949	T22w=	0.89627100197695309

TTw=	0.79983709215392085	TSw=	0.54503025190934662

FATw=	0.31857342294340496	T11/T22=	0.216217533

Time (units of 2NT generations) =	9.00000036E-02

Alleles sampled from different populations

T11b=	0.23659605004186735	T12b=	1.0553859694012977	T22b=	0.98366384483999247

TTb=	0.83275795842111378	TSb=	0.61012994744092985

FATb=	0.26733819680604498	T11/T22=	0.240525305

Alleles sampled from the same population

T11w=	0.21417550653491046	T12w=	1.0556737051246459	T22w=	0.89038558919646904

TTw=	0.80397712649516784	TSw=	0.55228054786568981

FATw=	0.31306435262246335	T11/T22=	0.240542427

Time (units of 2NT generations) =	0.100000001

Alleles sampled from different populations

T11b=	0.25829747009029735	T12b=	1.0565515127297429	T22b=	0.97742640825735627

TTb=	0.83720672595178480	TSb=	0.61786193917382681

FATb=	0.26199596823424254	T11/T22=	0.264262825

Alleles sampled from the same population

T11w=	0.23381864743049152	T12w=	1.0568388642781115	T22w=	0.88473981162506299

TTw=	0.80805904690294428	TSw=	0.55927922952777731

FATw=	0.30787331486314984	T11/T22=	0.264279544

Time (units of 2NT generations) =	0.125000000

Alleles sampled from different populations

T11b=	0.30917531737170822	T12b=	1.0600057782125325	T22b=	0.96294329058788253

TTb=	0.84803254109616399	TSb=	0.63605930397979538

FATb=	0.24995884809133939	T11/T22=	0.321073234

Alleles sampled from the same population

T11w=	0.27987099906486179	T12w=	1.0602922256342775	T22w=	0.87163053988330930

TTw=	0.81802149755418152	TSw=	0.57575076947408554

FATw=	0.29616670075843476	T11/T22=	0.321089029

Time (units of 2NT generations) =	0.150000006

Alleles sampled from different populations

T11b=	0.35561067087572562	T12b=	1.0641418670700828	T22b=	0.94995827687335632

TTb=	0.85846317047231191	TSb=	0.65278447387454097

FATb=	0.23958942406883921	T11/T22=	0.374343455

Alleles sampled from the same population

T11w=	0.32190224827144398	T12w=	1.0644274835581160	T22w=	0.85987731101653320

TTw=	0.82765863160105235	TSw=	0.59088977964398859

FATw=	0.28607066116020519	T11/T22=	0.374358356

Time (units of 2NT generations) =	0.200000003

Alleles sampled from different populations

T11b=	0.43684521770118501	T12b=	1.0740682555313095	T22b=	0.92806321492917698

TTb=	0.87826123592324523	TSb=	0.68245421631518099

FATb=	0.22294849368164138	T11/T22=	0.470706314

Alleles sampled from the same population

T11w=	0.39543232488890911	T12w=	1.0743523981065564	T22w=	0.84005939196807011

TTw=	0.84604912826752299	TSw=	0.61774585842848961

FATw=	0.26984635077461283	T11/T22=	0.470719486

Time (units of 2NT generations) =	0.250000000

Alleles sampled from different populations

T11b=	0.50497119107722477	T12b=	1.0856636144945628	T22b=	0.91095435373449207

TTb=	0.89681319345021071	TSb=	0.70796277240585836

FATb=	0.21057944109609816	T11/T22=	0.554332018

Alleles sampled from the same population

T11w=	0.45709722223529203	T12w=	1.0859464900253426	T22w=	0.82457378847364327

TTw=	0.86339099768990513	TSw=	0.64083550535446765

FATw=	0.25776906746874650	T11/T22=	0.554343641

Time (units of 2NT generations) =	0.300000012

Alleles sampled from different populations

T11b=	0.56239012723222181	T12b=	1.0984159909317368	T22b=	0.89792817610523312

TTb=	0.91428757130023208	TSb=	0.73015915166872747

FATb=	0.20139004992668863	T11/T22=	0.626319706

Alleles sampled from the same population

T11w=	0.50907067312427889	T12w=	1.0986977640703923	T22w=	0.81278368731709261

TTw=	0.87981247214553904	TSw=	0.66092718022068575

FATw=	0.24878630259818046	T11/T22=	0.626329839

Time (units of 2NT generations) =	0.349999994

Alleles sampled from different populations

T11b=	0.61104608495399448	T12b=	1.1119334478700065	T22b=	0.88836246798968777

TTb=	0.93081886217092380	TSb=	0.74970427647184112

FATb=	0.19457554316924131	T11/T22=	0.687834203

Alleles sampled from the same population

T11w=	0.55311232904952379	T12w=	1.1122142504056374	T22w=	0.80412587240089284

TTw=	0.89541667556542281	TSw=	0.67861910072520826

FATw=	0.24211920634973083	T11/T22=	0.687842965

Time (units of 2NT generations) =	0.400000006

Alleles sampled from different populations

T11b=	0.65251462993885012	T12b=	1.1259168773039494	T22b=	0.88171181505184348

TTb=	0.94651504989964808	TSb=	0.76711322249534675

FATb=	0.18953932895554271	T11/T22=	0.740054309

Alleles sampled from the same population

T11w=	0.59064830334210172	T12w=	1.1261968154147053	T22w=	0.79810664724372138

TTw=	0.91028714535380839	TSw=	0.69437747529291149

FATw=	0.23718853019392860	T11/T22=	0.740061879

Time (units of 2NT generations) =	0.449999988

Alleles sampled from different populations

T11b=	0.68807420165026389	T12b=	1.1401389400021165	T22b=	0.87750170351178070

TTb=	0.96146344629156943	TSb=	0.78278795258102229

FATb=	0.18583701169265543	T11/T22=	0.784128606

Alleles sampled from the same population

T11w=	0.62283576809323782	T12w=	1.1404180998797744	T22w=	0.79429649439420680

TTw=	0.92449211556174826	TSw=	0.70856613124372236

FATw=	0.23356173696172955	T11/T22=	0.784135103

Time (units of 2NT generations) =	0.500000000

Alleles sampled from different populations

T11b=	0.71876341032787061	T12b=	1.1544277525496232	T22b=	0.87532195812940516

TTb=	0.97573521838913047	TSb=	0.79704268422863789

FATb=	0.18313629639760387	T11/T22=	0.821141779

Alleles sampled from the same population

T11w=	0.65061481593503112	T12w=	1.1547062047627463	T22w=	0.79232413507233579

TTw=	0.93808784013321489	TSw=	0.72146947550368346

FATw=	0.23091479855316122	T11/T22=	0.821147263

Time (units of 2NT generations) =	0.600000024

Alleles sampled from different populations

T11b=	0.76875332383894435	T12b=	1.1827224140244095	T22b=	0.87569433569142585

TTb=	1.0024731218947973	TSb=	0.82222382976518515

FATb=	0.17980461340341858	T11/T22=	0.877878606

Alleles sampled from the same population

T11w=	0.69586456935521290	T12w=	1.1829996162915020	T22w=	0.79266246503242943

TTw=	0.96363156674266159	TSw=	0.74426351719382122

FATw=	0.22764722236151358	T11/T22=	0.877882600

Time (units of 2NT generations) =	0.699999988

Alleles sampled from different populations

T11b=	0.80753543104290171	T12b=	1.2101210473723238	T22b=	0.88058401367622074

TTb=	1.0270903848659425	TSb=	0.84405972235956117

FATb=	0.17820307268310254	T11/T22=	0.917045295

Alleles sampled from the same population

T11w=	0.73096951171590707	T12w=	1.2103971658875943	T22w=	0.79708959940371260

TTw=	0.98721336072370203	TSw=	0.76402955555980978

FATw=	0.22607453874032013	T11/T22=	0.917048097

Time (units of 2NT generations) =	0.800000012

Alleles sampled from different populations

T11b=	0.83847786792073331	T12b=	1.2362680216252184	T22b=	0.88837542669918101

TTb=	1.0498473344675878	TSb=	0.86342664730995722

FATb=	0.17756932940366099	T11/T22=	0.943832815

Alleles sampled from the same population

T11w=	0.75897830278552592	T12w=	1.2365431790192147	T22w=	0.80414318953493036

TTw=	1.0090519625897214	TSw=	0.78156074616022808

FATw=	0.22545044741366882	T11/T22=	0.943834782

Time (units of 2NT generations) =	0.899999976

Alleles sampled from different populations

T11b=	0.86384043843932390	T12b=	1.2609987433024936	T22b=	0.89792414386242070

TTb=	1.0709405172266828	TSb=	0.88088229115087224

FATb=	0.17746851764277916	T11/T22=	0.962041676

Alleles sampled from the same population

T11w=	0.78193640174228007	T12w=	1.2612730344839846	T22w=	0.81278734381480300

TTw=	1.0293174536312630	TSw=	0.79736187277854154

FATw=	0.22534892421615926	T11/T22=	0.962043047

Time (units of 2NT generations) =	1.00000000

Alleles sampled from different populations

T11b=	0.88515005108665057	T12b=	1.2842599630063116	T22b=	0.90843173066793981

TTb=	1.0905254269418034	TSb=	0.89679089087729524

FATb=	0.17765247033973741	T11/T22=	0.974371552

Alleles sampled from the same population

T11w=	0.80122589080209916	T12w=	1.2845334647810698	T22w=	0.82229935506301965

TTw=	1.0481480438568147	TSw=	0.81176262293255941

FATw=	0.22552674911689041	T11/T22=	0.974372506

Time (units of 2NT generations) =	1.25000000

Alleles sampled from different populations

T11b=	0.92674766776547424	T12b=	1.3361353942417780	T22b=	0.93572573947112736

TTb=	1.1336860489300393	TSb=	0.93123670361830080

FATb=	0.17857619885400200	T11/T22=	0.990405202

Alleles sampled from the same population

T11w=	0.83888039826428540	T12w=	1.3364071868424996	T22w=	0.84700691991666344

TTw=	1.0896754229664869	TSw=	0.84294365909047442

FATw=	0.22642684112698463	T11/T22=	0.990405619

Time (units of 2NT generations) =	1.50000000

Alleles sampled from different populations

T11b=	0.95796400749372845	T12b=	1.3798195638300950	T22b=	0.96142620493831177

TTb=	1.1697573350230575	TSb=	0.95969510621602017

FATb=	0.17957761196932065	T11/T22=	0.996398866

Alleles sampled from the same population

T11w=	0.86713790153481707	T12w=	1.3800899443882957	T22w=	0.87027171728285768

TTw=	1.1243973768985664	TSw=	0.86870480940883743

FATw=	0.22740409462267486	T11/T22=	0.996399045

Time (units of 2NT generations) =	1.75000000

Alleles sampled from different populations

T11b=	0.98274925612165132	T12b=	1.4164665901435221	T22b=	0.98408437701823948

TTb=	1.1999417033567337	TSb=	0.98341681656994540

FATb=	0.18044617182741340	T11/T22=	0.998643279

Alleles sampled from the same population

T11w=	0.88957397201349098	T12w=	1.4167357936949077	T22w=	0.89078245967435843

TTw=	1.1534570047694161	TSw=	0.89017821584392465

FATw=	0.22825193122661969	T11/T22=	0.998643339

Time (units of 2NT generations) =	2.00000000

Alleles sampled from different populations

T11b=	1.0029922339922224	T12b=	1.4471714074902371	T22b=	1.0035070942325852

TTb=	1.2252105358013203	TSb=	1.0032496641124038

FATb=	0.18116141283730325	T11/T22=	0.999486923

Alleles sampled from the same population

T11w=	0.90789832886414601	T12w=	1.4474396269928675	T22w=	0.90836435576039198

TTw=	1.1777854846525684	TSw=	0.90813134231226900

FATw=	0.22895013213704518	T11/T22=	0.999486983

Time (units of 2NT generations) =	2.50000000

Alleles sampled from different populations

T11b=	1.0337053314692866	T12b=	1.4944203652427031	T22b=	1.0337818958684388

TTb=	1.2640819894557827	TSb=	1.0337436136688627

FATb=	0.18221790810110827	T11/T22=	0.999925911

Alleles sampled from the same population

T11w=	0.93570048154195795	T12w=	1.4946870717253369	T22w=	0.93576978397881871

TTw=	1.2152111022428627	TSw=	0.93573513276038833

FATw=	0.22998141554718976	T11/T22=	0.999925911

Time (units of 2NT generations) =	3.00000000

Alleles sampled from different populations

T11b=	1.0551143400098140	T12b=	1.5275494587424150	T22b=	1.0551257258321360

TTb=	1.2913347458316951	TSb=	1.0551200329209749

FATb=	0.18292291264770710	T11/T22=	0.999989212

Alleles sampled from the same population

T11w=	0.95508038263319794	T12w=	1.5278151045904258	T22w=	0.95509068853569734

TTw=	1.2414503200874367	TSw=	0.95508553558444764

FATw=	0.23066954824484642	T11/T22=	0.999989212

Time (units of 2NT generations) =	4.00000000

Alleles sampled from different populations

T11b=	1.0806145996447023	T12b=	1.5670620536113411	T22b=	1.0806148514352649

TTb=	1.3238383895756625	TSb=	1.0806147255399836

FATb=	0.18372609976482157	T11/T22=	0.999999762

Alleles sampled from the same population

T11w=	0.97816377573664282	T12w=	1.5673264344882496	T22w=	0.97816400364543288

TTw=	1.2727451620896437	TSw=	0.97816388969103785

FATw=	0.23145346073439443	T11/T22=	0.999999762

Time (units of 2NT generations) =	5.00000000

Alleles sampled from different populations

T11b=	1.0931469251010963	T12b=	1.5864849506193919	T22b=	1.0931469306692914

TTb=	1.3398159392522930	TSb=	1.0931469278851940

FATb=	0.18410664042760749	T11/T22=	1.00000000

Alleles sampled from the same population

T11w=	0.98950831119773486	T12w=	1.5867487096855573	T22w=	0.98950831623779922

TTw=	1.2881285117016621	TSw=	0.98950831371776704

FATw=	0.23182484920654967	T11/T22=	1.00000000

**Scaled recombination rate= 0.4**

Equilibrium results for subdivided population

Alleles sampled from different populations

T11b=	1.1052631578947369	T12b=	6.1052631578947372	T22b=	1.1052631578947369

TTb=	3.6052631578947367	TSb=	1.1052631578947369

FATb=	0.69343065693430650	T11/T22=	1.00000000

Alleles sampled from the same population

T11w=	1.0004761904761905	T12w=	6.1055263157894739	T22w=	1.0004761904761905

TTw=	3.5530012531328321	TSw=	1.0004761904761905

FATw=	0.71841378057662431	T11/T22=	1.00000000

Time (units of 2NT generations) =	2.49999994E-03

Alleles sampled from different populations

T11b=	2.9621816027880499E-003	T12b=	1.0542973071743118	T22b=	1.0503715762159567

TTb=	0.79048209304184214	TSb=	0.52666687890937236

FATb=	0.33373964629266484	T11/T22=	2.82012741E-03

Alleles sampled from the same population

T11w=	2.7032803389637206E-003	T12w=	1.0545617100166762	T22w=	0.95076866561592654

TTw=	0.76564884149706069	TSw=	0.47673597297744513

FATw=	0.37734383291785423	T11/T22=	2.84325774E-03

Time (units of 2NT generations) =	4.99999989E-03

Alleles sampled from different populations

T11b=	5.9710119171541010E-003	T12b=	1.0565331667619544	T22b=	1.0481270834587249

TTb=	0.79179110722494694	TSb=	0.52704904768793948

FATb=	0.33435846540998670	T11/T22=	5.69683965E-03

Alleles sampled from the same population

T11w=	5.4267406509178606E-003	T12w=	1.0567991954804992	T22w=	0.94873663594111701

TTw=	0.76694044188825827	TSw=	0.47708168829601744

FATw=	0.37794167286131031	T11/T22=	5.71996532E-03

Time (units of 2NT generations) =	7.49999983E-03

Alleles sampled from different populations

T11b=	8.9658667585945651E-003	T12b=	1.0587681075783868	T22b=	1.0458949615360937

TTb=	0.79309926086286553	TSb=	0.52743041414734415

FATb=	0.33497553184765616	T11/T22=	8.57243501E-03

Alleles sampled from the same population

T11w=	8.1375502508707065E-003	T12w=	1.0590341344436141	T22w=	0.94671623584813303

TTw=	0.76823051374655793	TSw=	0.47742689304950187

FATw=	0.37853693063927585	T11/T22=	8.59555416E-03

Time (units of 2NT generations) =	9.99999978E-03

Alleles sampled from different populations

T11b=	1.1946815755699003E-002	T12b=	1.0610021215489480	T22b=	1.0436751498866816

TTb=	0.79440655218506917	TSb=	0.52781098282119032

FATb=	0.33559084908173231	T11/T22=	1.14468718E-02

Alleles sampled from the same population

T11w=	1.0835772940586130E-002	T12w=	1.0612681465240961	T22w=	0.94470697842536988

TTw=	0.76951976110353715	TSw=	0.47777137568297801

FATw=	0.37913046573641540	T11/T22=	1.14699826E-02

Time (units of 2NT generations) =	1.49999997E-02

Alleles sampled from different populations

T11b=	1.7867273000917727E-002	T12b=	1.0654673692161127	T22b=	1.0392722088989867

TTb=	0.79701855508303243	TSb=	0.52856974094995224

FATb=	0.33681626659885422	T11/T22=	1.71921011E-02

Alleles sampled from the same population

T11w=	1.6194708121018975E-002	T12w=	1.0657333904229063	T22w=	0.94072166664316870

TTw=	0.77209578890250008	TSw=	0.47845818738209384

FATw=	0.38031239872166522	T11/T22=	1.72151960E-02

Time (units of 2NT generations) =	1.99999996E-02

Alleles sampled from different populations

T11b=	2.3732933756757113E-002	T12b=	1.0699289103084082	T22b=	1.0349177665893932

TTb=	0.79962713024074161	TSb=	0.52932535017307514

FATb=	0.33803477876780819	T11/T22=	2.29321923E-02

Alleles sampled from the same population

T11w=	2.1504044120062016E-002	T12w=	1.0701949277624880	T22w=	0.93678025353107741

TTw=	0.77466853829402882	TSw=	0.47914214882556971

FATw=	0.38148753287343495	T11/T22=	2.29552705E-02

Time (units of 2NT generations) =	2.99999993E-02

Alleles sampled from different populations

T11b=	3.5302038752325071E-002	T12b=	1.0788408750863061	T22b=	1.0263524270427806

TTb=	0.80483405399192942	TSb=	0.53082723289755285

FATb=	0.34045132625206265	T11/T22=	3.43956314E-02

Alleles sampled from the same population

T11w=	3.1975885426361872E-002	T12w=	1.0791068850813925	T22w=	0.92902735739986897

TTw=	0.77980425324725389	TSw=	0.48050162141311542

FATw=	0.38381764473300206	T11/T22=	3.44186686E-02

Time (units of 2NT generations) =	3.99999991E-02

Alleles sampled from different populations

T11b=	4.6658422486818285E-002	T12b=	1.0877380207871363	T22b=	1.0179752779614442

TTb=	0.81002743550563383	TSb=	0.53231685022413122

FATb=	0.34284096205723036	T11/T22=	4.58345339E-02

Alleles sampled from the same population

T11w=	4.2255181541540621E-002	T12w=	1.0880040233841495	T22w=	0.92144480222188252

TTw=	0.78492700763293055	TSw=	0.48184999188171157

FATw=	0.38612127344833613	T11/T22=	4.58575301E-02

Time (units of 2NT generations) =	5.00000007E-02

Alleles sampled from different populations

T11b=	5.7806291258628573E-002	T12b=	1.0966203527391256	T22b=	1.0097825427957192

TTb=	0.81520738488314981	TSb=	0.53379441702717390

FATb=	0.34520414445989478	T11/T22=	5.72462752E-02

Alleles sampled from the same population

T11w=	5.2345739806005653E-002	T12w=	1.0968863479979554	T22w=	0.91402916964478020

TTw=	0.79003690136167415	TSw=	0.48318745472539293

FATw=	0.38839887871997947	T11/T22=	5.72692230E-02

Time (units of 2NT generations) =	5.99999987E-02

Alleles sampled from different populations

T11b=	6.8749767622480196E-002	T12b=	1.1054878766846494	T22b=	1.0017705202045666

TTb=	0.82037401029908641	TSb=	0.53526014391352339

FATb=	0.34754131994213000	T11/T22=	6.86282590E-02

Alleles sampled from the same population

T11w=	6.2251291759396477E-002	T12w=	1.1057538646641110	T22w=	0.90677710939148048

TTw=	0.79513403261977467	TSw=	0.48451420057543848

FATw=	0.39065090827633031	T11/T22=	6.86511472E-02

Time (units of 2NT generations) =	7.00000003E-02

Alleles sampled from different populations

T11b=	7.9492892057628772E-002	T12b=	1.1143405987710748	T22b=	0.99393558255884140

TTb=	0.82552741803965501	TSb=	0.53671423730823509

FATb=	0.34985292362215215	T11/T22=	7.99779147E-02

Alleles sampled from the same population

T11w=	7.1975494650564631E-002	T12w=	1.1146065795289797	T22w=	0.89968533790539018

TTw=	0.80021849790347854	TSw=	0.48583041627797741

FATw=	0.39287779831280811	T11/T22=	8.00007433E-02

Time (units of 2NT generations) =	7.99999982E-02

Alleles sampled from different populations

T11b=	9.0039624602931134E-002	T12b=	1.1231785255415412	T22b=	0.98627417447436794

TTb=	0.83066771254009542	TSb=	0.53815689953864954

FATb=	0.35213937966479791	T11/T22=	9.12926942E-02

Alleles sampled from the same population

T11w=	8.1521932917550943E-002	T12w=	1.1234444991346884	T22w=	0.89275063702261281

TTw=	0.80529039205238506	TSw=	0.48713628497008188

FATw=	0.39507997391033933	T11/T22=	9.13154557E-02

Time (units of 2NT generations) =	9.00000036E-02

Alleles sampled from different populations

T11b=	0.10039384645937455	T12b=	1.1320016639262915	T22b=	0.97878281137418077

TTb=	0.83579499642153454	TSb=	0.53958832891677766

FATb=	0.35440110167322014	T11/T22=	0.102570094

Alleles sampled from the same population

T11w=	9.0894119638121884E-002	T12w=	1.1322676304104853	T22w=	0.88596985267055839

TTw=	0.81034980828241276	TSw=	0.48843198615434014

FATw=	0.39725784943467513	T11/T22=	0.102592789

Time (units of 2NT generations) =	0.100000001

Alleles sampled from different populations

T11b=	0.11055936156062929	T12b=	1.1408100212338796	T22b=	0.97145807807938778

TTb=	0.84090937052694414	TSb=	0.54100871982000853

FATb=	0.35663849306258422	T11/T22=	0.113807648

Alleles sampled from the same population

T11w=	0.10009549795137906	T12w=	1.1410759806639739	T22w=	0.87933989359245990

TTw=	0.81539683821794673	TSw=	0.48971769577191948

FATw=	0.39941182891731641	T11/T22=	0.113830268

Time (units of 2NT generations) =	0.125000000

Alleles sampled from different populations

T11b=	0.13517184132375570	T12b=	1.1627662985655425	T22b=	0.95385344821836093

TTb=	0.85363947166830045	TSb=	0.54451264477105832

FATb=	0.36212808469728286	T11/T22=	0.141711324

Alleles sampled from the same population

T11w=	0.12237363772741205	T12w=	1.1630322405941786	T22w=	0.86340512232151689

TTw=	0.82796081030932145	TSw=	0.49288938002446447

FATw=	0.40469479486556403	T11/T22=	0.141733736

Time (units of 2NT generations) =	0.150000006

Alleles sampled from different populations

T11b=	0.15868425609594894	T12b=	1.1846303728283010	T22b=	0.93721901356169379

TTb=	0.86629100382856117	TSb=	0.54795163482882137

FATb=	0.36747394073451456	T11/T22=	0.169313952

Alleles sampled from the same population

T11w=	0.14365605080575494	T12w=	1.1848962977781188	T22w=	0.84834852499153834

TTw=	0.84044929283838277	TSw=	0.49600228789864664

FATw=	0.40983674788572022	T11/T22=	0.169336125

Time (units of 2NT generations) =	0.174999997

Alleles sampled from different populations

T11b=	0.18115010183534341	T12b=	1.2064023870425684	T22b=	0.92150674778634711

TTb=	0.87886540592670681	TSb=	0.55132842481084521

FATb=	0.37268161757999230	T11/T22=	0.196580335

Alleles sampled from the same population

T11w=	0.16399115917301255	T12w=	1.2066682952226273	T22w=	0.83412663046962487

TTw=	0.85286359502197306	TSw=	0.49905889482131871

FATw=	0.41484324370949255	T11/T22=	0.196602240

Time (units of 2NT generations) =	0.200000003

Alleles sampled from different populations

T11b=	0.20262025187205546	T12b=	1.2280824968144950	T22b=	0.90667098043305150

TTb=	0.89136405648352413	TSb=	0.55464561615255348

FATb=	0.37775635878716241	T11/T22=	0.223477155

Alleles sampled from the same population

T11w=	0.18342501093823516	T12w=	1.2283483885208870	T22w=	0.82069810003828425

TTw=	0.86520497200457336	TSw=	0.50206155548825970

FATw=	0.41971952111527411	T11/T22=	0.223498762

Time (units of 2NT generations) =	0.224999994

Alleles sampled from different populations

T11b=	0.22314308557721096	T12b=	1.2496708696241638	T22b=	0.89266828143812860

TTb=	0.90378827656591687	TSb=	0.55790568350766978

FATb=	0.38270311977544280	T11/T22=	0.249973133

Alleles sampled from the same population

T11w=	0.20200139679798390	T12w=	1.2499367451407419	T22w=	0.80802362287915286

TTw=	0.87747462748965521	TSw=	0.50501250983856838

FATw=	0.42447052710419009	T11/T22=	0.249994427

Time (units of 2NT generations)=	0.224999994

Alleles sampled from different populations

T11b=	0.22314308557721096	T12b=	1.2496708696241638	T22b=	0.89266828143812860

TTb=	0.90378827656591687	TSb=	0.55790568350766978

FATb=	0.38270311977544280	T11/T22=	0.249973133

Alleles sampled from the same population

T11w=	0.20200139679798390	T12w=	1.2499367451407419	T22w=	0.80802362287915286

TTw=	0.87747462748965521	TSw=	0.50501250983856838

FATw=	0.42447052710419009	T11/T22=	0.249994427

Time (units of 2NT generations) =	0.250000000

Alleles sampled from different populations

T11b=	0.24276461071905397	T12b=	1.2711676841503001	T22b=	0.87945735132426739

TTb=	0.91613933258598046	TSb=	0.56111098102166068

FATb=	0.38752659004627998	T11/T22=	0.276039094

Alleles sampled from the same population

T11w=	0.21976196078722365	T12w=	1.2714335437491666	T22w=	0.79606581667892162

TTw=	0.88967371624111968	TSw=	0.50791388873307264

FATw=	0.42910093952307160	T11/T22=	0.276060045

Time (units of 2NT generations) =	0.275000006

Alleles sampled from different populations

T11b=	0.26152857981564637	T12b=	1.2925731296286687	T22b=	0.86699891677302010

TTb=	0.92841843896150089	TSb=	0.56426374829433323

FATb=	0.39223121319574328	T11/T22=	0.301648110

Alleles sampled from the same population

T11w=	0.23674630559622600	T12w=	1.2928389735708503	T22w=	0.78478913310655229

TTw=	0.90180334646111970	TSw=	0.51076771935138909

FATw=	0.43361518743996996	T11/T22=	0.301668674

Time (units of 2NT generations) =	0.300000012

Alleles sampled from different populations

T11b=	0.27947660077867975	T12b=	1.3138874052429790	T22b=	0.85525563131517446

TTb=	0.94062676064495310	TSb=	0.56736611604692710

FATb=	0.39682120498261708	T11/T22=	0.326775521

Alleles sampled from the same population

T11w=	0.25299209272006573	T12w=	1.3141532337788888	T22w=	0.77415976792293772

TTw=	0.91386458205019527	TSw=	0.51357593032150173

FATw=	0.43801746953653919	T11/T22=	0.326795727

Time (units of 2NT generations) =	0.324999988

Alleles sampled from different populations

T11b=	0.29664824212881524	T12b=	1.3351107195454466	T22b=	0.84419198088820713

TTb=	0.95276541552697880	TSb=	0.57042011150851124

FATb=	0.40130056967589511	T11/T22=	0.351399034

Alleles sampled from the same population

T11w=	0.26853513769450554	T12w=	1.3353765329153902	T22w=	0.76414557549601769

TTw=	0.92585844475532586	TSw=	0.51634035659526156

FATw=	0.44231177074621442	T11/T22=	0.351418823

Time (units of 2NT generations) =	0.349999994

Alleles sampled from different populations

T11b=	0.31308113304842955	T12b=	1.3562432899073036	T22b=	0.83377419402229491

TTb=	0.96483547672133296	TSb=	0.57342766353536223

FATb=	0.40567311487761393	T11/T22=	0.375498712

Alleles sampled from the same population

T11w=	0.28340950065898762	T12w=	1.3565090883420385	T22w=	0.75471598750545033

TTw=	0.93778591621212881	TSw=	0.51906274408221897

FATw=	0.44650187733806179	T11/T22=	0.375518084

Time (units of 2NT generations) =	0.375000000

Alleles sampled from different populations

T11b=	0.32881105852547998	T12b=	1.3772853419964877	T22b=	0.82397015642809390

TTb=	0.97683797473663736	TSb=	0.57639060747678694

FATb=	0.40994246499048514	T11/T22=	0.399057001

Alleles sampled from the same population

T11w=	0.29764757247627505	T12w=	1.3775511257176500	T22w=	0.74584193563153489

TTw=	0.94964793988577745	TSw=	0.52174475405390497

FATw=	0.45059139061928588	T11/T22=	0.399075955

Time (units of 2NT generations)=	0.400000006

Alleles sampled from different populations

T11b=	0.34387204982905151	T12b=	1.3982371092817720	T22b=	0.81474932977045122

TTb=	0.98877389954076167	TSb=	0.57931068979975131

FATb=	0.41411207347927215	T11/T22=	0.422058702

Alleles sampled from the same population

T11w=	0.31128015662662323	T12w=	1.3985028785023204	T22w=	0.73749577803306710

TTw=	0.96144542291608270	TSw=	0.52438796732984516

FATw=	0.45458373940835228	T11/T22=	0.422077209

Time (units of 2NT generations) =	0.425000012

Alleles sampled from different populations

T11b=	0.35829647054596014	T12b=	1.4190988325621374	T22b=	0.80608267442301607

TTb=	1.0006442025233127	TSb=	0.58218957248448810

FATb=	0.41818523405583374	T11/T22=	0.444490969

Alleles sampled from the same population

T11w=	0.32433654708398985	T12w=	1.4193645874867347	T22w=	0.72965122942849403

TTw=	0.97317923787148830	TSw=	0.52699388825624194

FATw=	0.45848219141124613	T11/T22=	0.444509000

Time (units of 2NT generations) =	0.449999988

Alleles sampled from different populations

T11b=	0.37211509839579326	T12b=	1.4398707595186151	T22b=	0.79794257600862295

TTb=	1.0124497983604117	TSb=	0.58502883720220811

FATb=	0.42216509090167287	T11/T22=	0.466343194

Alleles sampled from the same population

T11w=	0.33684460237109914	T12w=	1.4401365003441260	T22w=	0.72228329460377205

TTw=	0.98485022441578074	TSw=	0.52956394848743560

FATw=	0.46228986361700208	T11/T22=	0.466360778

Time (units of 2NT generations) =	0.474999994

Alleles sampled from different populations

T11b=	0.38535720303192855	T12b=	1.4605531442905813	T22b=	0.79030277553976858

TTb=	1.0241915667882149	TSb=	0.58782998928584851

FATb=	0.42605464802914006	T11/T22=	0.487607062

Alleles sampled from the same population

T11w=	0.34883081598120813	T12w=	1.4608188712063654	T22w=	0.71536820517887056

TTw=	0.99645919089320245	TSw=	0.53209951058003935

FATw=	0.46600973181543148	T11/T22=	0.487624168

Time (units of 2NT generations) =	0.500000000

Alleles sampled from different populations

T11b=	0.39805062002520208	T12b=	1.4811462470719468	T22b=	0.78313830298301979

TTb=	1.0358703542880290	TSb=	0.59059446150411099

FATb=	0.42985677786865861	T11/T22=	0.508276284

Alleles sampled from the same population

T11w=	0.36032038334458338	T12w=	1.4814119602602140	T22w=	0.70888335947343895

TTw=	1.0080069158346125	TSw=	0.53460187140901116

FATw=	0.46964463932633838	T11/T22=	0.508292913

Time (units of 2NT generations) =	0.574999988

Alleles sampled from different populations

T11b=	0.43309704305239283	T12b=	1.5423925483248953	T22b=	0.76426518151192346

TTb=	1.0705368303035268	TSb=	0.59868111228215815

FATb=	0.44076551564095601	T11/T22=	0.566684246

Alleles sampled from the same population

T11w=	0.39204300689146154	T12w=	1.5426582213578435	T22w=	0.69180058691551860

TTw=	1.0422900091306668	TSw=	0.54192179690349007

FATw=	0.48006620791128396	T11/T22=	0.566699445

Time (units of 2NT generations) =	0.600000024

Alleles sampled from different populations

T11b=	0.44384777460014124	T12b=	1.5626312331730370	T22b=	0.75877595203161463

TTb=	1.0819715482444574	TSb=	0.60131186331587794

FATb=	0.44424429247559161	T11/T22=	0.584952354

Alleles sampled from the same population

T11w=	0.40177414659168131	T12w=	1.5628968931427991	T22w=	0.68683208962740872

TTw=	1.0536000056261721	TSw=	0.54430311810954501

FATw=	0.48338732421887509	T11/T22=	0.584967077

Time (units of 2NT generations) =	0.625000000

Alleles sampled from different populations

T11b=	0.45416983018533030	T12b=	1.5827820150741951	T22b=	0.75365442855777964

TTb=	1.0933470722228751	TSb=	0.60391212937155503

FATb=	0.44764828597039530	T11/T22=	0.602623463

Alleles sampled from the same population

T11w=	0.41111726889177969	T12w=	1.5830476621322500	T22w=	0.68219642185676710

TTw=	1.0648522537532616	TSw=	0.54665684537427339

FATw=	0.48663596902999096	T11/T22=	0.602637708

Time (units of 2NT generations) =	0.649999976

Alleles sampled from different populations

T11b=	0.46408380325292709	T12b=	1.6028451831530672	T22b=	0.74888215122489898

TTb=	1.1046640801959902	TSb=	0.60648297723891309

FATb=	0.45097972486684779	T11/T22=	0.619702041

Alleles sampled from the same population

T11w=	0.42009101399575588	T12w=	1.6031108174455291	T22w=	0.67787687461640100

TTw=	1.0760473808758038	TSw=	0.54898394430607844

FATw=	0.48981433897524485	T11/T22=	0.619715810

Time (units of 2NT generations) =	0.675000012

Alleles sampled from different populations

T11b=	0.47360927879784076	T12b=	1.6228210302823838	T22b=	0.74444156682987606

TTb=	1.1159232265481211	TSb=	0.60902542281385841

FATb=	0.45424075032674671	T11/T22=	0.636194050

Alleles sampled from the same population

T11w=	0.42871310930679751	T12w=	1.6230866519503842	T22w=	0.67385755958664184

TTw=	1.0871859931985519	TSw=	0.55128533444671968

FATw=	0.49292454290657983	T11/T22=	0.636207342

Time (units of 2NT generations) =	0.699999988

Alleles sampled from different populations

T11b=	0.48276488283533481	T12b=	1.6427098528169966	T22b=	0.74031598438839397

TTb=	1.1271251432144305	TSb=	0.61154043361186439

FATb=	0.45743342050925961	T11/T22=	0.652106524

Alleles sampled from the same population

T11w=	0.43700041420554725	T12w=	1.6429754619967820	T22w=	0.67012336888710000

TTw=	1.0982686767715528	TSw=	0.55356189154632363

FATw=	0.49596860654028452	T11/T22=	0.652119339

Time (units of 2NT generations) =	0.725000024

Alleles sampled from different populations

T11b=	0.49156832944429851	T12b=	1.6625119503419912	T22b=	0.73648953286943108

TTb=	1.1382704407494280	TSb=	0.61402893115686474

FATb=	0.46055971483139535	T11/T22=	0.667447805

Alleles sampled from the same population

T11w=	0.44496896263141905	T12w=	1.6627775471652875	T22w=	0.66665993681995439

TTw=	1.1092959984454871	TSw=	0.55581444972568672

FATw=	0.49894847677754384	T11/T22=	0.667460203

Time (units of 2NT generations) =	0.750000000

Alleles sampled from different populations

T11b=	0.50003646550219494	T12b=	1.6822276254321595	T22b=	0.73294712100122517

TTb=	1.1493597093419348	TSb=	0.61649179325171000

FATb=	0.46362153793899563	T11/T22=	0.682227194

Alleles sampled from the same population

T11w=	0.45263400357451800	T12w=	1.6824932100263315	T22w=	0.66345360348822813

TTw=	1.1202685067788523	TSw=	0.55804380353137306

FATw=	0.50186602572990635	T11/T22=	0.682239115

Time (units of 2NT generations) =	0.774999976

Alleles sampled from different populations

T11b=	0.50818531322540217	T12b=	1.7018571834245284	T22b=	0.72967439904712450

TTb=	1.1603935197803958	TSb=	0.61892985613626328

FATb=	0.46662072341338512	T11/T22=	0.696454883

Alleles sampled from the same population

T11w=	0.46001003958116127	T12w=	1.7021227559127663	T22w=	0.66049138019712461

TTw=	1.1311867329009546	TSw=	0.56025070988914294

FATw=	0.50472305447539423	T11/T22=	0.696466386

Time (units of 2NT generations) =	0.800000012

Alleles sampled from different populations

T11b=	0.51603011062218429	T12b=	1.7214009322032515	T22b=	0.72665772245483662

TTb=	1.1713724243708810	TSb=	0.62134391653851040

FATb=	0.46955903723598336	T11/T22=	0.710141897

Alleles sampled from the same population

T11w=	0.46711086336993057	T12w=	1.7216664927047933	T22w=	0.65776091655105229

TTw=	1.1420511913326423	TSw=	0.56243588996049143

FATw=	0.50752129656798184	T11/T22=	0.710152984

Time (units of 2NT generations) =	0.824999988

Alleles sampled from different populations

T11b=	0.52358534996084560	T12b=	1.7408591819943782	T22b=	0.72388411728719748

TTb=	1.1822969578092000	TSb=	0.62373473362402154

FATb=	0.47243818103042068	T11/T22=	0.723299980

Alleles sampled from the same population

T11w=	0.47394959265124381	T12w=	1.7411247306246693	T22w=	0.65525046916320884

TTw=	1.1528623807659479	TSw=	0.56460003090722632

FATw=	0.51026242132030286	T11/T22=	0.723310590

Time (units of 2NT generations) =	0.850000024

Alleles sampled from different populations

T11b=	0.53086481435073662	T12b=	1.7602322451714931	T22b=	0.72134124734722926

TTb=	1.1931676380102381	TSb=	0.62610303084898300

FATb=	0.47525979510046801	T11/T22=	0.735941291

Alleles sampled from the same population

T11w=	0.48053870323870762	T12w=	1.7604977820424086	T22w=	0.65294887189875905

TTw=	1.1636207848055709	TSw=	0.56674378756873334

FATw=	0.51294803687832857	T11/T22=	0.735951483

Time (units of 2NT generations) =	0.875000000

Alleles sampled from different populations

T11b=	0.53788161252819044	T12b=	1.7795204360713663	T22b=	0.71901738291439765

TTb=	1.2039849668963303	TSb=	0.62844949772129399

FATb=	0.47802546128019308	T11/T22=	0.748078704

Alleles sampled from the same population

T11w=	0.48689006053570405	T12w=	1.7797859612913438	T22w=	0.65084550757638460

TTw=	1.1743268726736942	TSw=	0.56886778405604432

FATw=	0.51557969310465279	T11/T22=	0.748088539

Time (units of 2NT generations) =	0.899999976

Alleles sampled from different populations

T11b=	0.54464821193590640	T12b=	1.7987240708198700	T22b=	0.71690137101311291

TTb=	1.2147494311471898	TSb=	0.63077479147450966

FATb=	0.48073670561111848	T11/T22=	0.759725451

Alleles sampled from the same population

T11w=	0.49301494947724844	T12w=	1.7989895844941053	T22w=	0.64893028105674322

TTw=	1.1849810998805506	TSw=	0.57097261526699583

FATw=	0.51815888428553714	T11/T22=	0.759734869

Time (units of 2NT generations) =	0.925000012

Alleles sampled from different populations

T11b=	0.55117647017926208	T12b=	1.8178434671649386	T22b=	0.71498260713837425

TTb=	1.2254615029118785	TSb=	0.63307953865881816

FATb=	0.48339500085924592	T11/T22=	0.770894945

Alleles sampled from the same population

T11w=	0.49892410300275580	T12w=	1.8181089693954835	T22w=	0.64719359364988560

TTw=	1.1955839088609019	TSw=	0.57305884832632070

FATw=	0.52068705167477103	T11/T22=	0.770903945

Time (units of 2NT generations)=	0.949999988

Alleles sampled from different populations

T11b=	0.55747766493927320	T12b=	1.8368789443202500	T22b=	0.71325100836717015

TTb=	1.2361216404867359	TSb=	0.63536433665322167

FATb=	0.48600176888494540	T11/T22=	0.781600952

Alleles sampled from the same population

T11w=	0.50462772913182075	T12w=	1.8371444352061985	T22w=	0.64562631877696774

TTw=	1.2061357295802964	TSw=	0.57512702395439419

FATw=	0.52316558588764850	T11/T22=	0.781609595

Time (units of 2NT generations) =	0.975000024

Alleles sampled from different populations

T11b=	0.56356252241808513	T12b=	1.8558308228158422	T22b=	0.71169698778770174

TTb=	1.2467302889593679	TSb=	0.63762975510289344

FATb=	0.48855838287596587	T11/T22=	0.791857421

Alleles sampled from the same population

T11w=	0.51013553671170719	T12w=	1.8560963024534463	T22w=	0.64421977882480985

TTw=	1.2166369801108523	TSw=	0.57717765776825858

FATw=	0.52559582915548908	T11/T22=	0.791865706

Time (units of 2NT generations) =	1.00000000

Alleles sampled from different populations

T11b=	0.56944124438851995	T12b=	1.8746994243565025	T22b=	0.71031143018188425

TTb=	1.2572878808208521	TSb=	0.63987633728520210

FATb=	0.49106616945401338	T11/T22=	0.801678300

Alleles sampled from the same population

T11w=	0.51545675990130768	T12w=	1.8749648928393405	T22w=	0.64296572313485334

TTw=	1.2270880671787106	TSw=	0.57921124151808057

FATw=	0.52797907745139416	T11/T22=	0.801686227

Time (units of 2NT generations) =	1.10000002

Alleles sampled from different populations

T11b=	0.59108186323924672	T12b=	1.9493475288757507	T22b=	0.70628677066600876

TTb=	1.2990159229141891	TSb=	0.64868431695262774

FATb=	0.50063405266243310	T11/T22=	0.836886525

Alleles sampled from the same population

T11w=	0.53504515014438980	T12w=	1.9496129536253672	T22w=	0.63932312935391966

TTw=	1.2683985466872609	TSw=	0.58718413974915473

FATw=	0.53706653063996912	T11/T22=	0.836893141

Time (units of 2NT generations) =	1.20000005

Alleles sampled from different populations

T11b=	0.61012054667459736	T12b=	2.0226891586316205	T22b=	0.70433615816444650

TTb=	1.3399587555255712	TSb=	0.65722835241952193

FATb=	0.50951598345149240	T11/T22=	0.866234899

Alleles sampled from the same population

T11w=	0.55227838685966313	T12w=	2.0229545409590024	T22w=	0.63755785798286491

TTw=	1.3089363316901332	TSw=	0.59491812242126407

FATw=	0.54549498855067302	T11/T22=	0.866240442

Time (units of 2NT generations) =	1.29999995

Alleles sampled from different populations

T11b=	0.62700785857583208	T12b=	2.0947452113210359	T22b=	0.70405823469844153

TTb=	1.3801391289790863	TSb=	0.66553304663713675

FATb=	0.51777829302655631	T11/T22=	0.890562475

Alleles sampled from the same population

T11w=	0.56756429904434191	T12w=	2.0950105524149389	T22w=	0.63730661904496833

TTw=	1.3487230057297972	TSw=	0.60243545904465512

FATw=	0.55332899603156405	T11/T22=	0.890567064

Time (units of 2NT generations) =	1.39999998

Alleles sampled from different populations

T11b=	0.64211265334709167	T12b=	2.1655366156955109	T22b=	0.70512514529782033

TTb=	1.4195777575089836	TSb=	0.67361889932245600

FATb=	0.52547939289743839	T11/T22=	0.910636425

Alleles sampled from the same population

T11w=	0.58123675618011572	T12w=	2.1658019166428124	T22w=	0.63827265397601463

TTw=	1.3877783108604387	TSw=	0.60975470507806517

FATw=	0.56062528120935240	T11/T22=	0.910640240

Time (units of 2NT generations) =	1.50000000

Alleles sampled from different populations

T11b=	0.65573696248980184	T12b=	2.2350842574056591	T22b=	0.70726914625905091

TTb=	1.4582936558900428	TSb=	0.68150305437442638

FATb=	0.53267090505273884	T11/T22=	0.927139223

Alleles sampled from the same population

T11w=	0.59356914284887374	T12w=	2.2353495192080466	T22w=	0.64021361405881372

TTw=	1.4261204488309451	TSw=	0.61689137845384368

FATw=	0.56743388753766399	T11/T22=	0.927142322

Time (units of 2NT generations) =	1.75000000

Alleles sampled from different populations

T11b=	0.68483609602318307	T12b=	2.4036479124793582	T22b=	0.71600404875747103

TTb=	1.5520339924348425	TSb=	0.70042007239032711

FATb=	0.54870829131035792	T11/T22=	0.956469595

Alleles sampled from the same population

T11w=	0.61990904674405545	T12w=	2.4039130803593958	T22w=	0.64812078875943802

TTw=	1.5189639990555712	TSw=	0.63401491775174668

FATw=	0.58260043151387997	T11/T22=	0.956471503

Time (units of 2NT generations) =	2.00000000

Alleles sampled from different populations

T11b=	0.70890607369286873	T12b=	2.5648874729559763	T22b=	0.72775723051550512

TTb=	1.6416095625300815	TSb=	0.71833165210418692

FATb=	0.56242235151391307	T11/T22=	0.974096894

Alleles sampled from the same population

T11w=	0.64169676603543613	T12w=	2.5651525519064524	T22w=	0.65875993272801958

TTw=	1.6076904506440903	TSw=	0.65022834938172780

FATw=	0.59555127722427770	T11/T22=	0.974098027

Time (units of 2NT generations) =	2.25000000

Alleles sampled from different populations

T11b=	0.72965300416491685	T12b=	2.7191141523152780	T22b=	0.74105465497534284

TTb=	1.7272339909427039	TSb=	0.73535382957012985

FATb=	0.57425928772465717	T11/T22=	0.984614313

Alleles sampled from the same population

T11w=	0.66047659019163485	T12w=	2.7193791467524910	T22w=	0.67079681988254314

TTw=	1.6925079258947902	TSw=	0.66563670503708905

FATw=	0.60671575308270287	T11/T22=	0.984614968

Time (units of 2NT generations) =	2.50000000

Alleles sampled from different populations

T11b=	0.74812080185909069	T12b=	2.8666287442561851	T22b=	0.75501680524105397

TTb=	1.8090987739031286	TSb=	0.75156880355007227

FATb=	0.58456176390603409	T11/T22=	0.990866423

Alleles sampled from the same population

T11w=	0.67719342203370392	T12w=	2.8668936581881113	T22w=	0.68343535489364338

TTw=	1.7736040233258925	TSw=	0.68031438846367365

FATw=	0.61642261772278995	T11/T22=	0.990866840

Time (units of 2NT generations) =	2.75000000

Alleles sampled from different populations

T11b=	0.76495244072221968	T12b=	3.0077208258930783	T22b=	0.76912331592569838

TTb=	1.8873793521085187	TSb=	0.76703787832395909

FATb=	0.59359633903642672	T11/T22=	0.994577110

Alleles sampled from the same population

T11w=	0.69242925216788320	T12w=	3.0079856630231996	T22w=	0.69620452917186149

TTw=	1.8511512768465361	TSw=	0.69431689066987234

FATw=	0.62492698497734256	T11/T22=	0.994577348

Time (units of 2NT generations) =	3.00000000

Alleles sampled from different populations

T11b=	0.78054794798199767	T12b=	3.1426684940543366	T22b=	0.78307059755150854

TTb=	1.9622388834105446	TSb=	0.78180927276675316

FATb=	0.60157283632668634	T11/T22=	0.996778488

Alleles sampled from the same population

T11w=	0.70654616738344833	T12w=	3.1429332578466287	T22w=	0.70882954927435948

TTw=	1.9253105580877663	TSw=	0.70768785832890391

FATw=	0.63242924350252094	T11/T22=	0.996778667

Time (units of 2NT generations) =	3.50000000

Alleles sampled from different populations

T11b=	0.80895152019584915	T12b=	3.3951860396040185	T22b=	0.80987433889212879

TTb=	2.1022994845740040	TSb=	0.80941292954398891

FATb=	0.61498685820778620	T11/T22=	0.998860538

Alleles sampled from the same population

T11w=	0.73225686002192536	T12w=	3.3954506663483599	T22w=	0.73309215142551509

TTw=	2.0640625860360400	TSw=	0.73267450572372028

FATw=	0.64503280536139362	T11/T22=	0.998860598

Time (units of 2NT generations) =	4.00000000

Alleles sampled from different populations

T11b=	0.83447352891560633	T12b=	3.6261822097339476	T22b=	0.83481110824355564

TTb=	2.2304122641567643	TSb=	0.83464231857958104

FATb=	0.62579011423472064	T11/T22=	0.999595642

Alleles sampled from the same population

T11w=	0.75535920222605635	T12w=	3.6264467112182954	T22w=	0.75566476290856333

TTw=	2.1909793468928029	TSw=	0.75551198256730978

FATw=	0.65517156351165085	T11/T22=	0.999595642

Time (units of 2NT generations) =	4.50000000

Alleles sampled from different populations

T11b=	0.85765174344444661	T12b=	3.8374899198441454	T22b=	0.85777523444341486

TTb=	2.3476017043940378	TSb=	0.85771348894393074

FATb=	0.63464267071431379	T11/T22=	0.999856055

Alleles sampled from the same population

T11w=	0.77633996532589455	T12w=	3.8377543067837729	T22w=	0.77645174347936607

TTw=	2.3070750805932017	TSw=	0.77639585440263037

FATw=	0.66347178688133490	T11/T22=	0.999856055

Time (units of 2NT generations) =	5.00000000

Alleles sampled from different populations

T11b=	0.87879276045688737	T12b=	4.0307868057639258	T22b=	0.87883793510344121

TTb=	2.4548010767720450	TSb=	0.87881534778016435

FATb=	0.64200140040032583	T11/T22=	0.999948621

Alleles sampled from the same population

T11w=	0.79547667353201401	T12w=	4.0310510879361585	T22w=	0.79551756346421720

TTw=	2.4132741032171370	TSw=	0.79549711849811566

FATw=	0.67036603200704059	T11/T22=	0.999948621
